# Regio-selective sulfation of heparin mimicking polymers defines in vivo activity

**DOI:** 10.64898/2026.07.09.737190

**Authors:** Madeline K. Loffredo, Hyun Ok Ham, Maria Varghese, Carolyn A Haller, Elliot L. Chaikof, Mark W. Grinstaff

## Abstract

Heparin, a naturally derived glycosaminoglycan, is the most commonly used anti-thromboembolic in the world. However, the biological origin of heparin inherently results in batch-to-batch variability, large dispersity indexes, and potential contamination, leading to inconsistent activity and patient-dependent dose-response. As such, new synthetic anticoagulants are of keen interest, particularly those that mimic heparin while being amenable to alterations in polymer structure and composition for performance optimization. Herein, we report the strategy, synthesis, and evaluation of well-defined, regioselectively functionalized di-sulfated polyamidosaccharides (**disulPASs**) including exploration of the structure-function relationship of molecular weight and sulfation density on anticoagulant activity. Polymerization of an orthogonally protected ý-lactam monomer via anionic ring-opening, followed by selective deprotection and sulfation reactions affords disulPAS. Similar to heparin, **disulPAS**s elongate clotting time through the intrinsic and extrinsic pathways, showing molecular weight and dose-dependent responses in clotting time; are non-cytotoxic and non-hemolytic, partially neutralized by protamine sulfate, and unlike heparin, are not degraded by heparinases. As compared to less sulfated and randomly sulfated iterations of PAS, **disulPAS** performs superiorly, with *in vitro* and *in vivo* clotting activity most similar to native heparin.

## Introduction

Heparin is a heterogeneously sulfated glycosaminoglycan extracted from porcine and bovine mucosa. Discovered in the early 20^th^ century and clinically approved since the 1930s,^1^ this anticoagulant is still widely used due to its favorable properties such as the rapid onset of anticoagulation following intravenous administration, reversibility, and widespread availability. Approximately 175 metric tons of heparin are utilized every year worldwide for the treatment of thromboembolic disorders including venous thromboembolism, deep vein thrombosis, and pulmonary embolism.^2^ However, while heparin is widely used and is listed as an “essential medicine” by the World Health Organization,^3^ it possesses several drawbacks, primarily linked to its heterogeneous structure and animal origin.

Heparin is a polysaccharide (average molecular weight between 12-16 kDa),^4^ composed of repeating disaccharide subunits of 1→4 linked α-D-glucosamine and iduronic acid. It is variably sulfated, with a majority of the highly sulfated (>2 sulfate groups per monomer) sequences concentrated at the non-reducing end of the polysaccharide, and less sulfated monomer sequences (<2 sulfate groups per monomer) concentrated at the reducing end, with sequences of intermediate sulfation between.^5^ Heparin exerts its anticoagulant effect most significantly by binding to and enhancing the activity of antithrombin-III (AT-III), a serine protease inhibitor, via a unique pentasaccharide sequence known as the antithrombin binding domain.^6,7^ The proportions of variably charged domains and the number of antithrombin binding regions in each batch of heparin vary significantly depending on the source animal and organ, as well as on the extraction and purification processes. Due to this inconsistency in structure, heparin exhibits batch-to-batch variation in activity, bioavailability, and patient-dependent dose-response.^8,9^ Additionally, the animal origin of heparin can lead to harmful biological and chemical impurities.^10,11^ In 2008, chondroitin sulfate contamination of heparin led to the death of ∼100 individuals.^11^ Moreover, the ongoing epidemic of African Swine Fever, which has wiped out over one-quarter of the world’s pig population, is resulting in a shortage of heparin and thus further reiterating the dangers of relying on animal tissue for a pharmaceutical agent.^12,13^ While small molecule synthetic anticoagulants are known,^14–17^ the organic routes are often arduous and the resulting drugs are either not as efficient as heparin or lack key favorable qualities such as reversibility, leading to increased bleeding risks in patients.^18^ Thus, there is demand for synthetic anticoagulants with selective and consistent functionalization and which retain the favorable safety profile and efficacy of heparin.

Previously, we reported a heparin mimicking synthetic anticoagulant, sulfated polyamidosaccharides (sulPASs).^19^ SulPASs exhibit promising properties such as dose-dependent increases in clotting time *ex vivo* and *in vivo*, reversal of activity by protamine sulfate, and safety profiles similar to heparin *in vivo*. However, these sulPASs are randomly sulfated and thus suffer from the same heterogeneity as heparin. In this work, we report the strategy, synthesis, characterization, and activity of enantiopure 3,6-disulfated polyamidosaccharides (**disulPASs**), which overcome these limitations. We highlight the synthesis and selection of orthogonally protected β-lactam monomers as well as their polymerization efficacy, kinetics, and post-polymerization modifications. Expansion of the **disulPAS** polymer library and further evaluation illustrates molecular weight, sulfation density, and dose-dependent anticoagulant activity by *ex vivo* and *in vivo* clotting and pharmacokinetic assays, mechanism of action by amidolytic activity inhibition assays, and safety and biocompatibility studies.

## Results

### Monomer design and synthesis

Anticoagulant activity of unfractionated heparin (UFH) relies primarily on the serine protease inhibitor antithrombin-III (AT-III). Mechanistically, a sulfated pentasaccharide sequence in UFH binds to AT-III resulting in a conformational change that exposes a reactive loop, which then binds to and inactivates enzymatic factors IIa and Xa, and to a lesser extent factors VIIa, IXa, XIa, and XIIa in the coagulation cascade.^20^ Elucidation of heparin’s anticoagulant activity has led to the development of FXa inhibitors such as Fondaparinux^16^ (K_D_ for AT-III binding of 36 nM)^21^ (Figure 1A), which is structurally the isolated AT-III pentasaccharide binding domain of heparin, and idraparinux,^15^ the methylated counterpart to fondaparinux.

**Figure 1.**
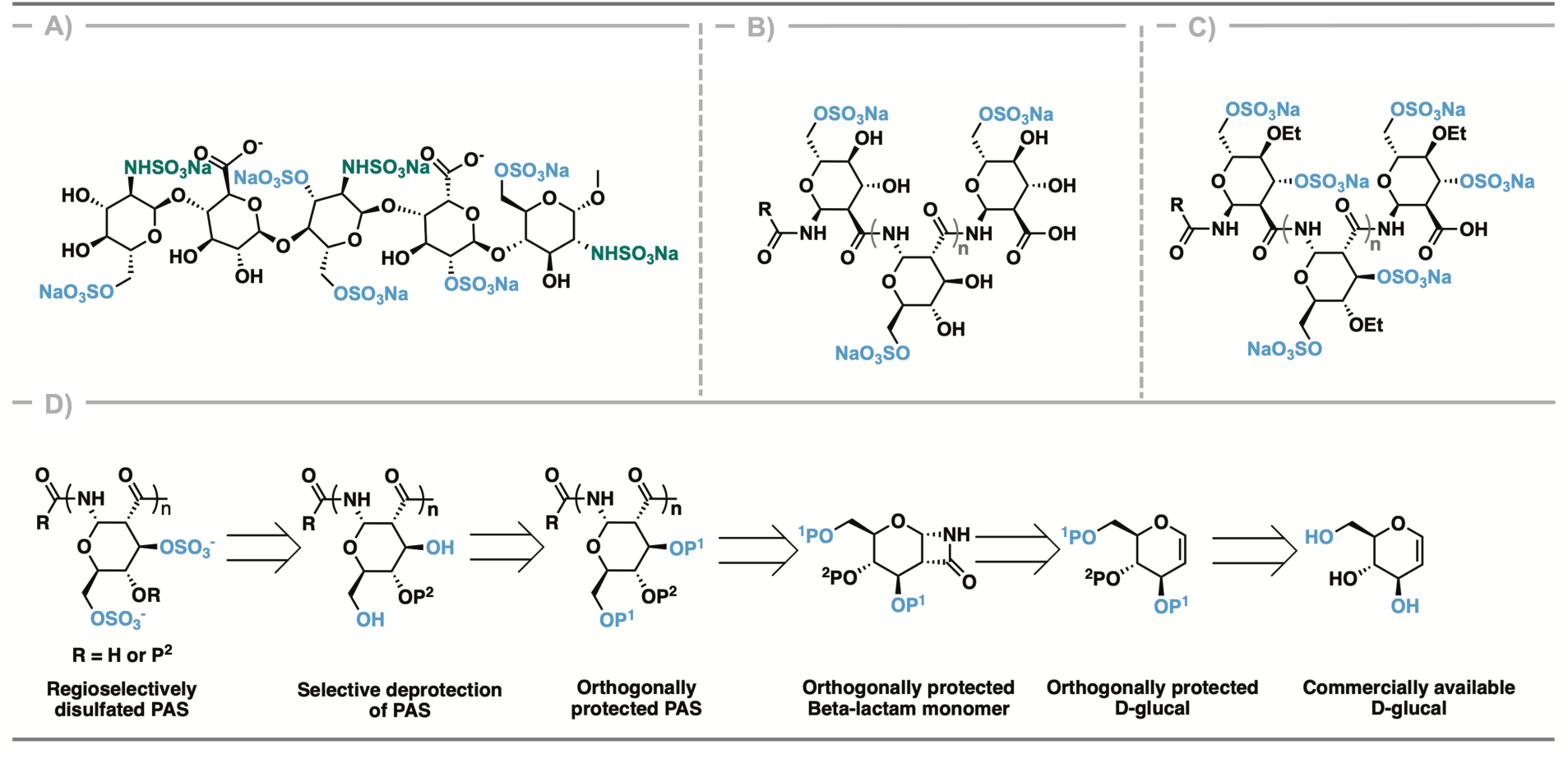
**A**) Structure of fondaparinux; **B**) Structure of 6-sulfate-PAS (6-sulPAS); **C**) Structure of 3,6-disulfate-4-*O*-ethyl-PAS (disulPAS); and **D**) Retrosynthetic scheme for disulPAS synthesis, where P^1^ and P^2^ are orthogonal protecting groups.

The AT-III binding domain of UFH comprises iduronic acid and glucosamine with an average degree of sulfation of 2 per monomer, and with sulfate groups typically at the O-6 and O-2 positions. Based on these observations and with the goal of achieving a defined sulfation pattern, we synthesized PASs that possess regioselective sulfation at the O-3 and O-6 positions, since O-2 is inaccessible for sulfation in PASs (Figure 1B, C). To prepare 3,6-disulfated PAS, we utilized a combination of pre-polymerization and post-polymerization modification reactions, starting from commercially available *D*-glucal. Beginning with orthogonally protected β-lactam monomers, we performed an anionic ring opening polymerization followed by selective deprotection of O-3 and O-6 and subsequent sulfation (Figure 1D).

Thus, we sought to determine the most effective orthogonal protecting groups for the selective protection and deprotection reactions, exploring the following combinations: tert-butyldimethylsilyl (TBS) & *para*-methoxybenzyl (PMB), triethylsilyl (TES) & PMB, trimethylsilyl (TMS) & PMB, PMB & benzyl (Bn), PMB & ethyl (Et), and PMB & triisopropylsilyl (TIPS) to synthesize monomers **1**, **2**, **3**, **4**, **5**, and **6** respectively (Figure 2A). The synthesis of monomers **1**, **2**, **3**, **4**, and **5** all begin with 3,6-di-*O*-TBS-4-OH *D*-glucal (Compound 1, Scheme S1). Taking advantage of the steric bulk of the TBS groups, we selectively protected the O-3 and O-6 positions^22^ leaving the O-4 hydroxyl available for subsequent protection with an orthogonal protecting group (PMB, Bn, Et). We then selectively removed the TBS groups using tetrabutylammonium fluoride (TBAF) and replaced them with TES, TMS, or PMB to obtain fully protected *D*-glucals (Schemes S1-5). Our efforts to synthesize monomer **3** by replacing the bulky TBS groups with TMS groups were unsuccessful due to the inherent instability of the trimethyl silyl groups. Initial attempts to prepare 3,6-di-*O*-TMS-4-*O*-PMB-*D*-glucal were hindered by the cleavage of the TMS groups in the presence of water, leading to their removal during the aqueous work up after the silylation reaction; therefore, we did not pursue this monomer further. For monomer **6**, because TIPS is also cleaved by TBAF, we instead began with 4,6-*para*-methoxybenzylidene-3-*O*-PMB *D*-glucal and selectively opened the benzylidene at O-6 to reveal a hydroxyl at O-4 via a Lewis acid mediated acetal hydrolysis. To favor formation of the thermodynamic product bearing the 4-hydroxy group, rather than the more kinetically favorable 6-hydroxy product, we cooled the solution to -78°C for the duration of the reaction.^23^ We confirmed the structure of the desired product via ^1^H NMR. The NMR spectrum shows the disappearance of the benzylidene proton at 5.57 ppm and a slight upfield shift of the H-6 protons from 4.33 and 3.86 to 4.06 and 3.76 ppm in the product. H-4 remains at approximately 4.0 ppm, however a hydroxyl proton emerges at 2.59 which couples to H-4 as seen in the [^1^H, ^1^H] COSY NMR (Small Molecule NMRs in Supplemental Information). We then functionalized the available 4-hydroxyl group with TIPS, yielding our final fully protected glucal (Scheme S6).

**Figure 2.**
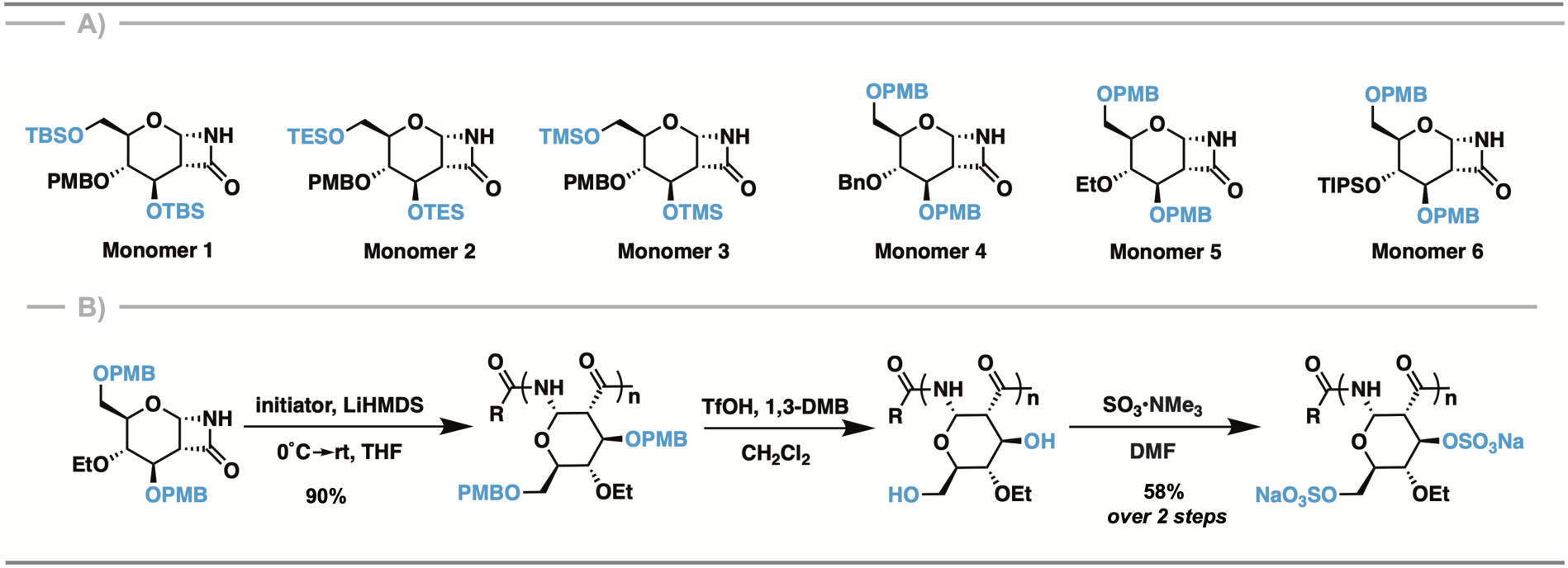
**A)** Monomers **1** – **6**; **B)** Synthetic scheme of 3,6-disulfate-4-*O*-ethyl-PAS.

The general synthetic procedure to obtain β-lactam monomers **1**, **2**, **4**, and **5** from the corresponding *D*-glucals involves a (2+2) cycloaddition using chlorosulfonyl isocyanate (CSI), followed by *in situ* reductive removal of the sulfonyl group via Red-Al (see Supplemental Information for detailed small molecule synthetic procedures and characterization). Monomer **6** requires a modified cycloaddition using trichloroacetyl isocyanate (TCAI). We synthesized all monomers with >95% purity, with isolated yields ranging from 18 - 70%.

### Polymerization and post-polymerization modifications

Once we established the library of orthogonally protected monomers, we determined the polymerization efficiency, as well as the post-polymerization deprotection efficiency and selectivity at the O-3 and O-6 positions. While monomer syntheses were straightforward and resulted in moderate to high-yields, polymerization and post-polymerization reactions were more challenging. We first conducted trial polymerizations of all monomers by dissolving the monomer under anhydrous conditions with 0.04 equivalents of initiator (Cbz-6-aminohexanoic acid pentafluorophenol ester) in THF, followed by the addition of 0.1 equivalents of lithium bis-(trimethylsilyl)amide (LiHMDS) in THF. Our previous experimental and modeling studies on the 3,4,6-tri-*O*-TBS β-lactam indicated that the steric bulk of the monomer, specifically the TBS group at the O-3 position, poses a steric impediment to polymerization.^24^ In line with this result, monomer **1**, with bulky TBS groups at O-3 and O-6, does not polymerize even after >48 hours of stirring or after treating with higher loadings of LiHMDS (0.2 and 0.5 equivalents).

Polymerization of monomer **2**, bearing the less bulky TES groups at O-3 and O-6, proceeds under these conditions; however, it does not reach full conversion. Approximately 33% of the monomer remains unreacted even after 106 hours, as estimated via ^1^H NMR by comparing the anomeric proton peak of the polymer at 5.74 ppm to the anomeric proton peak of the monomer at 5.46 ppm (Figure S1A).

The final monomer containing a bulky silyl group, monomer **6**, does polymerize under initial conditions to give the protected polymer in quantitative yield. This result indicates that although large silyl groups at O-3 are detrimental to polymerization, they do not pose the same impediment when present instead at O-4. As before, we monitored the polymerization via ^1^H NMR by observing the disappearance of the lactam peak at 6.06 ppm and the emergence of the broad amide backbone peak at 8.12 ppm (Figure S1D). Unfortunately, the next step, which involves the selective deprotection of O-3 and O-6, was unsuccessful. PMB deprotection using DDQ/H_2_O^25^ or TfOH/1,3-dimethoxy benzene^26^ results in complete deprotection of PMB groups at O-3 and O-6 as well as partial or complete deprotection of the TIPS group at O-4 (Figures S2 and S3). PMB deprotection by Birch reduction was also ineffective.

Monomer **4**, which incorporates a highly stable benzyl protecting group at O-4 instead of TIPS, polymerizes efficiently. Monitoring the reaction by ^1^H NMR reveals complete loss of the anomeric proton of the monomer at 5.40 ppm and its replacement by the broad anomeric proton of the polymer at 5.67 ppm after only 30 seconds (Figure S1B). As anticipated, selective PMB deprotection under oxidizing conditions (DDQ/H_2_O) also proceeds efficiently. We then sulfated the deprotected polymer using SO_3_•pyridine to obtain a white fluffy disulfated polymer. However, removal of the benzyl group at O-4 by Birch reduction proved difficult due to the low solubility of the highly sulfated polymer in THF, the co-solvent required for the reaction. Nevertheless, we evaluated the anticoagulant activity of 3,6-disulfate-4-*O*-benzyl-PAS with a degree of polymerization of 50 (N=50) using activated partial thromboplastin time (aPTT), an *ex vivo* clotting assay (Figure S4). At a concentration of 100 µg/mL, the clotting time increased by 110 seconds relative to the buffer control, confirming our hypothesis that 3,6-disulfated PASs are anticoagulants and are more potent than the previously reported randomly sulfated analogue of the same size, glu-PAS-50-HS (HS refers to “high sulfation” sulPAS, where glucose PAS was reacted with 5 eq. of SO_3_•NMe_3_ at 50°C for 5 days).^19^ Due to concerns about the potential toxic degradation products of this protected polymer (e.g., benzyl alcohol, benzene, toluene), we next turned to monomer **5,** in which the O-4 position is protected by a more biocompatible ethyl group.

Polymerization of monomer **5** proceeds efficiently and in high yields, similar to monomer **4**. Kinetic analysis indicates complete monomer consumption by 30 seconds with a rate constant > 0.11 s^−1^ as determined via ^1^H NMR by observing the H-1 peak of the monomer at 5.52 ppm as it broadens and shifts to 5.65 ppm when the lactam polymerizes (Figure S1C). The ^1^H NMR spectrum possesses distinct peaks corresponding to the polymeric amide peak (8.09 - 7.68 ppm), aromatic groups (7.42 - 6.49 ppm), ethyl CH_3_ groups (0.86 ppm) and anomeric proton peak (5.65 ppm), with additional broad, overlapping peaks between 4.77 ppm to 2.62 ppm. The IR spectrum contained peaks corresponding to the amide NH stretch (3326 cm^−1^), aromatic and aliphatic CH stretch (2915 cm^−1^) and amide C=O stretch (1675 cm^−1^) (Figure S5). Next, we selectively deprotected the PMB groups at O-3 and O-6 by treating the polymers with TfOH/1,3-dimethoxybenzene (Figure 2B). ^1^H NMR analysis on the deprotected polymers shows an absence of aromatic protons (6-8 ppm) consistent with the removal of PMB groups (SI, Polymer NMRs). Finally, we sulfated the resulting polymers using SO_3_•NMe_3_ in anhydrous dimethyl formamide before dialyzing against deionized water to obtain **3,6-disulfate-4-*O*-ethyl-PAS** (**disulPAS**).

Initial aPTT results on **disulPAS** (N=50) show comparable anticoagulant activity to its benzyl counterpart, with no significant difference in clotting time (Figure S4). These findings encouraged us to proceed with **monomer 5**.

We next optimized for bioactivity as a heparin mimetic by preparing a polymer library with **disulPASs** of varying molecular weights. Specifically, we varied the equivalents of initiator used for the polymerization reaction, using [I]_0_ = [M]_0_/N where [I]_0_ and [M]_0_ are initial concentrations of initiator and monomer respectively, and N is equal to the theoretical degree of polymerization (5, 12, 25, 50, 70, 100, 120). GPC analysis of the initial, protected library reveals polymers with narrow polydispersities (*Đ* ≤ 1.2) and molecular weights in good agreement with the theoretical values (Figure 3A, Table 1). We analyzed the protected polymers using a refractive index (RI) detector with THF (1.0 mL/min) as the mobile phase.

**Figure 3.**
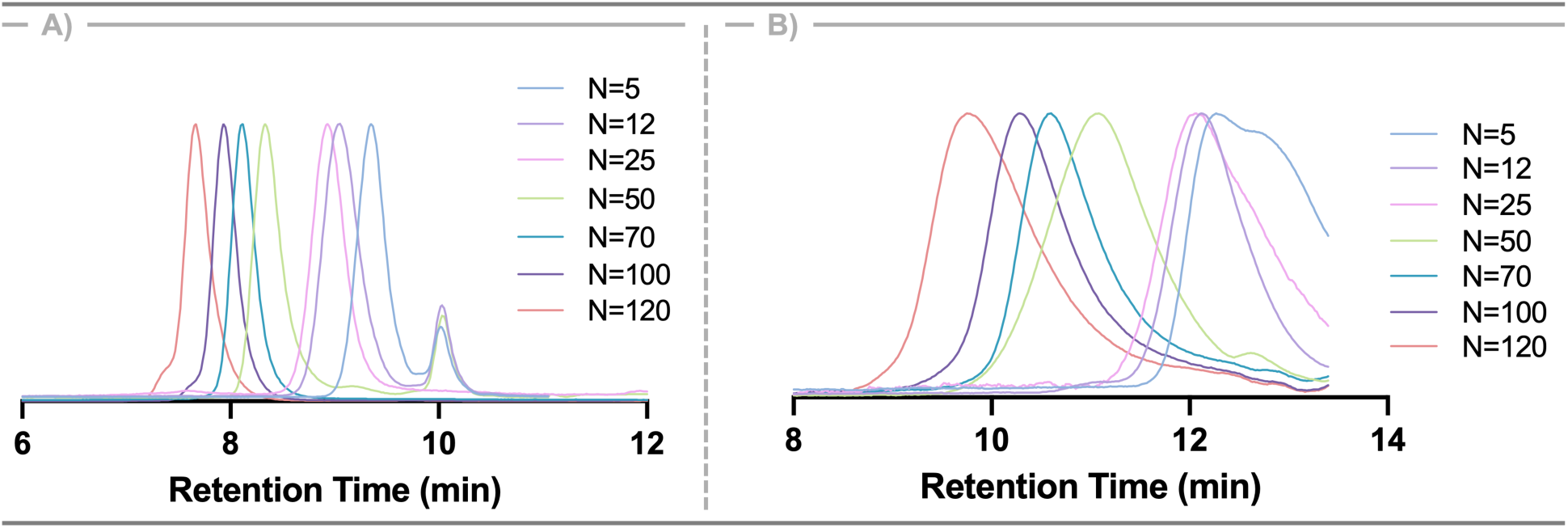
GPC chromatograms of: **A**) **3,6-di-*O*-PMB-4-*O*-Et-PAS** with N = 5, 12, 25, 50, 70, 100, 120; **B**) **3,6-disulfate-4-*O*-Et-PAS** with N = 5, 12, 25, 50, 70, 100, 120.

**Table 1.** GPC analysis of **3,6-di-*O*-PMB-4-*O*-Et PAS** and **3,6-disulfate-4-*O*-Et-PAS.**

| N Theory | THF |  |  |  | Aqueous |  |  |  | <sup>1</sup>H NMR |
| --- | --- | --- | --- | --- | --- | --- | --- | --- | --- |
|  | M<sub>n</sub> Theory (kDa) | M<sub>n</sub> Exp (kDa) | *D* | N Exp | M<sub>n</sub> Theory (kDa) | M<sub>n</sub> Exp (kDa) | *D* | N Exp | N Exp |
| 5 | 2.3 | 2.5 | 1.2 | 5 | 2.1 | 2.2 | 1.1 | 5 | 7 |
| 12 | 5.5 | 4.5 | 1.1 | 10 | 5.1 | 4.7 | 1.1 | 11 | 12 |
| 25 | 11.4 | 9.7 | 1.1 | 21 | 10.5 | 10.5 | 1.1 | 25 | 25 |
| 50 | 22.3 | 15.3 | 1.2 | 33 | 21 | 10.1 | 1.4 | 24* | 32 |
| 70 | 32.0 | 34.0 | 1.0 | 74 | 29.5 | 22.5 | 1.1 | 53* | 71 |
| 100 | 45.7 | 44.5 | 1.1 | 97 | 42.1 | 23.3 | 1.2 | 55* | 95 |
| 120 | 54.9 | 51.6 | 1.2 | 113 | 67.4 | 38.8 | 1.3 | 92* | 118 |

Following deprotection and sulfation, we analyzed the final polymers by aqueous GPC using a high salt buffer (0.5 M NaNO_3_, 0.01 M Na_2_HPO_4_, pH 7.4; 0.7 mL/min) as the mobile phase, again with RI detection. It is important to note that for N = 50 - 120 the M_n_ deduced from the GPC trace is approximately half of the expected value, however, no substantial increase in PDI or secondary peak formation was observed that would suggest polymer degradation. Molecular weight measurement efforts using SEC-MALS as well as different buffers (0.01 M - 0.75 M NaNO_3_, 1 mM - 75 mM Na_2_HPO_4_, 1x PBS) at different flow rates (0.7 - 1.0 mL/min) and temperatures (25 - 50°C) yield varying MWs. Thus, we hypothesized that the larger charged polymers interact significantly with the column, skewing the MW readout (Figure 3B, Table 1). In support of this hypothesis, the ratio of an initiator peak at 2.72 ppm to the standalone methyl peak of the ethyl group at 1.20 ppm in the ^1^H NMR spectra of the final polymers indicate degrees of polymerization consistent with those of the protected polymers and expected molecular weights (Figure S6, Table 1).

We further characterized the **disulPASs** using 1D and 2D NMR, IR, and CD (Figures S7, S8, S9, and Polymer NMRs). Compared to 3,6-di-OH-4-*O*-Et-PAS and to non-sulfated glucose PAS, **disulPAS** exhibits a downfield shift of the H-3 and H-6 protons in the ^1^H NMR spectrum. Specifically, H-3 appears at 4.94 ppm (4.15 ppm in glucose PAS (gluPAS), 3,6-di-OH-4-*O*-Et-PAS, and 6-sulfate-PAS (6-sulPAS)) and H-6 appears at 4.44 ppm and 4.27 ppm (similar to 6-sulPAS which appear at 4.30 ppm and 4.15 ppm, and downfield from 3.77 ppm in gluPAS and 3.68 ppm for 3,6-di-OH-4-*O*-Et-PAS).^24,27^ This downfield shift of the carbohydrate proton peaks after sulfation is due to the electron withdrawing effect of the sulfate groups at O-3 and O-6. In contrast, H-1, H-2, and H-4 for **disulPAS** exhibit comparable chemical shifts to those of gluPAS and 6-sulPAS. Immediately after dialysis, the ^1^H NMR also shows a sharp peak at 2.93 ppm, indicative of residual NMe_3_ from the SO_3_•NMe_3_ reagent. We removed this small molecule impurity by treating the polymer with 1M NaOH under high vacuum for 3 hours.^19^ To confirm that **disulPAS** is not degraded by the NaOH treatment, we performed GPC analysis before and after base exposure. Both polymers elute at the same retention time, confirming the absence of degradation (Figure S7). We used NaOH treated polymers for all subsequent studies.

### Evaluation of anticoagulant activity

The effect of anticoagulants on the coagulation cascade is clinically assessed by two *ex vivo* clotting assays: activated partial thromboplastin time (aPTT) and prothrombin time (PT). aPTT and PT examine the effect of the anticoagulants on the intrinsic and extrinsic pathway of coagulation, respectively (Figure 4). Prolongation of clotting time on either of these tests demonstrates an anticoagulant effect. Elongation observed in both aPTT and PT, as seen for UFH, indicates an effect on the common pathway, which is downstream to both intrinsic and extrinsic pathways. Control test groups include HBS buffer without polymer, 5.5 µg/mL (1 IU/mL) of UFH (8-15 kDa), and 100 µg/mL of glu-PAS-50,^24^ glu-PAS-HS-50,^19^ and 6-sulPAS-50^27^ (suffix refers to degree of polymerization, N; concentrations of polymer are prior to addition to plasma). aPTT results indicate glu-PAS-50, 6-sulPAS-50, glu-PAS-HS-50, **disulPAS-5**, and **disulPAS-12** do not significantly elongate clotting time at 100 µg/mL compared to the negative control (HBS without polymer). In contrast, **disulPAS-25, -50,** and **-70**, produce statistically significant increases in clotting time at 100 µg/mL. While **disulPAS-100** and **disulPAS-120** also significantly increase clotting time, they are less effective than **disulPAS-70**, indicating a decrease in anticoagulant activity at higher molecular weights, which we suspect is due to steric hindrance (Figure 5A). Both 3,6-disulfate-4-*O*-Bn-PAS-50 and 3,6-disulfate-4-*O*-Et-PAS-50 exhibit similar clotting times, suggesting the identity of a hydrophobic group at O-4 minimally impacts anticoagulant activity (Figure S4). Additionally, both 3,6-disulfated polymers show 1.5-fold greater anticoagulant activity than the mono-sulfated 6-sulPAS-50 and randomly sulfated glu-PAS-HS-50, leading us to conclude that sulfation at the 3 position is crucial for increased anticoagulant activity in sulfated PASs (Figure 5E).

**Figure 4.**
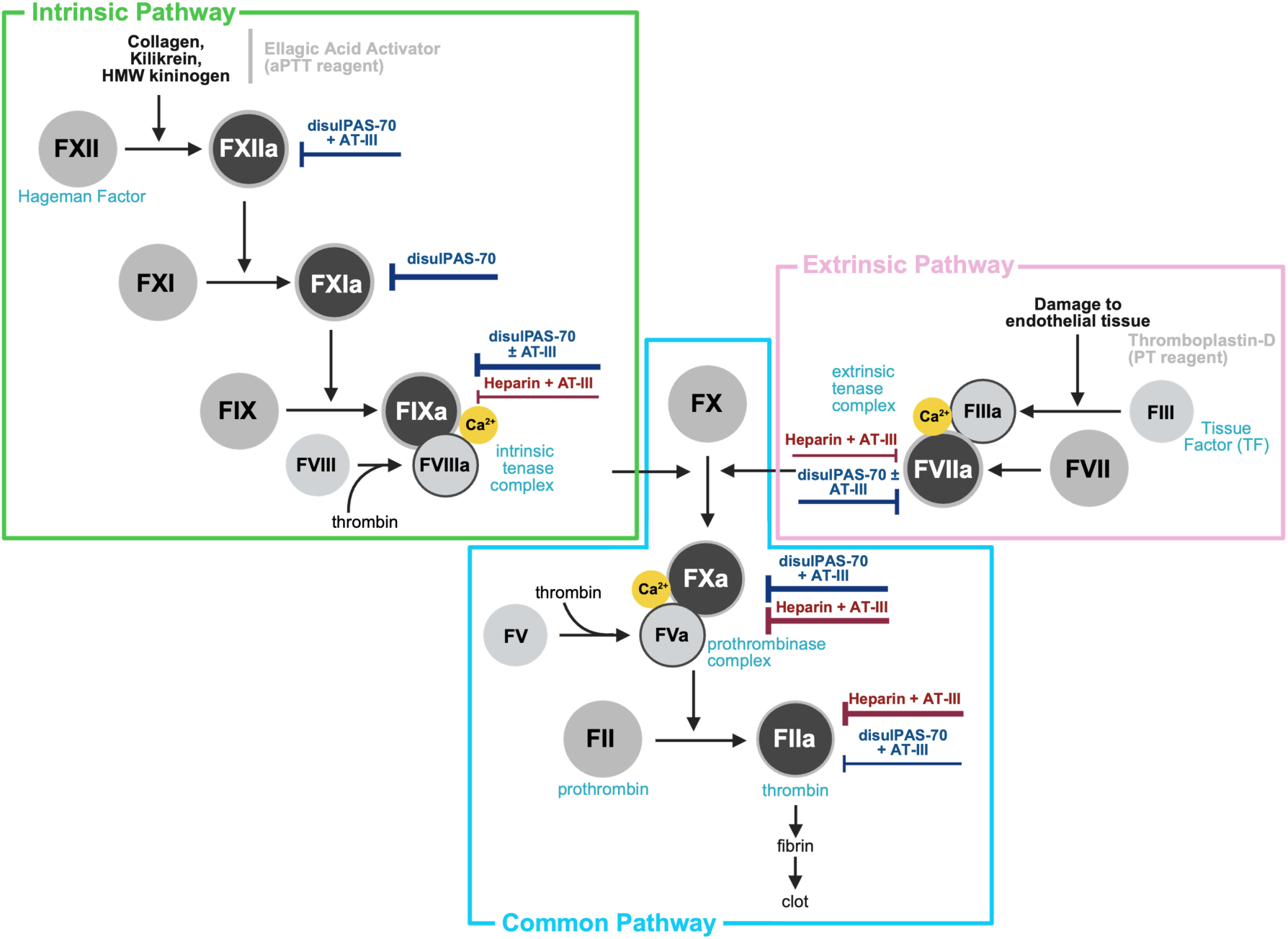
The coagulation cascade indicating the effect of **disulPAS-70** on different clotting factors.

**Figure 5:**
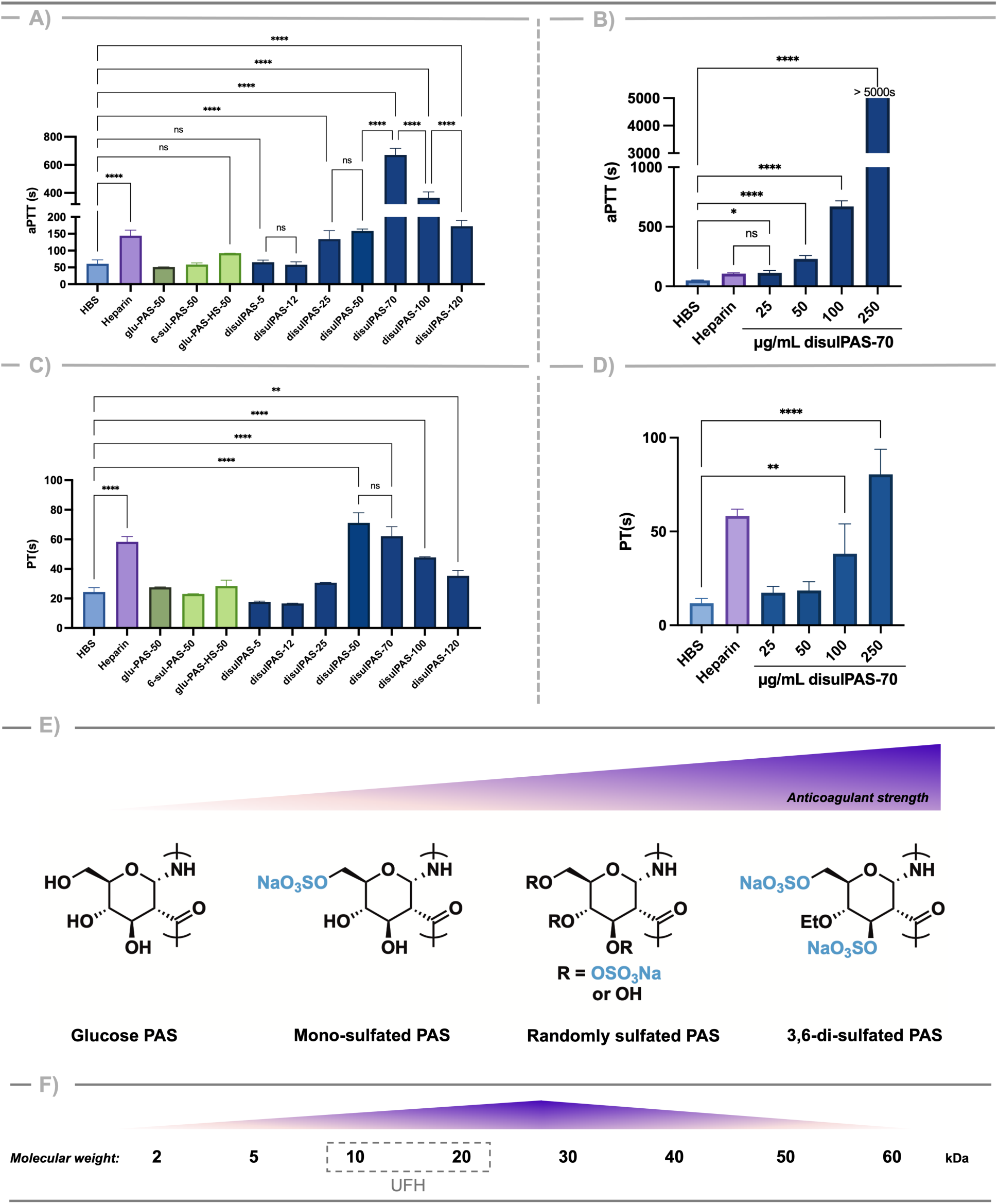
**A**) aPTT assay of control polymers and **disulPASs** at a concentration of 100 µg/mL; **B**) Dose dependent aPTT on **disulPAS-70**; **C**) PT assay of control polymers and **disulPASs** at a concentration of 100 µg/mL; **D**) Dose dependent PT on **disulPAS-70**. All concentrations listed are prior to addition of analyte to plasma (1:2). Heparin control is 1 IU/mL or 5.5 µg/mL before addition to plasma. Data represent mean ± SD, n = 3, *P ≤ 0.05, **P ≤ 0.01 ***P ≤ 0.001, ****P ≤ 0.0001, ns= not statistically significant, unpaired t-test. **E**) Sulfation density dependence of anticoagulant strength for all sulfated PASs; **F**) Molecular weight dependence of anticoagulant strength for **disulPAS**.

With this knowledge in hand, we next examined the effect of **disulPAS-70** concentration on anticoagulant activity. **DisulPAS-70** exhibits a dose-dependent prolongation of clotting time, with a statistically significant increase relative to HBS control at concentrations as low as 25 µg/mL (Figure 5B). At 250 µg/mL, clotting does not occur even after 5000 seconds, indicating the high potency of this anticoagulant. Notably, the clotting time at 25 µg/mL **disulPAS-70** is equivalent to that of 5.5 µg/mL UFH, thus benchmarking the anticoagulant activity of **disulPAS-70** at approximately one-fifth of UFH on a mass basis, or one-half on a molar basis (approximately 847 nM disulPAS compared to 478 nM UFH, assuming 29.5 and 11.5 kDa MW, respectively).

We next evaluated PT in the presence of **disulPAS**. Similar to aPTT, the PT data exhibit a bell-curve dependence on the degree of polymerization; prolongation in PT time increases with degree of polymerization, reaches a maximum at approximately N = 70, and then reduces for N = 100, and 120 (Figure 5C). Interestingly, the smallest polymers (N = 5 and 12) decrease clotting time relative to the buffer at 100 µg/mL, acting as mild procoagulants. This procoagulant effect in the PT assay, along with the absence of aPTT prolongation, indicates that the 5-mer and 12-mer are poor anticoagulants. Consistent with the aPTT results, **disulPAS-70** produces a dose dependent increase in PT (Figure 5D), indicating that the **disulPASs** exert a more pronounced effect on the extrinsic and common pathways of coagulation than our previous generation of randomly sulfated PASs. When compared to other sulfated polysaccharides and heparin mimetics, **disulPASs** performs favorably. λ-carrageenan, a sulfated polysaccharide isolated from seaweed, reaches the activity of 5 µg UFH in aPTT and PT at 20 µg.^28^ Fucosylated chondroitin sulfates also exhibit some anticoagulant activity but overall are not as effective as heparin at any concentration tested;^29^ the same effect is reported for sulfated ginger polysaccharides.^30^ Depolymerized fucosylated chondroitin sulfate matches UFH activity at 50 µg/mL and outperforms heparin at higher concentrations, but the same health risks remain as with chondroitin sulfate.^31^ Synthetically, sulfated polyureas and sulfated pseudo-polysaccharides demonstrate anticoagulant activity about one-tenth that of UFH.^32,33^ The most structurally similar synthetic mimetic to heparin, a sulfated polymer made from glucose-based cyclic thionocarbonates, exhibits comparable elongation in aPTT clotting time to the UFH control (5.5 µg/mL) at 50 µg/mL.^33^ Overall, these results distinguish **disulPAS-70** as a strong contender for a heparin mimetic, and, thus, we next investigated its mechanism of action and evaluated its *in vivo* performance.

### Elucidation of mechanism of action

Given the strong *ex vivo* anticoagulant activity exhibited by the aPTT and PT tests, we next evaluated the effects of **disulPAS-70** on various clotting factors of the intrinsic and extrinsic pathways of coagulation using amidolytic activity inhibition assays to ascertain its mechanism of action. Due to the structural similarity of **disulPAS** and heparin, we hypothesized that its mechanism of action would also be AT-III-mediated. To test this hypothesis, we evaluated inhibition of serine proteases FIIa, FVIIa, FIXa, FXa, FXIa, and FXIIa in the presence and absence of AT-III (Figure 6). We incubated varying concentrations of **disulPAS-70** (4.44 nM - 44.4 µM) with each coagulation factor in the presence or absence of 0.8 µM AT-III and determined percent inhibition using factor-specific colorimetric substrates.

**Figure 6:**
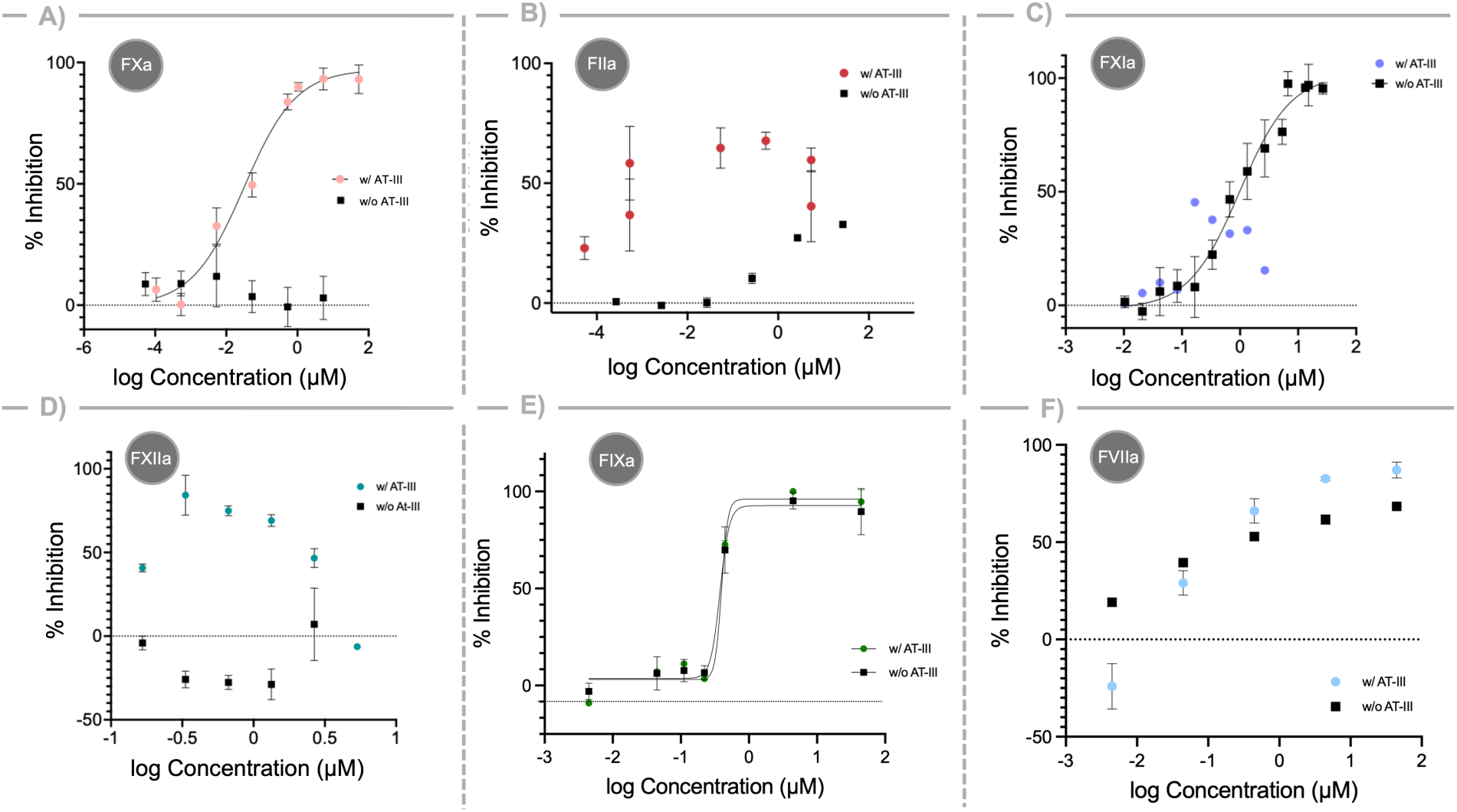
Activity inhibition of coagulation factors by **disulPAS-70**. Amidolytic activity was monitored at 405 nm by incubating each coagulation factor with the designated concentration of polymer with or without AT-III, followed by addition of enzyme-specific chromogenic substrate. **A**) FXa; **B**) FIIa; **C**) FXIa; **D)** FXIIa; and **E**) FIXa; and, **F**) FVIIa. Data represents mean ± SD, n = 3.

### Inhibition of clotting factors in the common pathway (factor Xa and factor IIa)

**DisulPAS-70** does not inhibit FXa in the absence of AT-III even at 4.44 µM (Figure 6A, black), indicating that the formation of a FXa:disulPAS complex does not contribute to its anticoagulant effect. In the presence of AT-III, however, **disulPAS-70** does inhibit FXa with an IC_50_ of 0.16 µM (Figure 6A, pink), consistent with a heparin-like, AT-III–mediated mechanism, albeit with lower potency than heparin (IC_50_ = 0.02 µM).^19^

For FIIa, **disulPAS-70** partially inhibits in the absence of AT-III, reaching a maximum of ∼33% at 22.2 µM (Figure 6B, black). In the presence of AT-III, the inhibition curve displays an ascending limb with a maximum inhibition of 68% at 0.44 µM, followed by a pseudo-plateau and a descending limb at higher concentrations (Figure 6B, red). This non-monotonic profile mirrors that observed for heparin^34^ and for our earlier randomly sulfated PAS, glu-PAS-HS-50, indicating a template mechanism of inhibition.^19^

### Inhibition of clotting factors in the intrinsic pathway (factor XIa, factor XIIa, factor IXa)

AT-III weakly but progressively inhibits FXIa,^35^ with inhibition accelerated by 40-fold in the presence of heparin.^36^ While heparin does not directly inhibit FXIa, dextran sulfate, a heparin mimic, inhibits FXIa by up to 50%.^37^ Consistent with dextran sulfate, randomly sulfated PASs directly inhibit FXIa in the absence of AT-III, with glu-PAS-HS-50 exhibiting an IC_50_ of 15 µM.^19^ Further, we found that **disulPAS-70** also inhibits FXIa in the absence of AT-III with 15-fold higher efficacy than glu-PAS-HS-50 (IC_50_ = 0.98 µM) (Figure 6C, black). Interestingly, in the presence of AT-III, maximum inhibition of FXIa by **disulPAS-70** diminishes to 50% (Figure 6C, purple), indicating that AT-III does not mediate FXIa inhibition in this system. Factor XIa inhibitors are of increasing interest as anticoagulants as they act on the intrinsic pathway and may reduce bleeding risks often associated with sulfated anticoagulants.^38^ Many potent FXIa inhibitors, such as milvexian and asundexian, bind directly to the FXIa catalytic pocket,^39,40^ whereas heparin binds both allosterically and to a basic region of the active site.^41,42^ Additional work is ongoing to deduce where on the enzyme **disulPAS-70** binds, and its mechanism of inhibition.

For FXIIa, which lies upstream of FXIa, AT-III inhibits the protease and this inhibition enhances 4-fold by heparin.^37^ **DisulPAS-70** does not directly inhibit FXIIa in the absence of AT-III at concentrations of up to 2.2 µM, but slightly activates FXIIa at lower concentrations (Figure 6D, black). In the presence of AT-III, however, **disulPAS-70** inhibits FXIIa with a maximum inhibition of 84% at 0.33 µM (Figure 6D, green).

FIXa, the final factor examined in the intrinsic pathway, lies downstream of FXIIa and FXIa and forms a complex with FVIIIa to activate FX. Heparin weakly inhibits FIXa in the presence of AT-III with an inhibition profile similar to that for FIIa.^43^ In contrast, treatment with **disulPAS-70** affords complete inhibition of FIXa with an IC_50_ of 0.38 µM in both the presence and absence of AT-III (Figure 6E), indicating a largely AT-III-independent mechanism.

### Inhibition of clotting factors in the extrinsic pathway (factor VIIa)

Finally, we examined the extrinsic pathway by assessing FVIIa inhibition by **disulPAS-70** in the presence or absence of AT-III. FVIIa associates with tissue factor to form the extrinsic tenase complex, which activates FX and initiates clot formation. Heparin inhibits FVIIa activity by up to 40% in the presence of AT-III.^44^ Treatment with **disulPAS-70** at 4.4 µM along with AT-III results in 82% inhibition of FVIIa activity, which plateaus at 87% at 44.4 µM (Figure 6F, blue). In the absence of AT-III, a slight activation of FVIIa occurs at 4.4 nM, followed by increasing inhibition up to 68% at 44 µM (Figure 6F, black). The ability of **disulPAS-70** to inhibit FVIIa in the absence of AT-III contrasts with heparin, which shows limited inhibition only in the presence co-factors.

To summarize the mechanism, we observed that **disulPAS-70** with or without AT-III leads to moderate (60 - 99%) or complete (100%) inhibition of every factor in the coagulation cascade. In the presence of AT-III, all coagulation factors: IIa, VIIa, IXa, Xa, XIa, and XIIa, are inhibited by **disulPAS-70** at least partially. Without AT-III, **disulPAS-70** inhibits factors VIIa, IXa, and XIa. Comparatively, heparin exhibits inhibitory effects on the coagulation cascade only in the presence of cofactors such as AT-III, primarily acting on factors Xa and IIa. A weak inhibitory effect occurs with factors IXa and VIIa, and heparin is not reported as a significant inhibitor of factors XIa and XIIa. These results are summarized in Figure 4.

### Safety and biocompatibility of disulPAS

#### Cytocompatibility and hemocompatibility

A major shortcoming of animal-derived heparin and heparin mimics is contamination with toxic impurities such as chondroitin sulfate, which can cause severe adverse effects.^30^ Therefore, we evaluated the cytocompatibility and hemocompatibility of **disulPAS-70** using a standard MTS assay and a hemolysis assay. We treated NIH-3T3 fibroblasts with varying concentrations (0.0002 - 3.0 mg/mL) of sterile **disulPAS-70**, dextran (*Leuconostoc spp*, molecular weight ∼40 kDa), glu-PAS-100, or UFH, and measured the cell viability after 24 hours. **DisulPAS-70** is non-cytotoxic up to 1 mg/mL, similar to the control polymers (dextran, glu-PAS-100 and UFH). At 3 mg/mL, cell viability decreases to 50% with **disulPAS-70** and dextran, while cytotoxic effects remained minimal for glu-PAS-100 and UFH (Figure 7A). Importantly, at 25 µg/mL, the concentration at which **disulPAS-70** shows comparable anticoagulant activity to 1 IU/ mL UFH (113 s as compared to 107 s for UFH), no cytotoxicity is present.

**Figure 7:**
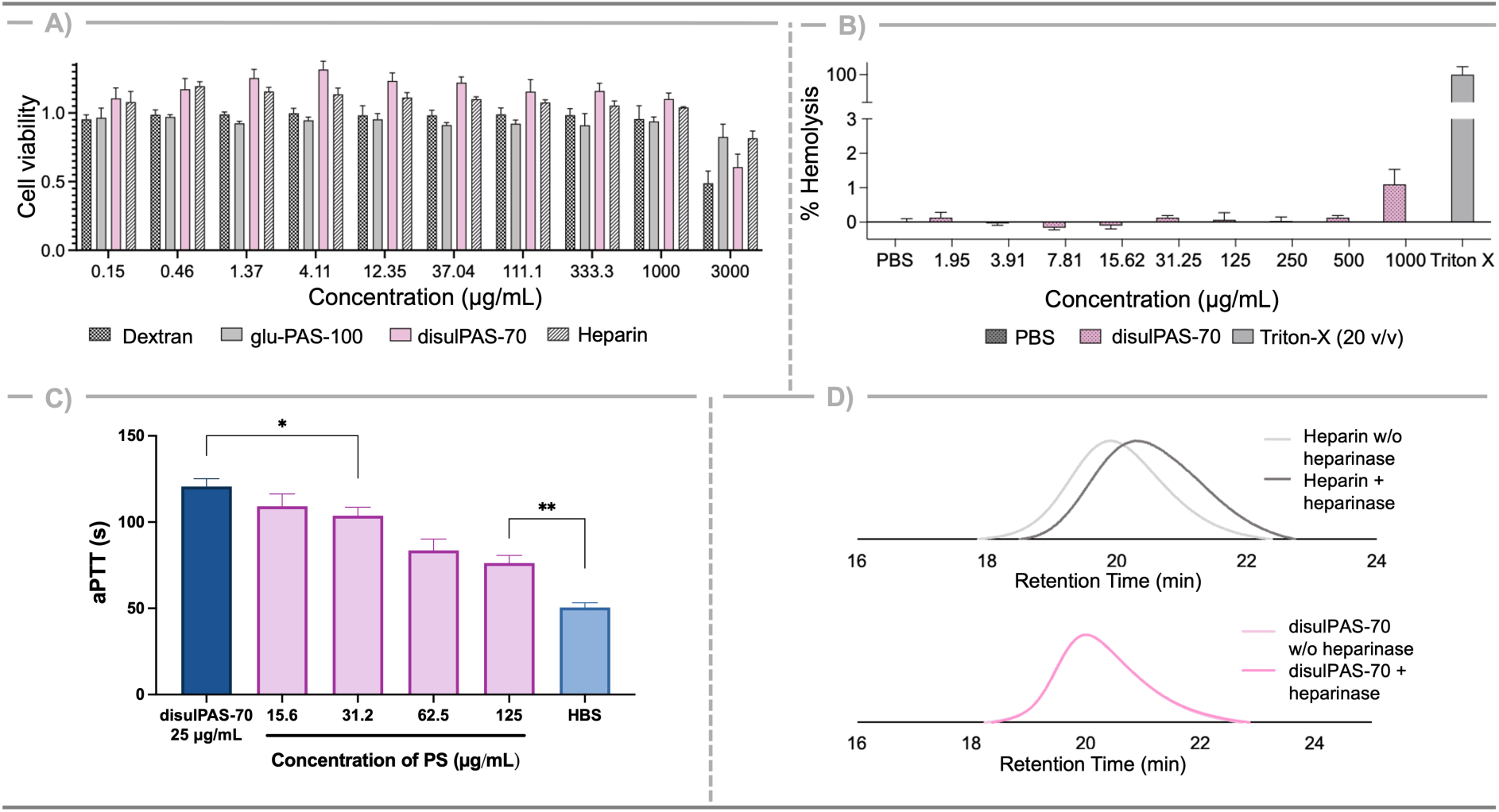
**A**) Cytotoxicity assay; **B**) Hemolysis assay; **C**) Reversibility by protamine sulfate (PS); **D**) Heparinase degradation. Data represent mean ± SD, n = 3, *P ≤ 0.0114, **P ≤ 0.001, unpaired t-test.

For hemocompatibility, we evaluated the propensity of **disulPAS-70** to lyse red blood cells (RBCs). Human RBCs were incubated with **disulPAS-70** (1.95 - 1000 µg/mL) at 37°C for 24 hours. **DisulPAS-70** does not induce hemolysis at concentrations≤ 500 µg/mL and causes only very minimal hemolysis (∼1%) at 1000 µg/mL (Figure 7B).

#### Reversal of anticoagulant effect by protamine sulfate

One of the major advantages of UFH over other synthetic FXa inhibitors, such as fondaparinux (Arixtra), is that protamine sulfate (PS), a cationic polypeptide, halts anticoagulant activity *in vivo* in cases of excessive bleeding. To investigate whether PS neutralizes **disulPAS**, we incubated 25 µg/mL **disulPAS-70** in normal human plasma with varying concentrations of PS (15.6 - 125 µg/mL) and determined the aPTT. 25 µg/mL **disulPAS** was again chosen given it exhibits the same prolongation in aPTT as 1 IU/mL UFH. We observed that clotting time decreases from 109 seconds at 31.25 µg/mL PS to 77 seconds at 125 µg/mL PS (Figure 7C). The aPTT at 125 µg/mL PS remains statistically higher than that of the HBS control, and no further decrease in clotting time is present at higher PS concentrations. Thus, protamine sulfate partially neutralizes the anticoagulant effect of **disulPAS-70**.

#### Heparinase degradation

Another drawback of UFH is its susceptibility to degradation by heparinase (HPSE), an enzyme elevated in the plasma of patients with diabetes and renal insufficiency. Further, interactions between heparin and HPSE play critical roles in the progression of a variety of diseases, including cancer.^45^ Similar to our earlier work showing that sulPASs are resistant to HPSE, we used size-exclusion chromatography to demonstrate that **disulPAS-70** is likewise not degraded by HPSE (Figure 7D).

#### In vivo anticoagulant activity of disulPAS in murine deep vein thrombosis (DVT) model

Following the *in vitro* and mechanistic studies, we evaluated the *in vivo* performance of **disulPAS-70** after intravenous (IV) administration (0.75, 1.5, and 4.5 µg/g). As shown in Figure 8A, treatment with **disulPAS-70** affords a dose-dependent increase in aPTT in mice, consistent with the *in vitro* dose-response behavior observed in Figure 5B. We next assessed the pharmacokinetic/ pharmacodynamic (PK/PD) profile of **disulPAS-70** using aPTT following both IV and subcutaneous (SC) administration. 2 minutes after IV administration of 0.75 µg/g **disulPAS-70**, the aPTT value is 168 seconds; this prolongation then rapidly declines over time and returns to baseline at ∼2 hours post-administration (Figure 8B). Pilot studies for SC administration (1.5 - 7.5 µg/g, SC; Figure S10) demonstrate that peak aPTT occurs at 2 hours for all doses, with the anticoagulant effect dissipating by ∼6 hours. Thus, we performed a more detailed study at a single SC dose (7.5 µg/g), revealing a maximum aPTT of 174 seconds at 2 hours (Figure 8C), a value expected to fall within the therapeutic range for anticoagulation.

**Figure 8.**
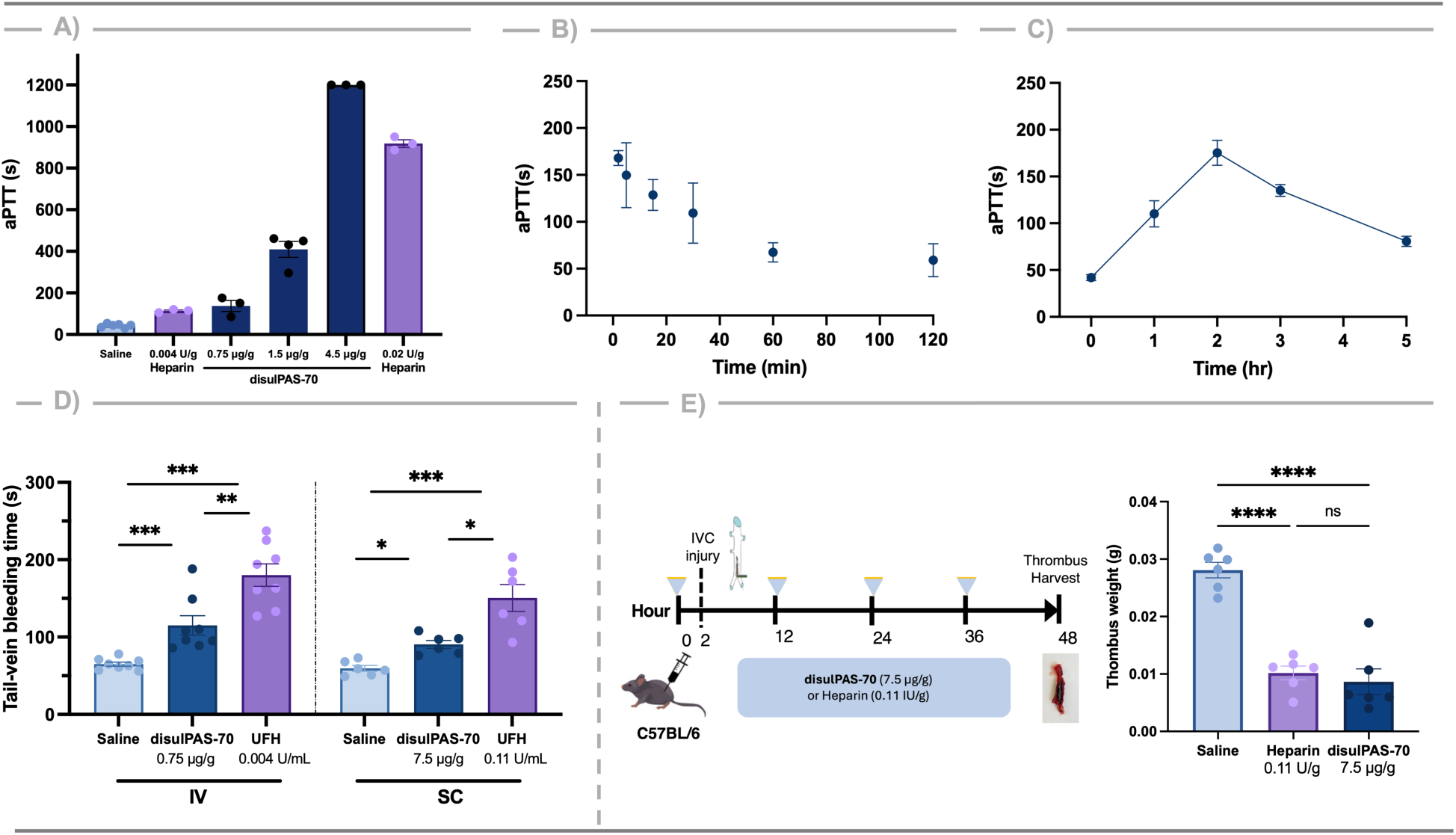
Evaluation of anticoagulant and antithrombogenic disulPAS-70 in murine deep vein thrombosis (DVT) model. (**A**) HBS, heparin (0.004 U/g, 0.02 U/g), or **disulPAS-70** (0.75 – 4.5 µg/g) were administered (IV) to C57BL/6 mice 2 minutes prior to blood draw, and plasma clotting time was measured. **DisulPAS-70** demonstrated a dose-dependent increase in aPTT. (**B**) Plasma clotting time (aPTT) was determined over a 2 hour period after IV administration of **disulPAS-70** (0.75 µg/g, n ≥ 3 mice/ time point), or (**C**) over a 5 hour period after SC administration of **disulPAS-70** (7.5 µg/g, n ≥ 5 mice/time point). (**D**) Tail vein bleeding time was performed 2 min post-IV administration or 2 hours post-SC administration (n = 5 - 8 mice per agent). (**E**) The thrombus weight was obtained 2 days after electrolytic injury of the vena cava following SC administration of saline vehicle (n = 6), heparin (0.11 U/g, 2 hr-pre + every 12 hr thereafter, n = 6), or **disulPAS-70** (7.5 µg/g, 2 hr-pre+ every 12 hr thereafter, n = 6). Data represent mean ± SEM, *P ≤ 0.05, ***P ≤ 0.001, ****P ≤ 0.0001 (ANOVA and post hoc Student’s t-test).

We next determined the effect of **disulPAS-70** on hemostasis using a mouse tail-transection bleeding time assay following IV (0.75 µg/g) or SC (7.5 µg/g) administration. UFH (0.004 U/g IV; 0.11 U/g SC) served as a clinical reference. Both heparin and **disulPAS-70** significantly increase bleeding time (151 and 99 sec, respectively) compared to the saline vehicle at 60 sec (Figure 8D). The antithrombotic and bleeding profiles of heparin and **disulPAS-70** are summarized in Table 2. These results suggest that **disulPAS-70** can be dosed to achieve a clinically relevant increase in aPTT with a bleeding risk less than that of heparin.

**Table 2.** Summary of anticoagulant properties of **disulPAS-70** *in vivo*.

|  | Saline | UFH | **disulPAS-70** |
| --- | --- | --- | --- |
| Dosage | n/a | 0.11 U/g | 7.5 µg/g |
| Administration | 2 hr pre, SC | 2 hr pre, SC<br>+ every 12 hr | 2 hr pre, SC<br>+ every 12 hr |
| Avg. bleeding time (sec) | 60 ± 9 | 151 ± 42 | 99 ± 35 |
| Avg. thrombus weight (mg) | 0.0195 ±<br>0.0036 | 0.0085 ± 0.0060 | 0.0093 ± 0.0032 |
| % Thrombus weight reduction | n/a | 64% | 69% |

Finally, we evaluated the efficacy of **disulPAS-70** for venous thromboprophylaxis in a murine model of non-occlusive venous thrombosis. We selected the dose and regimen based on the PK/PD data. We administered **disulPAS-70** (7.5 µg/g, SC) and UFH (0.11 U/g, SC) 2 hours prior to electrolytic injury of the vena cava and every 12 hours thereafter until termination. Treatment with **disulPAS-70** results in a 69% reduction in thrombus formation relative to the saline vehicle, comparable to a 64% reduction observed with heparin (Figure 8E).

## Discussion

In an effort to develop a more consistent, homogenous, synthetic heparin mimetic, we prepared polyamidosaccharides bearing regioselective sulfation at the 3 and 6 positions using an iterative sequence of monomer synthesis, polymerization, and post-polymerization modification. We confirmed polymer composition, structure, and molecular weight by NMR, IR, GPC and CD. Functionally, **disulPASs** exhibit chain-length dependent and dose dependent prolongation of clotting time in both the extrinsic and intrinsic pathways, with **disulPAS-70** showing anticoagulant activity comparable to heparin at 25 µg/mL. Mechanistically, **disulPAS-70**, directly inhibits FVIIa, FIXa, and FXIa, while inhibition of FXa, FIIa, and FXIIa is AT-III-mediated. In *in vitro* assays, **disulPASs** are cytocompatible and hemocompatible, and their anticoagulant effects are partially reversed by protamine sulfate, supporting their safety as therapeutic candidates. *In vivo*, **disulPAS-70** prolongs clotting time in a dose-dependent manner and reduces thrombus formation in a model of non-occlusive venous thrombosis to an extent comparable to heparin. The limitations of animal-derived heparin underscore the need for novel heparin analogs, including bioengineered glycans, small-molecule agents, and polymeric anticoagulants. Overall, **disulPAS-70** is a promising new anticoagulant, possessing favorable properties of both heparin and synthetic anticoagulants, while overcoming many of their respective drawbacks. In summary, **disulPAS** represents a potential therapeutic platform for the treatment of thromboembolic disorders.

## ACKNOWLEDGMENTS

We acknowledge funding in part for this work from the National Institutes of Health (R01HL164650 ELC, MWG), National Cancer Institute of the National Institutes of Health (1F31CA298728-01, MKL), and the William Fairfield Warren Distinguished Professorship (MWG). The content of this work is solely the responsibility of the authors and does not necessarily represent the official views of the National Institutes of Health.

## DATA AVAILABILITY

The raw data required to reproduce these findings are available from the authors upon request. All additional figures, synthetic methods, small molecule characterization, and NMR spectra can be found in the Supplemental Information (PDF).

## AUTOR CONTRIBUTIONS

Conceptualization: MKL, MV, ELC, MWG

Methodology: MKL, MV, HH, CH

Investigation: MKL, MV, HH

Visualization: MKL, MV

Supervision: ELC, MWG

Writing—original draft: MKL, MV, HH

Writing—review & editing: All authors

Funding: MKL, ELC, MWG

## COMPETEING INTERESTS

All other authors declare they have no other competing interests.

