## Supplementary material for "Regio-selective sulfation of heparin mimicking polymers defines in vivo activity": SI

#### Table of Contents


**SUPPLEMENTAL FIGURES** 4

**GENERAL METHODS AND MATERIALS** 10

SIZE EXCLUSION CHROMATOGRAPHY 10

CIRCULAR DICHROISM 10

EX VIVO APTT AND PT ASSAYS 11

GENERAL AMIDOLYTIC ACTIVITY INHIBITION ASSAYS 11

CYTOTOXICITY 12

HEMOLYTIC ASSAY 13

REVERSAL OF ANTICOAGULANT ACTIVITY OF DISULPAS BY PROTAMINE SULFATE 13

EVALUATION OF DEGRADATION OF DISULPAS BY HEPARANASE 13

**IN VIVO STUDIES** 14

PHARMACOKINETIC/PHARMACODYNAMIC ANALYSIS WITH APTT ASSAY 14

*IN VIVO* EVALUATION OF ANTI-THROMBOTIC EFFICACY WITH A NON-OCCLUSIVE VENOUS THROMBOSIS MURINE MODEL 14

BLEEDING TIME MEASUREMENT 15

**SMALL MOLECULE SYNTHETIC PROCEDURES** 15

GENERAL PROCEDURE FOR CSI CYCLOADDITION 15

GENERAL PROCEDURE FOR TCAI CYCLOADDITION 16

*MONOMER 1* 16

*MONOMER 2* 18

*MONOMER 3* 19

*MONOMER 4* 19

*MONOMER 5* 21

*MONOMER 6* 22

**POLYMERIZATION AND POST-POLYMERIZATION SYNTHETIC PROCEDURES** 25

GENERAL PROCEDURE FOR POLYMERIZATION 25

GENERAL PROCEDURE FOR PMB DEPROTECTION USING TFOH/1,3-DMB 26

GENERAL PROCEDURE FOR PMB DEPROTECTION USING DDQ/$H_2O$ 26

GENERAL PROCEDURE FOR SULFATION AND NAOH WORKUP 27

**SMALL MOLECULE NMRS** 28

**POLYMER NMRS** 57

**REFERENCES** 69

#### Supplemental Figures

**Figure S1**. Evaluation of polymerization by $^{1}$H NMR; blue dots highlight polymer peaks, green dots highlight corresponding monomer peaks. **A**) **monomer 2**; **B**) **monomer 4**; **C**) **monomer 5**; **D**) **monomer 6**.

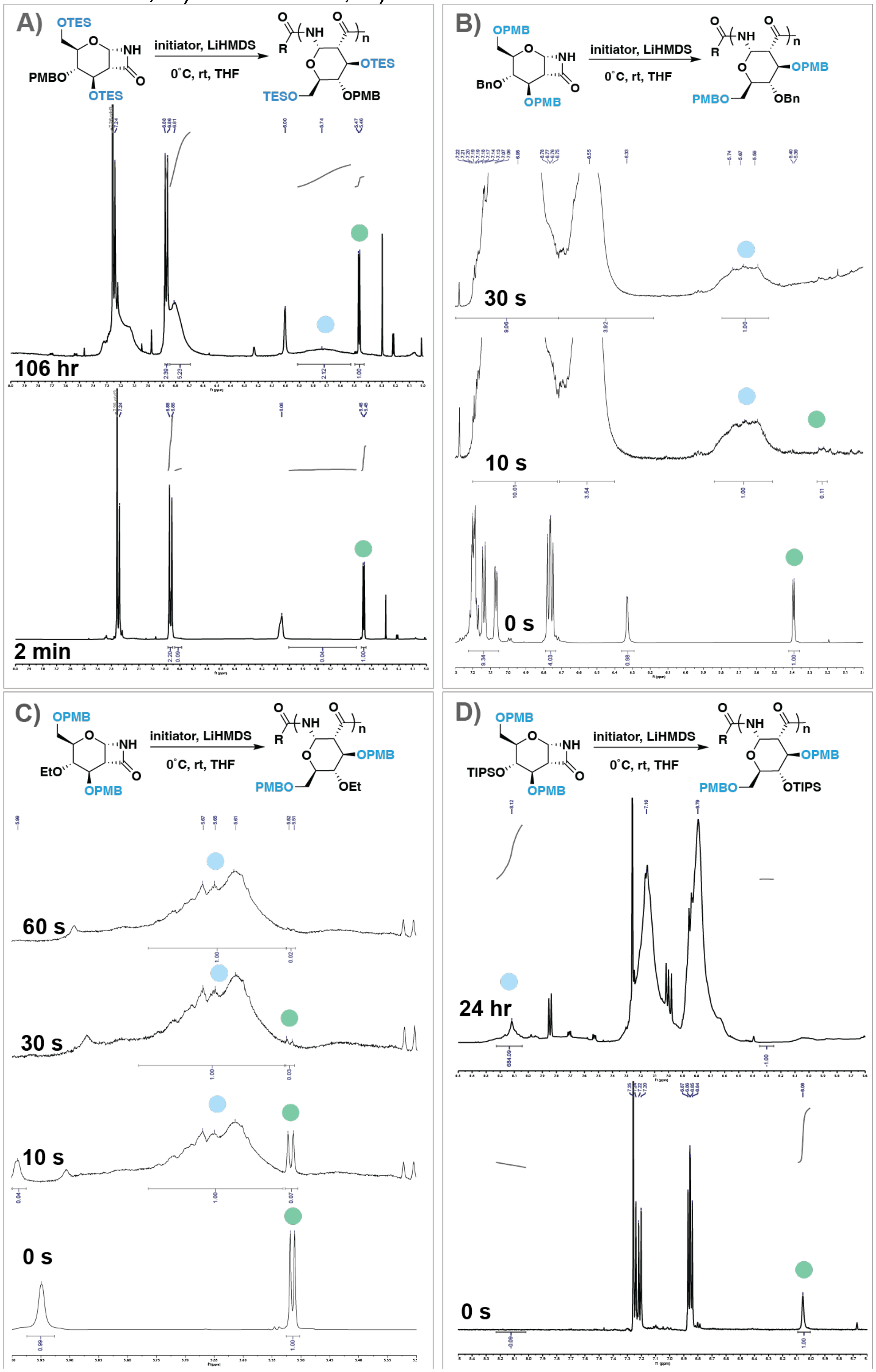

**Figure S2.** [1]H NMR of selective PMB deprotection of **3,6-di-*O*-PMB-4-*O*-TIPS-PAS** using DDQ/$H_2O$.

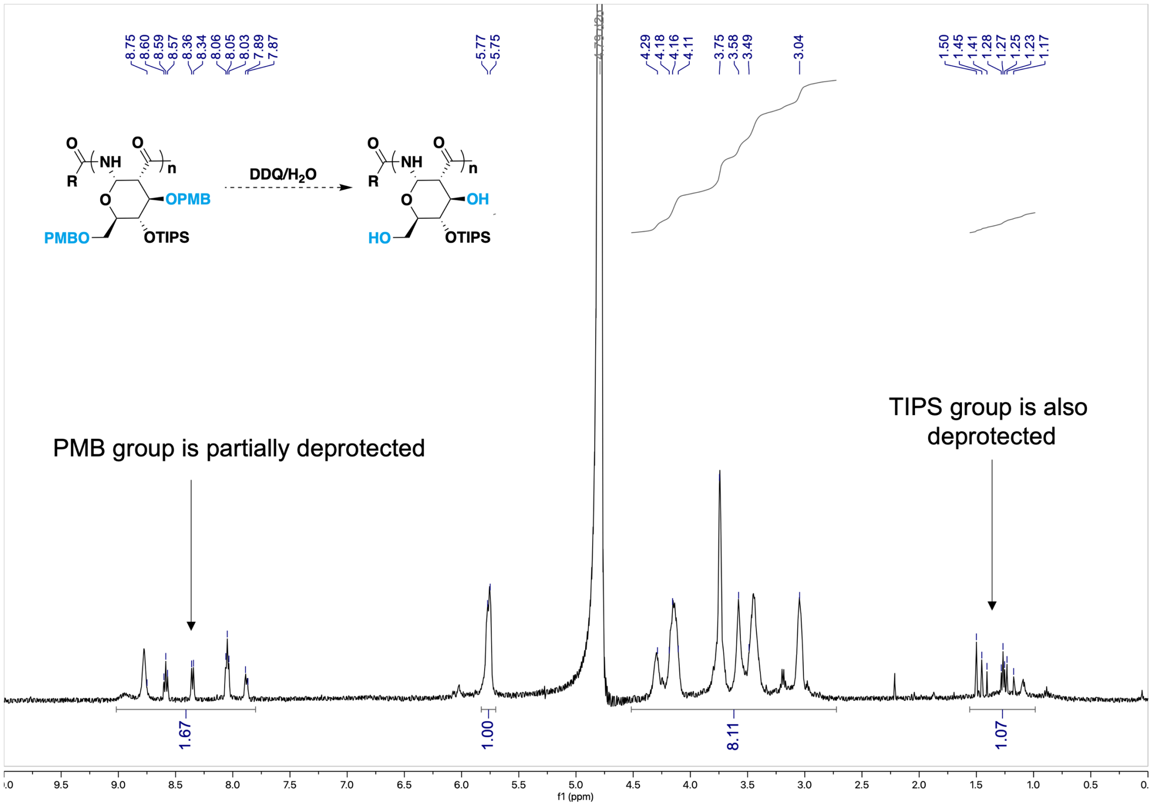


**Figure S3.** [1]H NMR of selective PMB deprotection of **3,6-di-*O*-PMB-4-*O*-TIPS-PAS** using TfOH/ 1,3-dimethoxy benzene.

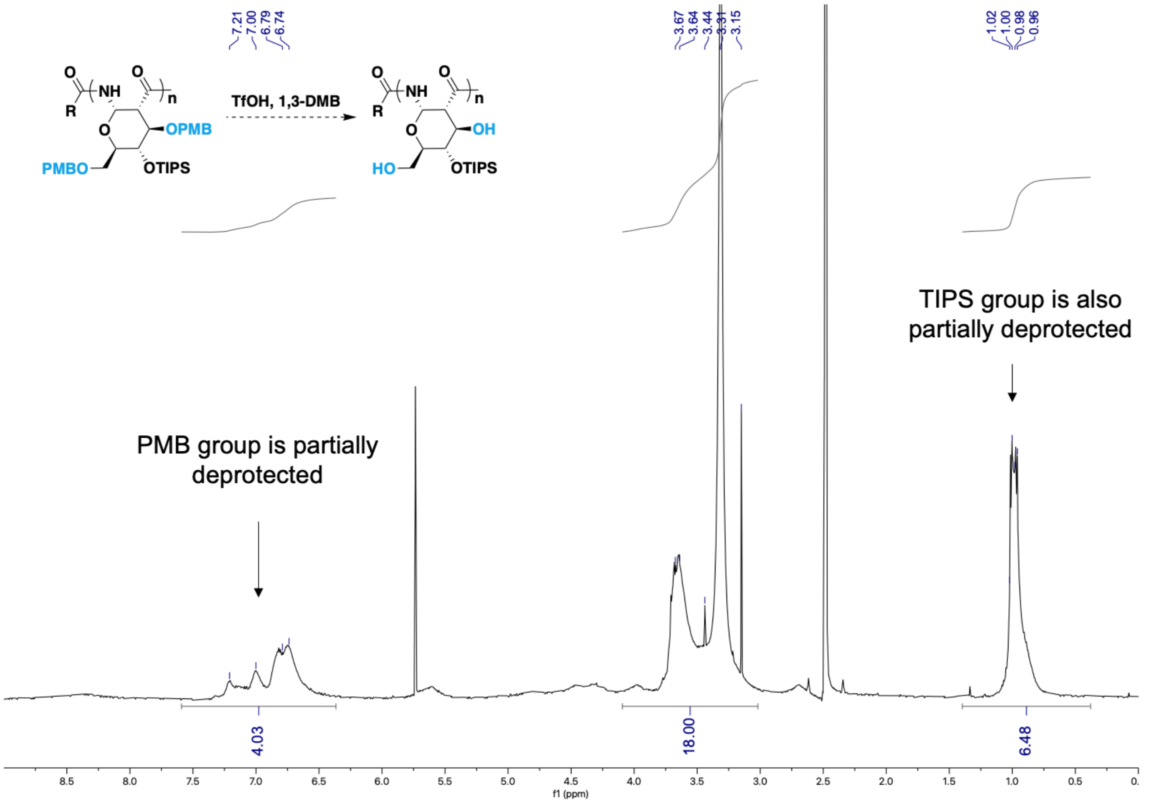

**Figure S4.** aPTT on **3,6-di-sulfate-4-*O*-Bn-PAS-50.**

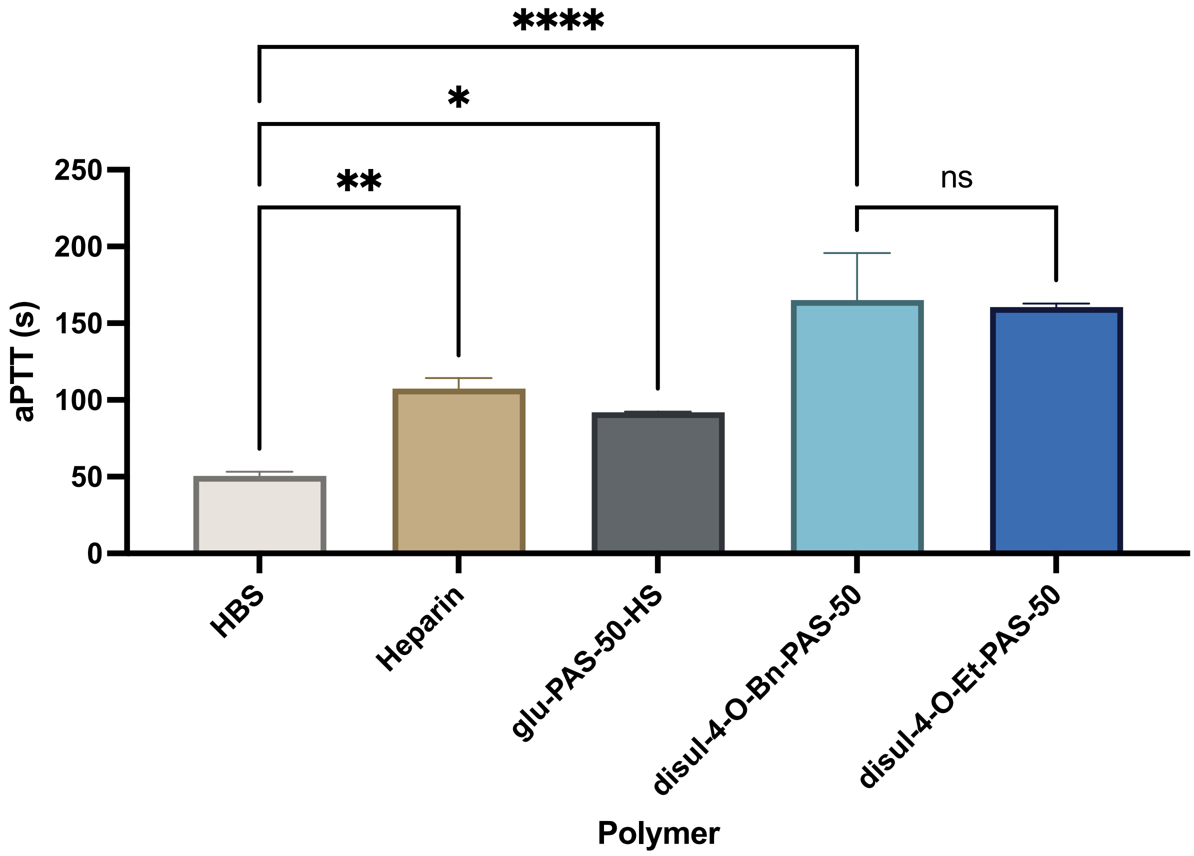


**Figure S5.** IR spectra of **3,6-di-*O*-PMB-4-*O*-Et-PAS.**

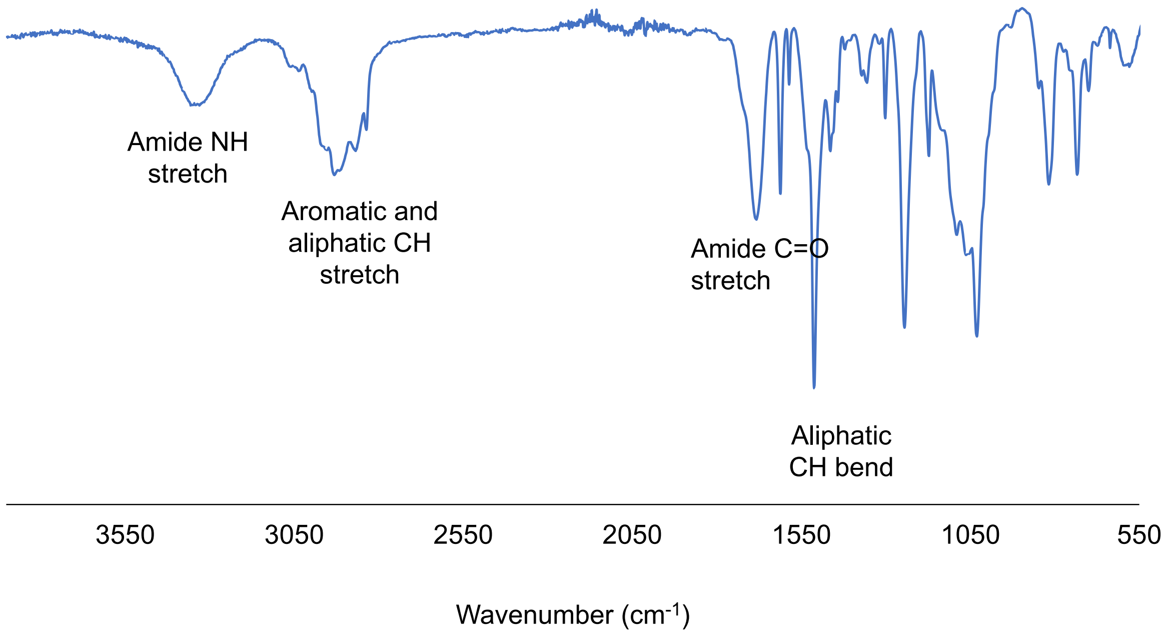

**Figure S6**. Evaluation of degree of polymerization of **disulPASs** by $^1$H NMR; light blue dots highlight initiator peak, dark blue dots highlight polymer $CH_3$ of 4-*O*-ethyl. **A**) N=5; **B**) N=12; **C**) N=25; **D**) N=50; **E**) N=70; **F**) N=100; **G**) N=120.

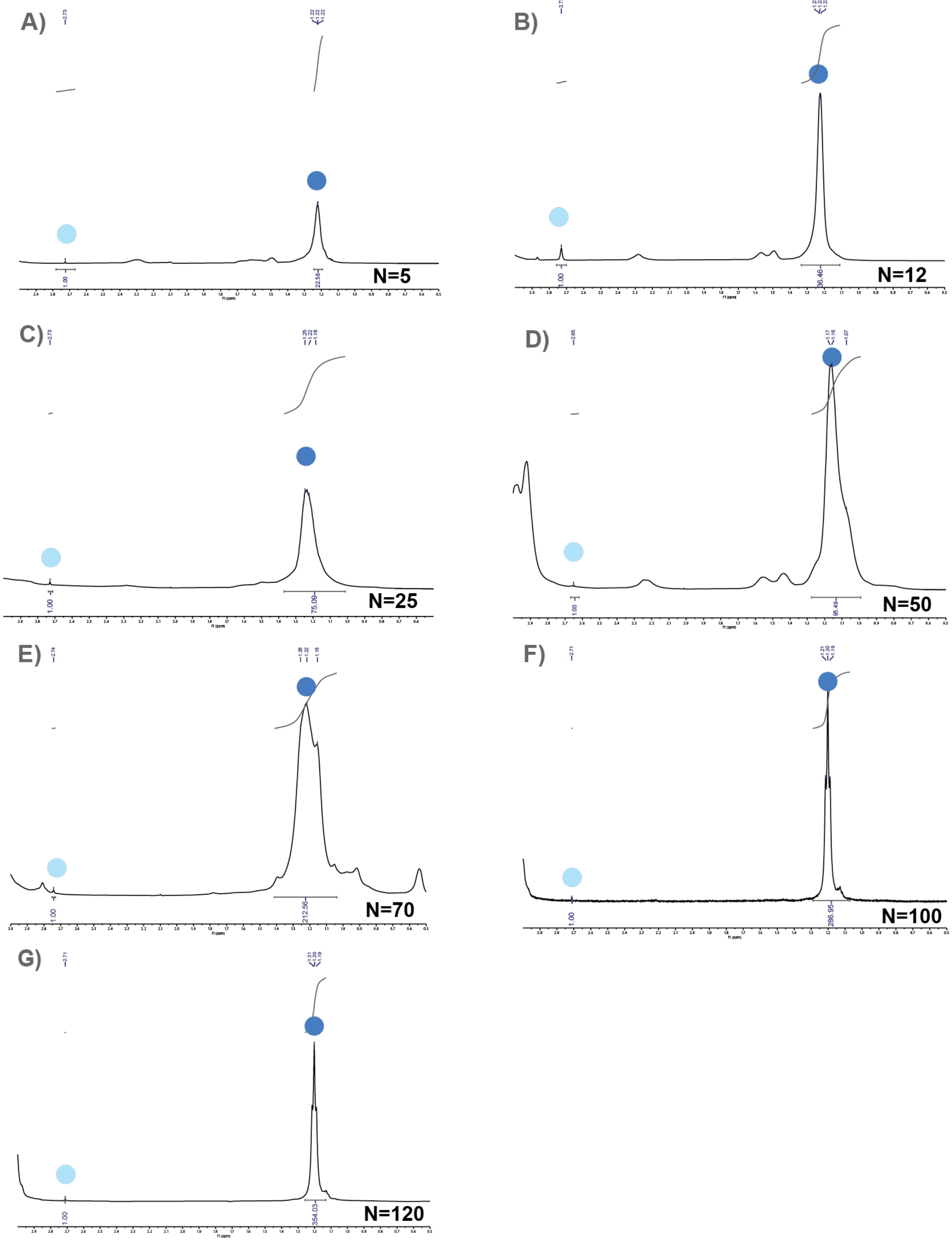

**Figure S7.** GPC chromatogram of **3,6-disulfate-4-*O*-Et-PAS** *before* and *after* NaOH treatment.

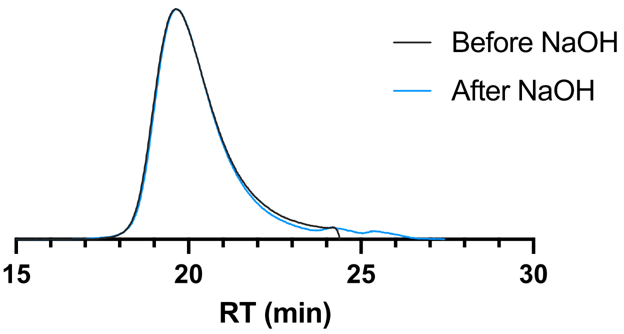


**Figure S8**. CD spectra of **3,6-disulfate-4-*O*-Et-PAS; A)** *before* and *after* NaOH treatment; **B)** After NaOH treatment and at varying temperatures; **C)** After NaOH treatment and at varying pH.

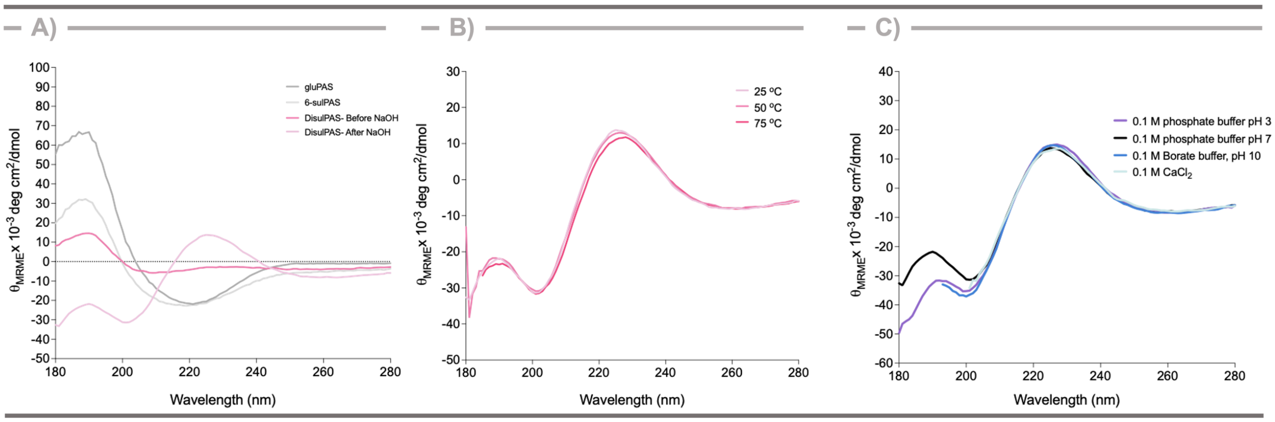


**Figure S9**. IR spectra of **3,6-di-sulfate-4-*O*-Et-PAS** *after* NaOH treatment.

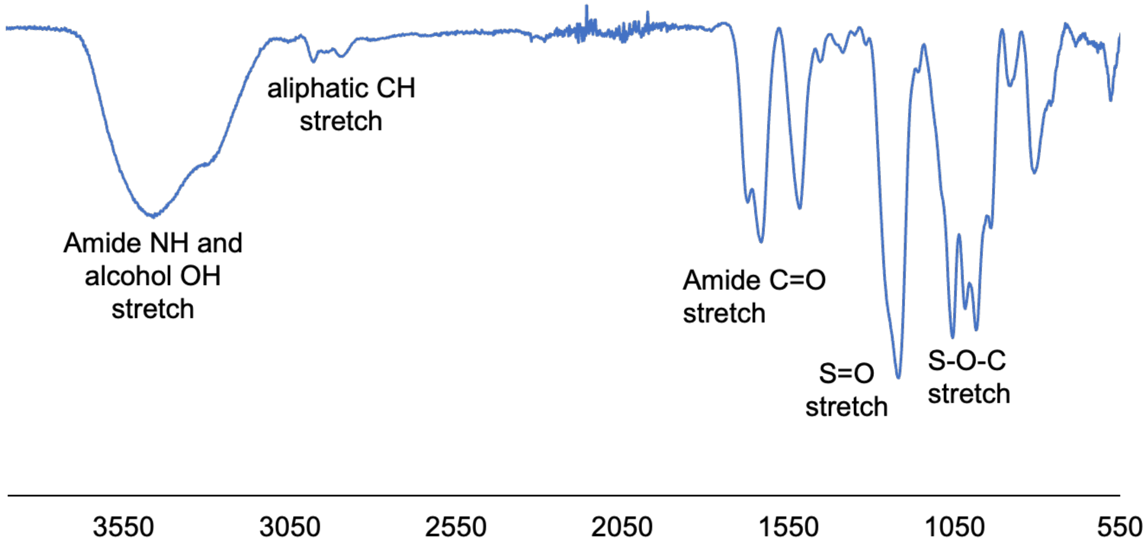

**Figure S10.** Plasma clotting time (aPTT) screening after: **A**) Intravenous administration of **disulPAS-70** (0.75 - 1.5 μg/g per mouse, n ≥ 3 mice/time point) or **B**) subcutaneous administration of varying amounts of **disulPAS-70** (1.5 - 7.5 μg/g per mouse, n ≥ 5 mice/time point). Data represent mean ± SEM.

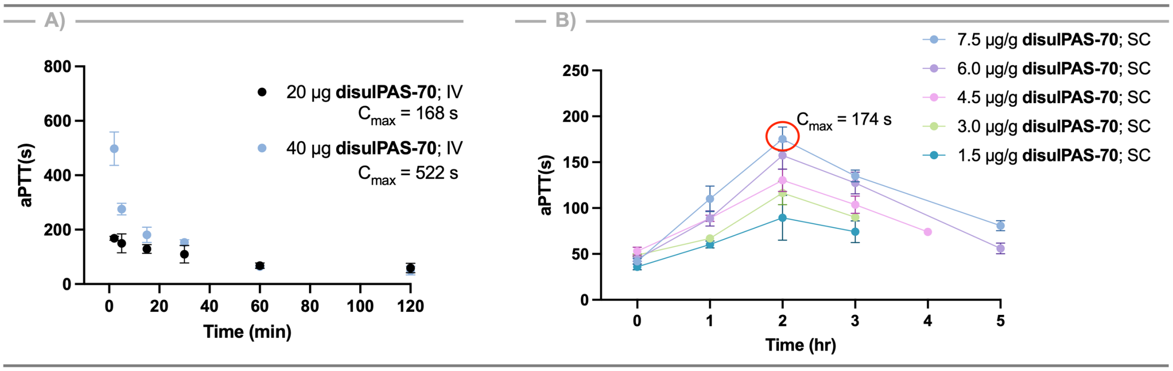

**General Methods and Materials**

D-glucal was purchased from Carbosynth, LLC (San Diego, California). All other reagents were purchased from Millipore Sigma (Burlington, MA), TCI (Portland, OR), or Chem-Impex (Wood Dale, Illinois). Solvents were purchased from Thermo Fisher Scientific (Waltham, MA). Pooled normal human plasma, APTT-XL and PT reagent were purchased from Thermo Fisher Scientific (Waltham, MA). Chromogenix Coatest® Heparin kit, Bovine thrombin (FIIa), FIIa substrate (Chromogenix S-2238™), FXIa substrate (Chromogenix S-2366™) and FXIIa substrate (Chromogenix S-2302™) were purchased from Diapharma (West Chester, Ohio). FXIIa was purchased from Biovision (Milpitas, California). Human Coagulation Factor XIa was purchased from Haematologic Technologies, Inc. (Essex Junction, VT). FIX (11503-H08H) and AT-III (10142-H08H) were purchased from SinoBiological (Wayne, PA). FVIIa was purchased from Sigma Aldrich (Burlington, MA). Aniara Diagnostica Biophen FVIIa and FIXa Assay kits were purchased from Thermo Fisher Scientific (Waltham, MA). Recombinant Human Active Heparanase/HPSE Protein, CF were purchased from R&D Systems, Inc. (Minneapolis, MN). Reactions were monitored by thin-layer chromatography (TLC) analysis and stained by potassium permanganate. Single Donor Human Whole Blood (sodium citrate anticoagulant) was purchase from Innovative Research (Novi, MI). NMR spectra were obtained with compounds dissolved $CDCl_3$ (D, 99.8 % +0.03% V/V TMS), DMSO-D6 (D, 99%) or $D_2O$ (D, 99.9%). Chemical shifts (δ) are recorded in ppm, coupling constants (*J*) are reported in Hz. NMR spectra were recorded on a Varian INOVA 500 MHz spectrometer, Varian INOVA 400 MHz spectrometer, or 600 MHz Bruker NMR Spectrometer. Infrared spectroscopy (IR) was performed on a Nicolet FT-IR with a horizontal attenuated total reflectance (ATR) adapter plate. aPTT and PT absorbance measurements were recorded on a Beckman Coulter AD340 plate-reader at 405 nm. Unless otherwise noted, all reactions were performed under a nitrogen atmosphere using anhydrous solvents and oven-dried glassware and stir bars. Lyophilization was performed using a Virtis Benchtop 4K freeze dryer Model 4BT4K2L-105 at -40˚C.

**Size exclusion chromatography**

Polymer molecular weights were determined by size exclusion chromatography using an Agilent Infinity II 1260 instrument equipped with refractive index and dual angle light scattering detectors. Protected polymers were run through an Agilent PLgel MIXED-C 5 µm column (300 x 7.5 mm) fitted with a guard column with THF as the eluent at 1.0 mL/ min. Deprotected polymers were run through an Agilent PL aquagel-OH MIXED-M 8 µm column (300 x 7.5 mm) fitted with a guard column with aqueous buffer (0.5 M $NaNO_3$, 0.01 M $Na_2HPO_4$, pH 7.4) as the eluent at 0.7 mL/ min at 35˚C. Analysis using refractive index was performed using the Agilent GPC/SEC software 2.2. Polystyrene standards were used for organic GPC and poly(ethylene glycol) and pullulan standards were used for aqueous GPC.

**Circular dichroism**

For initial CD experiments, gluPAS-100, 6-sulPAS-100, or 3,6-disulfate-4-*O*-Et-PAS-70 (**disulPAS-70**) before or after NaOH treatment were dissolved in DI water at 0.2 mg/mL and measured through a 1 mm path length cuvette. For **disulPAS-70** secondary structure stability experiments, polymer (after NaOH treatment) was dissolved to 0.2 mg/mL in 0.1M buffer at varying pH and measured as described previously, or dissolved in DI water and heated to 25˚C, 50˚C or 75˚C and measured.

**Ex vivo aPTT and PT assays** [1,2] 25 µL of **disulPAS-70** (25 - 250 µg/mL) dissolved in HBS at the aforementioned concentrations was added to 50 µL of Normal Human Plasma to give final concentrations [in plasma] of 8.33 - 83.33 µg/mL. This was incubated for 5 min at 37˚C before addition of 50 µL of aPTT-XL reagent and incubation for another 3 min at ambient temperature, with shaking. After this time, 25 µL of 0.04 M $CaCl_2$ solution was added to induce clotting, and absorbance was measured immediately at 405 nm every 6 s for a total time of 90 min. As clotting occurs, the absorbance value increases; aPTT was calculated as the time (s) taken for the absorption to increase by half the maximum value using a non-linear sigmoidal fit on GraphPad Prism 8. Heparin solution (25 µL of 1 IU/mL= 5.5 µg/mL in HBS prior to addition to plasma) was used as a positive control; HBS was used as a negative control.

Prothrombin time was evaluated by adding 100 µL of PT reagent (Thromboplastin-D) to a mixture of Normal Human Plasma (50 µL) and the analyte (25 µL of varying concentrations of **disulPAS-70** noted above, in HBS), pre-incubated at 37˚C (5 min), followed by measuring absorbance of all the samples at 405 nm every 6 s for a total time of 6 min. Clotting time was calculated from the absorbance graph as mentioned above for aPTT, and the same controls were used.

**General amidolytic activity inhibition assays**
All assays were measured kinetically, and the slope of time vs absorbance was calculated to give: $A_i$, for varying concentrations of analyte, $A_0$, for no analyte (positive control), and $A_b$, for when the coagulation factor was completely inhibited by 2% citric acid or 20% acetic acid prior to addition of substrate (negative control). Percent inhibition was calculated as follows:

$$\%\,\boldsymbol{inhibition} = \frac{\boldsymbol{Ao - Ai}}{\boldsymbol{Ao - Ab}} * \mathbf{100}$$

Each assay was repeated with the addition of buffer in place of AT-III for "without AT-III" data points.

***FXa amidolytic activity inhibition assay:*** Reagents in Chromogenix Coatest® Heparin kit and Bovine thrombin were used for the FXa amidolytic activity inhibition assay. A mixture 10 µL of AT-III (0.8 µM) and 10 µL of varying concentrations (0.1-10000 µg/mL) of analyte (**disulPAS-70**) was incubated at 37˚C for 5 min. Then, 10 µL of FXa (80 nM) was added, mixed, and incubated at 37˚C for 1 min. Last, 20 µL of S-2222 (1 mM) was added and the absorbance at 405 nm was measured every 60 s for 4 min. FXa was completely inhibited by 10 µL of 20% acetic acid prior to addition of substrate.

***FIIa amidolytic activity inhibition assay:*** Reagents in Chromogenix Coatest® Heparin kit and Bovine thrombin were used for the FIIa amidolytic activity inhibition assay.

Initially, 10 µL of AT-III (0.8 µM) and 10 µL of varying concentrations (0.1- 10000 µg/mL) of **disulPAS-70** were incubated together at 37˚C for 5 min. 10 µL of FIIa (80 nM) was then added, mixed well, and incubated at 37˚C for 1 min. Next, 20 µL of S-2238 (1 mM) was added and the absorbance at 405 nm was measured every 60 s for 4 min. FIIa was completely inhibited by 10 µL of 20% acetic acid prior to addition of substrate.

***FVIIa amidolytic activity inhibition assay:*** Reagents in Aniara Diagnostica LLC Biophen Factor VIIa Assay kit were used. Appropriate concentration of FVIIa was determined via a standard titration from 0 - 200 nM protein under normal assay conditions. Briefly, 10 µL of FVIIa and 10 µL of analyte were mixed to form final concentrations of 20 nM FVIIa and 0.1 - 1000 µg/mL **disulPAS**; 20 µL of AT-III (0.8 µM) was added and the assay plate was incubated for 5 min at 37˚C. 20 µL of R2 (Thromboplastin Calcium) and 40 µL of R1 (Factor X) were then added and the plate was incubated for 7 min at 37˚C with brief shaking. 40 µL of R3 (substrate SXa-II) was added and the absorbance of each well was read every 10 sec for 6 min at 405 nm. Complete inhibition was achieved with 20 µL of 2% citric acid prior to addition of R3.

***FIXa amidolytic activity inhibition assay:*** In buffer of 0.02 M Tris-HCl pH 7.5, 0.1 M NaCl, 0.1 mg/mL BSA, 33% ethylene glycol, and 5 mM $CaCl_2$, FIX was activated by incubating 500 µg/mL with 30 µg/mL Thermolysin (Sigma) for 1 hr at 37˚C. 10 mM (final) of 1,10 Phenanthroline was added and reaction was incubated at ambient temperature for 5 min to stop the activation. FIXa was then diluted to 50 nM and used without purification. For the chromogenic assay, reagents in Aniara Diagnostica LLC Biophen Factor IXa Assay kit were used. Briefly, 25 µL of FIXa with (0.1 - 1000 µg/mL) **disulPAS-70** was added to the plate and mixed with 25 µL of R1 (FX:FVIII:C) and 25 µL of AT-III (0.8 µM); this was incubated at 37˚C for 2 min. 25 µL of R2 (Activator reagent) was then added followed by another incubation at 37˚C for 3 min. Finally, R3 (substrate) was added, and absorbance was read at 405 nm every 20 s for 5 min. Complete inhibition was achieved with 25 µL of 2% citric acid prior to addition of R3.

***FXIa amidolytic activity inhibition assay:*** 20 µL of varying concentration (1.95 - 5000 µg/mL) of **disulPAS-70** was prepared, mixed with 20 µL of AT-III (0.8 µM) and made up to 60 µL by buffer. 20 µL of FXIa (2.5 µg/mL, 16 nM) was added to the solution and incubated at 37˚C for 5 min. Next, 20 µL of S-2366 (250 µM) was added and the absorbance at 405 nm was measured every 60 s for 5 min. FXIa was completely inhibited by 10 µL of 20% acetic acid prior to addition of substrate.

***FXIIa amidolytic activity inhibition assay:*** 20 µL of varying concentration of **disulPAS-70** (0.05 - 500 µg/mL) was mixed with 20 µL of FXIIa (80 nM) and 20 µL of AT-III (0.8 µM) and incubated at 37˚C for 5 min. Then, 20 µL of S-2302 (16 mM) was added and the absorbance at 405 nm was measured every 60 s for 5 min. FXIIa was completely inhibited by 10 µL of 20% acetic acid prior to addition of substrate.

**Cytotoxicity**

NIH 3T3 fibroblast cells were obtained from ATCC (Manassas, VA) and all experiments were conducted on cells between P3 and P10. Cells were cultured in DMEM media

(Invitrogen, Carlsbad, CA) supplemented with 10% bovine calf serum (BCS) and 1x Penicillin / Streptomycin (PS) in a 37˚C incubator with 5% $CO_2$. For MTS experiments, cells were cultured for one day at 5000 cells/well in 96 well plates. The media was then removed and 150 µL of a 3 mg/mL solution of polymer (**disulPAS-70**, gluPAS-100, heparin or dextran) dissolved in media and filtered through a 0.22 µm filter was added to column 1 and 100 µL of media alone was added to columns 2-12. 50 µL of the solution from column 1 was serially diluted (1:3 dilution) across columns 2-10 and the final 50 µL was discarded. The cells were incubated in these polymer solutions for 24 hr, followed by aspiration of the media and the addition of a 6:1 solution of media to MTS reagent (Promega, Madison, WI). The plates were then incubated at 37˚C for 3 hr, and absorbance was read at 492 nm. Cells exposed to media alone (n=8) were considered 100% proliferative, and wells containing no cells (n=8) were considered 0% proliferative. The absorbance of each experimental well (n=8) was normalized to this range.

**Hemolytic assay**
[3] Single donor human whole blood was purchased from Innovative Research (Novi, MI). 2 mL whole blood was centrifuged at 1500 rpm for 10 min at 4˚C. The plasma supernatant was removed, and erythrocytes were resuspended in ice cold PBS. Cells were centrifuged at 1500 rpm for 10 min at 4˚C. This was repeated two more times to ensure the removal of any released hemoglobin. After the supernatant was removed following the last wash, the cells were resuspended in PBS to get a 20% v/v RBC solution. 100 µL of varying concentrations (1 - 0.125 mg/mL) of the polymer or heparin prepared in PBS were incubated with 100 µL of the 20% v/v RBC solution in a 96-well plate for 24 hr at 37˚C. Complete hemolysis was attained using a 2% v/v Triton-X yielding the 100% control value (positive control). After incubation, the plates were centrifuged, and the supernatants were transferred to a new 96-well plate. The release of hemoglobin was determined by spectrophotometric analysis of the supernatant at 550 nm. Results were expressed as the amount of hemoglobin release induced by the conjugates as a percentage of the total.

**Reversal of anticoagulant activity of disulPAS by Protamine sulfate**
aPTT assay was repeated with 12.5 µL of **disulPAS-70** (25 µg/mL) and 12.5 µL of varying concentrations (15.6 - 125 µg/mL) of Protamine Sulfate (PS) in place of the original assay's 25 µL analyte. The concentration of PS required to bring the aPTT down to negative control was estimated.

**Evaluation of degradation of disulPAS by heparanase**
[2] Heparin (200 µg for each trial) or **disulPAS-70** (200 µg for each trial) were incubated for 24 hr at 37˚C with (n = 3) or without (n = 1) Recombinant Human Active Heparanase/ HPSE Protein (550 ng for each trial) in a final volume of 200 µL reaction solution (0.15 M NaCl, 20 mM phosphate-citrate buffer, pH 5.8, 1 mM dithiothreitol (DTT), 1 mM CaCl2). After 24 hr, the samples were lyophilized, re-dissolved in SEC buffer and retention times were evaluated.

#### In vivo studies

##### Animals

C57BL/6 mice (25-30 g, 9 weeks old, male, Jackson Laboratory, Bar Harbor, ME) were used for all animal experiments. Protocols were approved by the Institutional Animal Care and Use Committee of Beth Israel Deaconess Medical Center.

##### Pharmacokinetic/pharmacodynamic analysis with aPTT assay

For pharmacokinetic analysis with intravenous (IV) administration, mice were anesthetized with an intraperitoneal injection of ketamine and xylazine. Mice received **disulPAS-70** (0.75 µg/g in 50 µL of saline) IV, via the penile vein. Two minutes after injection, blood samples were collected from the mice via cardiac puncture into a citrated tube (10% v/v). For pharmacokinetic analysis following subcutaneous (SC) administration, blood samples were collected into microtubes containing citrate (10% v/v) by cheek puncture from the submandibular vein at 1, 2, 3, and 5 hours after SC administration of **disulPAS-70** (1.5 - 7.5 µg/g in 100 µL of saline). Plasma was obtained by centrifuging at 800 g for 10 minutes at room temperature. The plasma was either used immediately or stored at -80˚C. For the aPTT assay, 50 µL of mouse plasma was pre-incubated at 37˚C for 5 minutes. Afterward, 50 µL of APTT-XL reagent was added, and the mixture was incubated at room temperature with shaking for 5 minutes. Subsequently, 25 µL of 25 mM $CaCl_2$ solution was added to trigger clotting, and the absorbance at 405 nm was measured every 6 seconds for up to 20 minutes at 37°C; absorbance increased as clotting occurred. The aPTT was measured as the time to half-maximum absorbance and plotted using GraphPad Prism 11 (GraphPad Software, San Diego, CA, USA).

##### *In vivo* evaluation of anti-thrombotic efficacy with a non-occlusive venous thrombosis murine model

The prophylactic efficacy of **disulPAS-70** was evaluated in a preclinical mouse model of non-occlusive venous thrombosis.[4] Polymer was administered (7.5 µg/g in 100 µL saline, SC, n=6) to male C57BL/6 mice 2 hours prior to electrolytic injury and every 12 hours thereafter until termination. Unfractionated heparin was administered (0.11 U/g, SC, n=6) as a clinically relevant reference for prophylactic dosing, 2 hours prior to electrolytic injury and every 12 hours until termination. An additional control group was enrolled with saline administered SC (n=6).

Non-occlusive venous thrombosis was induced by electrolytic injury of the inferior vena cava. Mice were anesthetized with 2% isoflurane, and the inferior vena cava was approached via a midline laparotomy. Venous side branches were ligated or cauterized, while posterior branches were left patent. A 25-gauge stainless steel needle, attached to a silver-coated copper wire, was inserted into the exposed caudal vena cava and positioned against the anterior wall (anode). A second wire was implanted subcutaneously to complete the circuit (cathode), and a 250 µA current was applied for 15 min. Subsequently, the needle was removed, and a cotton swab was gently applied to the puncture site to prevent bleeding. The vena cava and associated thrombus, immediately below the renal veins to just above the bifurcation, was excised 48 hours after initial drug administration. Reported thrombus weight is the total vessel weight of the excised IVC with thrombus minus the mean vessel weight (n=12) of healthy IVC. %

reduction in thrombus weight is reported relative to thrombus weight measured in saline control mice.

**Bleeding time measurement**
The effect of **disulPAS-70** on bleeding time was evaluated using a murine tail-vein transection model, administered either IV or SC (n=8, or n=6 mice per group, respectively). C57BL/6 mice were anesthetized with an intraperitoneal injection of ketamine HCl (Zoetis, 125 mg/kg) and xylazine (Akorn Inc., 12.5 mg/kg). Two minutes after IV injection of saline, **disulPAS-70** (0.75 μg/g), or heparin (0.004 U/g), the tail vein was transected 10 mm from the tip of the tail. The tail was immediately submerged vertically in 50 mL warm saline (37˚C) with the tip 2 cm below the body horizon. Bleeding time was defined as the time required for bleeding to stop for 30 seconds after injury. For SC administration groups, saline, **disulPAS-70** (7.5 μg/g), or heparin (0.11 U/g) was administered via intraperitoneal injection 2 hours before the tail-vein bleeding test. Following anesthesia, the tail-vein bleeding time for each mouse was measured as described above.

**Statistical Analysis**
Data are presented as mean ± SEM unless otherwise noted. Group comparisons were conducted using an unpaired Student's t-test. Significance was indicated as *$p \leq 0.05$, **$p \leq 0.01$, and ***$p \leq 0.001$; p-values less than 0.05 were considered statistically significant.

#### Small Molecule Synthetic Procedures

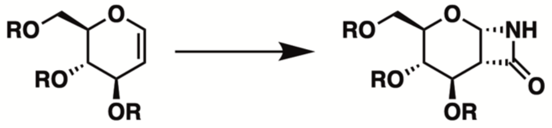


**General procedure for CSI cycloaddition**
Protocol was adapted from literature with minor modifications.[5] At ambient temperature, chlorosulfonyl isocyanate (0.22 mL, 2.52 mmol, 1.4 eq), which had previously been stored over oven-dried sodium carbonate for > 1 week at 4˚C, was dissolved in 2.8 mL of anhydrous toluene over oven-dried sodium carbonate (0.32 g, 3.0 eq) and under nitrogen. The solution was cooled to -78˚C in an acetone/ dry ice bath, and a solution of compound 4 (0.89 g, 1.8 mmol, 1.0 eq) dissolved in 2.8 mL of anhydrous toluene was added dropwise. The solution was warmed to -55˚C by transferring the reaction flask to a chloroform/ dry ice cooling bath. After 3 hr at -55˚C, the reaction was cooled to -78˚C and diluted dropwise with an additional 8.4 mL of anhydrous toluene followed by the dropwise addition of Red-Al (>65 wt% solution in toluene; 0.95 mL, 5.27 mmol, 2.9 eq) via syringe pump. After 30 min at -78˚C, the reaction was warmed to -10˚C in an ice/ brine bath for 5-10 min. After this time, 1.0 mL of water was added to quench the reaction, and it was stirred for an additional 30 min while warming to 0˚C. If possible, the solution was filtered to remove solids. If an emulsion formed, a saturated solution of

tartrate could be added to complex aluminum ions. After the addition of 50 mL of diethyl ether, the organic phase was washed with saturated bicarbonate and brine, dried over Na2SO4, and the solvent was removed. The crude material was purified with silica gel chromatography (40% EtOAc in cyclohexane) to afford a white solid (0.97 g, 57%).

**General procedure for TCAI cycloaddition**

The procedure was adapted from the literature with minor changes.[6] Compound 7 (1.0 g, 1.84 mmol, 1.0 eq), was dissolved to 1 M in MeCN (1.8 mL) and cooled to 0˚C. Trichloroacetyl isocyanate (2.5 eq, 0.57 mL, 4.6 mmol, 2.56 eq) was added dropwise and the reaction mixture was allowed to warm to ambient temperature. The reaction was monitored by TLC (permanganate stain) for 3 days or until the starting material was completely consumed. The reaction mixture was then cooled to -20˚C with a water/ methanol/ dry ice cooling bath, and benzylamine (0.55 mL, 5.06 mmol, 2.81 eq) dissolved in 3.6 mL of anhydrous acetonitrile was added dropwise by syringe pump over the course of 1 hr. The reaction mixture was then allowed to warm to ambient temperature while monitoring; once its temperature reached ~0˚C, a white precipitate was formed in the yellow solution. The mixture was filtered to remove the white precipitate and the flask and filtered solid were quickly washed with hexanes. The filtrate was then poured into 6 mL of 0.1 M HCl and extracted 4 times with EtOAc. The combined organic extracts were washed 2 more times with 0.1 M HCl, followed by brine, and dried over sodium sulfate before evaporating the solvent. The crude product was purified by column chromatography (40% EtOAc in cyclohexane) to yield monomer 3 as a pale foam (0.38 g, 35%).

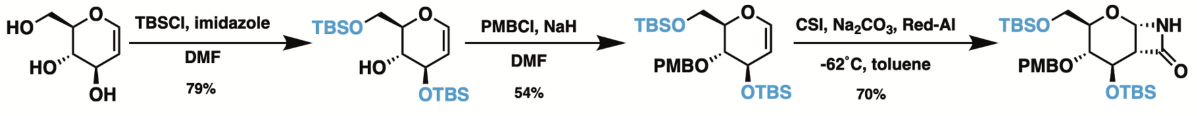


**Scheme S1**: Synthesis of monomer **1**

***Monomer 1***

Compound **1**: Protocol was adapted from literature with minor changes.[7] To a 0˚C solution of D-glucal (5.0 g, 34.2 mmol, 1.0 eq) and imidazole (11.9 g, 175.5 mmol, 5.13 eq) in anhydrous DMF (0.2 M; 180 mL) under $N_2$ protection, tert-Butyldimethylsilyl chloride (TBSCl) (10.98 g, 72.9 mmol, 2.13 eq) was added. The reaction was stirred at 0˚C for 4 hr, after which time it was placed under high vacuum for 30 min at ambient temperature. Reaction was then transferred to a separatory funnel containing 5x reaction volumes of water and the solution was extracted 3x with EtOAc before the combined organic layer was washed with brine and dried over sodium sulfate. Product was purified via column chromatography (5% EtOAc in cyclohexane) (12.8 g, 79%). **HRMS (m/z):** $[M^+Na]^+$ calcd. for $C_{18}H_{38}O_4Si_2$ Na, 397.2206; found, 397.2200. **$^1$H NMR** (600 MHz, $CDCl_3$) δ 6.27 (d, *J* = 6.1 Hz, 1H, H-1), 4.62 (dd, *J* = 6.1, 2.4 Hz, 1H, H-2), 4.25 – 4.20 (m, 1H, H-3), 3.97 (dd, *J* = 11.3, 4.9 Hz, 1H, H-6), 3.88 (dd, *J* = 11.3, 3.8 Hz, 1H, H-6'), 3.85 – 3.80 (m, 1H, H-5), 3.79 – 3.72 (m, 1H, H-4), 2.50 (d, *J* = 3.6 Hz, 1H, -O*H*), 0.91 (d, *J* = 2.9 Hz, 18H, $Si(CH_3)_2$ *i*-Pr), 0.10 (dd, *J* = 19.1, 1.7 Hz, 12H, $Si(CH_3)_2$ *i*-Pr) ppm; **$^{13}$C NMR** (151 MHz, CDCl3) δ 143.3 (C-1), 103.5 (C-2), 77.9 (C-5),

71.1 (C-4), 69.9 (C-3), 63.2 (C-6), 25.94 (SiC(*C*$H_3$)$_3$), 25.86 (SiC(*C*$H_3$)$_3$), 25.84 (SiC(*C*$H_3$)$_3$), 18.4 (Si*C*($CH_3$)$_3$), 18.2 (Si*C*($CH_3$)$_3$), -4.4 (Si(*C*$H_3$)$_2$), -4.5 (Si(*C*$H_3$)$_2$), -5.3 (Si(*C*$H_3$)$_2$), -5.4 (Si(*C*$H_3$)$_2$) ppm.

Compound **2** was prepared following reported procedures.[7,8] Briefly, compound **1** (5.5 g, 14.7 mmol, 1.0 eq) was dissolved in anhydrous DMF (0.5 M; 30 mL) under $N_2$ and cooled to 0˚C, followed by the portionwise addition of NaH (0.71 g, 29.4 mmol, 2.0 eq). After stirring for 30 min at 0˚C, *para*-methoxybenzyl chloride (PMBCl) (2.43 mL, 2.81 g, 17.93 mmol, 1.22 eq) was added dropwise and then stirred for 2 hr at ambient temperature. After this time, excess NaH was quenched with MeOH before the reaction was poured into an equal volume of ice water and extracted with EtOAc 2x. The combined organic layer was washed with brine and dried over sodium sulfate. Product was purified via column chromatography (4% EtOAc in cyclohexane) to afford a clear oil (3.9 g, 54%). **HRMS (m/z):** [$M^+$H]$^+$ calcd for: $C_{26}H_{46}O_5Si_2$ H, 495.1999; found, 495.1939. **$^1$H NMR** (600 MHz, $CDCl_3$) δ 7.27 (d, *J* = 8.6 Hz, 2H, $H_{Ar}$), 6.87 (d, *J* = 8.7 Hz, 2H, $H_{Ar}$), 6.29 (dd, *J* = 6.1, 1.4 Hz, 1H, H-1), 4.74 (d, *J* = 10.8 Hz, 1H, OC*H*$_2$Ph), 4.64 (d, *J* = 10.8 Hz, 1H, OC*H*$_2$Ph), 4.62 (dd, *J* = 6.1, 2.7 Hz, 1H, H-2), 4.33 – 4.29 (m, 1H, H-3), 3.94 (dd, *J* = 11.4, 4.7 Hz, 1H, H-6), 3.87 (ddd, *J* = 7.5, 4.6, 2.4 Hz, 1H, H-5), 3.82 (dd, *J* = 11.4, 2.3 Hz, 1H, H-6'), 3.80 (s, 3H, PhOC*H*$_3$), 3.62 (dd, *J* = 8.2, 6.1 Hz, 1H, H-4), 0.91 (d, *J* = 8.7 Hz, 18H, Si($CH_3$)$_2$ *i*-Pr), 0.12 – 0.06 (m, 12H, Si(C*H*$_3$)$_2$ *i*-Pr) ppm; **$^{13}$C NMR** (151 MHz, $CDCl_3$) δ 159.3 ($C_{Ar}$), 143.4 (C-1), 129.6 ($C_{Ar}$), 113.8 ($C_{Ar}$), 103.3 (C-2), 78.2 (C-5), 76.2 (C-4), 73.6 (O*C*$H_2$Ph), 69.0 (C-3), 62.0 (C-6), 55.3 (PhO*C*$H_3$), 26.00 (SiC(*C*$H_3$)$_3$), 25.98 (SiC(*C*$H_3$)$_3$), 25.86 (SiC(*C*$H_3$)$_3$), 18.4 (Si*C*($CH_3$)$_3$), 18.1 (Si*C*($CH_3$)$_3$), -4.4 (Si(*C*$H_3$)$_2$), -4.6 (Si(*C*$H_3$)$_2$), -5.1 (Si(*C*$H_3$)$_2$), -5.3 (Si(*C*$H_3$)$_2$) ppm.

**Monomer 1** was synthesized from compound **2** using the CSI cycloaddition reaction. Product was purified via column chromatography (30% EtOAc in cyclohexane) to afford a white solid (70%). **HRMS (m/z):** [$M^+$Na]$^+$ calcd. for $C_{27}H_{47}NO_6Si_2$ Na, 560.2840; found, 560.2852. **$^1$H NMR** (500 MHz, $CDCl_3$) δ 7.24 (d, *J* = 8.6 Hz, 2H, $H_{Ar}$), 6.86 (d, *J* = 8.6 Hz, 2H, $H_{Ar}$), 6.28 (s, 1H, N*H*), 5.43 (d, *J* = 4.5 Hz, 1H, H-1), 4.67 (d, *J* = 10.9 Hz, 1H, OC*H*$_2$Ph), 4.56 (d, *J* = 10.9 Hz, 1H, OC*H*$_2$Ph), 4.31 (dd, *J* = 6.1, 2.9 Hz, 1H, H-3), 3.85 – 3.81 (m, 1H, H-5), 3.79 (s, 4H, PhOC*H*$_3$, H-6), 3.70 (dd, *J* = 10.6, 3.7 Hz, 1H, H-6'), 3.45 (t, *J* = 6.5 Hz, 1H, H-4), 3.26 – 3.12 (m, 1H, H-2), 0.90 (d, *J* = 5.1 Hz, 18H, Si($CH_3$)$_2$ *i*-Pr), 0.18 – 0.01 (m, 12H, Si(C*H*$_3$)$_2$ *i*-Pr) ppm; **$^{13}$C NMR** (126 MHz, $CDCl_3$) δ 166.0 (C=O), 159.2 ($C_{Ar}$), 130.4 ($C_{Ar}$), 129.3 ($C_{Ar}$), 113.8 ($C_{Ar}$), 77.2 (C-4), 76.0 (C-1), 73.2 (O*C*$H_2$Ph), 72.1 (C-5), 69.1 (C-3), 63.4 (C-6), 57.1 (PhO*C*$H_3$), 55.2 (C-2), 25.9 (SiC(*C*$H_3$)$_3$), 25.8 (SiC(*C*$H_3$)$_3$), 18.4 (Si*C*($CH_3$)$_3$), 17.8 (Si*C*($CH_3$)$_3$), -4.7(Si(*C*$H_3$)$_2$), -4.8 (Si(*C*$H_3$)$_2$), -5.2 (Si(*C*$H_3$)$_2$), -5.3 (Si(*C*$H_3$)$_2$) ppm.

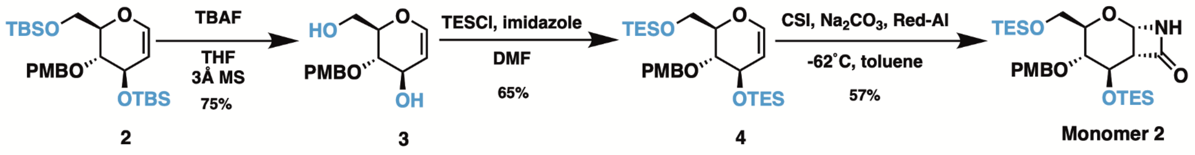


**Scheme S2**: Synthesis of monomer **2**.

***Monomer 2***

The protocol to yield compound **3** was adapted from literature with modifications.[9,10] Briefly, a flask containing only dry compound **2** (4.94 g, 10.0 mmol, 1.0 eq) and 3Å molecular sieves was cooled to 0˚C under $N_2$, followed by the dropwise addition of TBAF (1 M solution in THF; 40 mL, 40 mmol, 4.0 eq). After stirring overnight at ambient temperature, the reaction was filtered and solvent was removed before purifying via column chromatography (60% EtOAc in cyclohexane) to afford a white solid (2.0 g, 75%). **HRMS (m/z):** $[M^{-}H]^{-}$ calcd. for $C_{14}H_{17}O_5$ 265.1092; found, 265.1076. **$^1$H NMR** (600 MHz, $CDCl_3$) δ 7.31 (d, *J* = 8.6 Hz, 2H, $H_{Ar}$), 6.90 (d, *J* = 8.6 Hz, 2H, $H_{Ar}$), 6.35 (dd, *J* = 6.0, 1.7 Hz, 1H, H-1), 4.75 (s, 2H, O$CH_2$Ph), 4.73 (dd, *J* = 6.0, 2.5 Hz, 1H, H-2), 4.39 – 4.30 (m, 1H, H-3), 3.96 – 3.91 (m, 1H, H-6), 3.89 – 3.82 (m, 2H, H-5, H-6'), 3.81 (s, 3H, PhO$CH_3$), 3.61 (dd, *J* = 9.4, 6.8 Hz, 1H, H-4), 1.88 (dd, *J* = 8.2, 5.1 Hz, 1H, 6-O*H*), 1.79 (d, *J* = 5.7 Hz, 1H, 3-O*H*) ppm; **$^{13}$C NMR** (151 MHz, $CDCl_3$) δ 144.2 (C-1), 129.8 ($C_{Ar}$), 114.1 ($C_{Ar}$), 103.3 (C-2), 77.4 (C-5), 76.7 (C-4), 73.6 (O$CH_2$Ph), 69.5 (C-3), 61.9 (C-6), 55.3 (PhO$CH_3$) ppm.

Compound **4**: Protocol was adapted from literature with modifications.[7] 0.18 g (0.676 mmol, 1.0 eq) of compound **3** and 0.23 g of imidazole (3.38 mmol, 5.0 eq) were dissolved in anhydrous DMF (0.34 M; 2 mL) under $N_2$ and cooled to 0˚C. Next, chlorotriethylsilane (TESCl) (0.25 mL, 1.48 mmol, 2.2 eq) was added dropwise and the reaction was warmed to ambient temperature and stirred overnight. After completion of reaction (confirmed by TLC), the reaction mixture was transferred to a separatory funnel containing 10 mL of DI water, and the product was extracted with diethyl ether 5x. The combined organic layer was washed with brine, dried over sodium sulfate, and the solvent was evaporated. The crude product was purified using column chromatography (4% EtOAc in cyclohexane) to yield a colorless oil (0.22 g, 65%). **HRMS (m/z):** $[M^{+}Na]^{+}$ calcd. for $C_{26}H_{46}O_5Si_2$ Na, 517.2782; found, 517.2789. **$^1$H NMR** (600 MHz, $CDCl_3$) δ 7.27 (d, *J* = 8.6 Hz, 2H, $H_{Ar}$), 6.88 (d, *J* = 8.6 Hz, 2H, $H_{Ar}$), 6.31 (dd, *J* = 6.1, 1.4 Hz, 1H, H-1), 4.76 (d, *J* = 11.0 Hz, 1H, O$CH_2$Ph), 4.67 – 4.61 (m, 2H, O$CH_2$Ph, H-2), 4.33 (ddd, *J* = 6.0, 2.9, 1.4 Hz, 1H, H-3), 3.91 – 3.88 (m, 2H, H-5, H-6), 3.84 – 3.81 (m, 4H, H-6', PhO$CH_3$), 3.65 – 3.59 ( ap t, 1H, H-4), 0.97 (dt, *J* = 12.6, 7.9 Hz, 18H, $Si(CH_2CH_3)_3$), 0.62 (dq, *J* = 19.5, 8.2 Hz, 12H, $Si(CH_2CH_3)_3$) ppm; **$^{13}$C NMR** (151 MHz, $CDCl_3$) δ 159.3 ($C_{Ar}$), 143.5, 130.6 ($C_{Ar}$), 129.6 ($C_{Ar}$), 113.8 ($C_{Ar}$), 103.2 (C-2), 78.1 (C-5), 76.2 (C-4), 73.5 (O$CH_2$Ph), 68.9 (C-3), 61.8 (C-6), 55.3 (PhO$CH_3$), 6.9 ($Si(CH_2CH_3)_3$), 6.76 ($Si(CH_2CH_3)_3$), 6.75 ($Si(CH_2CH_3)_3$), 5.1 ($Si(CH_2CH_3)_3$), 4.5 ($Si(CH_2CH_3)_3$) ppm.

**Monomer 2:** A CSI cycloaddition was carried out on compound **4** and purified via column chromatography (30% EtOAc in cyclohexane) to obtain **monomer 2** as a colorless oil (57%). **HRMS (m/z):** $[M^{+}Na]^{+}$ calcd. for $C_{27}H_{47}NO_6Si_2$ Na, 560.2840; found, 560.2855. **$^1$H NMR** (500 MHz, CDCl3) δ 7.26 – 7.24 (ap d, 2H, $H_{Ar}$), 6.87 (d, *J* = 8.6 Hz, 2H, $H_{Ar}$), 6.03 (s, 1H, N*H*), 5.46 (d, *J* = 4.5 Hz, 1H, H-1), 4.68 (d, *J* = 11.1 Hz, 1H, O$CH_2$Ph), 4.54 (d, *J* = 11.1 Hz, 1H, O$CH_2$Ph), 4.33 (dd, *J* = 5.9, 2.9 Hz, 1H, H-3), 3.89 – 3.83 (m, 1H, H-5), 3.80 (s, 3H, PhO$CH_3$), 3.76 (dd, *J* = 10.8, 5.4 Hz, 1H, H-6), 3.69 (dd, *J* = 10.8, 4.0 Hz, 1H, H-6'), 3.45 (t, *J* = 6.4 Hz, 1H, H-4), 3.22 (d, *J* = 4.5 Hz, 1H, H-2), 1.00 – 0.92 (m, 18H, $Si(CH_2CH_3)_3$), 0.63 (dq, *J* = 32.4, 7.9 Hz, 12H, $Si(CH_2CH_3)_3$) ppm; **$^{13}$C NMR** (126 MHz, $CDCl_3$) δ 168.2 (C=O), 166.9 ($C_{Ar}$), 159.2 ($C_{Ar}$), 130.4 ($C_{Ar}$), 129.4

($C_{Ar}$), 113.7 ($C_{Ar}$), 77.0 (C-4), 75.9 (C-1), 73.1 (O*C*$H_2$Ph), 72.3 (C-5), 69.0 (C-3), 63.3 (C-6), 57.0 (C-2), 55.3 (PhO*C*$H_3$), 6.82 (Si($CH_2$*C*$H_3$)$_3$), 6.77 (Si($CH_2$*C*$H_3$)$_3$), 6.73 (Si($CH_2$*C*$H_3$)$_3$), 4.7 (Si(*C*$H_2CH_3$)$_3$), 4.4 (Si(*C*$H_2CH_3$)$_3$) ppm.

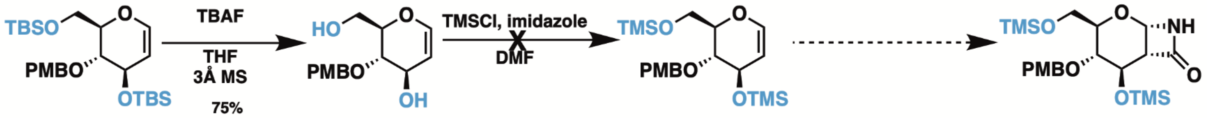


2 3 Monomer 3

**Scheme S3**: Synthesis of monomer **3**.

##### *Monomer 3*

Compounds **2** and **3** prepared as previously reported under **monomer 2**.
Synthesis of 3,6-di-*O*-TMS-4-*O*-PMB *D*-glucal was unsuccessful as TMS groups cleaved off during aqueous workup and glucal could not undergo cycloaddition.

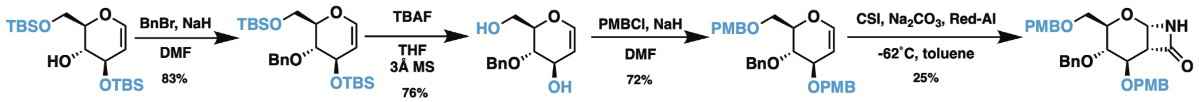


1 5 6 7 Monomer 4

**Scheme S4**: Synthesis of monomer **4**

##### *Monomer 4*

Compound **5** was synthesized as previously described for compound **2**,[9] but with benzyl bromide (BnBr) (0.175 mL, 1.47 mmol, 1.1 eq) in place of PMBCl; compound **1** (0.5 g, 1.33 mmol, 1.0 eq), NaH (0.04 g, 1.60 mmol, 1.2 eq), DMF (0.2 M; 6.7 mL), stirred overnight. Product eluted off column at 2% EtOAc in cyclohexane; pure product was a colorless oil (0.49 g, 79%). **HRMS (m/z):** $[M^{+}Na]^{+}$ calcd. for $C_{25}H_{44}O_4Si_2Na$, 487.2676; found, 487.2782. **$^{1}$H NMR** (600 MHz, $CDCl_3$) δ 7.35 (d, *J* = 5.2 Hz, 5H, $H_{Ar}$), 6.31 (dd, *J* = 6.1, 1.4 Hz, 1H, H-1), 4.82 (d, *J* = 11.2 Hz, 1H, OC*$H_2$*Ph), 4.73 (d, *J* = 11.2 Hz, 1H, OC*$H_2$*Ph), 4.64 (dd, *J* = 6.1, 2.7 Hz, 1H, H-2), 4.35 (dt, *J* = 6.3, 2.3 Hz, 1H, H-3), 3.96 (dd, *J* = 11.5, 4.7 Hz, 1H, H-6), 3.93 – 3.90 (m, 1H, H-5), 3.84 (dd, *J* = 11.4, 2.3 Hz, 1H, H-6’), 3.65 (dd, *J* = 8.2, 6.1 Hz, 1H, H-4), 0.91 (d, *J* = 6.5 Hz, 18H, Si($CH_3$)$_2$ *i*-Pr), 0.10 – 0.07 (m, 12H, Si(C*$H_3$*)$_2$ *i*-Pr) ppm; **$^{13}$C NMR** (151 MHz, $CDCl_3$) δ 143.5 (C-1), 129.0 ($C_{Ar}$), 128.8 ($C_{Ar}$), 128.4 ($C_{Ar}$), 127.9 ($C_{Ar}$), 127.7 ($C_{Ar}$), 103.3 (C-2), 78.1 (C-5), 76.6 (C-4), 73.9 (O*C*$H_2$Ph), 69.0 (C-3), 62.0 (C-6), 26.0 (SiC(*C*$H_3$)$_3$), 25.97 (SiC(*C*$H_3$)$_3$), 25.85 (SiC(*C*$H_3$)$_3$), 18.4 (Si*C*($CH_3$)$_3$), 18.1 (Si*C*($CH_3$)$_2$), -4.4 (Si(*C*$H_3$)$_2$), -4.6 (Si(*C*$H_3$)$_2$), -5.1 (Si(*C*$H_3$)$_2$), -5.3 (Si(*C*$H_3$)$_2$) ppm.

Compound **6** was prepared following previously reported procedure for compound **3**.[9] Compound **5** (0.49 g, 1.05 mmol, 1.0 eq), TBAF (4.21 mL, 4.21 mmol, 4.0 eq). Product eluted off column at 70% EtOAc in cyclohexane; pure product was a white solid (0.19 g, 76%). **HRMS (m/z):** $[M^{+}Na]^{+}$ calcd. for $C_{13}H_{16}O_4$ Na, 259.0946; found, 259.0947. **$^{1}$H NMR** (600 MHz, $CDCl_3$) δ 7.39 – 7.28 (m, 5H, $H_{Ar}$), 6.35 (dd, *J* = 6.0, 1.7 Hz, 1H, H-1),

4.85 – 4.77 (m, 2H, OC*H*$_2$Ph), 4.72 (dd, *J* = 6.0, 2.6 Hz, 1H, H-2), 4.37 (dt, *J* = 6.8, 2.1 Hz, 1H, H-3), 3.95 – 3.90 (m, 1H, H-6), 3.88 – 3.83 (m, 2H, H-5, H-6'), 3.62 (dd, *J* = 9.1, 6.7 Hz, 1H, H-4), 2.26 (s, 2H, O*H*) ppm; **$^{13}$C NMR** (151 MHz, $CDCl_3$) δ 144.2 (C-1), 128.66 ($C_{Ar}$), 128.07 ($C_{Ar}$), 128.05 ($C_{Ar}$), 102.6 (C-2), 77.4 (C-5), 77.0 (C-4), 73.9 (O*C*H$_2$Ph), 68.7 (C-3), 61.4 (C-6) ppm.

Compound **7**: Protocol was adapted from literature with changes.[8] Briefly, 1.35 g of compound **6** (5.72 mmol, 1.0 eq) was dissolved in anhydrous DMF (0.36 M; 16 mL) and cooled to 0˚C. Portionwise, NaH (0.33 g, 13.7 mmol, 2.4 eq) was added before the reaction mixture was warmed to ambient temperature and stirred for 15 min. The reaction was then cooled to 0˚C again, and PMBCl (1.8 mL, 20.2 mmol, 2.2 eq) was added followed by stirring at ambient temperature overnight. After completion of reaction (confirmed by TLC), the mixture was cooled to 0˚C and quenched with anhydrous MeOH added dropwise. The solution was then transferred to a separatory funnel containing 4x reaction volumes of DI water, and the aqueous layer was extracted with diethyl ether 5x. The combined organic layer was washed with brine and dried over sodium sulfate. Solvent was evaporated under reduced pressure and the crude product was purified by column chromatography (30% EtOAc in cyclohexane) to obtain compound 8 as a white solid product (1.96 g, 72%). **HRMS (m/z):** $[M^{+}Na]^{+}$ calcd. for $C_{29}H_{32}O_6$ Na, 499.2091; found 499.2097. **$^1$H NMR** ($CDCl_3$, 500 MHz) δ 7.39 – 7.24 (m, 9H, $H_{Ar}$), 6.89 (dd, *J* = 8.7, 2.8 Hz, 4H, $H_{Ar}$), 6.45 (dd, *J* = 6.1, 1.4 Hz, 1H, H-1), 4.89 (dd, *J* = 6.2, 2.7 Hz, 1H, H-2), 4.85 (d, *J* = 11.3 Hz, 1H, OC*H*$_2$Ph), 4.65 (d, *J* = 11.3 Hz, 1H, OC*H*$_2$Ph), 4.59 – 4.51 (m, 4H, OC*H*$_2$Ph), 4.24 – 4.21 (m, 1H, H-3), 4.07 (ddd, *J* = 8.3, 5.1, 2.8 Hz, 1H, H-5), 3.86 (dd, *J* = 8.8, 6.2 Hz, 1H, H-4), 3.82 (d, *J* = 6.1 Hz, 6H, PhOC*H*$_3$), 3.79 (d, *J* = 5.1 Hz, 1H, H-6), 3.76 (dd, *J* = 10.7, 2.8 Hz, 1H, H-6') ppm; **$^{13}$C NMR** (151 MHz, $CDCl_3$) δ 159.3 ($C_{Ar}$), 159.2 ($C_{Ar}$), 144.7 (C-1), 130.5 ($C_{Ar}$), 130.1 ($C_{Ar}$), 129.6 ($C_{Ar}$), 129.5 ($C_{Ar}$), 129.4 ($C_{Ar}$), 128.43 ($C_{Ar}$), 128.39 ($C_{Ar}$), 127.9 ($C_{Ar}$), 127.7 ($C_{Ar}$), 113.9 ($C_{Ar}$), 113.8 ($C_{Ar}$), 100.1 (C-2), 76.8 (C-5), 75.5 (C-3), 74.5 (C-4), 73.8 (O*C*H$_2$Ph), 73.2 (O*C*H$_2$Ph), 70.2 (O*C*H$_2$Ph), 68.2 (C-6), 55.30 (PhO*C*H$_3$), 55.27 (PhO*C*H$_3$) ppm.

**Monomer 4:** A CSI cycloaddition was carried out on **10** and purified via column chromatography (50% EtOAc in cyclohexane) to obtain **monomer 4** as a white solid (25%). **HRMS (m/z):** $[M^{+}Na]^{+}$ calcd. for $C_{30}H_{33}NO_7$ Na, 542.2155; found, 542.2134. **$^1$H NMR** (500 MHz, $CDCl_3$) δ 7.28 (dd, *J* = 8.1, 3.2 Hz, 5H, $H_{Ar}$), 7.22 (d, *J* = 8.6 Hz, 2H, $H_{Ar}$), 7.15 (d, *J* = 7.9 Hz, 2H, $H_{Ar}$), 6.88 – 6.82 (m, 4H, $H_{Ar}$), 6.06 (s, 1H, N*H*), 5.53 (d, *J* = 4.5 Hz, 1H, H-1), 4.72 (d, *J* = 11.2 Hz, 1H, OC*H*$_2$Ph), 4.61 (d, *J* = 11.4 Hz, 1H, OC*H*$_2$Ph), 4.52 – 4.40 (m, 4H, OC*H*$_2$Ph), 4.07 (dd, *J* = 6.7, 2.8 Hz, 1H, H-3), 4.00 – 3.94 (m, 1H, H-5), 3.79 (d, *J* = 5.5 Hz, 6H, PhOC*H*$_3$), 3.63 (t, *J* = 4.2 Hz, 2H, H-6), 3.59 (dd, *J* = 9.1, 6.8 Hz, 1H, H-4), 3.40 (dt, *J* = 4.9, 2.8 Hz, 1H, H-2) ppm; **$^{13}$C NMR** (126 MHz, $CDCl_3$) δ 167.0 (C=O), 159.4 ($C_{Ar}$), 159.3 ($C_{Ar}$), 138.0 ($C_{Ar}$), 129.9 ($C_{Ar}$), 129.79 ($C_{Ar}$), 129.75 ($C_{Ar}$), 129.7 ($C_{Ar}$), 129.5 ($C_{Ar}$), 128.3 ($C_{Ar}$), 127.8 ($C_{Ar}$), 127.7 ($C_{Ar}$), 113.9 ($C_{Ar}$), 113.8 ($C_{Ar}$), 76.3 (C-1), 75.9 (C-3), 75.7 (C-4), 73.6 (O*C*H$_2$Ph), 73.1 (O*C*H$_2$Ph), 70.8 (O*C*H$_2$Ph), 69.8 (C-5), 68.9 (C-6), 55.3 (PhO*C*H$_3$), 55.0 (C-2) ppm.

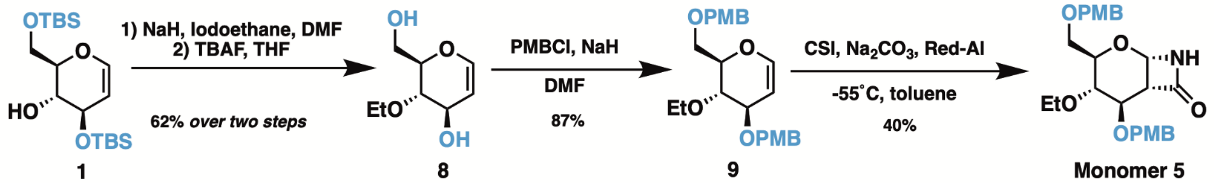


**Scheme S5**: Synthesis of monomer **5**.

***Monomer 5***

Compound **8**: Protocols were adapted from literature with changes.[7,9] Compound **1** (7.93 g, 21.2 mmol, 1.0 eq) was dissolved in anhydrous DMF (0.134 M; 158 mL) and cooled to 0˚C under $N_2$ protection. NaH (1.12 g, 46.64 mmol, 2.2 eq) was added portionwise, and the reaction mixture was stirred at ambient temperature for 15 min. The reaction was then cooled back to 0˚C, and ethyl iodide (8.5 mL, 106 mmol, 5.0 eq) was added dropwise before warming the solution back to ambient temperature and stirring overnight. After completion of reaction (confirmed by TLC), the reaction mixture was cooled to 0˚C and quenched with anhydrous MeOH dropwise. The mix was transferred to a separatory funnel containing 5x reaction volumes of DI water and the aqueous layer was extracted with diethyl ether 5x. The organic layer was washed with brine and dried over sodium sulfate. Solvent was evaporated under reduced pressure, and the crude product was a waxy white solid which was taken to the next step without further purification.

8.4 g (18.9 mmol, 1.0 eq) of crude material in a round bottom flask with 3Å molecular sieves was cooled to 0˚C and TBAF (1 M in THF; 75.6 mL, 75.6 mmol, 4.0 eq) was added dropwise. The reaction was stirred at ambient temperature overnight, after which the reaction was diluted with THF, filtered, and the solvent was evaporated. The crude product was purified by column chromatography (70% EtOAc in cyclohexane) to afford a white solid (2.3 g, 62% *over two steps*). **HRMS (m/z):** $[M^-H]^-$ calcd. for $C_8H_{13}O_4$, 173.0814; found, 173.0808. **$^1$H NMR** (600 MHz, $CDCl_3$) δ 6.35 (dd, *J* = 6.0, 1.7 Hz, 1H, H-1), 4.74 (dd, *J* = 6.0, 2.5 Hz, 1H, H-2), 4.31 (d, *J* = 4.9 Hz, 1H, H-3), 3.95 – 3.89 (m, 1H, $OCH_2CH_3$), 3.89 – 3.80 (m, 3H, H-6, $OCH_2CH_3$, H-5), 3.79 – 3.70 (m, 1H, H-6’), 3.47 (dd, *J* = 9.1, 6.8 Hz, 1H, H-4), 2.14 (s, 2H, O*H*), 1.23 (t, *J* = 7.0 Hz, 3H, $OCH_2CH_3$) ppm; **$^{13}$C NMR** (151 MHz, $CDCl_3$) δ 144.2 (C-1), 103.3 (C-2), 77.6 (C-4), 77.4 (C-5), 69.1 (C-3), 67.7 (C-6), 61.9 ($OCH_2CH_3$), 15.7 ($OCH_2CH_3$) ppm.

Compound **9**: Protocol was adapted from literature with changes.[8] Compound **8** (1.9 g, 11.4 mmol, 1.0 eq) was dissolved in anhydrous DMF (0.3 M; 38 mL) and cooled to 0˚C. NaH (0.65 g, 27.2 mmol, 2.4 eq) was added portionwise and the reaction mixture was warmed to ambient temperature and stirred for 15 min. The reaction mixture was then cooled to 0˚C again, and PMBCl (3.4 mL, 24.9 mmol, 2.2 eq) was added. The reaction was stirred at ambient temperature overnight. After completion of reaction (confirmed by TLC), the mixture was cooled back to 0˚C and quenched with anhydrous MeOH dropwise. The mix was transferred to a separatory funnel containing 5x reaction volumes of DI water and the aqueous layer was extracted with diethyl ether 5x. The organic layer was washed with brine and dried over sodium sulfate. Solvent was evaporated under reduced pressure and the product was purified by column chromatography (10% EtOAc in cyclohexane) to obtain compound **9** as a colorless oil (4.1 g, 87%). **HRMS (m/z):** $[M^{+}Na]^{+}$ calcd. for $C_{24}H_{30}O_6$ Na, 437.1940; found, 437.1956. **$^{1}$H NMR** (600 MHz, $CDCl_3$) δ 7.27 (dd, *J* = 8.7, 2.0 Hz, 4H, $H_{Ar}$), 6.88 (dd, *J* = 8.7, 1.6 Hz, 4H, $H_{Ar}$), 6.40 (dd, *J* = 6.1, 1.4 Hz, 1H, H-1), 4.83 (dd, *J* = 6.2, 2.7 Hz, 1H, H-2), 4.62 – 4.47 (m, 4H, $OCH_2Ph$), 4.09 (ddd, *J* = 6.2, 2.8, 1.4 Hz, 1H, H-3), 3.99 (ddd, *J* = 8.4, 5.2, 2.9 Hz, 1H, H-5), 3.82 (dd, *J* = 7.0, 2.3 Hz, 1H, $OCH_2CH_3$), 3.80 (d, *J* = 1.2 Hz, 6H, $PhOCH_3$), 3.77 (dd, *J* = 10.7, 5.2 Hz, 1H, H-6), 3.73 (dd, *J* = 10.7, 2.9 Hz, 1H, H-6'), 3.68 – 3.59 (m, 2H, H-4, $OCH_2CH_3$), 1.16 (t, *J* = 7.0 Hz, 3H, $OCH_2CH_3$) ppm; **$^{13}$C NMR** (126 MHz, $CDCl_3$) δ 159.25 ($C_{Ar}$), 159.21 ($C_{Ar}$), 144.6 (C-1), 130.6 ($C_{Ar}$), 130.2 ($C_{Ar}$), 129.62 ($C_{Ar}$), 129.58 ($C_{Ar}$), 129.49 ($C_{Ar}$), 129.44 ($C_{Ar}$), 129.37 ($C_{Ar}$), 129.27 ($C_{Ar}$), 113.82 ($C_{Ar}$), 113.77 ($C_{Ar}$), 100.2 (C-2), 76.9 (C-5), 75.3 (C-3), 74.8 (C-4), 73.2 ($OCH_2Ph$), 70.3 ($OCH_2Ph$), 68.2 (C-6), 67.3 ($OCH_2CH_3$), 55.27 ($PhOCH_3$), 55.25 ($PhOCH_3$), 15.7 ($OCH_2CH_3$) ppm.

**Monomer 5**: CSI cycloaddition reaction with a minor modification was carried out on **12** to obtain **monomer 5** as a white solid after column purification (40% EtOAc in cyclohexane), (40%). After addition of compound **12** to CSI, solution was stirred at -55˚C for 1.5 hr instead of 3 hr. **HRMS (m/z):** $[M^{+}Na]^{+}$ calcd. for $C_{25}H_{31}NO_7$ Na, 480.1998; found, 480.2001. **$^{1}$H NMR** (600 MHz, $CDCl_3$) δ 7.34 – 7.22 (m, 4H, $H_{Ar}$), 6.87 (dd, *J* = 8.6, 4.9 Hz, 4H, $H_{Ar}$), 5.95 (s, 1H, N*H*), 5.51 (d, *J* = 4.5 Hz, 1H, H-1), 4.57 (dd, *J* = 43.9, 11.6 Hz, 2H, $OCH_2Ph$), 4.48 (d, *J* = 11.6 Hz, 2H, $OCH_2Ph$), 3.95 (dd, *J* = 6.8, 2.8 Hz, 1H, H-3), 3.91 (ddd, *J* = 9.2, 5.1, 2.8 Hz, 1H, H-5), 3.80 (d, *J* = 1.3 Hz, 6H, $PhOCH_3$), 3.76 – 3.70 (m, 1H, $OCH_2CH_3$), 3.68 (dd, *J* = 10.5, 5.0 Hz, 1H, H-6), 3.64 (dd, *J* = 10.5, 2.7 Hz, 1H, H-6'), 3.51 – 3.44 (m, 1H, $OCH_2CH_3$), 3.44 – 3.39 (m, 1H, H-4), 3.37 (dt, *J* = 5.0, 2.8 Hz, 1H, H-2), 1.09 (t, *J* = 7.0 Hz, 3H, $OCH_2CH_3$) ppm; **$^{13}$C NMR** (151 MHz, $CDCl_3$) δ 166.9 (C=O), 159.3 ($C_{Ar}$), 158.7 ($C_{Ar}$), 133.4 ($C_{Ar}$), 133.2 ($C_{Ar}$), 130.0 ($C_{Ar}$), 129.7 ($C_{Ar}$), 129.6 ($C_{Ar}$), 129.5 ($C_{Ar}$), 113.85 ($C_{Ar}$), 113.78 ($C_{Ar}$), 76.4 (C-1), 76.0 (C-3), 75.96 (C-4), 73.2 ($OCH_2Ph$), 70.9 ($OCH_2Ph$), 69.9 (C-5), 69.0 (C-6), 67.2 ($OCH_2CH_3$), 55.3 ($PhOCH_3$), 55.1 (C-2), 15.6 ($OCH_2CH_3$) ppm.

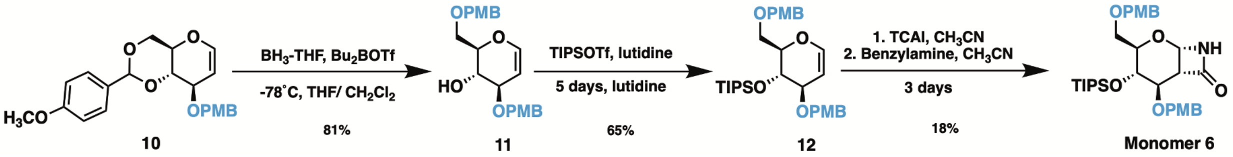


**Scheme S3**: Synthesis of monomer **6**

***Monomer 6***

Compound **10** was prepared following reported procedures with modifications.[8,11] Briefly, D-glucal (3.0 g, 20.5 mmol, 1.0 eq) was dissolved in anhydrous THF (0.7 M; 29.3 mL) under $N_2$ protection. To this was added *p*-anisaldehyde dimethyl acetal (5.26 mL, 28.7 mmol, 1.4 eq) followed by pyridinium *p*-toluenesulfonate (1.03 g, 4.1 mmol, 0.2 eq) and stirring for 2 hr. After 2 hr, the reaction was diluted in EtOAc and washed with brine, dried over sodium sulfate, and purified via column chromatography (20% EtOAc in cyclohexane) to afford a white solid (5.0 g, 92%). The dry product (1.0 g, 3.78 mmol, 1.0 eq) was then dissolved in anhydrous DMF (0.088 M; 43.2 mL) and cooled to 0˚C. To this, NaH (0.11 g, 4.54 mmol,1.2 eq) was added portionwise and the reaction was allowed to stir at ambient temperature for 15 min. The solution was then cooled back to 0˚C and PMBCl (0.561 mL, 4.16 mmol, 1.1 eq) was added before the reaction was stirred for 3 hr. Still at 0˚C, the reaction was quenched with 0.5 mL MeOH before adding to a separatory funnel containing 2x reaction volumes of water. The product was extracted with EtOAc 2x followed by washing the combined organic layer with brine and drying over sodium sulfate. The crude product was purified via precipitation (1:3 EtOAc:cyclohexane for 30 min at -20˚C) resulting in a white solid (0.99 g, 68%). **HRMS (m/z):** $[M^{+}Na]^{+}$ calcd. for $C_{22}H_{24}O_{6}$ Na, 407.1471; found, 407.1456. **$^{1}$H NMR** (600 MHz, $CDCl_3$) δ 7.44 (d, *J* = 8.9 Hz, 2H, $H_{Ar}$), 7.28 (d, *J* = 8.6 Hz, 2H, $H_{Ar}$), 6.92 (d, *J* = 8.8 Hz, 2H, $H_{Ar}$), 6.86 (d, *J* = 8.7 Hz, 2H, $H_{Ar}$), 6.35 – 6.32 (m, 1H, H-1), 5.60 (s, 2H, C*H*Ph), 4.79 (dd, *J* = 6.2, 2.0 Hz, 1H, H-2), 4.73 (d, *J* = 11.6 Hz, 1H, OC*H₂*Ph), 4.64 (d, *J* = 11.6 Hz, 1H, OC*H₂*Ph), 4.33 – 4.31 (m, 1H, H-6), 4.30 (dt, *J* = 7.4, 1.9 Hz, 1H, H-3), 4.00 (dd, *J* = 10.0, 7.5 Hz, 1H, H-4), 3.89 (td, *J* = 10.1, 4.9 Hz, 1H, H-5), 3.81 – 3.77 (m, 7H, H-6', PhOC*H₃*) ppm; **$^{13}$C NMR** (151 MHz, $CDCl_3$) δ 160.1 ($C_{Ar}$), 159.2 ($C_{Ar}$), 144.3 (C-1), 130.6 ($C_{Ar}$), 129.9 ($C_{Ar}$), 129.4 ($C_{Ar}$), 127.4 ($C_{Ar}$), 113.8 ($C_{Ar}$), 113.6 ($C_{Ar}$), 103.0 (C-2), 101.2 (*C*HPh), 80.0 (C-4), 72.3 (C-3), 71.7 (O*C*$H_2$Ph), 68.7 (C-5), 68.4 (C-6), 55.32 (PhO*C*$H_3$), 55.29 (PhO*C*$H_3$) ppm.

Synthesis of compound **11** was adapted from literature.[12] 1.1 g (2.84 mmol, 1.0 eq) of compound **10** was dissolved in anhydrous THF (0.05 M; 57 mL) under argon and cooled to -78˚C in an acetone/ dry ice bath, then treated with borane-THF complex (1 M solution in THF; 14.2 mL, 14.2 mmol, 5.0 eq). After 15 min, the mixture was treated with $Bu_2BOTf$ (1 M solution in $CH_2Cl_2$; 7.1 mL, 7.1 mmol, 2.5 eq) and stirred at -78˚C for 8 hr. The reaction was then quenched with $Et_3N$ (3.5 mL), followed by the dropwise addition of MeOH (105 mL) until effervescence ceased. The mixture was warmed to ambient temperature and stirred for 30 min, then concentrated by rotary evaporation and purified by column chromatography (40% EtOAc in cyclohexane) to yield compound **11** as a white powder (0.89 g, 81%). **HRMS (m/z):** $[M^{+}Na]^{+}$ calcd. for $C_{22}H_{26}O_{6}$ Na, 409.1627; found, 409.1634. **$^{1}$H NMR** (600 MHz, $CDCl_3$) δ 7.28 (t, *J* = 8.7 Hz, 4H, $H_{Ar}$), 6.88 (dd, *J* = 8.7, 2.2 Hz, 4H, $H_{Ar}$), 6.37 (dd, *J* = 6.1, 1.5 Hz, 1H, H-1), 4.82 (dd, *J* = 6.2, 2.3 Hz, 1H, H-2), 4.63 (d, *J* = 11.5 Hz, 1H, OC*H*$_2$Ph), 4.57 – 4.48 (m, 3H, *OCH*$_2$*Ph*), 4.08 – 4.03 (m, 1H, H-3), 3.96 – 3.92 (m, 2H, H-4, H-6), 3.80 (s, 6H, PhOC*H*$_3$), 3.76 (ap d, 2H, H-5, H-6'), 2.59 (s, 1H, O*H*) ppm; **$^{13}$C NMR** (151 MHz, $CDCl_3$) δ 159.34 ($C_{Ar}$), 159.29 ($C_{Ar}$), 144.61 (C-1), 130.5 ($C_{Ar}$), 129.9 ($C_{Ar}$), 129.7 ($C_{Ar}$), 129.6 ($C_{Ar}$), 129.5 ($C_{Ar}$), 129.4 ($C_{Ar}$), 129.3 ($C_{Ar}$), 129.1 ($C_{Ar}$), 113.92 ($C_{Ar}$), 113.87 ($C_{Ar}$), 100.2 (C-2), 76.9 (C-4), 76.0 (C-3), 73.3 (O*C*$H_2$Ph), 70.5 (O*C*$H_2$Ph), 69.0 (C-6), 68.9 (C-5), 55.3 (PhO*C*$H_3$) ppm.

Synthesis of compound **12** was adapted from literature.[5,13] Compound **11** (2.2 g, 5.95 mmol, 1.0 eq) was dissolved in anhydrous $CH_2Cl_2$ (0.13 M, 46 mL) under $N_2$. The reaction was cooled to 0˚C and 2,6-lutidine (4.13 mL, 35.7 mmol, 6.0 eq) dissolved in 10 mL of anhydrous $CH_2Cl_2$ was added. To this, tri-isopropyl triflate (7 mL, 26.2 mmol, 4.4 eq) dissolved in an additional 10 mL $CH_2Cl_2$ was added dropwise. The reaction was stirred at ambient temperature for 3 days, after which the reaction was quenched by addition of water. The organic layer was washed with water, brine, and dried over sodium sulfate. The crude product was purified using column chromatography (30% EtOAc in cyclohexane) to obtain **12** as a colorless oil (2.1 g, 65%). **HRMS (m/z):** $[M^{+}Na]^{+}$ calcd. for $C_{31}H_{46}O_{6}Si$ Na, 565.2961; found, 565.2980. **$^{1}$H NMR** (600 MHz, $CDCl_3$) δ 7.14 (d, *J* = 8.4 Hz, 2H, $H_{Ar}$), 7.09 (d, *J* = 8.4 Hz, 2H, $H_{Ar}$), 6.74 (t, *J* = 8.5 Hz, 4H, $H_{Ar}$), 6.31 (d, *J* = 6.3 Hz, 1H, H-1), 4.77 (dd, *J* = 6.4, 3.9 Hz, 1H, H-2), 4.47 – 4.41 (ap dd, 2H, C$H_2$Ph), 4.35 – 4.23 (ap dd, 2H, C$H_2$Ph), 4.08 (dt, *J* = 7.6, 4.4 Hz, 1H, H-5), 3.99 (t, *J* = 4.7 Hz, 1H, H-4), 3.71 (t, *J* = 4.0 Hz, 1H, H-3), 3.67 (s, 6H, PhOC$H_3$), 3.65 – 3.55 (m, 2H, H-6), 0.96 (d, *J* = 4.7 Hz, 21H, Si(*i*-Pr)$_3$) ppm; **$^{13}$C NMR** (151 MHz, $CDCl_3$) δ 159.2 ($C_{Ar}$), 159.1 ($C_{Ar}$), 144.3 (C-1), 130.6 ($C_{Ar}$), 130.3 ($C_{Ar}$), 129.6 ($C_{Ar}$), 129.5 ($C_{Ar}$), 129.4 ($C_{Ar}$), 129.3 ($C_{Ar}$), 129.2 ($C_{Ar}$), 129.0 ($C_{Ar}$), 113.72 ($C_{Ar}$), 113.68, 98.2 (C-2), 78.3 (C-5), 73.8 (C-3), 73.0 (*C*$H_2$Ph), 69.5 (*C*$H_2$Ph), 68.4 (C-4), 68.2 (C-6), 55.18 (PhO*C*$H_3$), 55.16 (PhO*C*$H_3$), 18.2 (Si(*i*-Pr)$_3$), 18.1 (Si(*i*-Pr)$_3$), 17.8 (Si(*i*-Pr)$_3$), 12.6 (Si(*i*-Pr)$_3$), 12.4 (Si(*i*-Pr)$_3$) ppm.

**Monomer 6:** TCAI cycloaddition reaction was carried out on compound **12** to obtain **monomer 6** as a pale foam after purification via column chromatography (50% EtOAc in cyclohexane), (18%). **HRMS (m/z):** $[M^{+}Na]^{+}$ calcd. for $C_{32}H_{47}NO_{7}Si$ Na, 608.3020; found, 608.3034. **$^{1}$H NMR** (500 MHz, $CDCl_3$) δ 7.22 (dd, *J* = 16.4, 8.6 Hz, 4H, $H_{Ar}$), 6.85 (dd, *J* = 8.6, 4.5 Hz, 4H, $H_{Ar}$), 6.54 (d, *J* = 2.6 Hz, 1H, N*H*), 5.44 (d, *J* = 4.5 Hz, 1H, H-1), 4.55 (dd, *J* = 11.6, 5.0 Hz, 2H, OC$H_2$Ph), 4.40 (dd, *J* = 25.7, 11.6 Hz, 2H, OC$H_2$Ph), 4.03 (td, *J* = 7.1, 2.8 Hz, 1H, H-5), 3.87 (d, *J* = 8.3 Hz, 1H, H-3), 3.79 (m, 7H, H-4, PhOC$H_3$), 3.68 (dd, *J* = 10.2, 2.9 Hz, 1H, H-6), 3.57 (dd, *J* = 10.2, 7.0 Hz, 1H, H-6'), 3.36 – 3.33 (m, 1H, H-2), 0.95 (d, *J* = 10.3 Hz, 21H, Si(*i*-Pr)$_3$) ppm; **$^{13}$C NMR** (126 MHz, $CDCl_3$) δ 166.8 (C=O), 159.6 ($C_{Ar}$), 159.32 ($C_{Ar}$), 159.25 ($C_{Ar}$), 159.22 ($C_{Ar}$), 130.0 ($C_{Ar}$), 129.9 ($C_{Ar}$), 129.8 ($C_{Ar}$), 129.7 ($C_{Ar}$), 129.6 ($C_{Ar}$), 129.5 ($C_{Ar}$), 129.2 ($C_{Ar}$), 114.0 ($C_{Ar}$), 113.8 ($C_{Ar}$), 113.8 ($C_{Ar}$), 113.7 ($C_{Ar}$), 77.2 (C-3), 75.8 (C-1), 73.1 (O*C*$H_2$Ph), 73.0 (O*C*$H_2$Ph), 72.5 (C-5), 70.5 (C-4), 69.8 (C-6), 55.30 (PhO*C*$H_3$), 55.27 PhO*C*$H_3$), 53.3 (C-2), 18.2 (Si(*i*-Pr)$_3$), 18.1 (Si(*i*-Pr)$_3$), 18.0 (Si(*i*-Pr)$_3$), 12.7 (Si(*i*-Pr)$_3$), 12.4 (Si(*i*-Pr)$_3$) ppm.

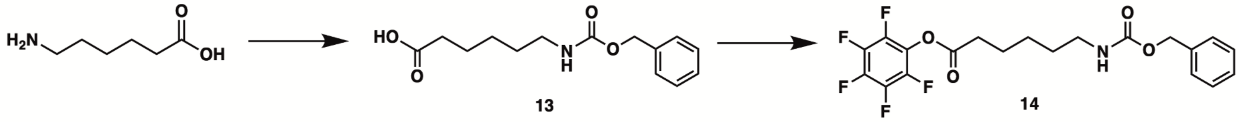


**Scheme S6:** Synthesis of pentafluorophenyl initiator

Compound **13:**[14] 6-aminohexanoic acid (6.42 g, 48.9 mmol, 1.0 eq) was dissolved in DI water (2 M; 24.5 mL) at ambient temperature. $NaHCO_3$ (6.73 g, 80.1 mmol, 1.64 eq) was added followed by cooling to 0˚C and the dropwise addition of benzyl chloroformate (7.6 mL, 53.8 mmol, 1.1 eq). The reaction was then allowed to warm back to ambient temperature and stirred for 16 hr - overnight. The following day, the reaction mixture was washed with 1M HCl before extracting the aqueous layer with chloroform 3x. Combined organic layers were washed with brine, dried over sodium sulfate, and solvent was removed. Product was purified via column chromatography (33% EtOAc in cyclohexane) to yield a white solid (2.58 g, 20%). Spectral data matches previously reported.[14]

Compound **14:**[15] DCC (2.20 g, 10.7 mmol, 1.1 eq) was added to a solution of compound **13** (2.58 g, 9.71 mmol, 1.0 eq) in anhydrous $CH_2Cl_2$ (0.5 M; 19.4 mL) under $N_2$ and stirred for 30 min or until fully dissolved. After this time, pentafluorophenol (1.97 g, 10.7 mmol, 1.1 eq) was added. The reaction was stirred overnight before removing solvent under vacuum. The crude was redissolved in EtOAc and filtered to remove DCU solids. The filtrate was concentrated and purified via column chromatography (10% EtOAc in cyclohexane) to yield a white solid (3.77 g, 90%). Spectral data matches previously reported.[15] **HRMS (m/z):** $[M^{+}H]^{+}$ calcd. for $C_{20}H_{18}F_5NO_4$, 432.1234, found 432.1229. **$^1$H NMR** (600 MHz, $CDCl_3$) δ 7.32 – 7.22 (m, 5H, $H_{Ar}$), 5.02 (s, 2H, O$CH_2$Bn), 4.70 (s, 1H, N*H*), 3.15 (d, *J* = 6.7 Hz, 2H, PfpOCOC$H_2$), 2.59 (t, *J* = 7.3 Hz, 2H, PfpOCOCH$_2$CH$_2$CH$_2$CH$_2$C$H_2$), 1.72 (p, *J* = 7.5 Hz, 2H, PfpOCOCH$_2$CH$_2$CH$_2$C$H_2$), 1.50 (q, *J* = 7.4 Hz, 2H, PfpOCOCH$_2$C$H_2$), 1.38 (p, *J* = 7.6, 7.1 Hz, 2H, PfpOCOCH$_2$CH$_2$C$H_2$) ppm; **$^{13}$C NMR** (151 MHz, $CDCl_3$) δ 169.33 ($C_{Ar}$F), 156.44 ($C_{Ar}$F), 136.57 ($C_{Ar}$F), 128.53 ($C_{Ar}$), 128.13 ($C_{Ar}$), 66.69 (O$C$H$_2$Bn), 40.75 (C-1), 33.16 (C-5), 29.58 (C-2), 25.92 (C-3), 24.34 (C-4) ppm; **$^{19}$F NMR** (565 MHz, $CDCl_3$) δ -152.84 (d, *J* = 17.5 Hz), -158.10 (t, *J* = 21.7 Hz), -162.36 (t, *J* = 19.6 Hz) ppm.

#### Polymerization and Post-polymerization Synthetic Procedures

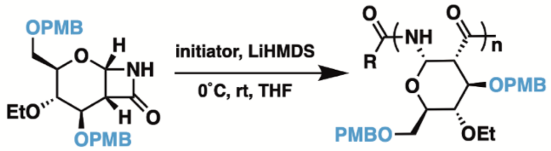


**General procedure for polymerization**

Protocol was adapted from literature with minor changes.[5,16] **Monomer 5** (0.125 g, 0.27 mmol, 1.0 eq) was dissolved in 2.7 mL* of distilled anhydrous THF (0.1 M) over 3Å MS, under $N_2$ protection, and at ambient temperature. Initiator **14**, previously dissolved to 0.5M in anhydrous THF, (0.083 eq for N=12, 0.04 eq for N=25, 0.02 eq for N=50, 0.014 eq for N=70, 0.01 eq for N=100, 0.005 eq for N=200) was added and the reaction was cooled to 0˚C. Next, LiHMDS, previously dissolved to 1M in anhydrous THF (0.108 mL, 0.018 g, 0.108 mmol, 0.4 eq) was added, and the solution was warmed to ambient temperature and stirred for 1 hr. To quench, a pipette full of saturated $NH_4Cl$ solution was added followed by the removal of THF under reduced pressure. The resulting solid was redissolved in $CH_2Cl_2$ and washed with 1M HCl, saturated $NaHCO_3$, and brine, and dried over sodium sulfate followed by removal of solvent. The crude polymer was redissolved in a minimal amount of $CH_2Cl_2$ and precipitated dropwise into cold *n*-pentane. Solid polymer was filtered out and dried under high vacuum to yield a white fluffy product (0.113 g, 90%).
* Volume of initiator and LiHMDS solution additions were accounted for in total volume of THF for a final monomer concentration of 0.1 M.

**Example for 3,6-di-*O*-PMB-4-*O*-Et-PAS: $^1$H NMR** (600 MHz, $CDCl_3$) δ 7.77 (br, 1H, N*H*), 7.19 – 7.00 (br, 4H, $H_{Ar}$), 6.86 – 6.80 (br, 4H, $H_{Ar}$), 5.64 (br, 1H, H-1), 4.63 – 3.78

(m, 6H, H-3, H-5, OC$H_2$Ph), 3.75 – 3.67 (br, 6H, PhOC$H_3$), 3.50 – 2.76 (m, 6H, H-2, H-4, H-6, OC$H_2$CH$_3$), 1.09 – 0.88 (br, 3H, OCH$_2$C$H_3$) ppm; **$^{13}$C NMR** (151 MHz, $CDCl_3$) δ 170.7 (C=O), 159.1 ($C_{Ar}$), 159.0 ($C_{Ar}$), 130.6 ($C_{Ar}$), 129.8 ($C_{Ar}$), 129.5 ($C_{Ar}$),129.3 ($C_{Ar}$), 128.9 ($C_{Ar}$), 114.0 ($C_{Ar}$),113.7 ($C_{Ar}$), 113.6 ($C_{Ar}$), 77.8 (C-1), 74.83, 73.6 (O$C$H$_2$Ph), 72.7 (O$C$H$_2$Ph), 68.0 (O$C$H$_2$CH$_3$), 67.6, 65.0, 64.1, 55.2, 55.1, 51.2 (C-2), 15.5 (OCH$_2$$C$H$_3$) ppm.

See **Table 1** for GPC data.

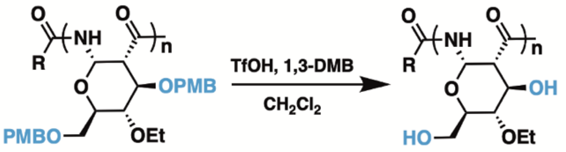


**General procedure for PMB deprotection using TfOH/1,3-DMB**

Protocol adapted from literature.[17] **3,6-di-*O*-PMB-4-*O*-Et-PAS** (0.115 g, 0.25 mmol of monomer, 1.0 eq) and 1,3-dimethoxy benzene (0.099 ml, 0.75 mmol, 3.0 eq) were dissolved in anhydrous $CH_2Cl_2$ (0.054 M; 4.6 mL), followed by the addition of trifluoromethanesulfonic acid (0.011 mL, 0.12 mmol, 0.5 eq). The reaction was stirred at ambient temperature for 20 min before quenching with MeOH (0.5 mL or until effervescence stopped and color disappeared). The solvent was removed, and the solid product was washed with $CH_2Cl_2$ until the pink color disappeared to obtain powdery white **3,6-di-OH-4-*O*-Et-PAS**. The polymer was dried and used for the next reaction without further purification.

**General procedure for PMB deprotection using DDQ/$H_2O$**

Protocol adapted from literature.[8,18] **3,6-di-*O*-PMB-4-*O*-Bn-PAS** (0.041g, 0.079 mmol of monomer, 1.0 eq) was dissolved in a 17:1 mixture of $CH_2Cl_2$: $H_2O$ (0.03 M; 2.63 mL total) and cooled to 0˚C. 2,3-dichloro-5,6-dicyano-p-benzoquinone (DDQ; 0.072 g, 0.316 mmol, 4.0 eq) was then added and the reaction was stirred for 30 min at 0˚C before warming to ambient temperature and stirring overnight. The reaction was added to ~10 mL of water and washed with $CH_2Cl_2$ 2x, before acidifying the aqueous layer with 1M HCl and washing again with EtOAc until the orange color disappeared from the aqueous layer (10x). The product was lyophilized to obtain a white powder and used without further purification.

**Example for 3,6-di-OH-4-*O*-Et-PAS: $^1$H NMR** (600 MHz, $D_2O$) δ 5.66 (s, 1H, H-1), 4.14 – 4.11 (br, 1H, H-3), 3.82 – 3.79 (br, 1H, OC$H_2$CH$_3$), 3.69 – 3.68 (br, 3H, H-5, H-6, OC$H_2$CH$_3$), 3.32 – 3.29 (br, 2H, H-6', H-4), 2.99 – 2.97 (br, 1H, H-2), 1.17 – 1.14 (br, 3H, OCH$_2$C$H_3$) ppm; **$^{13}$C NMR** (151 MHz, $D_2O$) δ 170.1 (C=O), 79.4 (C-4), 73.6 (C-1), 71.6 (C-6), 69.1 (O$C$H$_2$CH$_3$), 68.3 (C-3), 60.0 (C-5), 51.2 (C-2), 14.7 (OCH$_2$$C$H$_3$) ppm.

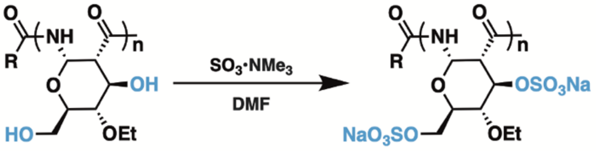

**General procedure for sulfation and NaOH workup**
Protocol adapted from literature.[16] **3,6-di-OH-4-*O*-Et-PAS** from the previous reaction (assume 100% yield: 0.25 mmol of monomer, 1.0 eq) was dissolved in anhydrous DMF (0.044 M; 5.7 mL) and $SO_3{\cdot}NMe_3$ (0.350 g, 2.5 mmol, 10.0 eq) was added. The reaction was heated to 50˚C and stirred for 24 hr. After 24 hr, the reaction was cooled to ambient temperature, and the polymer was precipitated out of the solution by addition of $CH_2Cl_2$. The precipitate obtained was dissolved in 1M NaOH and left under high vacuum with stirring for 3 hr. The polymer solution was then dialyzed against DI water for 24 hr and lyophilized to obtain white, fluffy **3,6-di-sulfate-4-*O*-Et-PAS** (67% *over two steps*).

**Example for 3,6-di-sulfate-4-*O*-Et-PAS: $^1$H NMR** (600 MHz, $D_2O$) δ 7.47 – 7.02 (br, 1H, N*H*), 6.17 – 5.83 (br, 1H, H-1), 5.14 ( br, 1H, H-3), 4.29 – 3.18 (m, 7H, H-6, H-5, OC$H_2$CH$_3$, H-4, H-2), 1.25 (s, 3H, OCH$_2$C$H_3$) ppm; **$^{13}$C NMR** (151 MHz, $D_2O$) δ 165.4 (C=O), 136.2, 131.4, 76.2 (C-2), 75.3 (C-3), 72.2 (C-1), 66.7, 66.8 (O*C*H$_2$CH$_3$), 66.2 (C-6), 51.6, 14.8, 14.5 (OCH$_2$*C*H$_3$) ppm.

See **Table 1** for GPC data.

**Table S1. Efficacy of sulfation reagents on 3,6-di-OH-4-*O*-Et-PAS.**

| Reagent | Conditions | % S | Polymer degradation (via GPC) |
|---|---|---|---|
| $SO_3 \cdot NMe_3$ | 10 equivalents, 50˚C, 24 hr | 12.81 | No degradation |
| $SO_3 \cdot NMe_3$ | 10 equivalents, 50˚C, 3 days | 8.59 | No degradation |
| $HSO_3Cl \cdot$ Pyridine | 157 equivalents, 60˚C, 3 hr | 9.94 | Some degradation |
| Concentrated $H_2SO_4$ | 78 equivalents, 0˚C, 30 min | 16.69 | Full degradation |
| Concentrated $H_2SO_4$ | 78 equivalents, 0˚C, 10 min | 7.03 | Full degradation |
| Concentrated $H_2SO_4$ | 50 equivalents, 0˚C, 20 min | 4.08 | Full degradation |
| $SO_3 \cdot$ Pyridine | 10 equivalents, 50˚C, 24 hr | - | Polymer not successfully purified. |
| $SO_3 \cdot$ DMF | 20 equivalents, 30˚C, 2 hr | 8.62 | No degradation |
| $SO_3 \cdot$ DMF | 50 equivalents, 70˚C, 24 hr | 10.48 | No degradation |
| $SO_3 \cdot$ DMF | 70 equivalents, 60˚C, 24 hr | 10.38 | No degradation |

Theoretical full sulfation at every 3 and 6 position on the polymer results in 16 %S. Experimental results from elemental analysis (Atlantic Microlab, Inc., Georgia, USA).

#### Small Molecule NMRs

##### Compound 1 - $^{1}H$ NMR (600 MHz, $CDCl_3$)

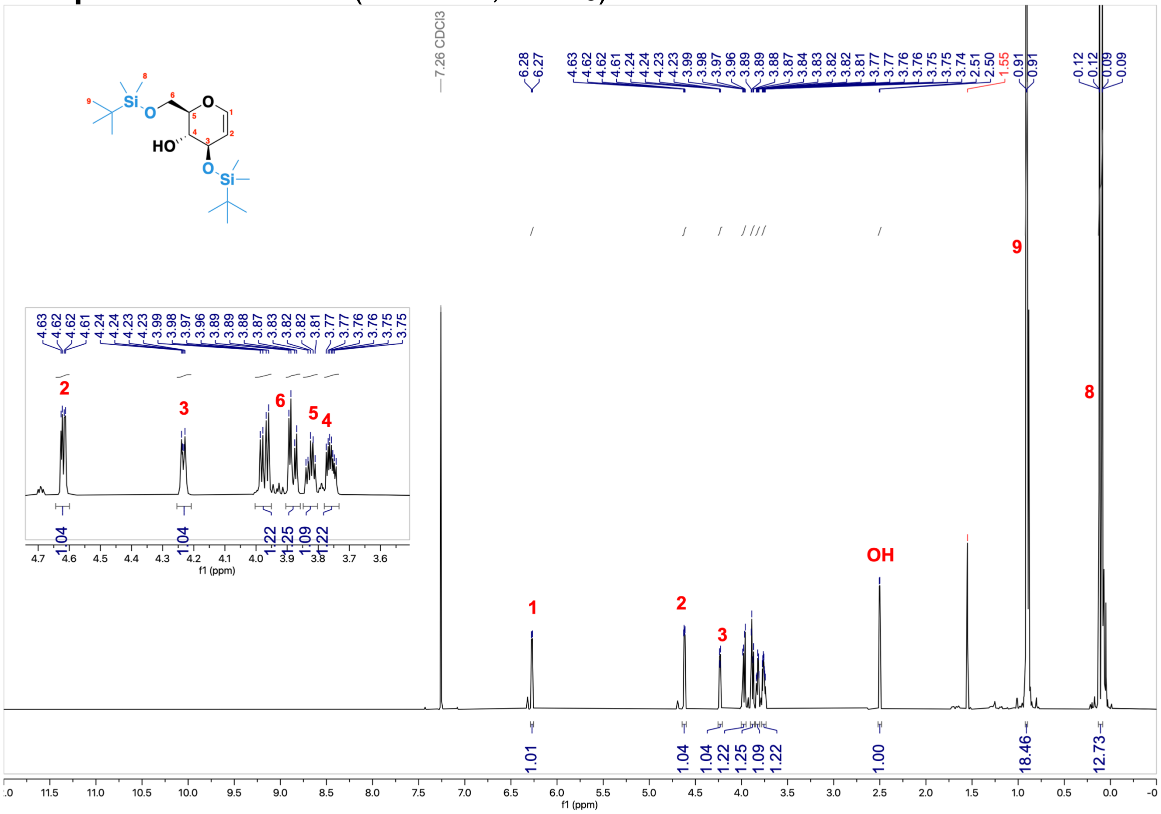

##### Compound 1 - $^{13}C$ NMR (151 MHz, $CDCl_3$)

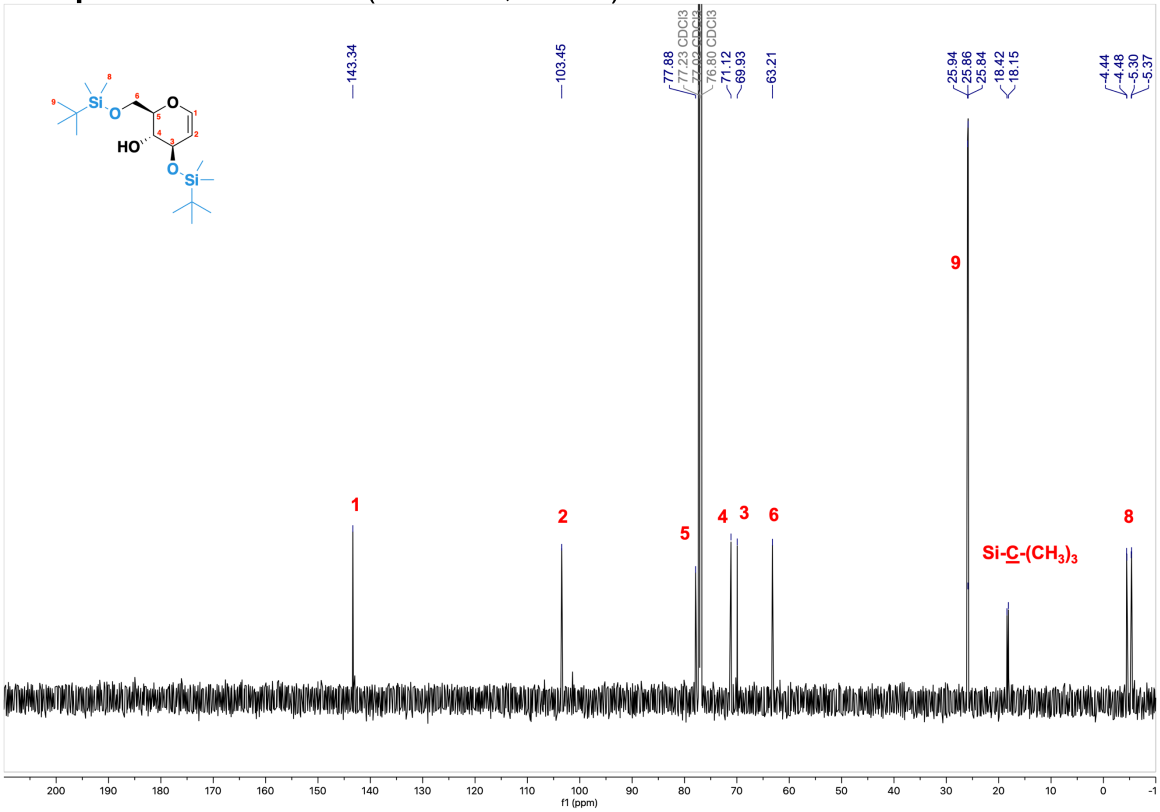

#### Compound 2 - $^{1}$H NMR (600 MHz, $CDCl_3$)

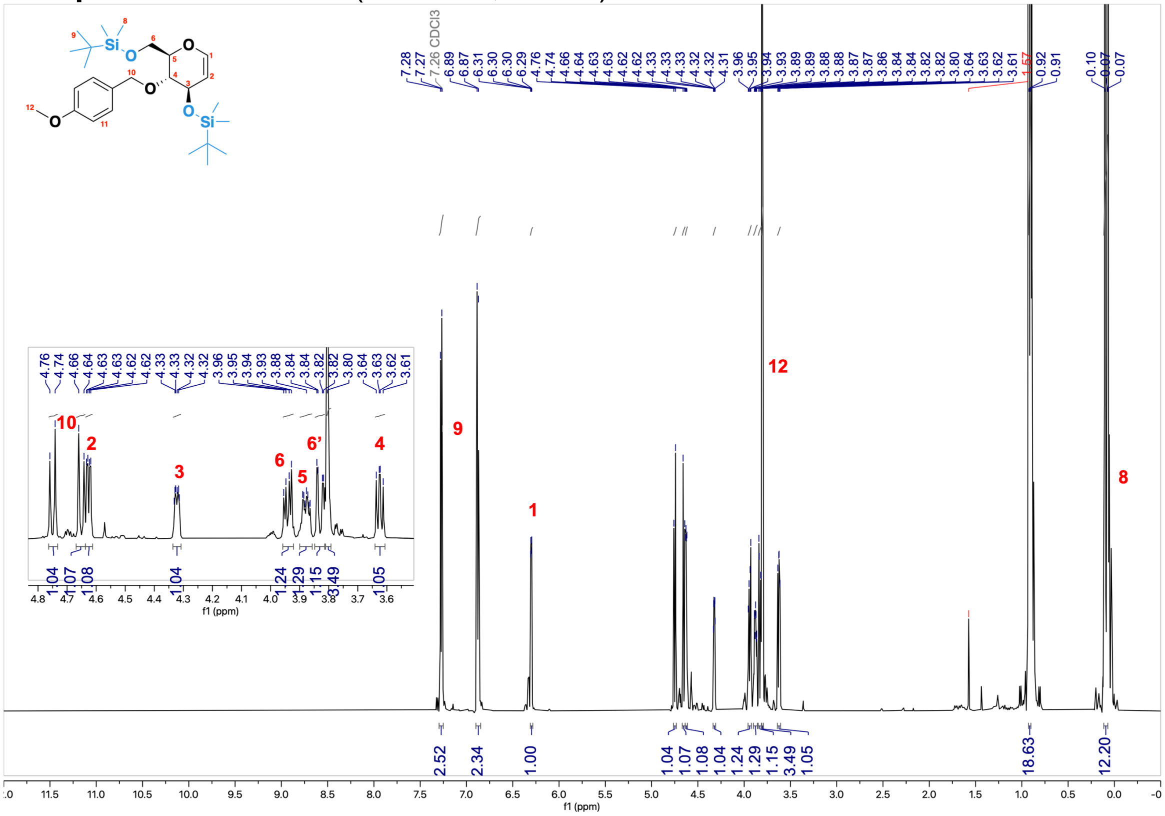


#### Compound 2 - $^{13}$C NMR (151 MHz, $CDCl_3$)

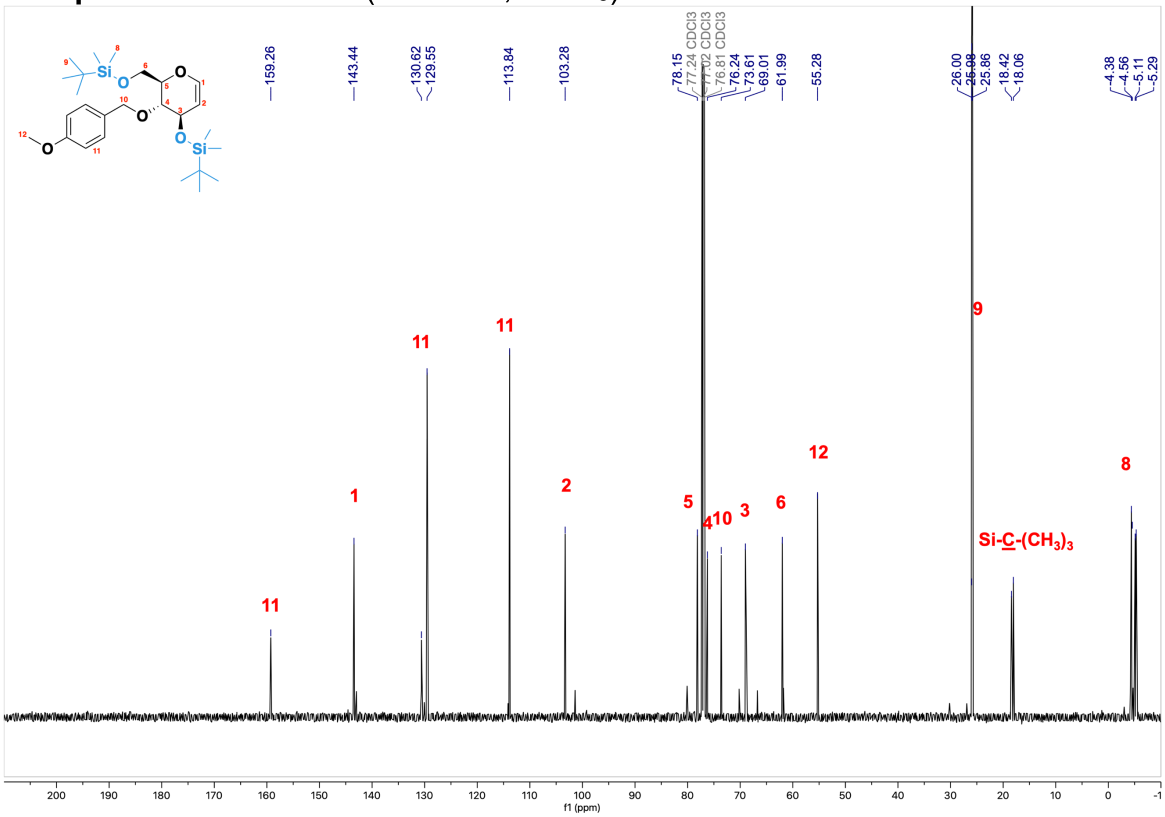

#### **Monomer 1** - $^{1}$H NMR (600 MHz, $CDCl_3$)

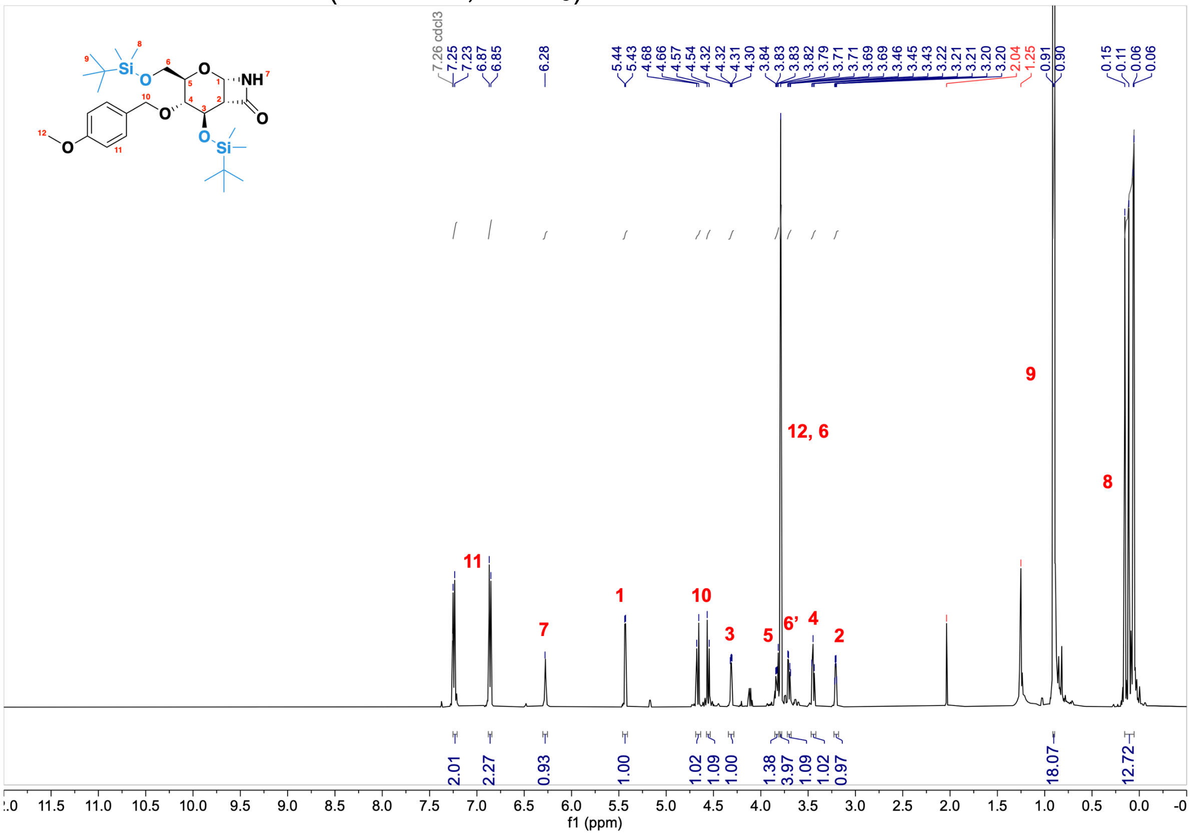


#### **Monomer 1** - $^{13}$C NMR (151 MHz, $CDCl_3$)

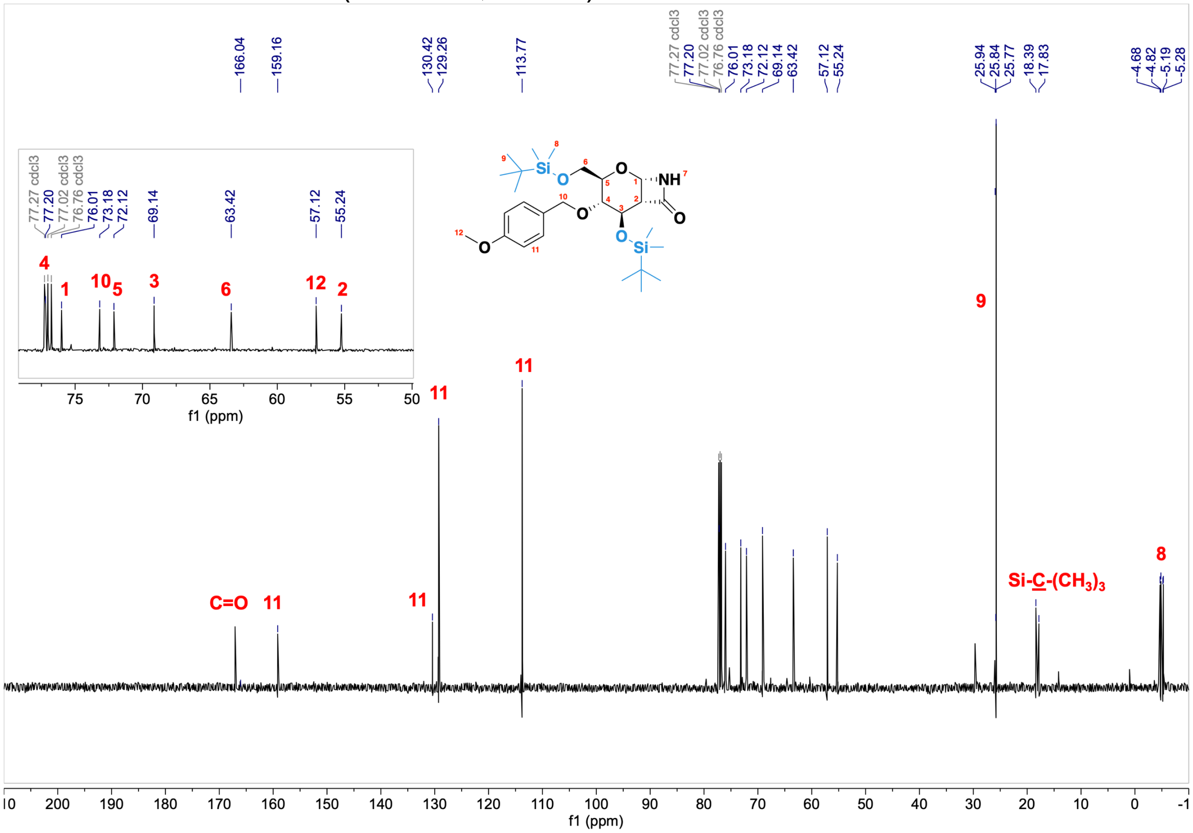

#### Monomer 1 - [$^{1}$H, $^{1}$H] COSY (2D, 600 MHz, $CDCl_3$)

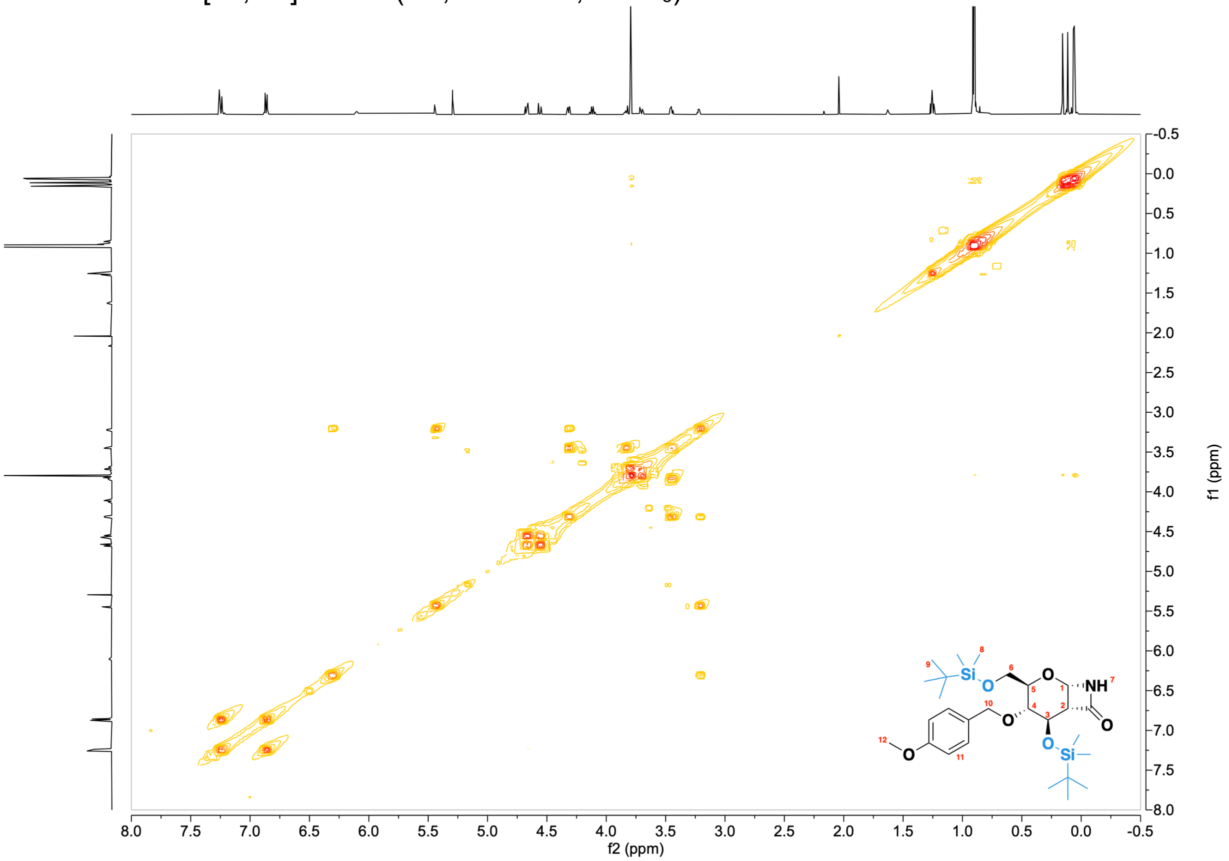


#### Monomer 1 - [$^{1}$H, $^{13}$C] HSQC (2D, 600 MHz, $CDCl_3$)

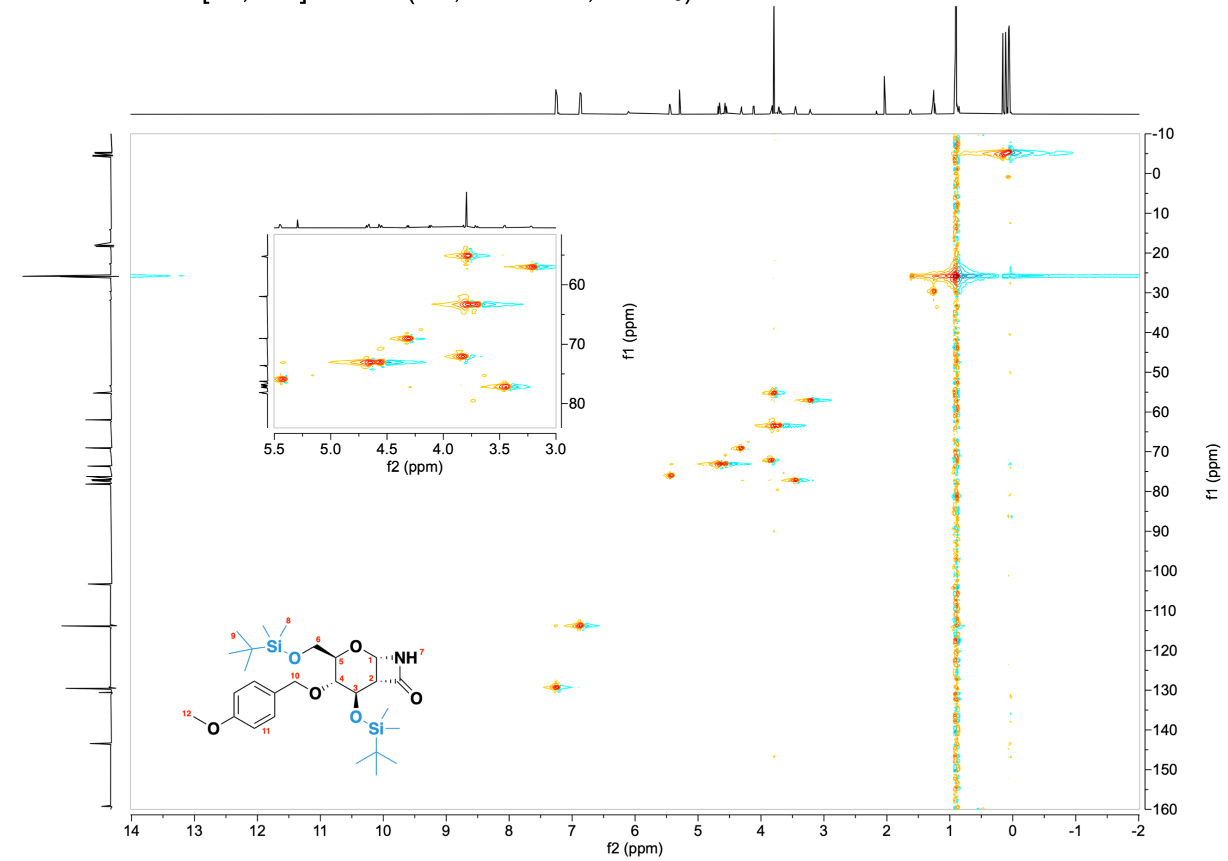

#### **Compound 3** - $^{1}$H NMR (600 MHz, $CDCl_3$)

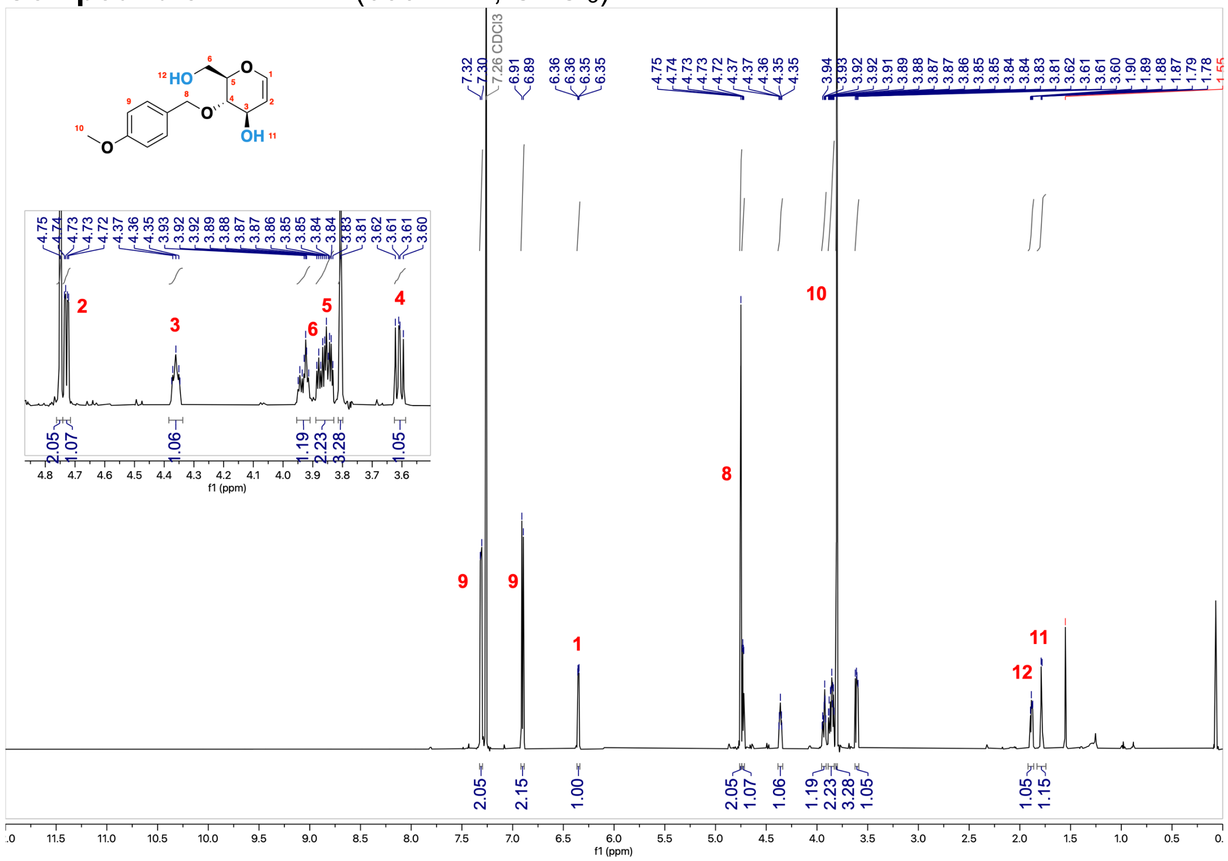


#### **Compound 3** - $^{13}$C NMR (151 MHz, $CDCl_3$)

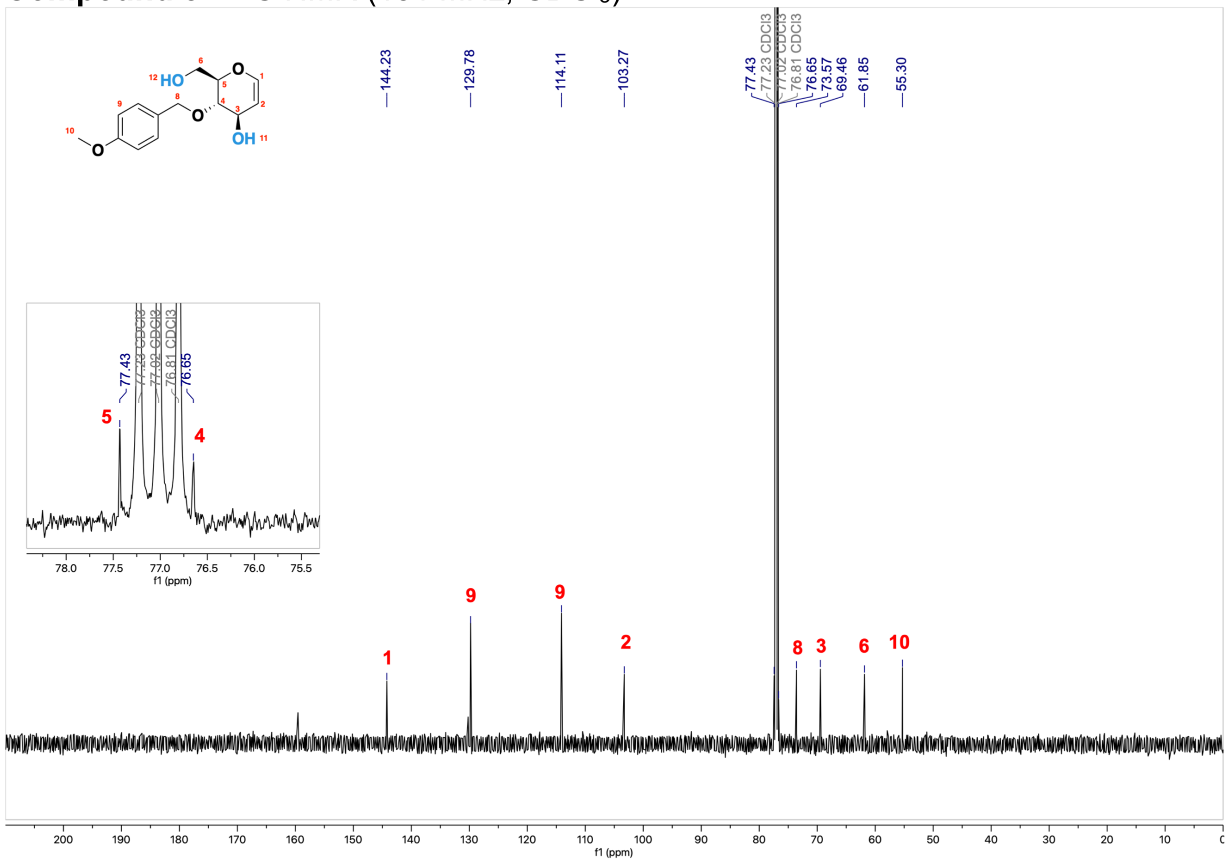

**Compound 4** - $^1$H NMR (600 MHz, $CDCl_3$)

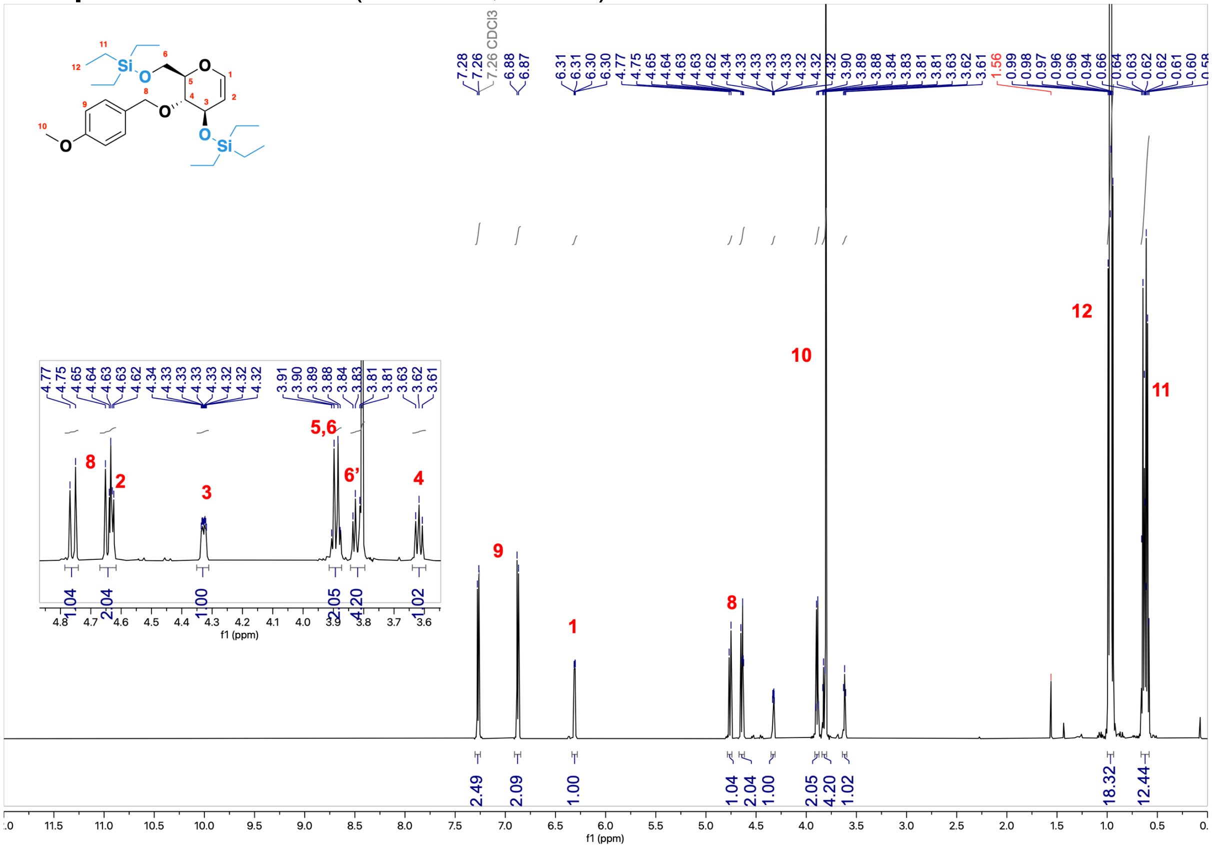

**Compound 4** - $^{13}$C NMR (151 MHz, $CDCl_3$)

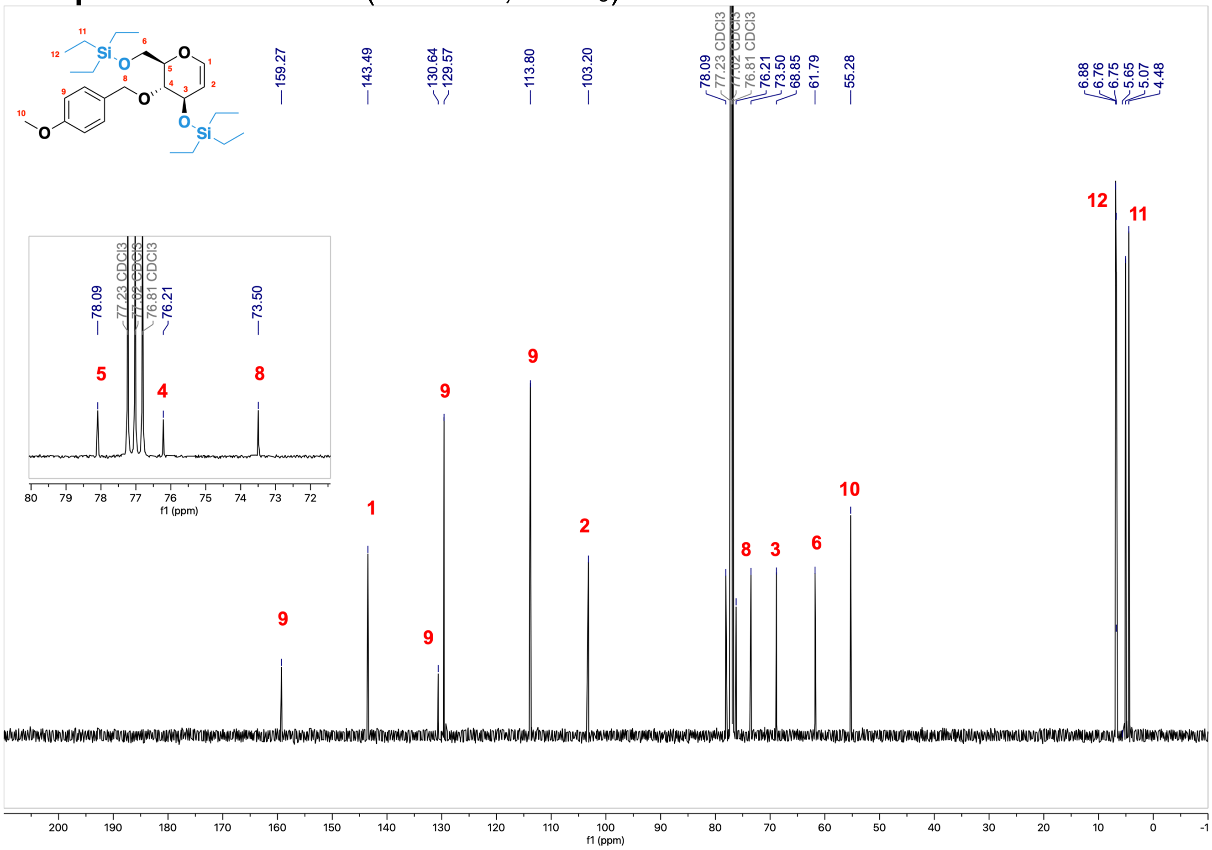

#### **Compound 4** - [$^1$H, $^1$H] COSY (2D, 600 MHz, $CDCl_3$)

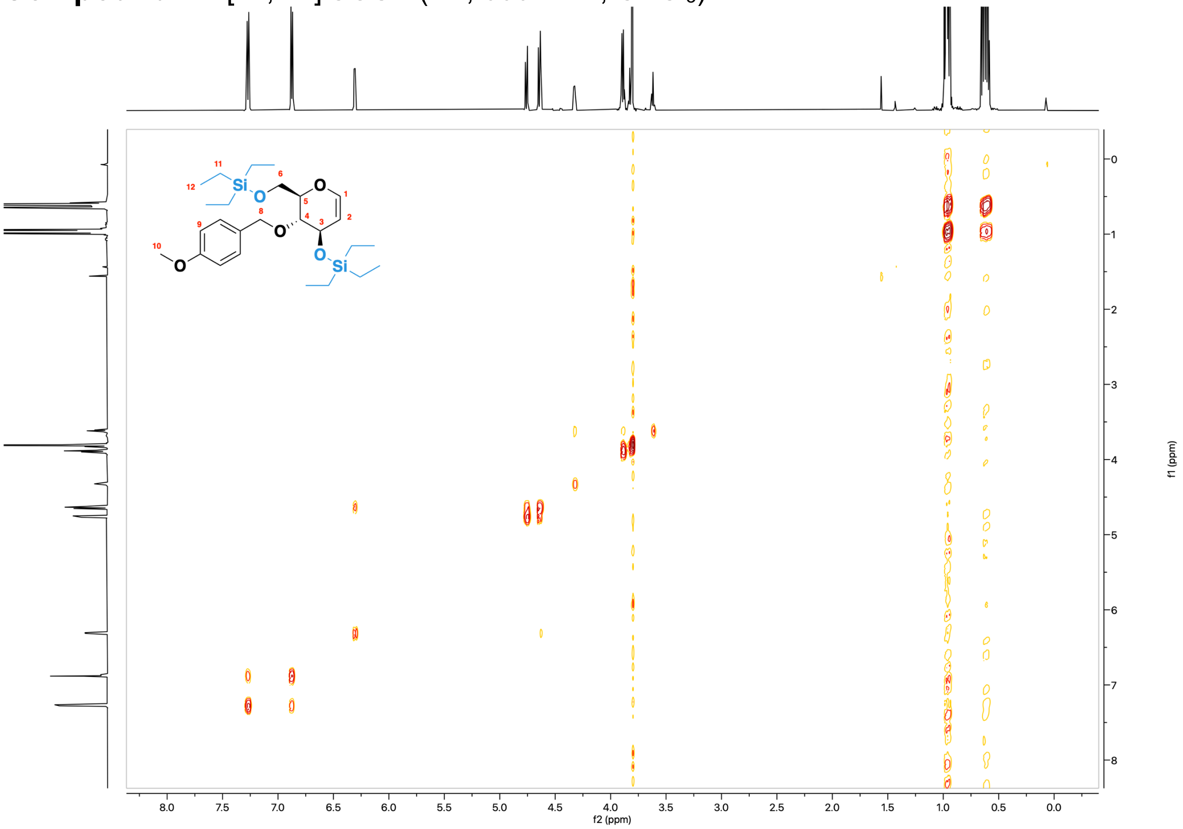


#### **Compound 4** - [$^1$H, $^{13}$C] HSQC (2D, 600 MHz, $CDCl_3$)

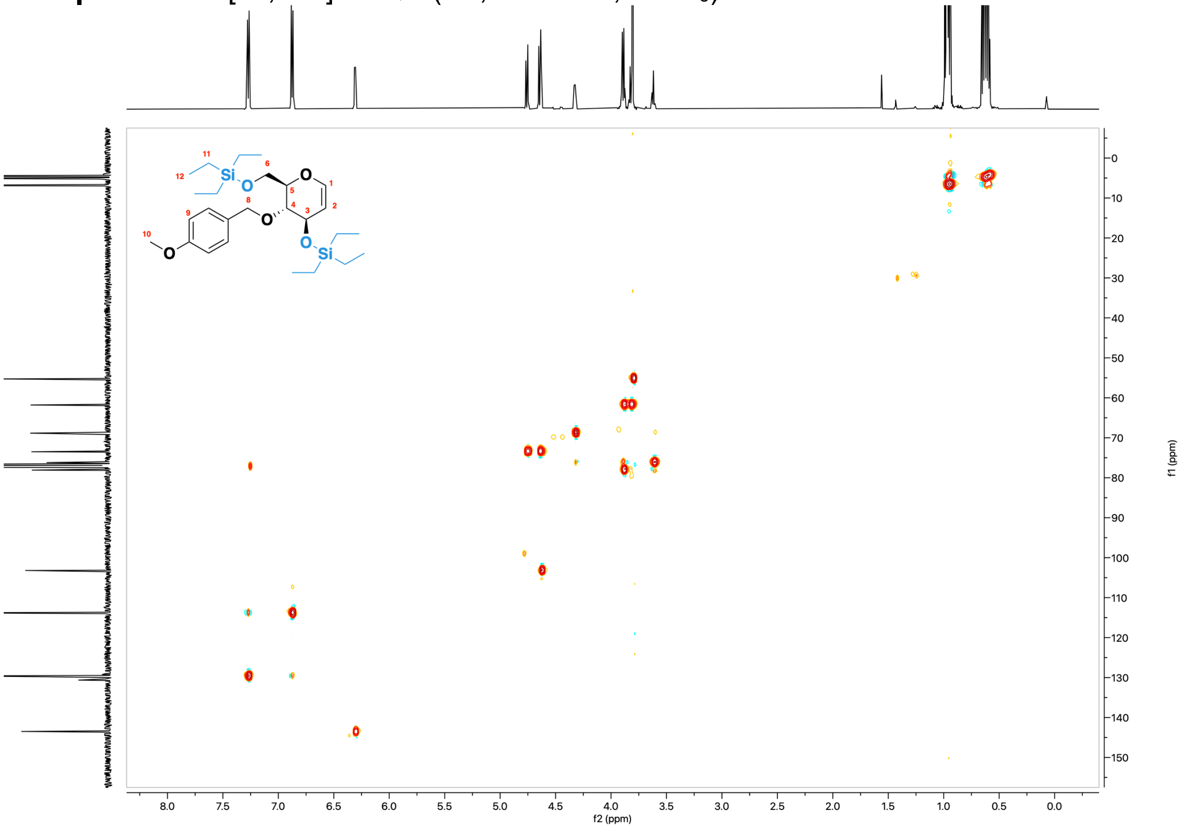

**Monomer 2** - $^{1}$H NMR (600 MHz, CDCl$_3$)

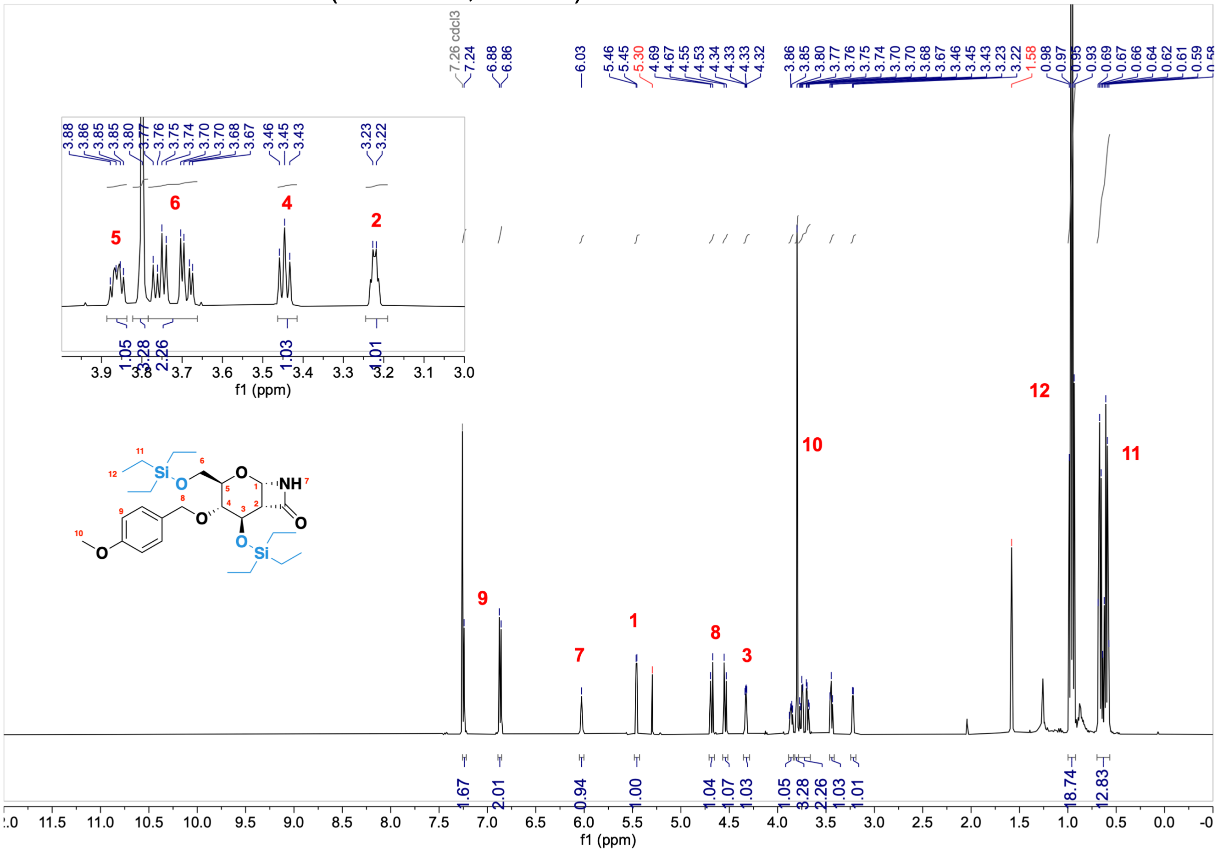


**Monomer 2** - $^{13}$C NMR (151 MHz, CDCl$_3$)

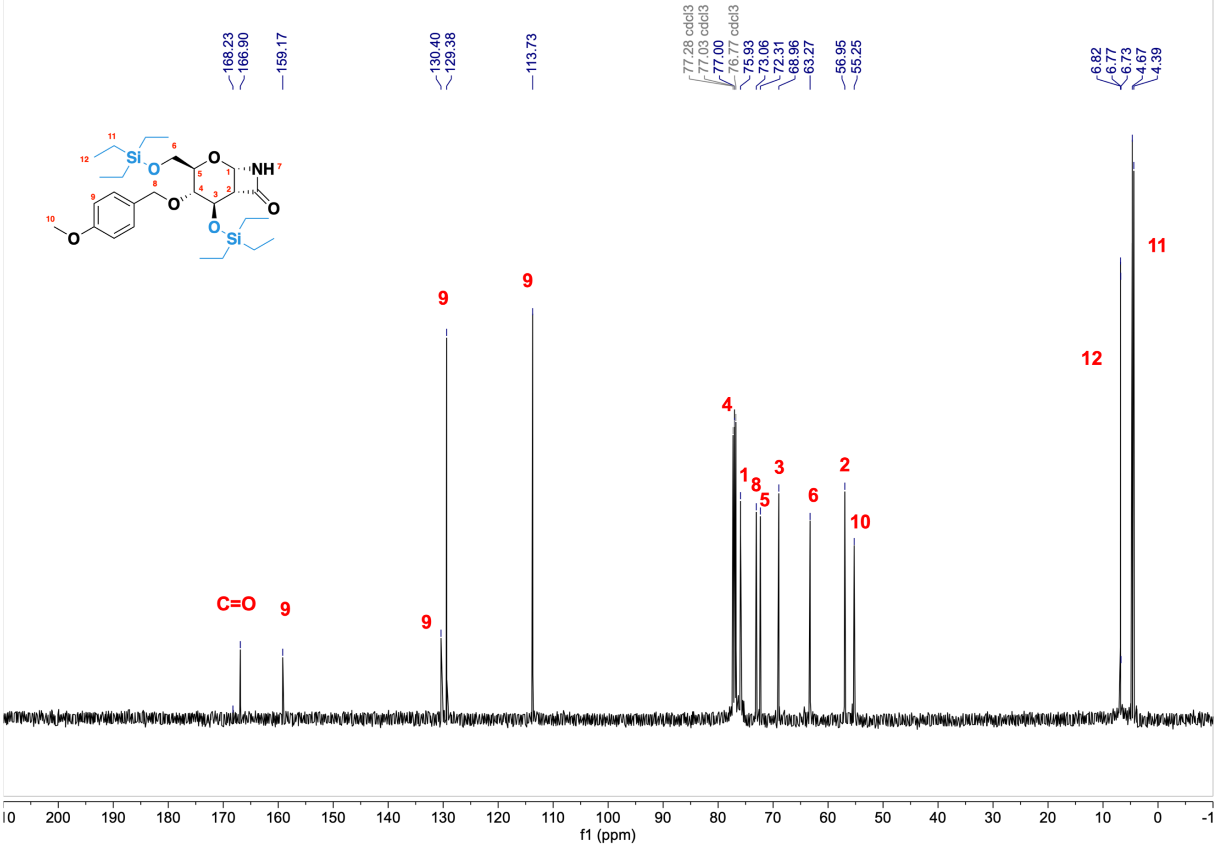

#### **Monomer 2** - [$^1$H, $^1$H] COSY (2D, 600 MHz, $CDCl_3$)

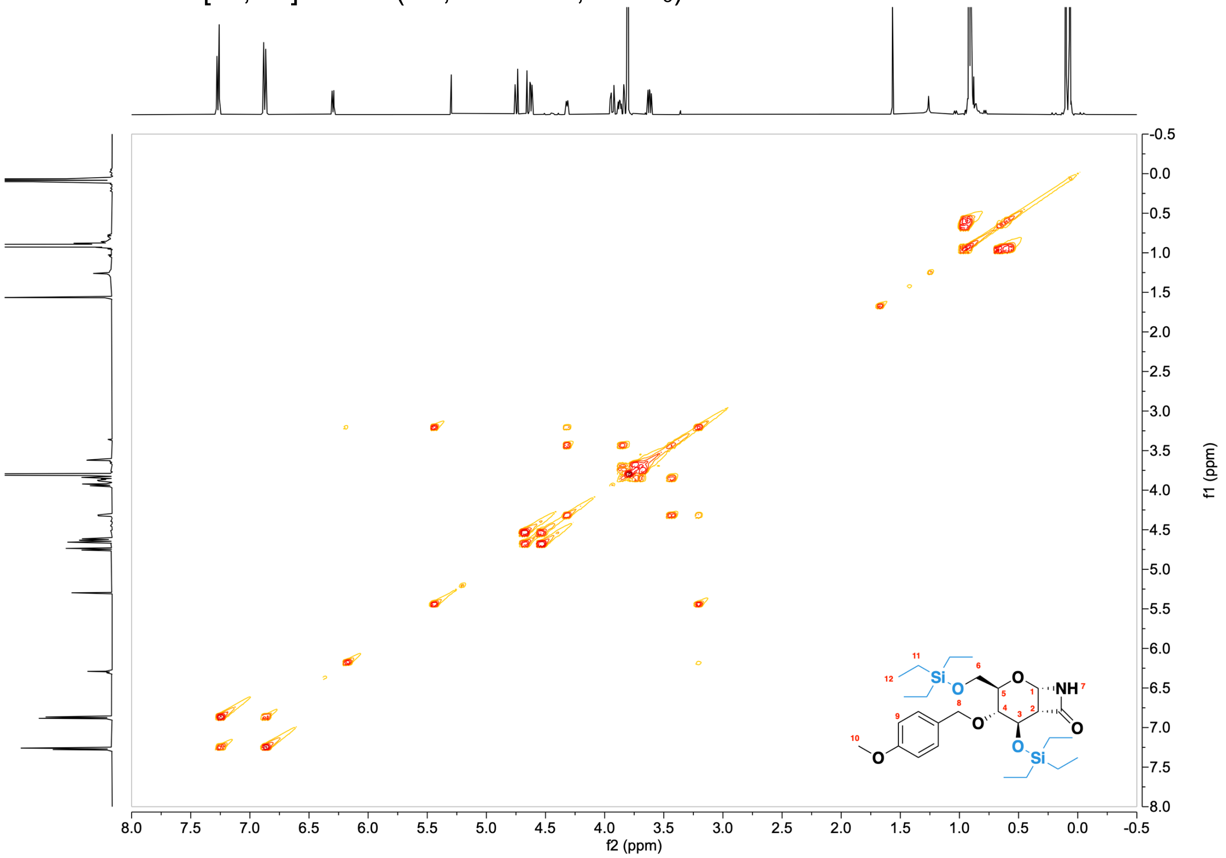


#### **Monomer 2** - [$^1$H, $^{13}$C] HSQC (2D, 600 MHz, $CDCl_3$)

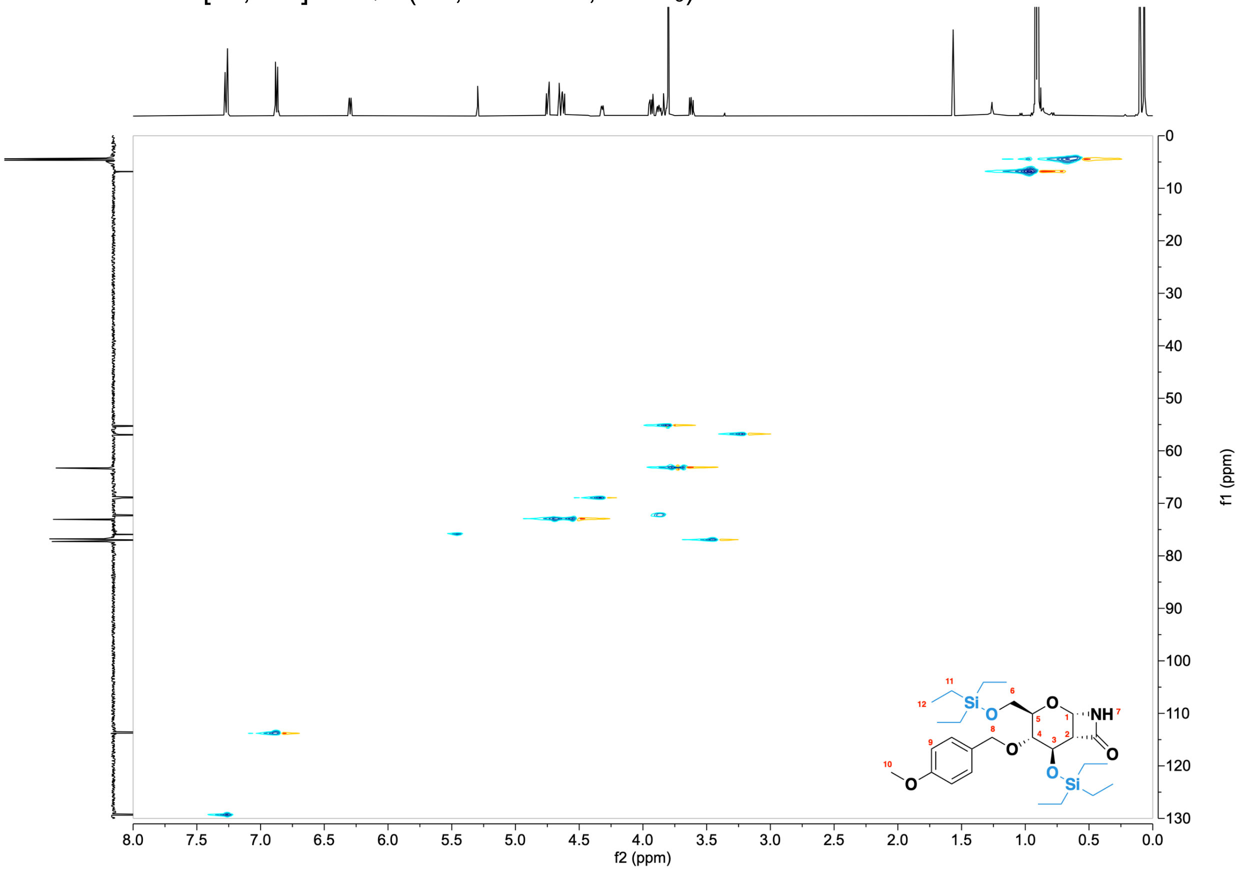

#### Compound 5 - $^{1}$H NMR (600 MHz, $CDCl_3$)

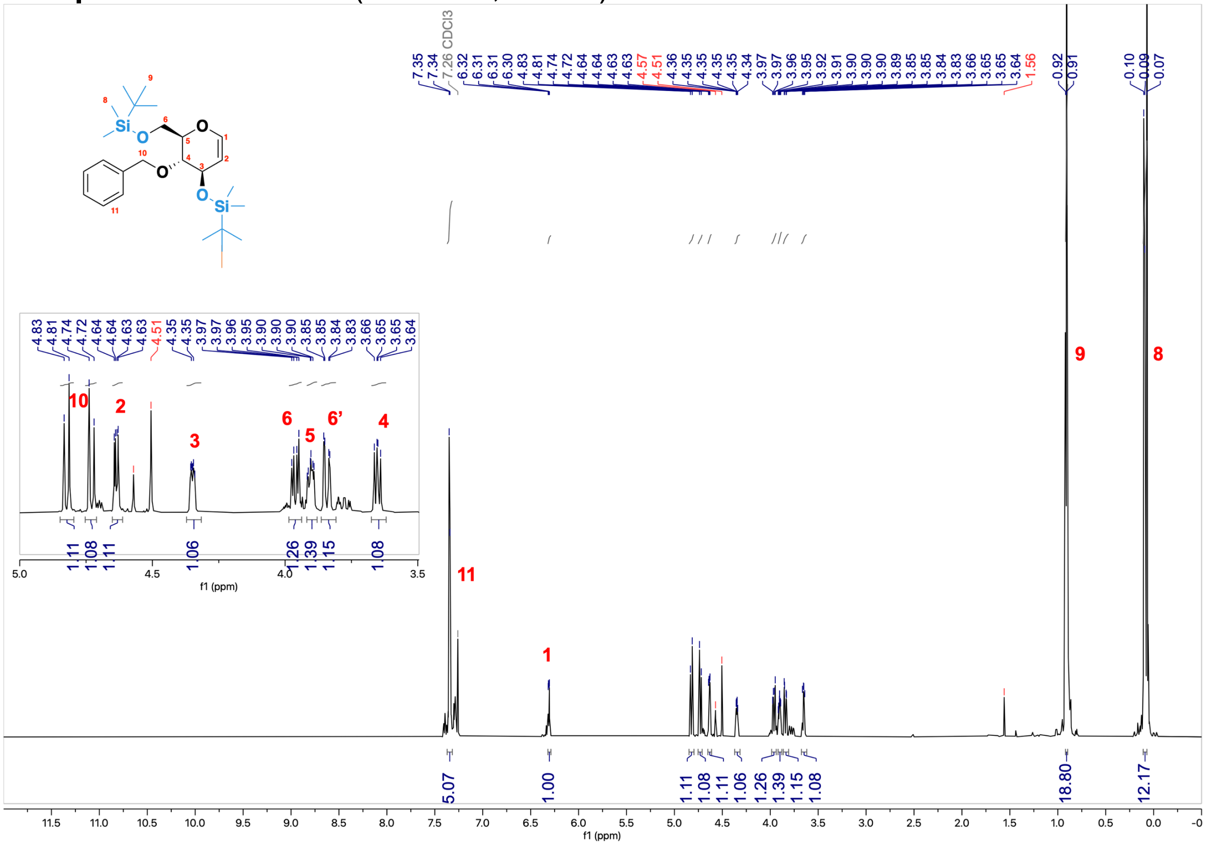


#### Compound 5 - $^{13}$C NMR (151 MHz, $CDCl_3$)

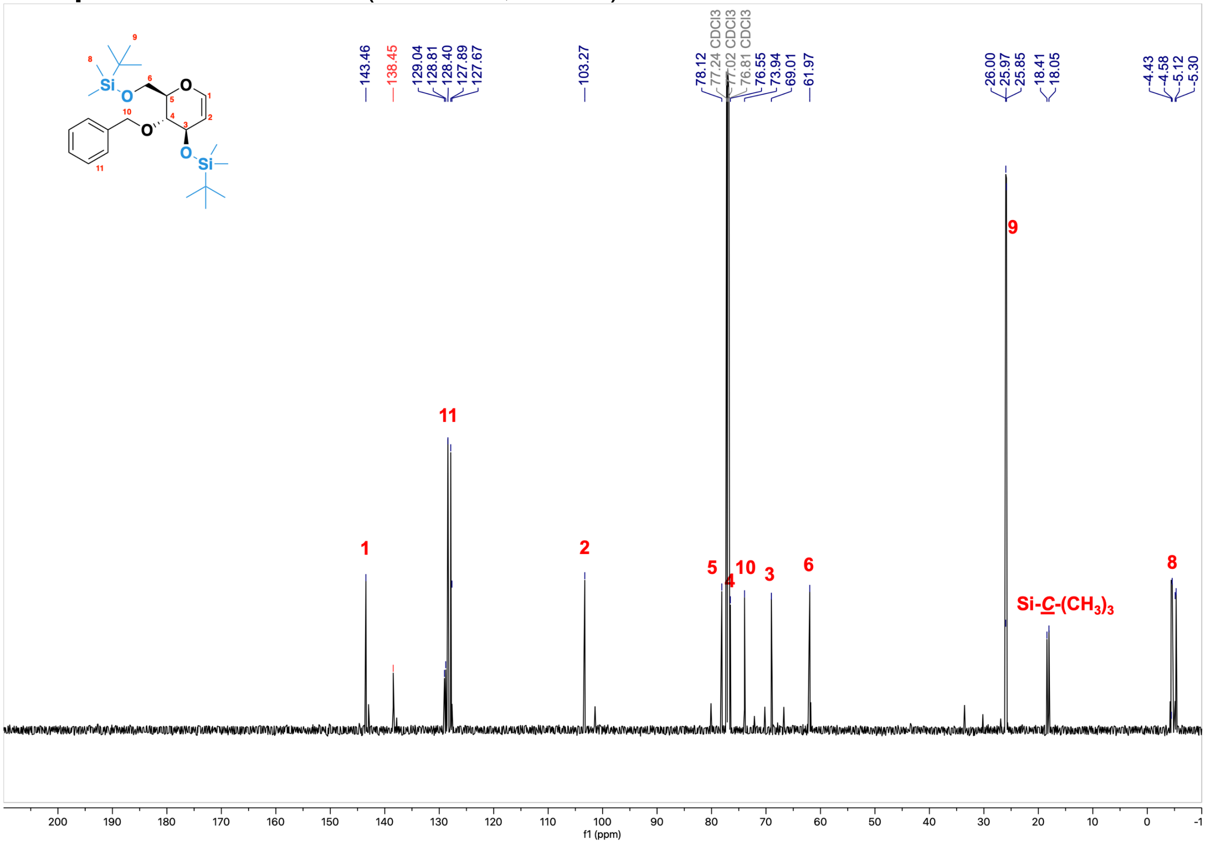

**Compound 6** - $^{1}$H NMR (600 MHz, $CDCl_3$)

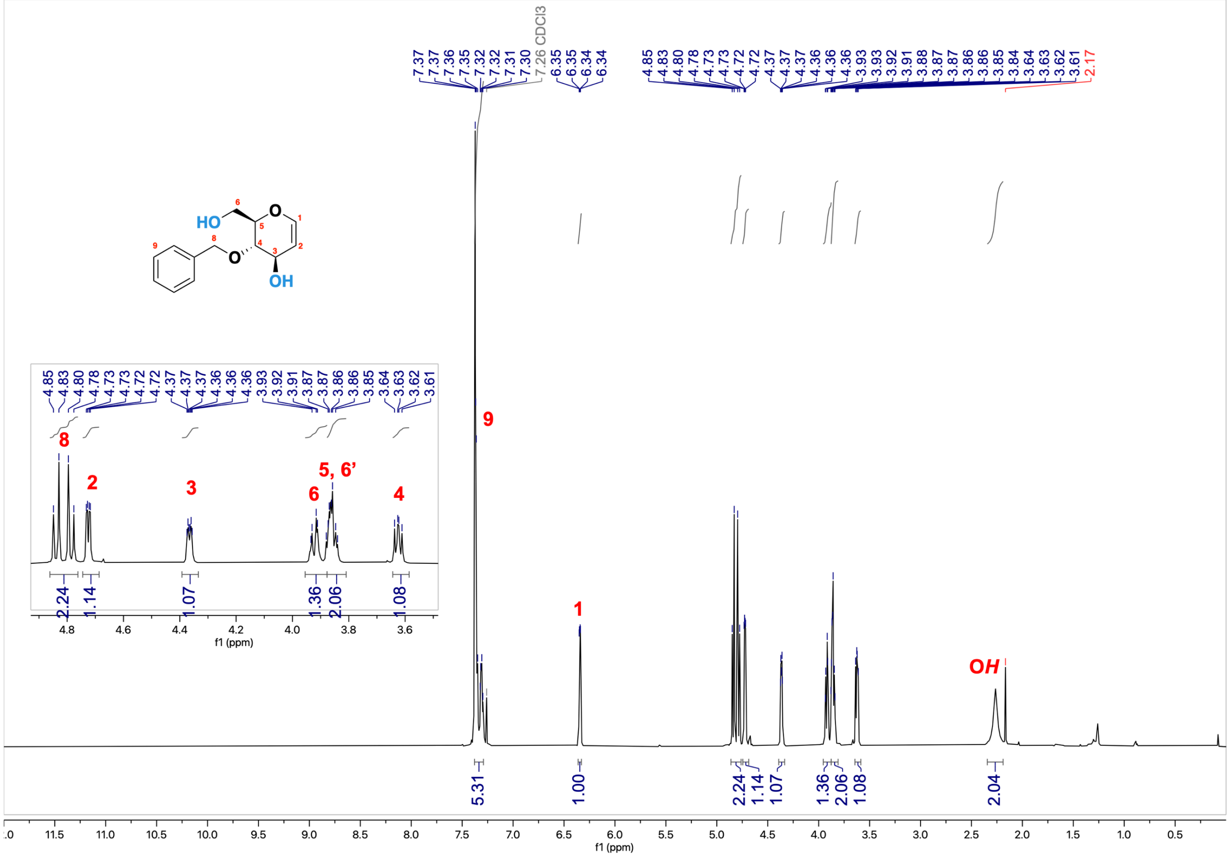

**Compound 6** - $^{13}$C NMR (151 MHz, $CDCl_3$)

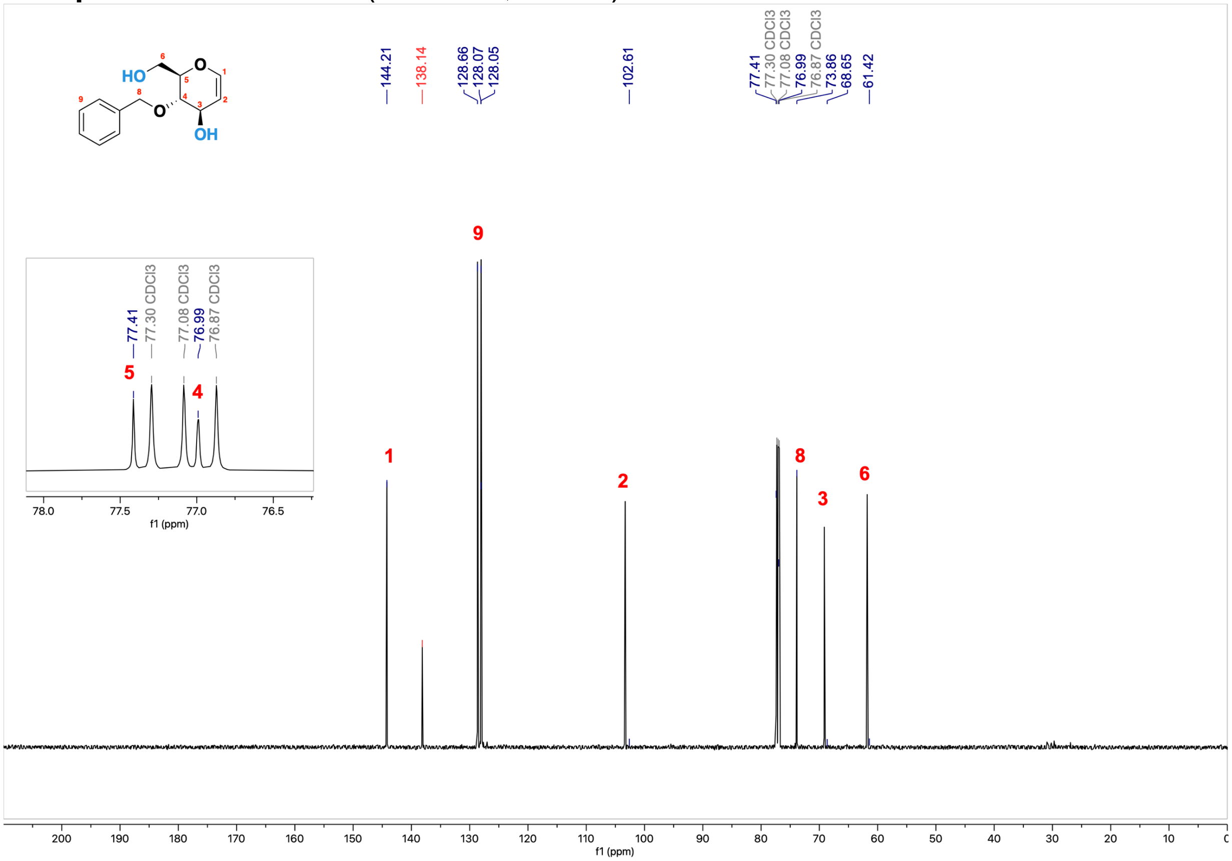

#### Compound 7 - $^1$H NMR (600 MHz, $CDCl_3$)

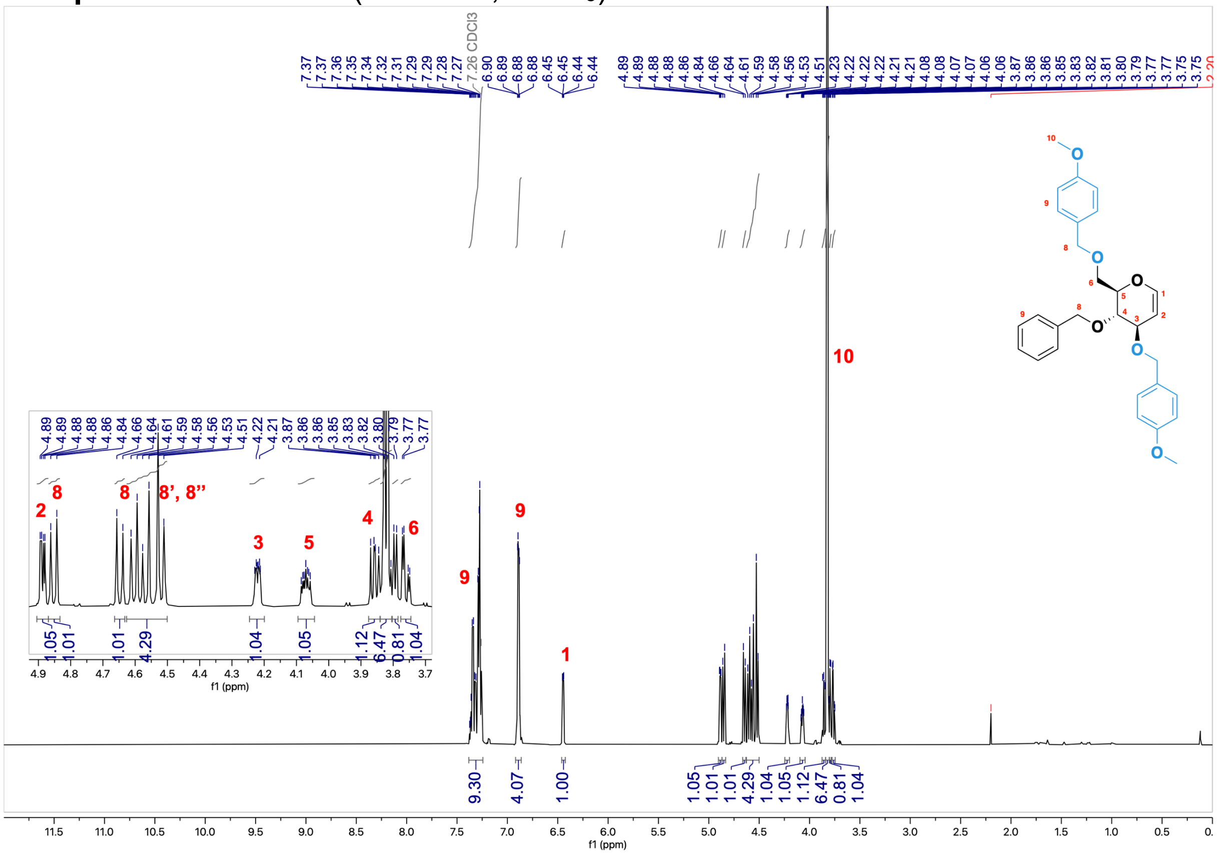


#### Compound 7 - $^{13}$C NMR (151 MHz, $CDCl_3$)

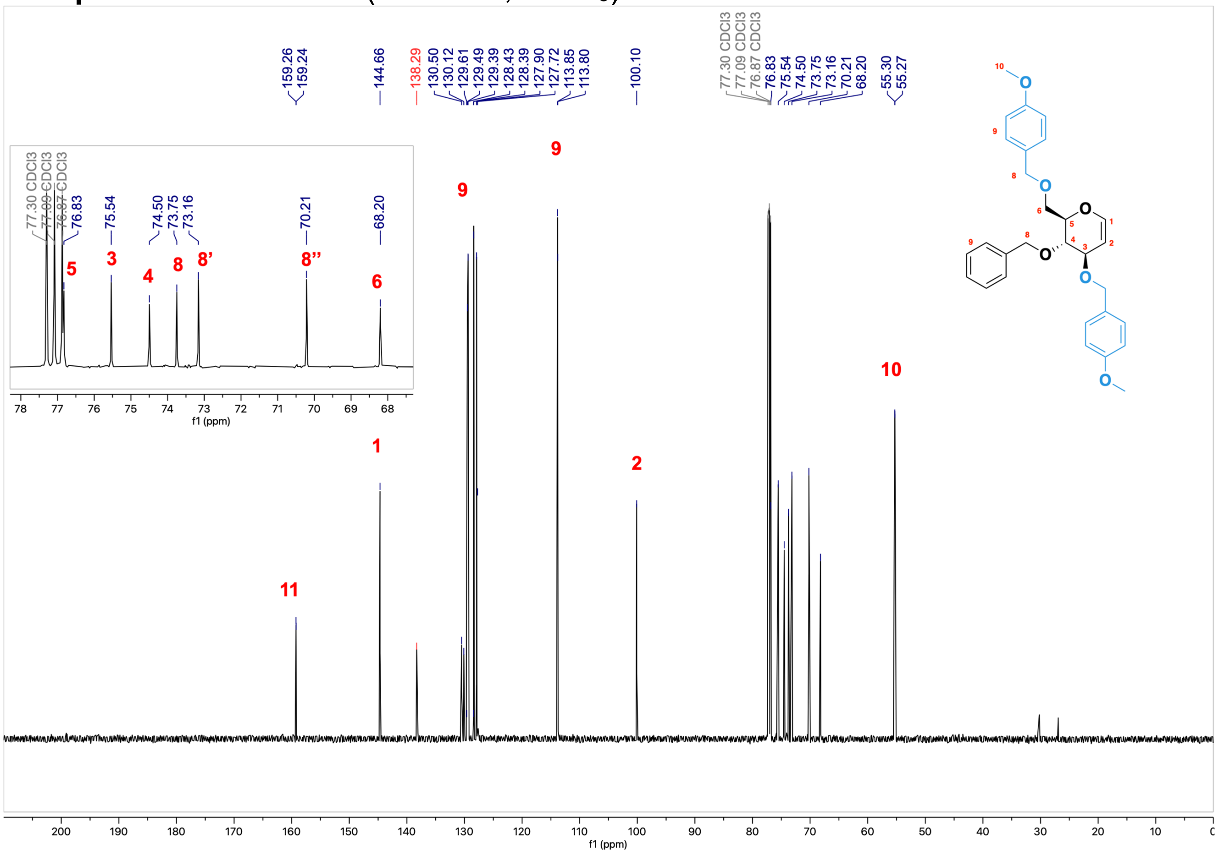

#### **Compound 7** - [$^1$H, $^1$H] COSY (2D, 600 MHz, $CDCl_3$)

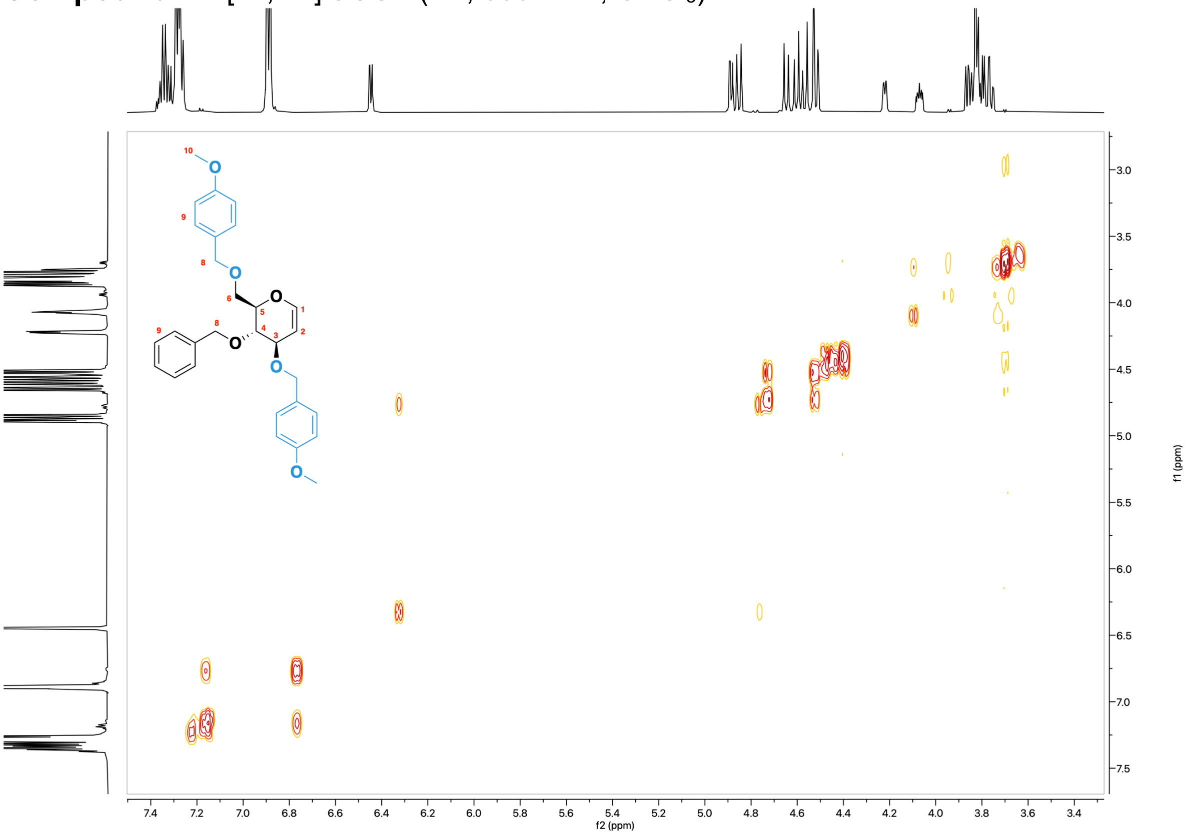

#### **Compound 7** - [$^1$H, $^{13}$C] HSQC (2D, 600 MHz, $CDCl_3$)

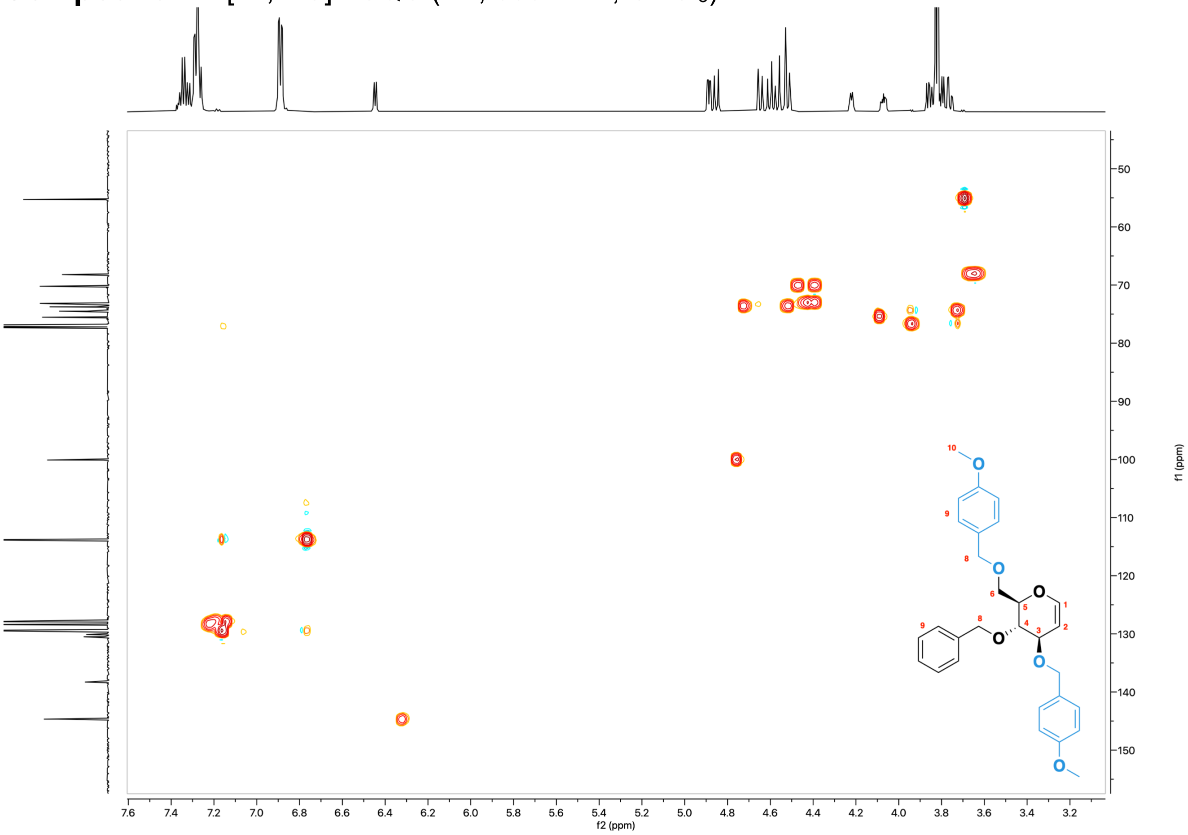

#### Monomer 4 - $^{1}$H NMR (600 MHz, $CDCl_3$)

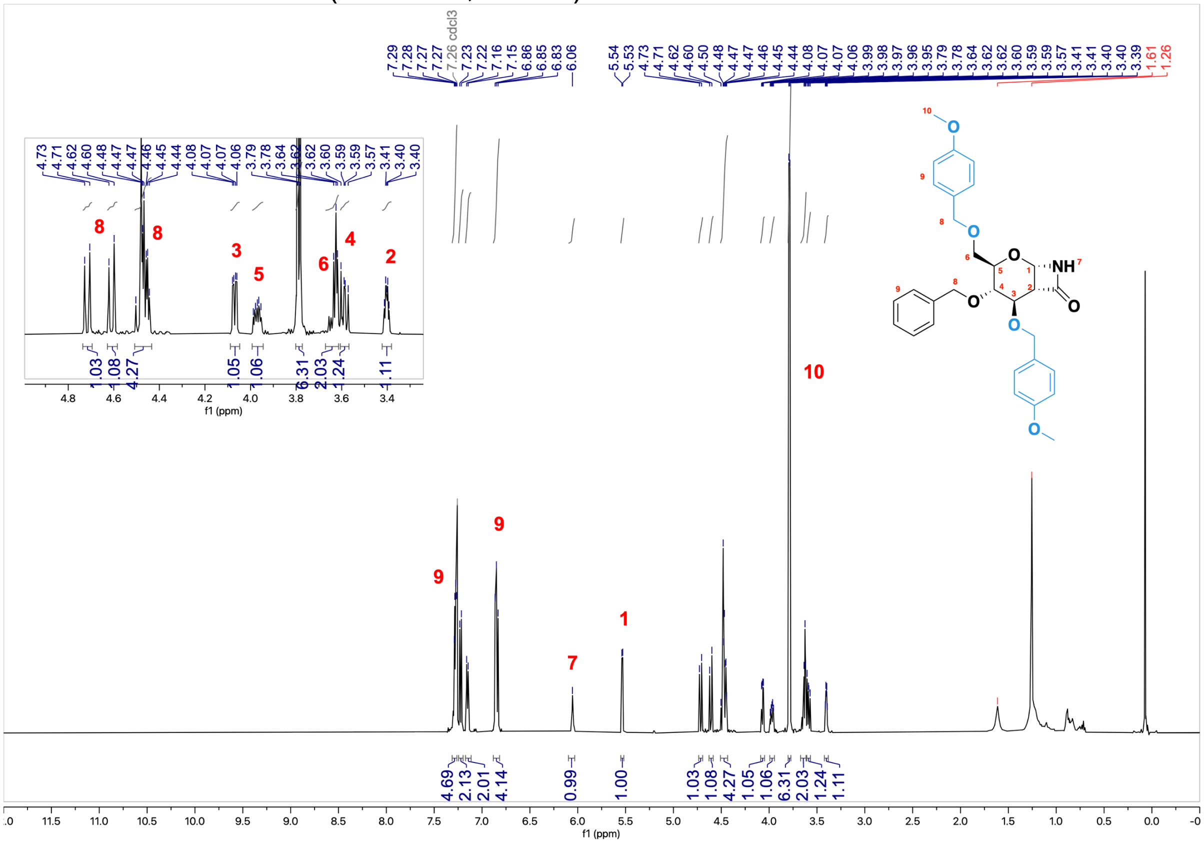

#### Monomer 4 - $^{13}$C NMR (151 MHz, $CDCl_3$)

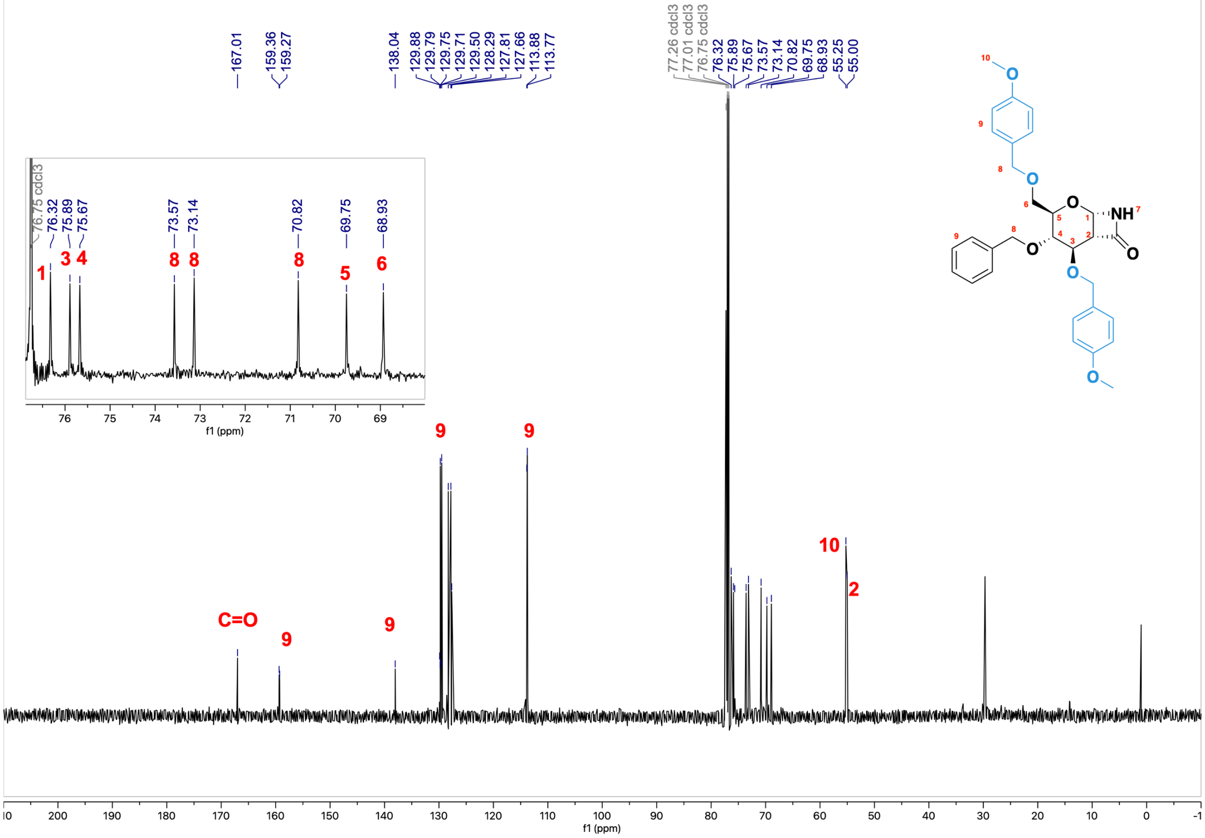

**Monomer 4** - [$^1$H, $^1$H] COSY (2D, 600 MHz, $CDCl_3$)

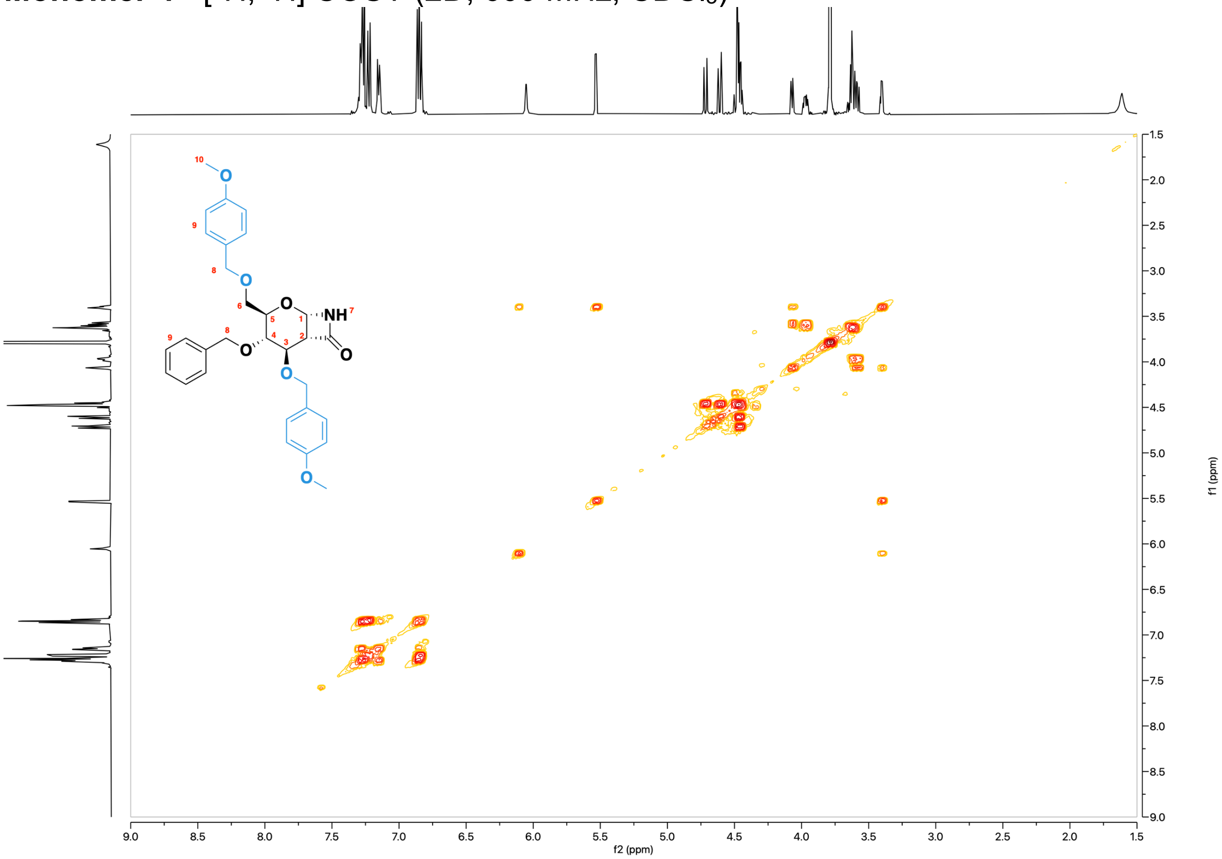


**Monomer 4** - [$^1$H, $^{13}$C] HSQC (2D, 151 MHz, $CDCl_3$)

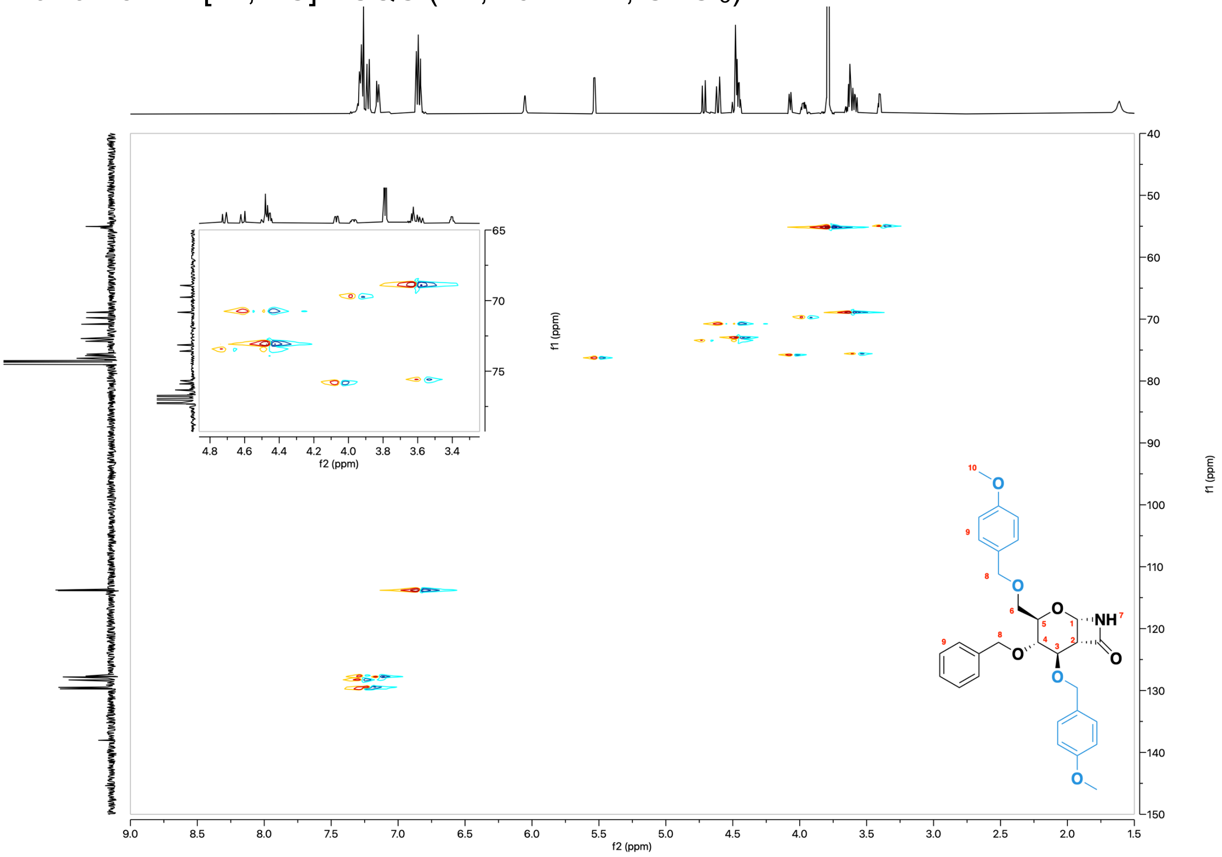

**Compound 8** - $^{1}$H NMR (600 MHz, $CDCl_3$)

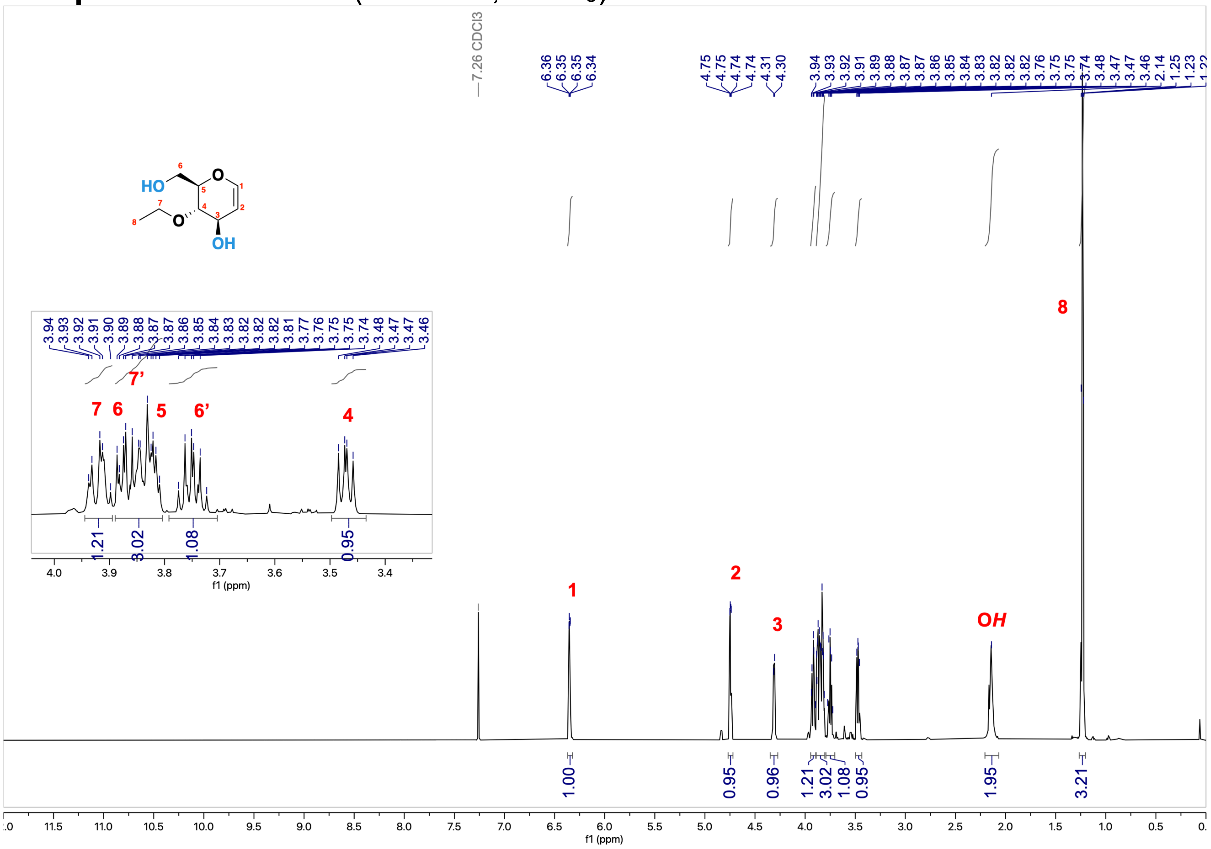

**Compound 8** - $^{13}$C NMR (151 MHz, $CDCl_3$)

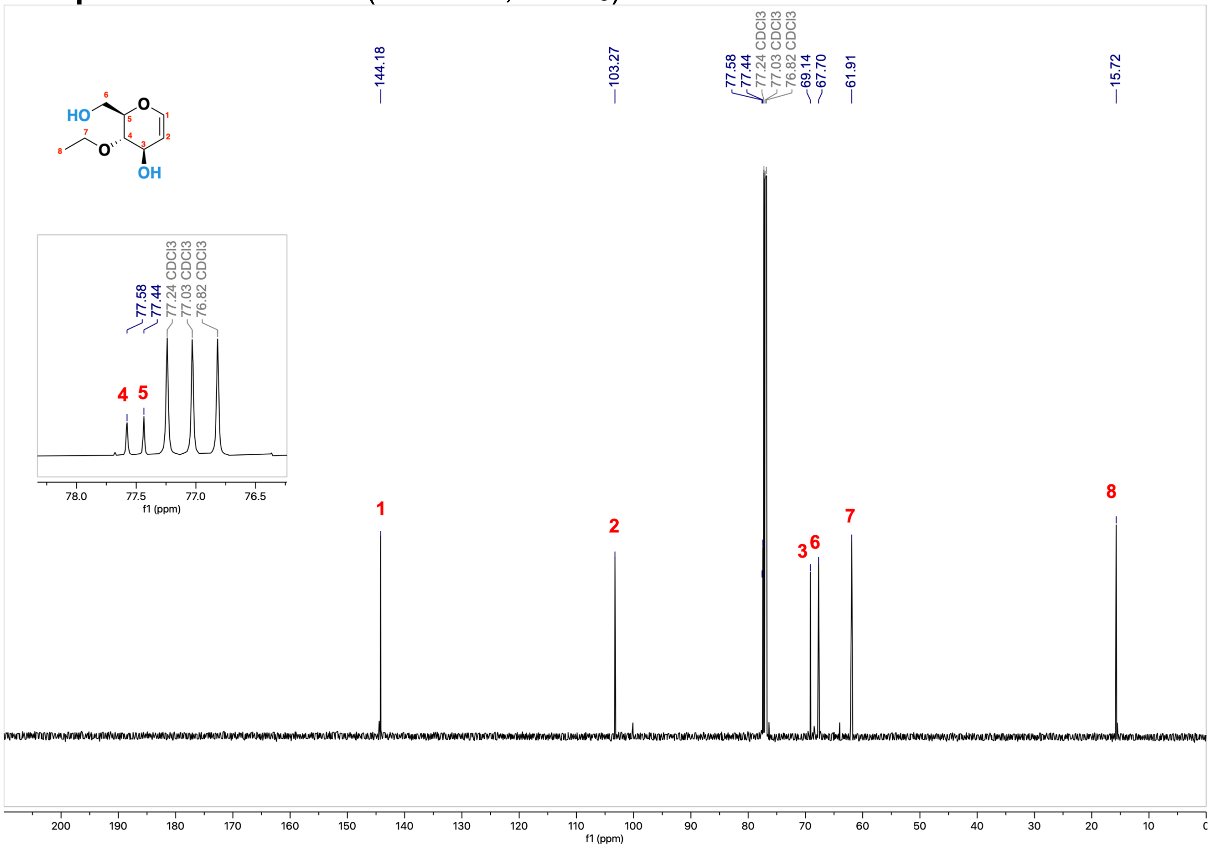

**Compound 8** - [1H, 1H] COSY (2D, 600 MHz, $CDCl_3$)

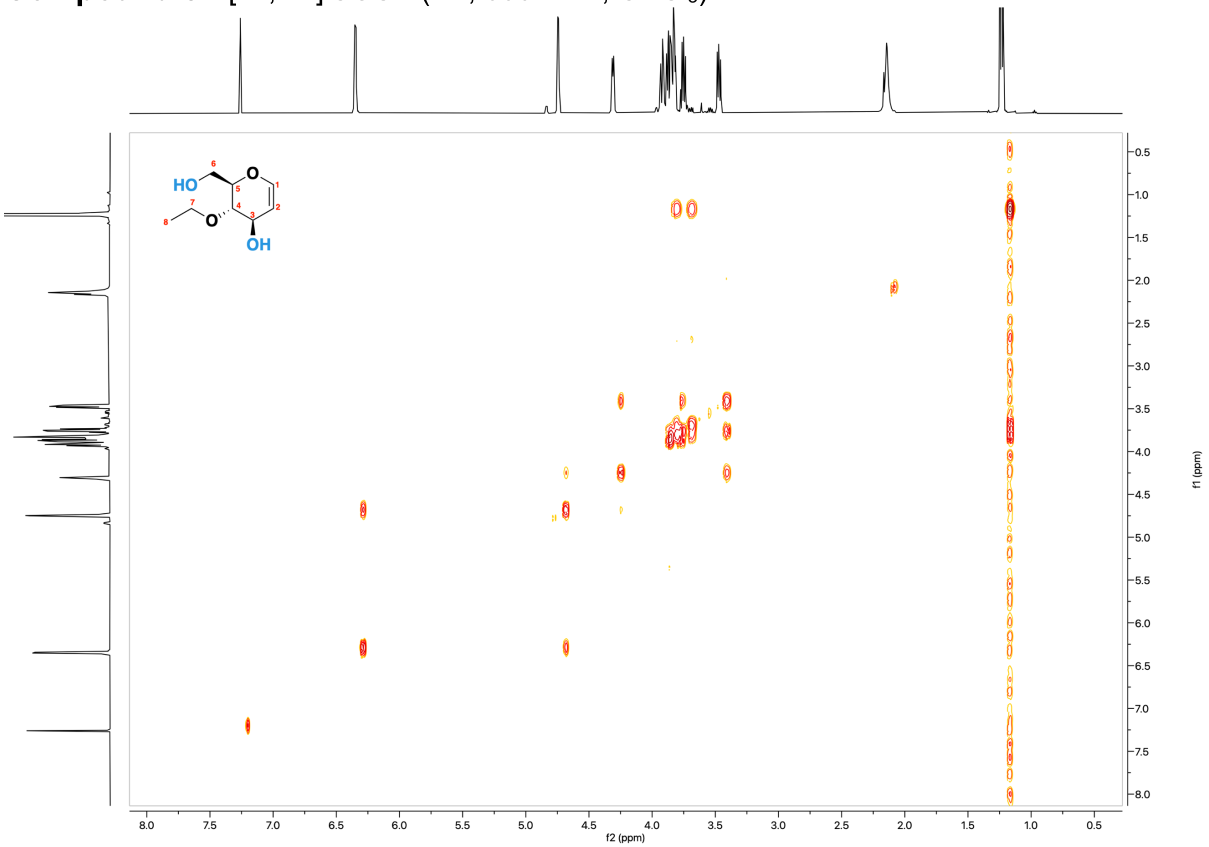


**Compound 8** - [1H, 13C] HSQC (2D, 600 MHz, $CDCl_3$)

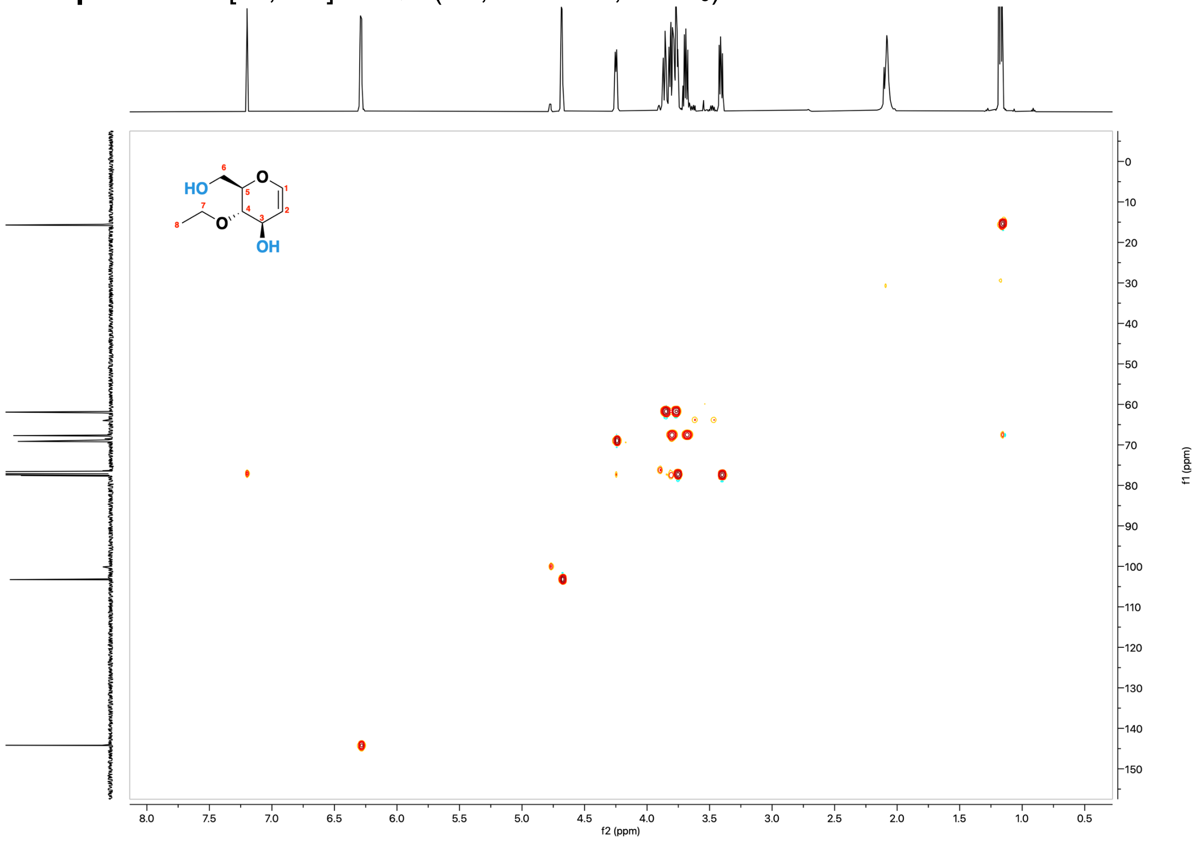

**Compound 9** - $^{1}$H NMR (600 MHz, $CDCl_3$)

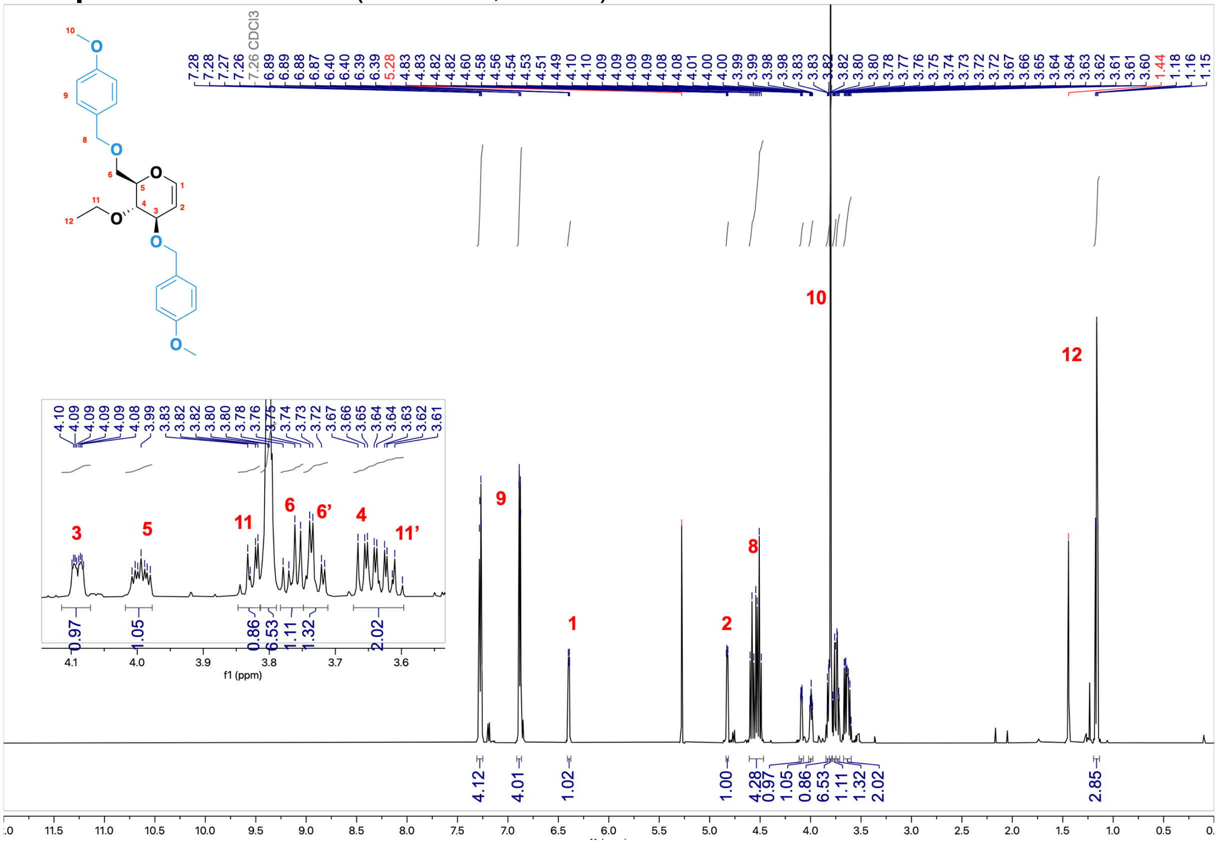

**Compound 9** - $^{13}$C NMR (151 MHz, $CDCl_3$)

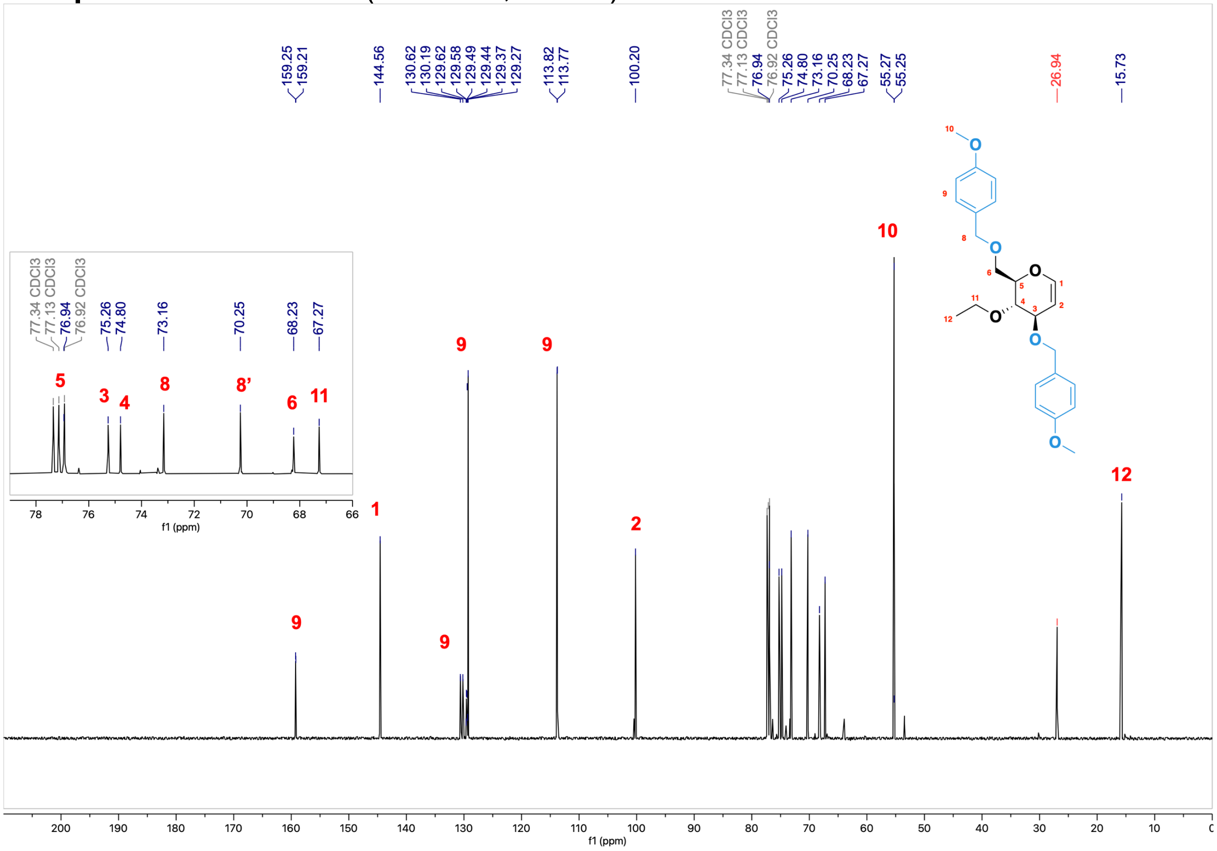

**Compound 9** - [[1]H, [1]H] COSY (2D, 600 MHz, $CDCl_3$)

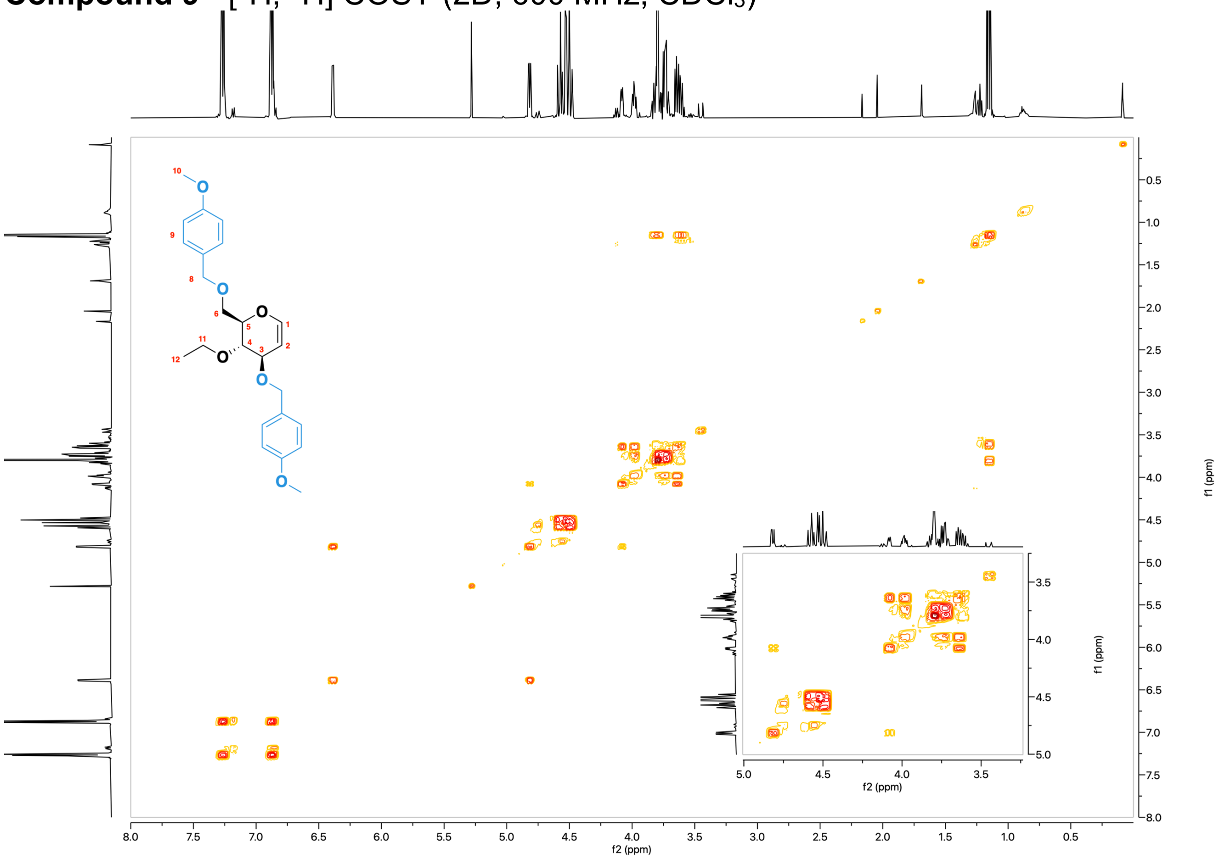


**Compound 9** - [[1]H, [13]C] HSQC (2D, 151 MHz, $CDCl_3$)

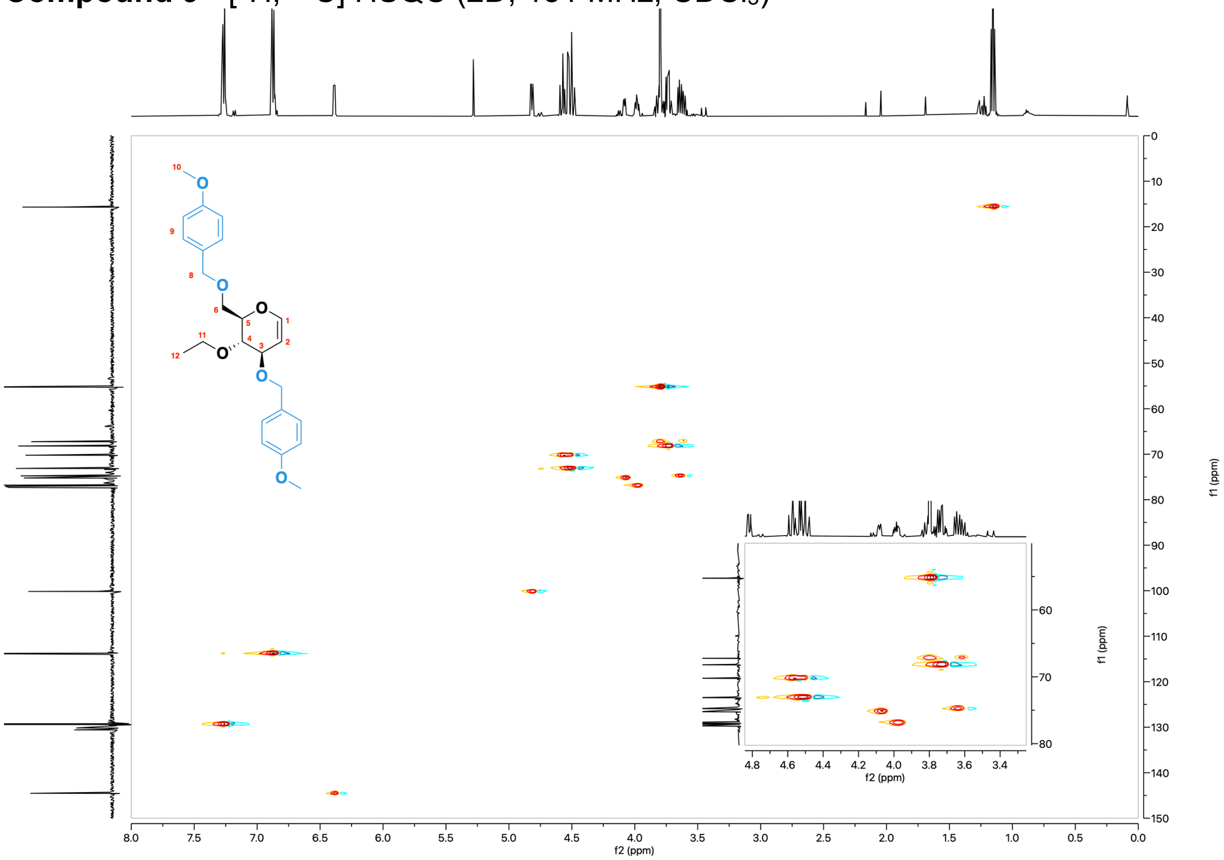

**Monomer 5** - $^{1}$H NMR (600 MHz, $CDCl_3$)

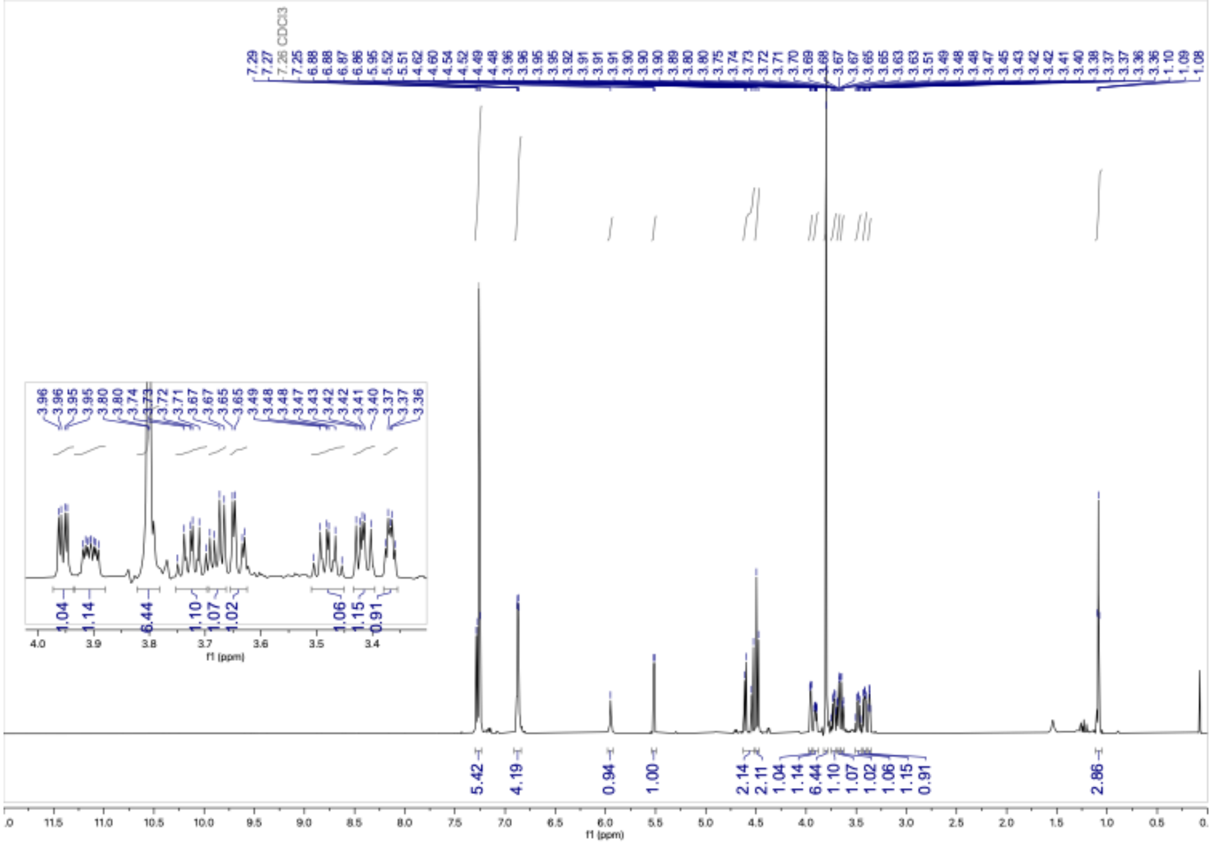

**Monomer 5** - $^{13}$C NMR (151 MHz, $CDCl_3$)

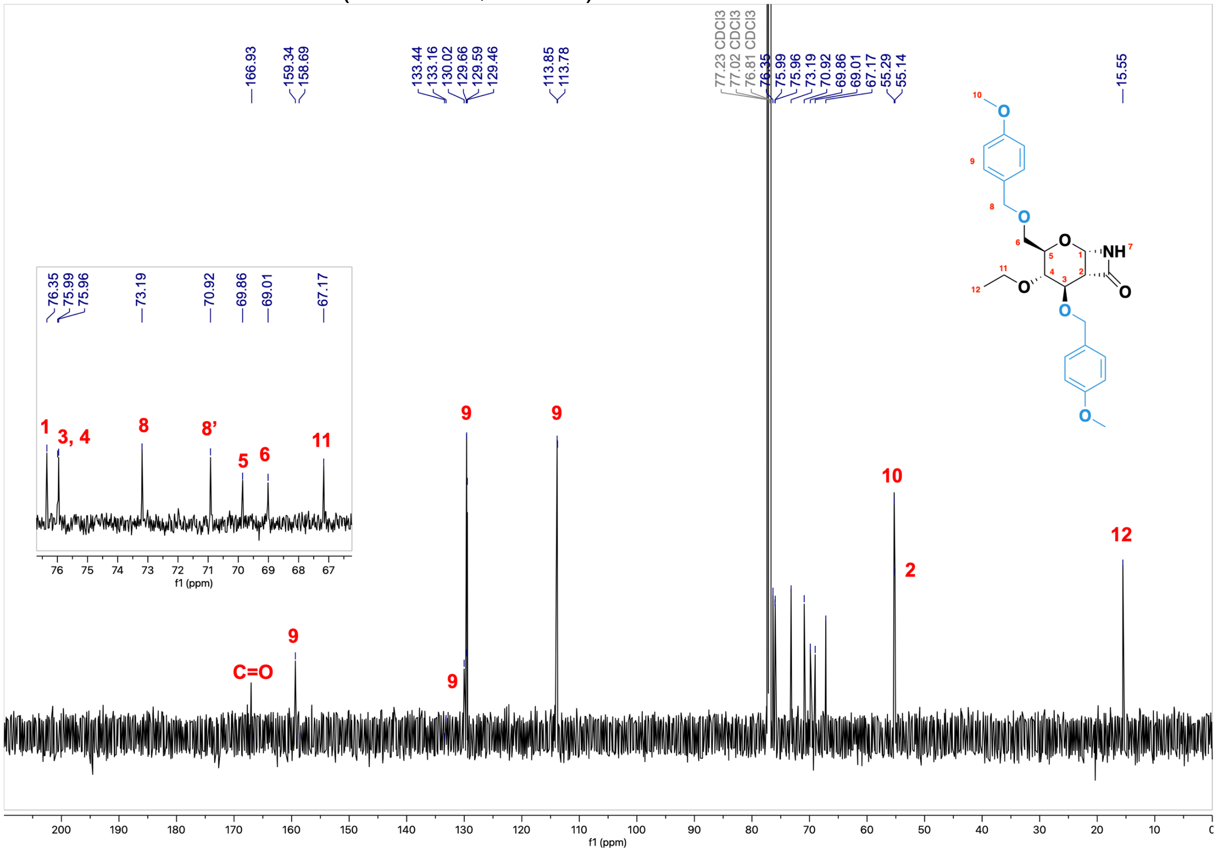

**Monomer 5** - [$^1$H, $^1$H] COSY (2D, 600 MHz, $CDCl_3$)

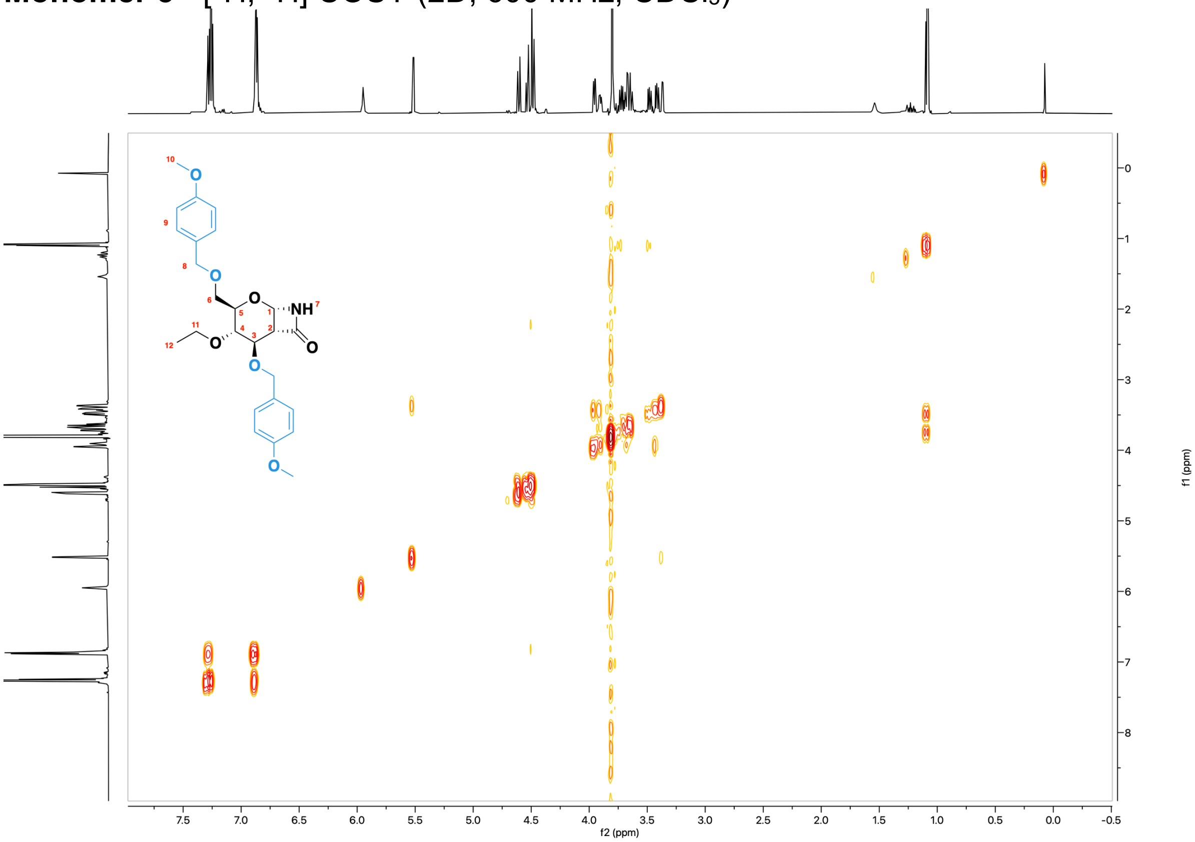


**Monomer 5** - [$^1$H, $^{13}$C] HSQC (2D, 151 MHz, $CDCl_3$)

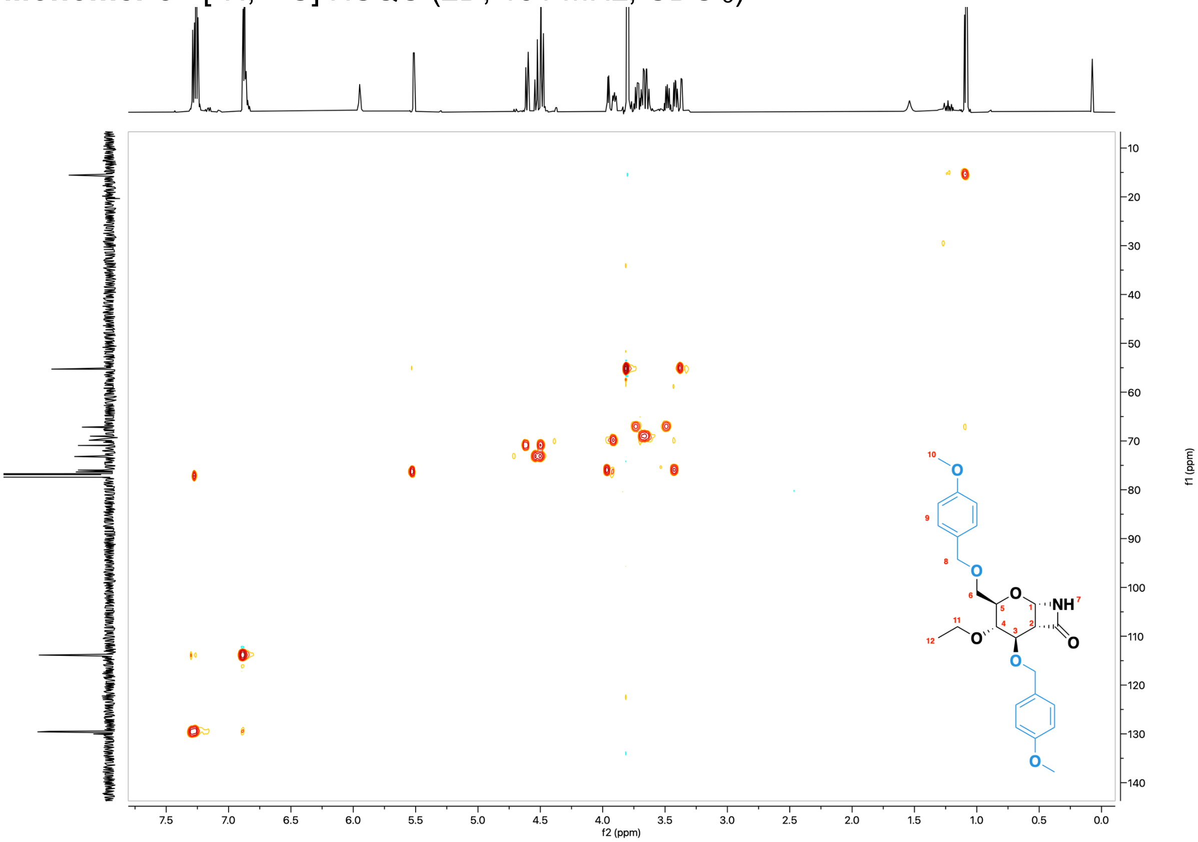

**Compound 10** - $^{1}$H NMR (600 MHz, $CDCl_3$)

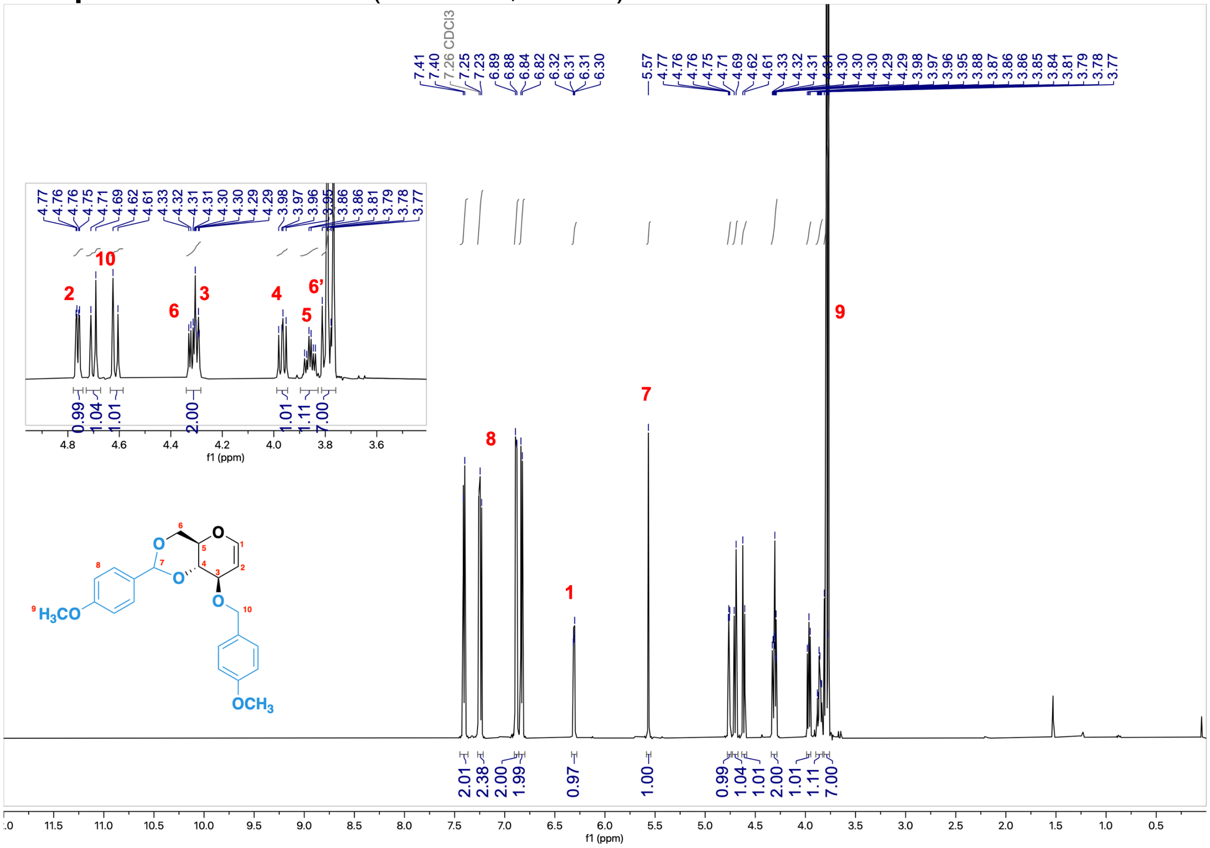


**Compound 10** - $^{13}$C NMR (151 MHz, $CDCl_3$)

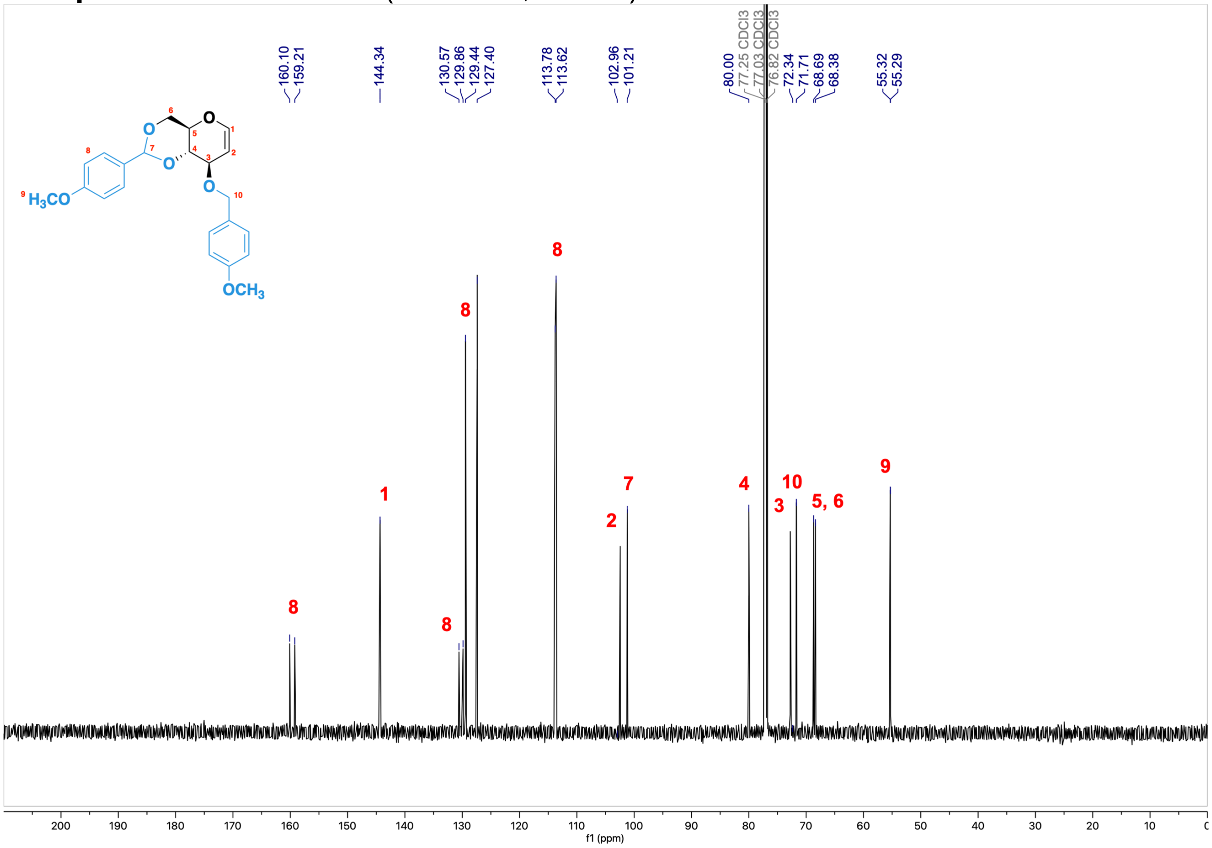

**Compound 11** - $^{1}H$ NMR (600 MHz, $CDCl_3$)

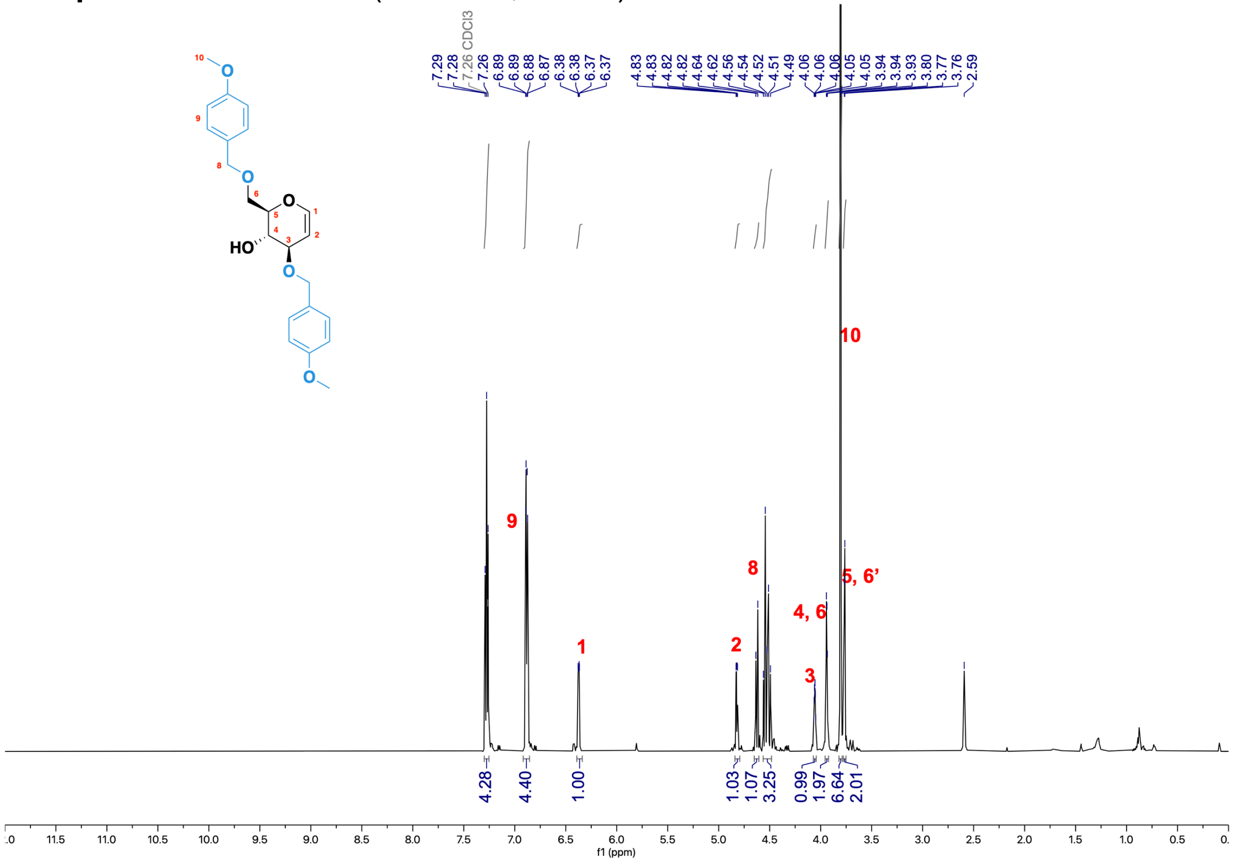

**Compound 11** - $^{13}C$ NMR (151 MHz, $CDCl_3$)

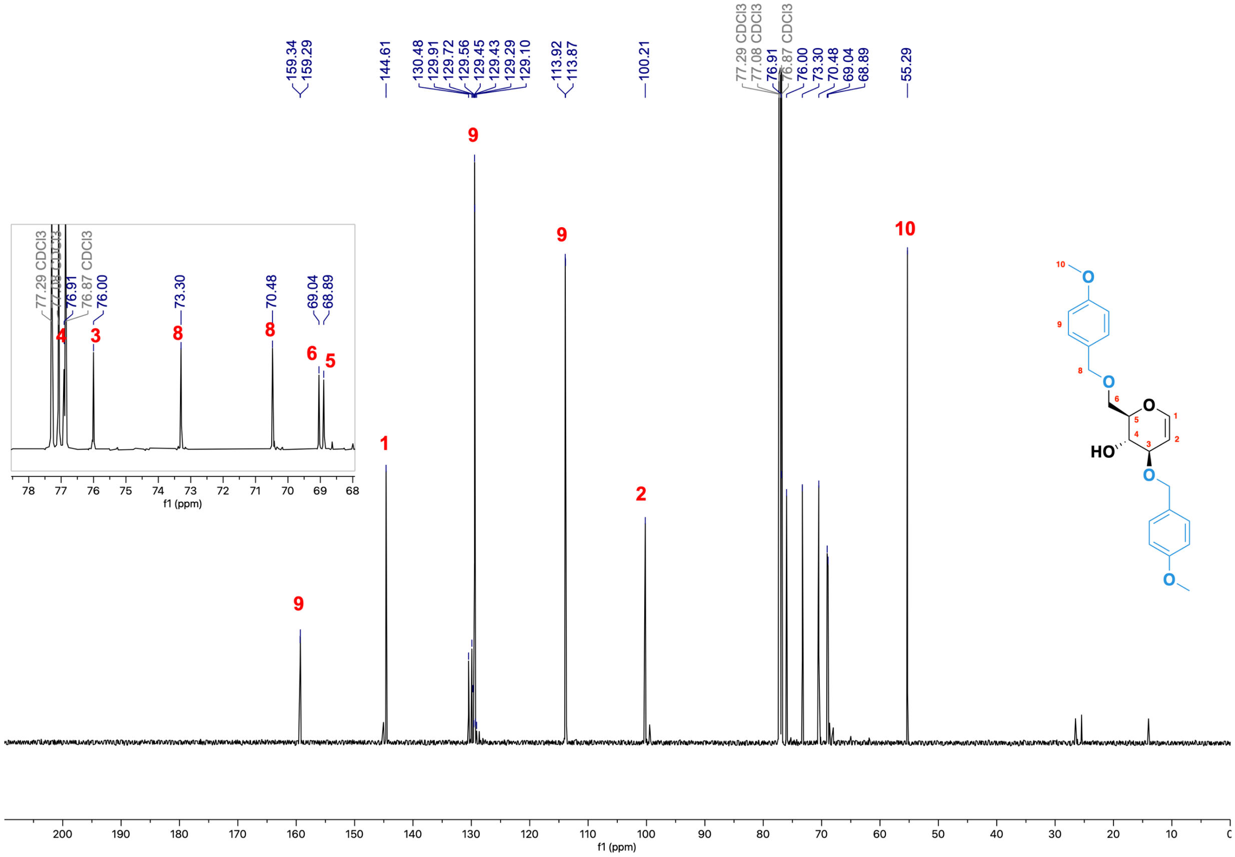

**Compound 11** - [¹H, ¹H] COSY (2D, 600 MHz, $CDCl_3$)

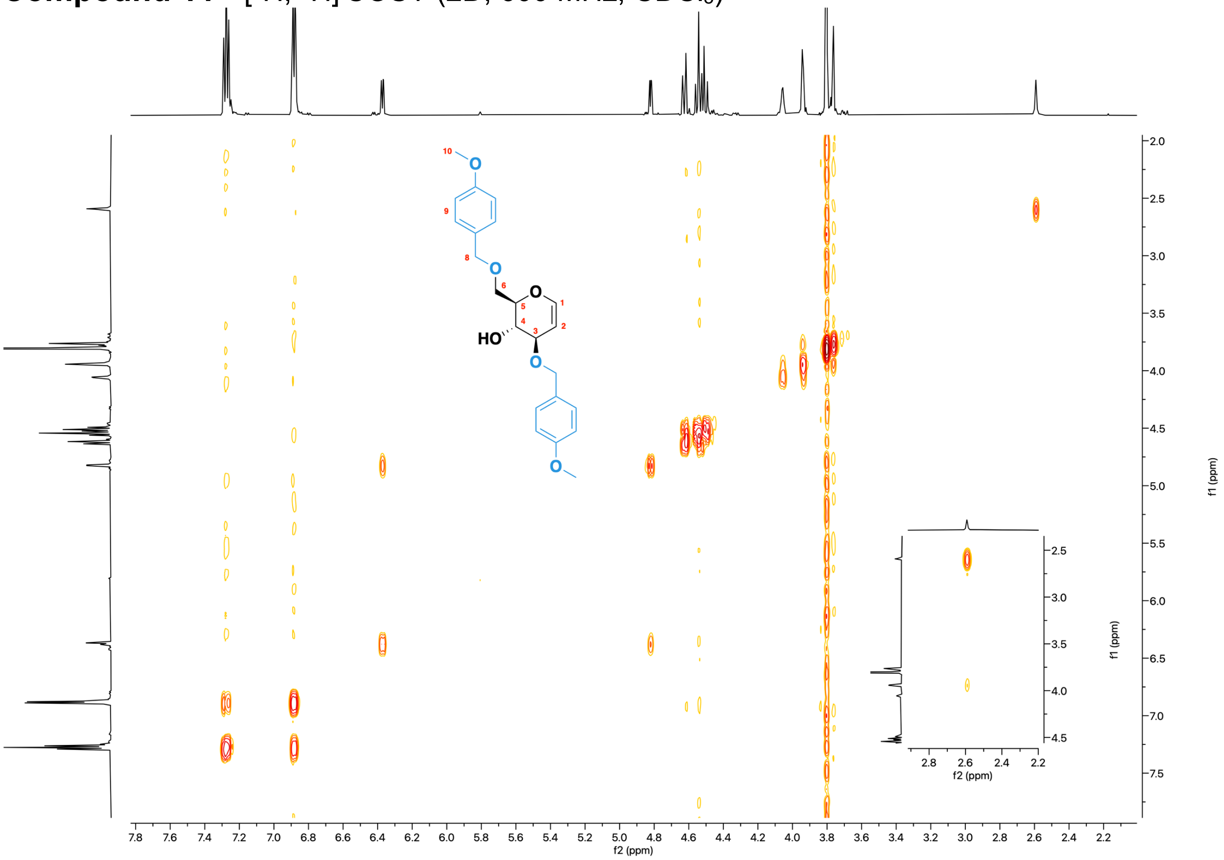

**Compound 11** - [¹H, ¹³C] HSQC (2D, 600 MHz, $CDCl_3$)

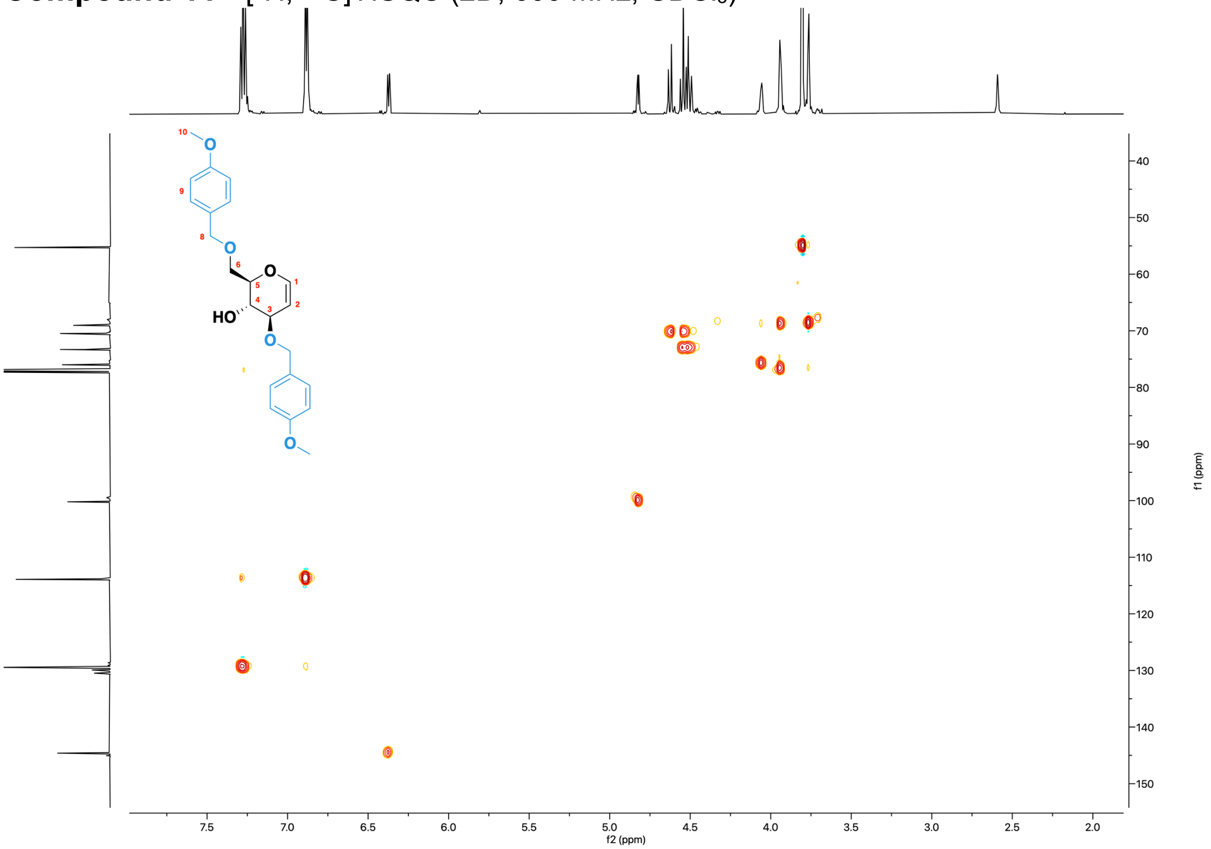

#### Compound 12 - $^{1}$H NMR (600 MHz, $CDCl_3$)

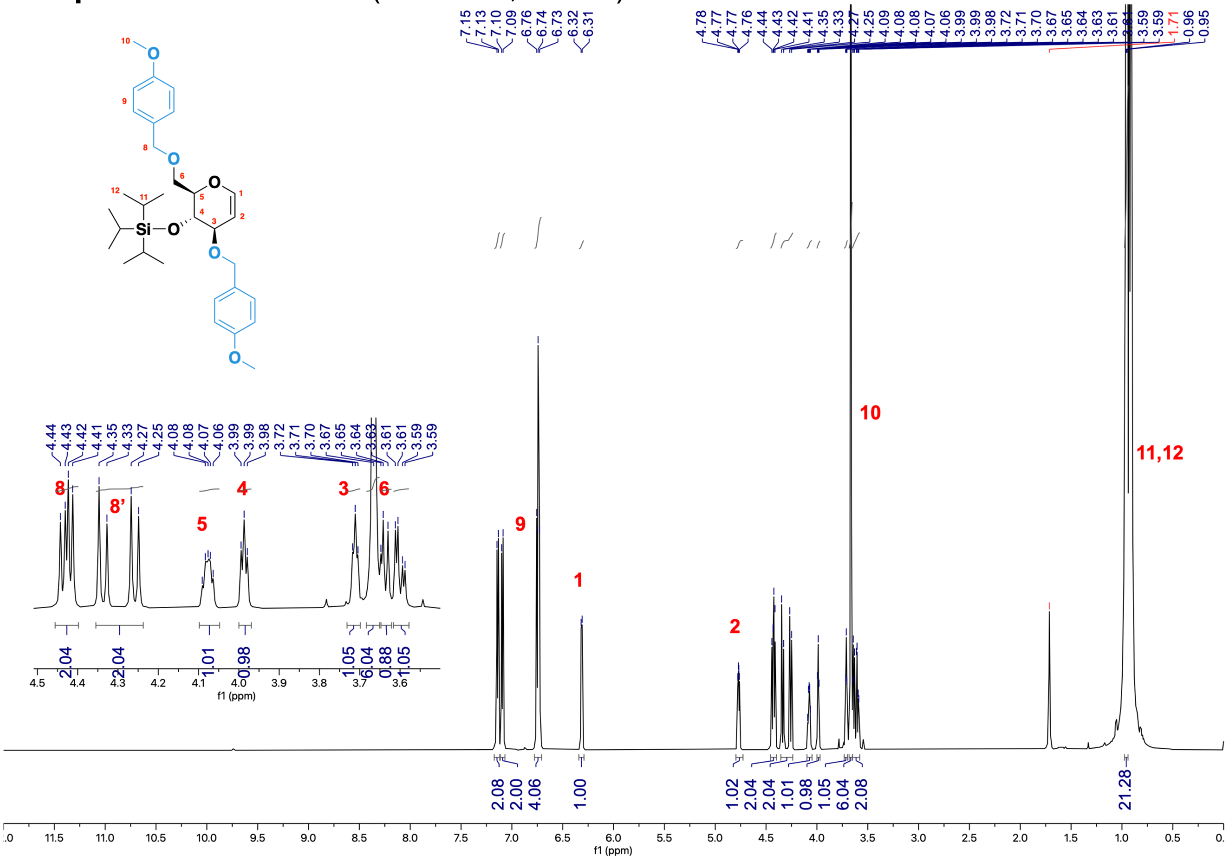

#### Compound 12 - $^{13}$C NMR (151 MHz, $CDCl_3$)

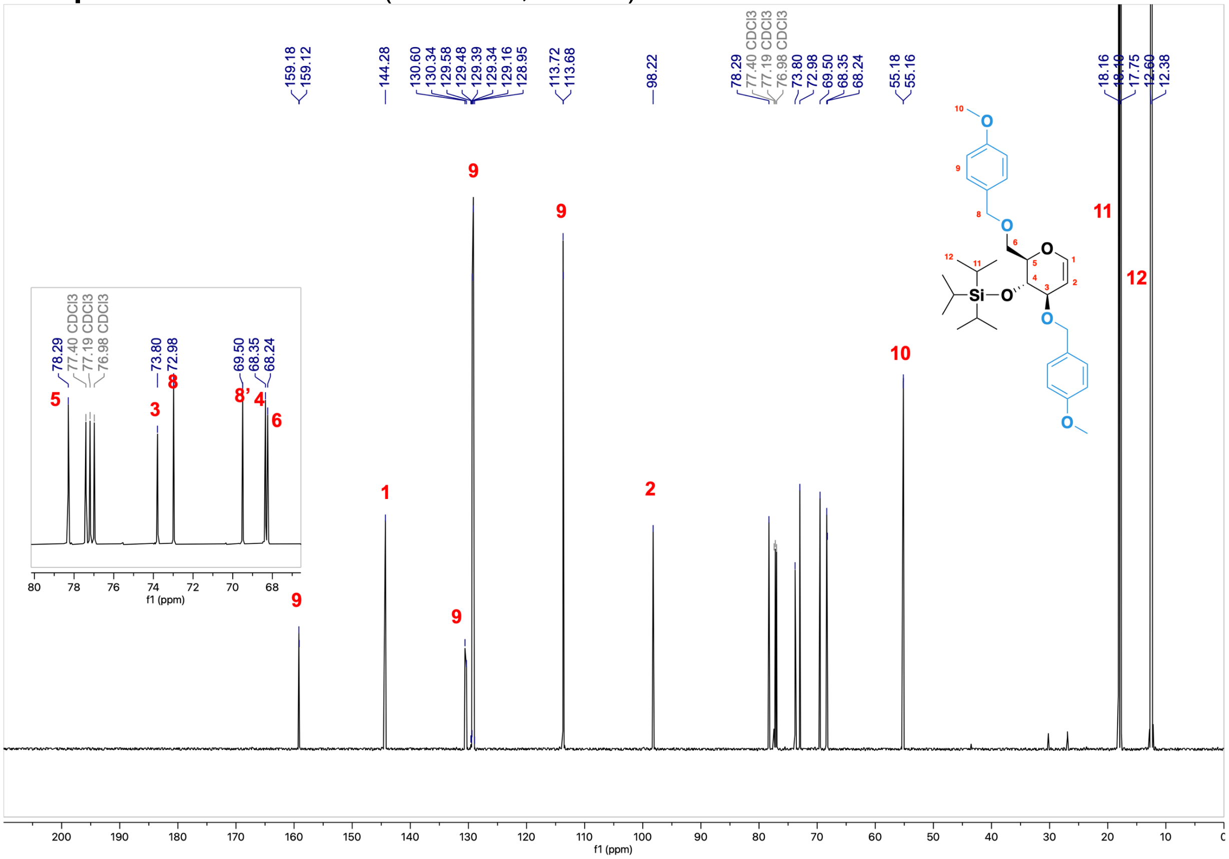

#### **Compound 12** - [$^{1}$H, $^{1}$H] COSY (2D, 600 MHz, $CDCl_3$)

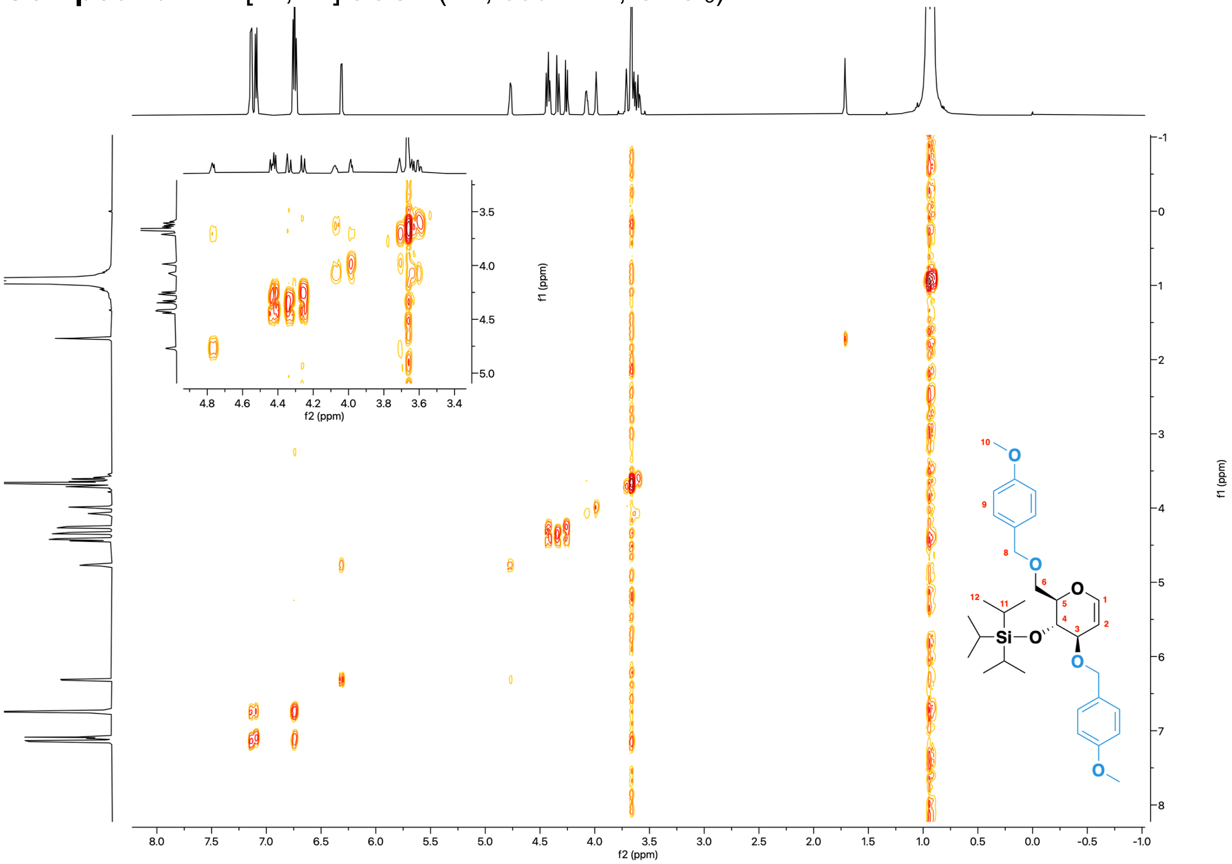


#### **Compound 12** - [$^{1}$H, $^{13}$C] HSQC (2D, 600 MHz, $CDCl_3$)

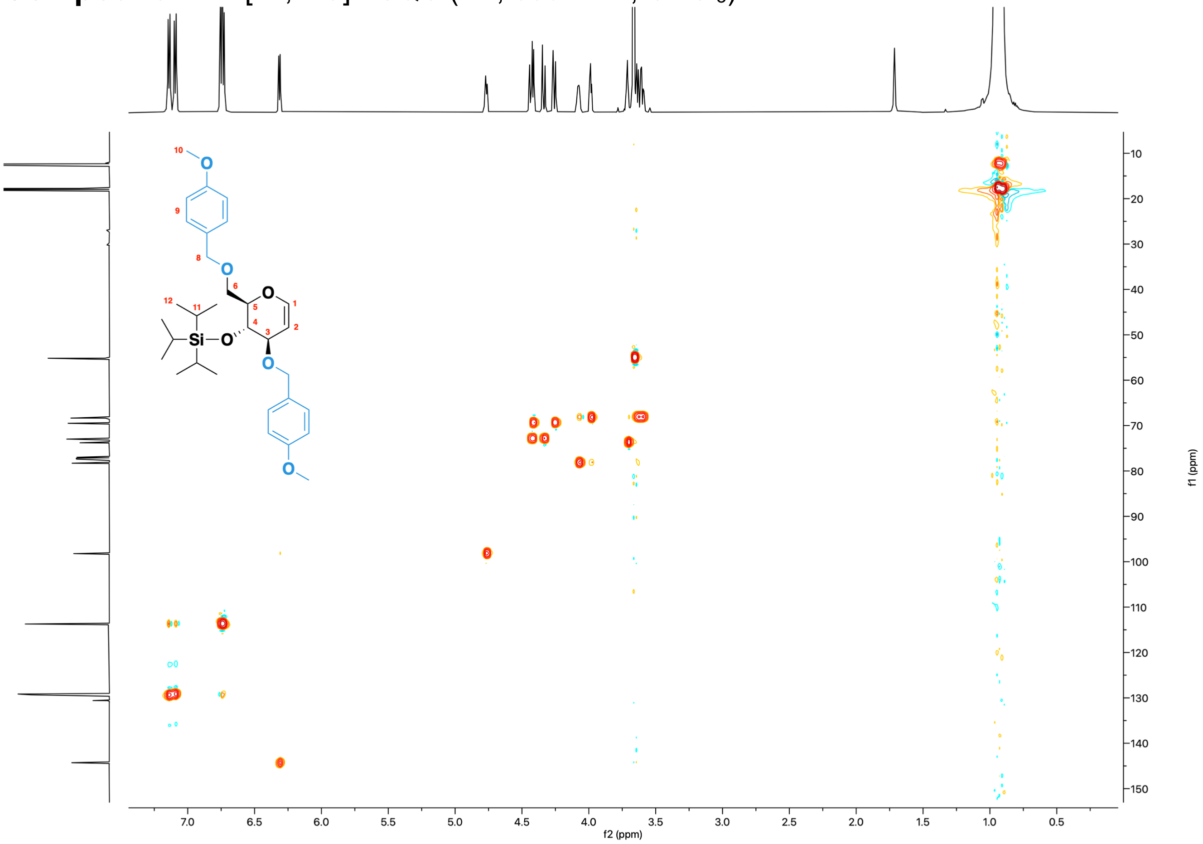

#### Monomer 6 - $^{1}$H NMR (600 MHz, $CDCl_3$)

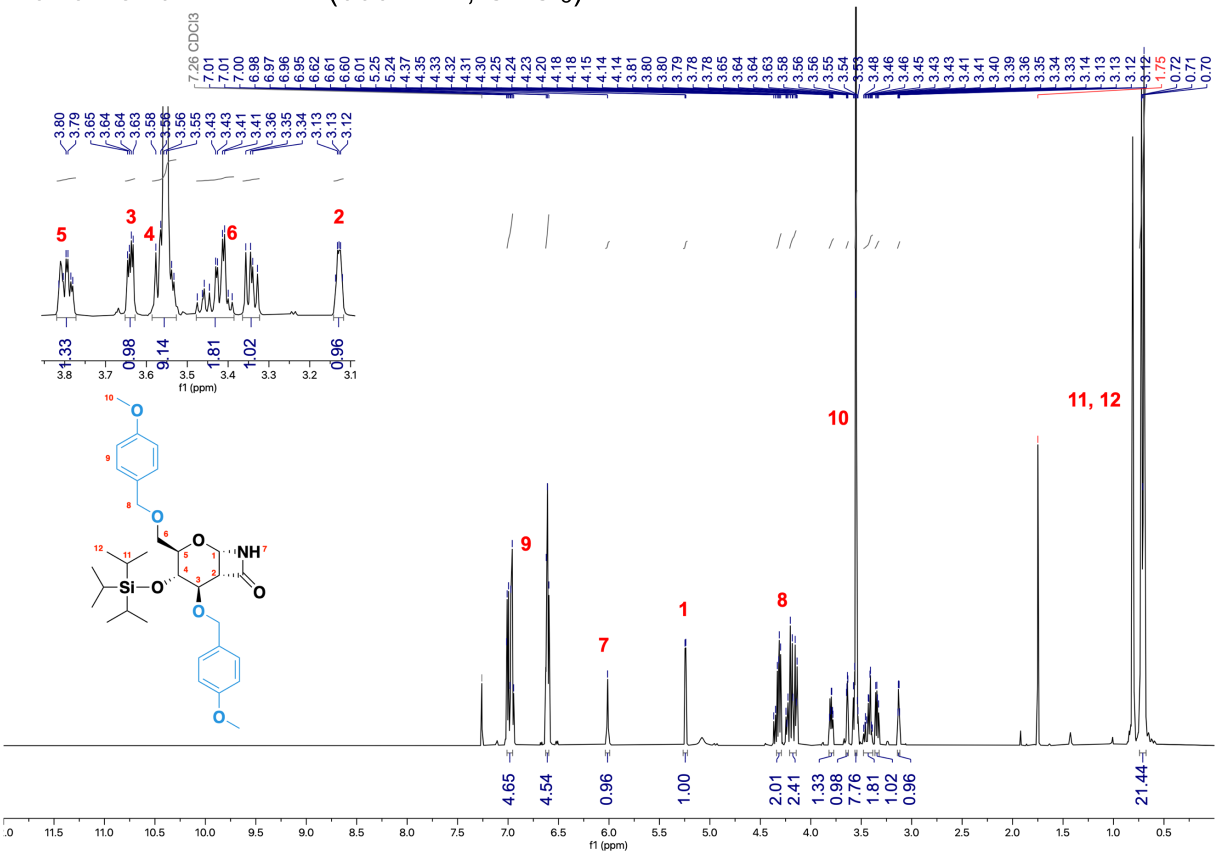


#### Monomer 6 - $^{13}$C NMR (151 MHz, $CDCl_3$)

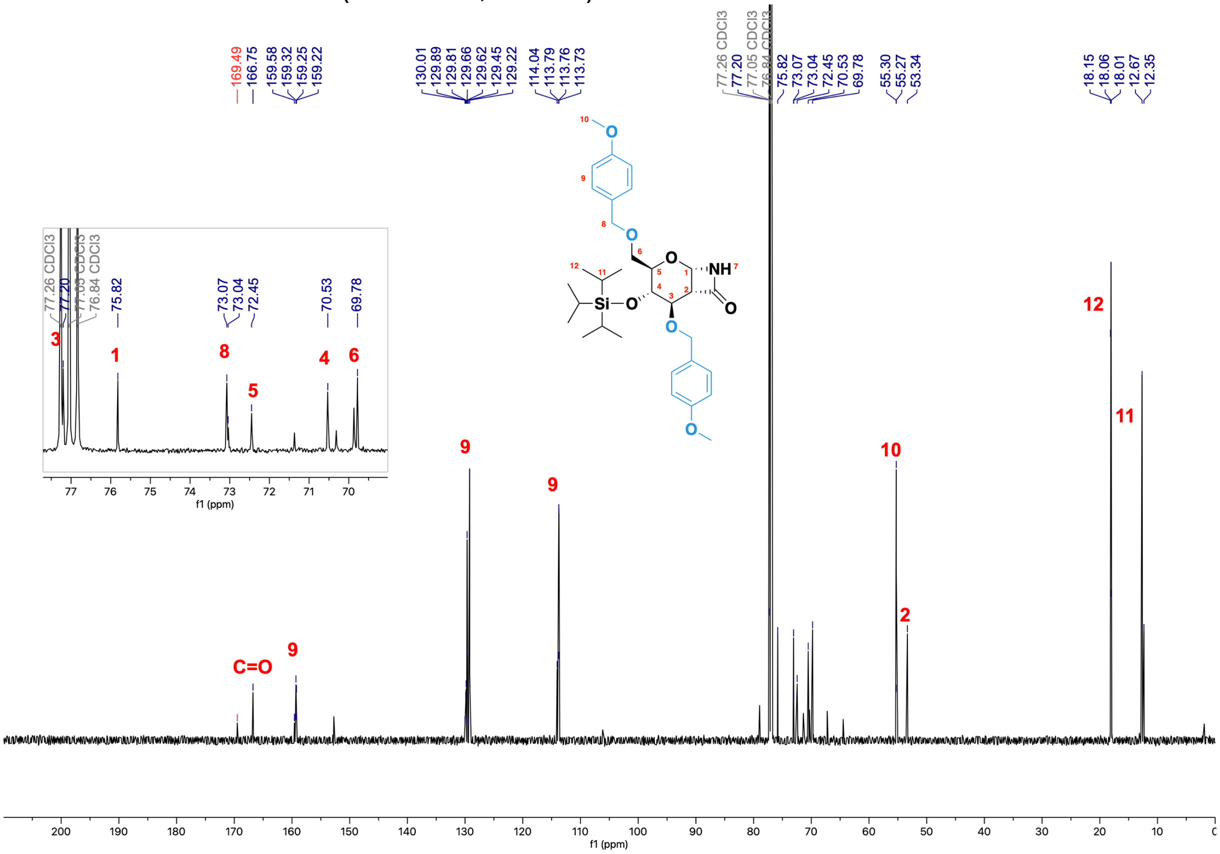

**Monomer 6** - [1H, 1H] COSY (2D, 600 MHz, $CDCl_3$)

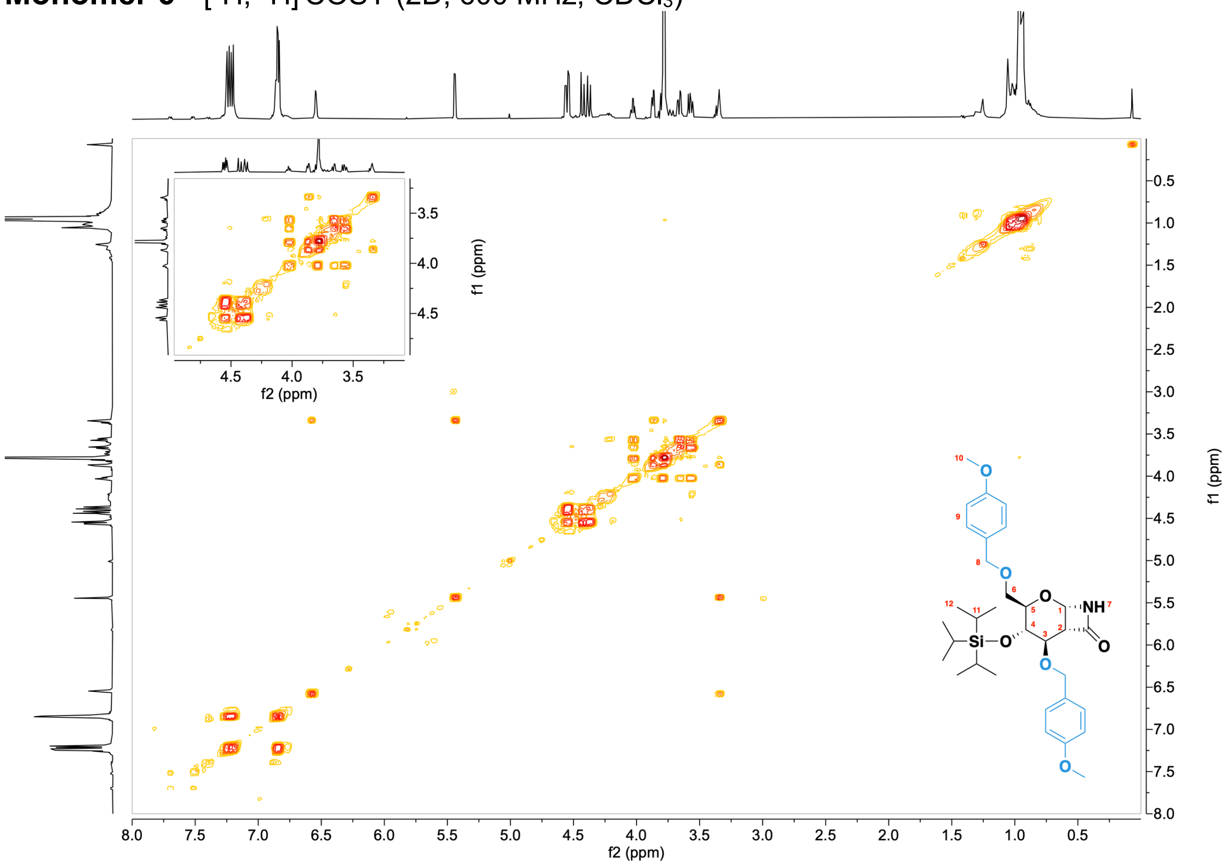

**Monomer 6** - [1H, 13C] HSQC (2D, 600 MHz, $CDCl_3$)

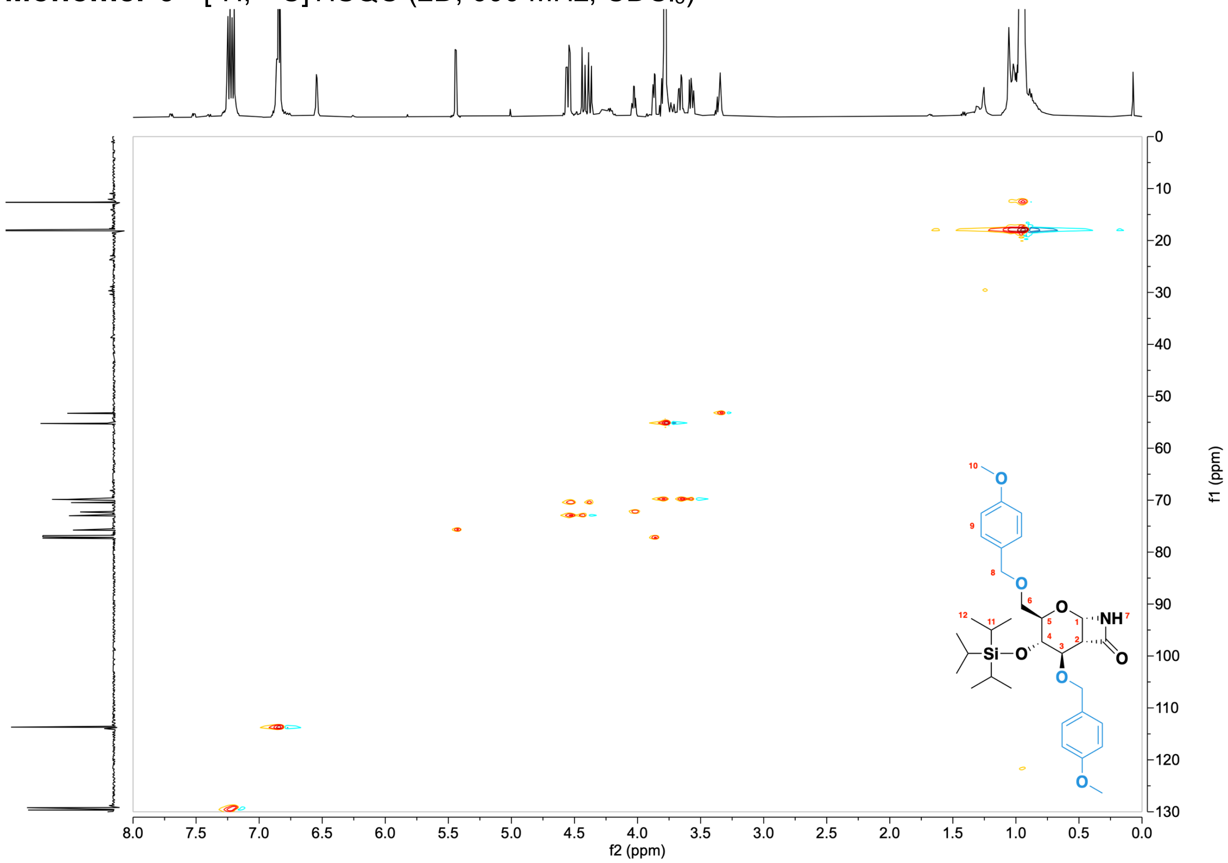

### Polymer NMRs

[1]H NMR of **3,6-di-*O*-PMB-4-*O*-TIPS-PAS.**

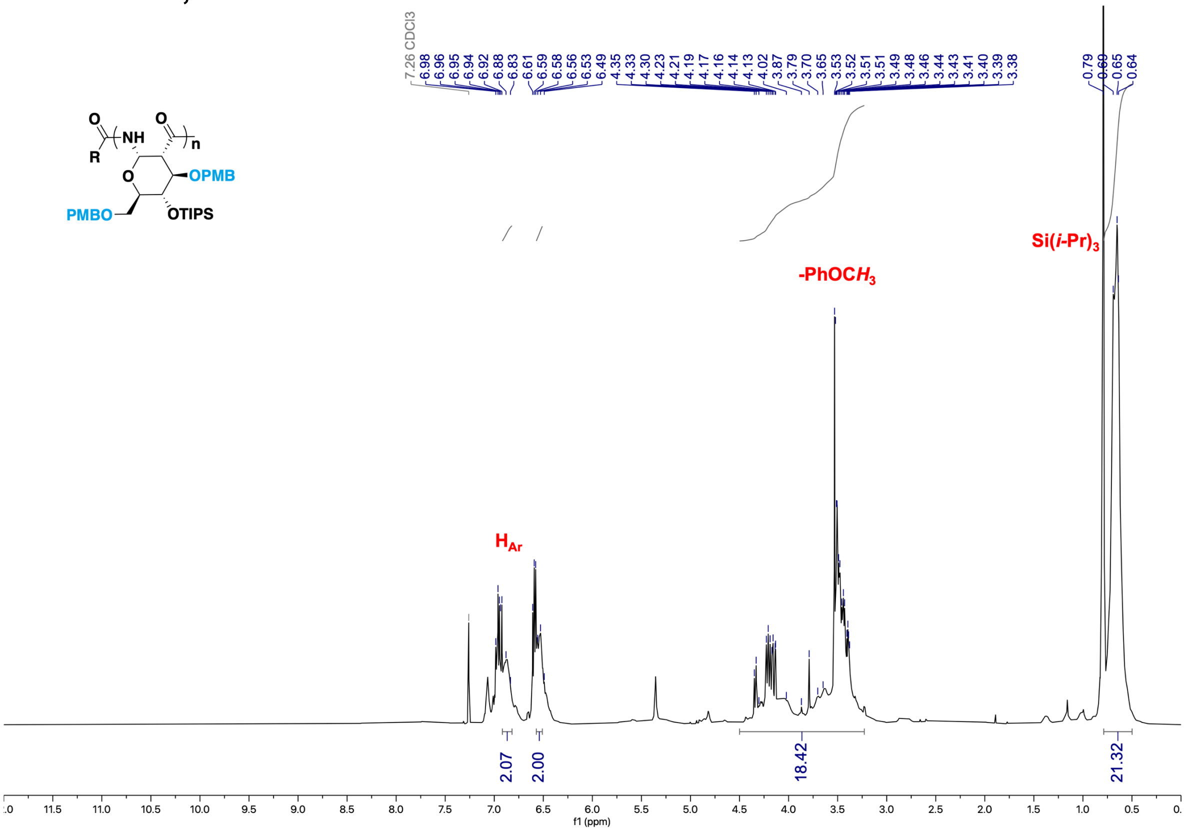


[13]C NMR of **3,6-di-*O*-PMB-4-*O*-TIPS-PAS.**

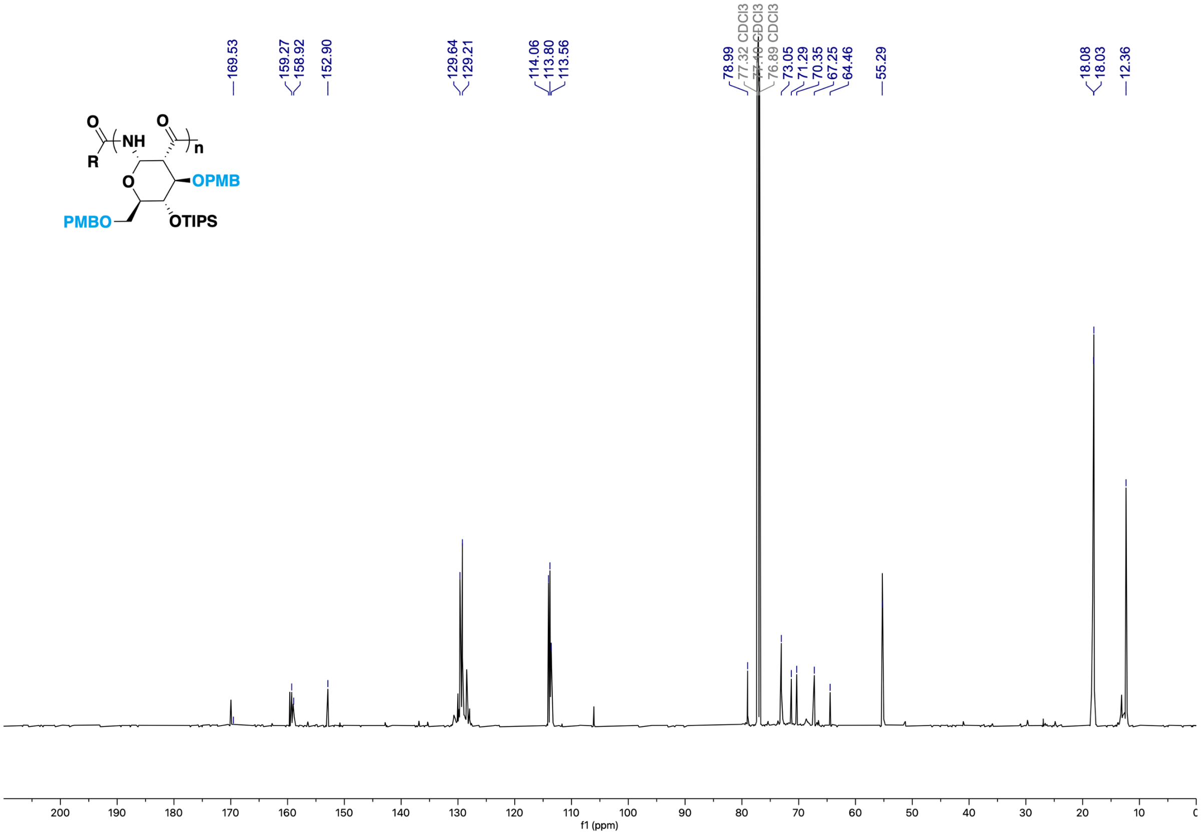

[1]H NMR of **3,6-di-*O*-PMB-4-*O*-Bn-PAS**.

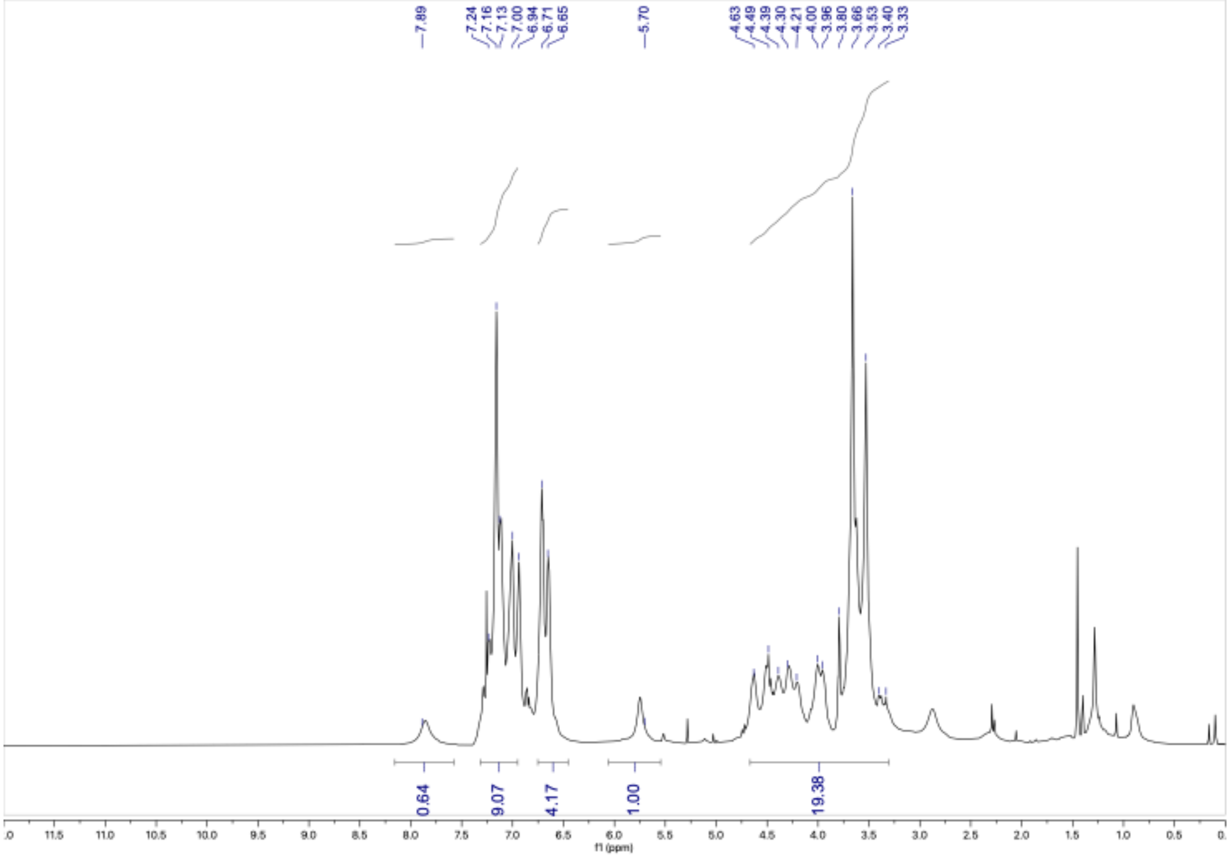


[13]C NMR of **3,6-di-*O*-PMB-4-*O*-Bn-PAS**.

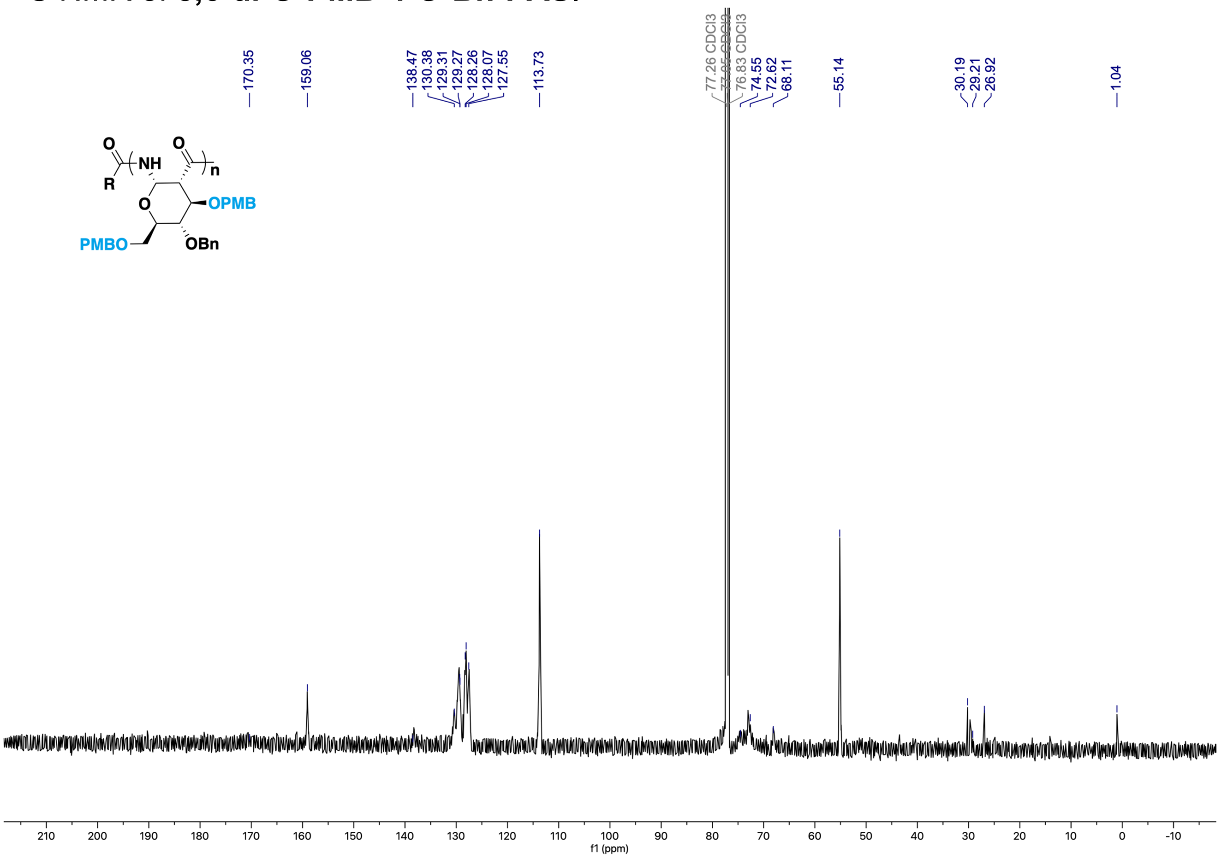

$^1$H NMR of **3,6-di-*O*-PMB-4-*O*-Et-PAS.**

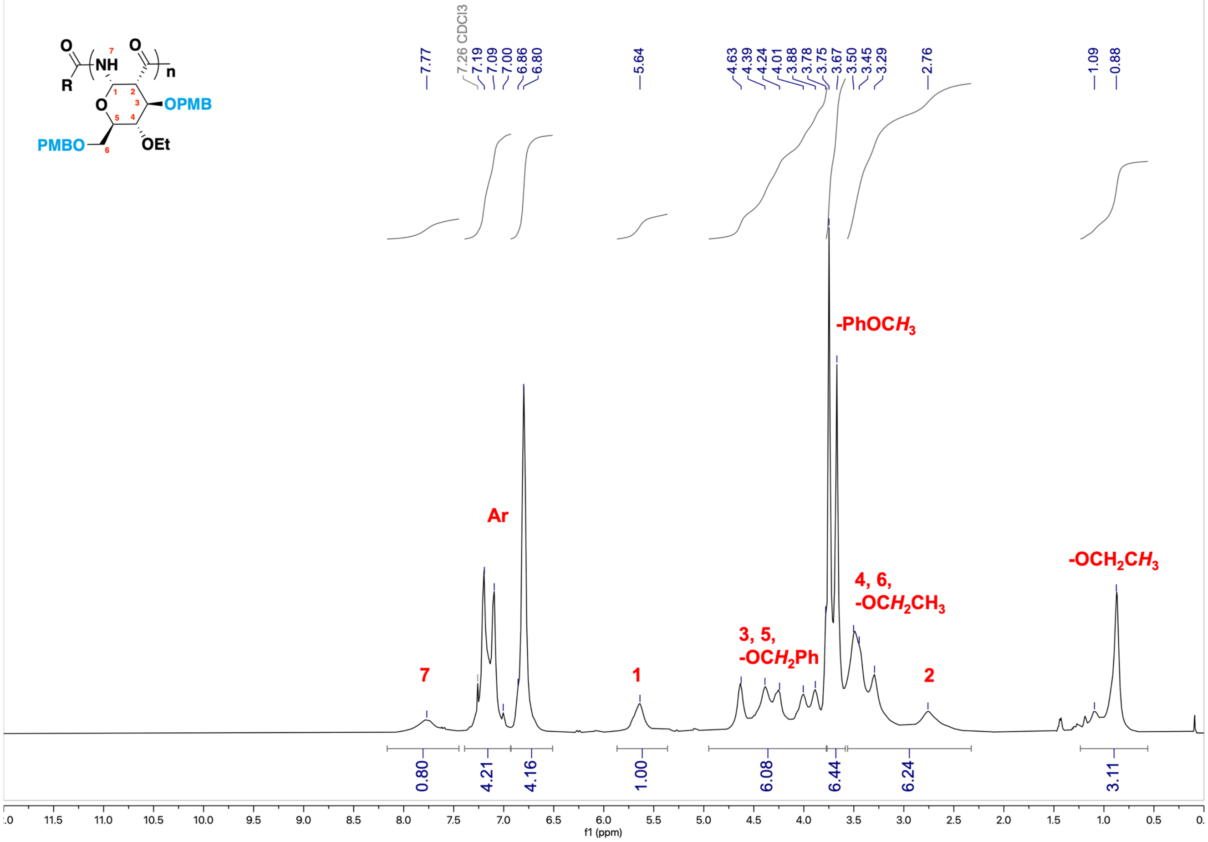


$^{13}$C NMR of **3,6-di-*O*-PMB-4-*O*-Et-PAS.**

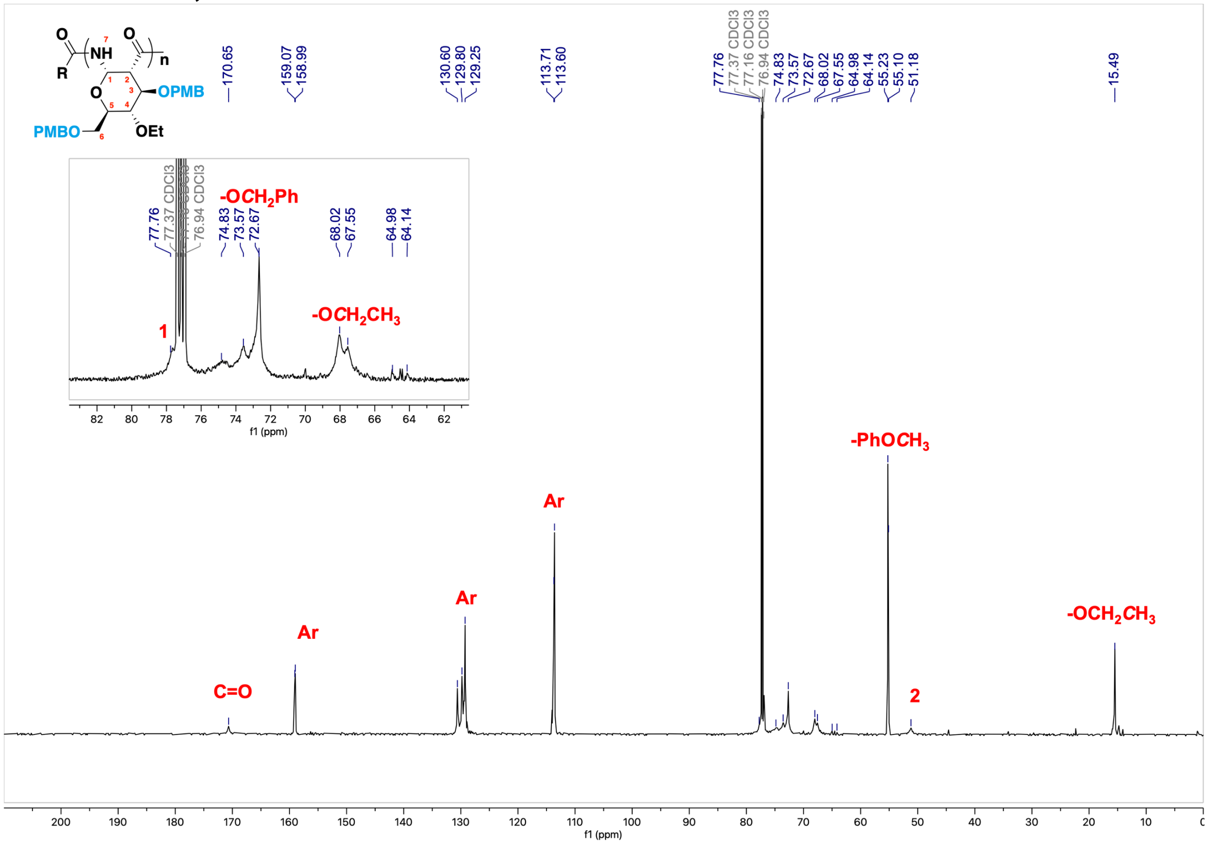

**3,6-di-*O*-PMB-4-*O*-Et-PAS** - [1H, 1H] COSY (2D, 600 MHz, $CDCl_3$)

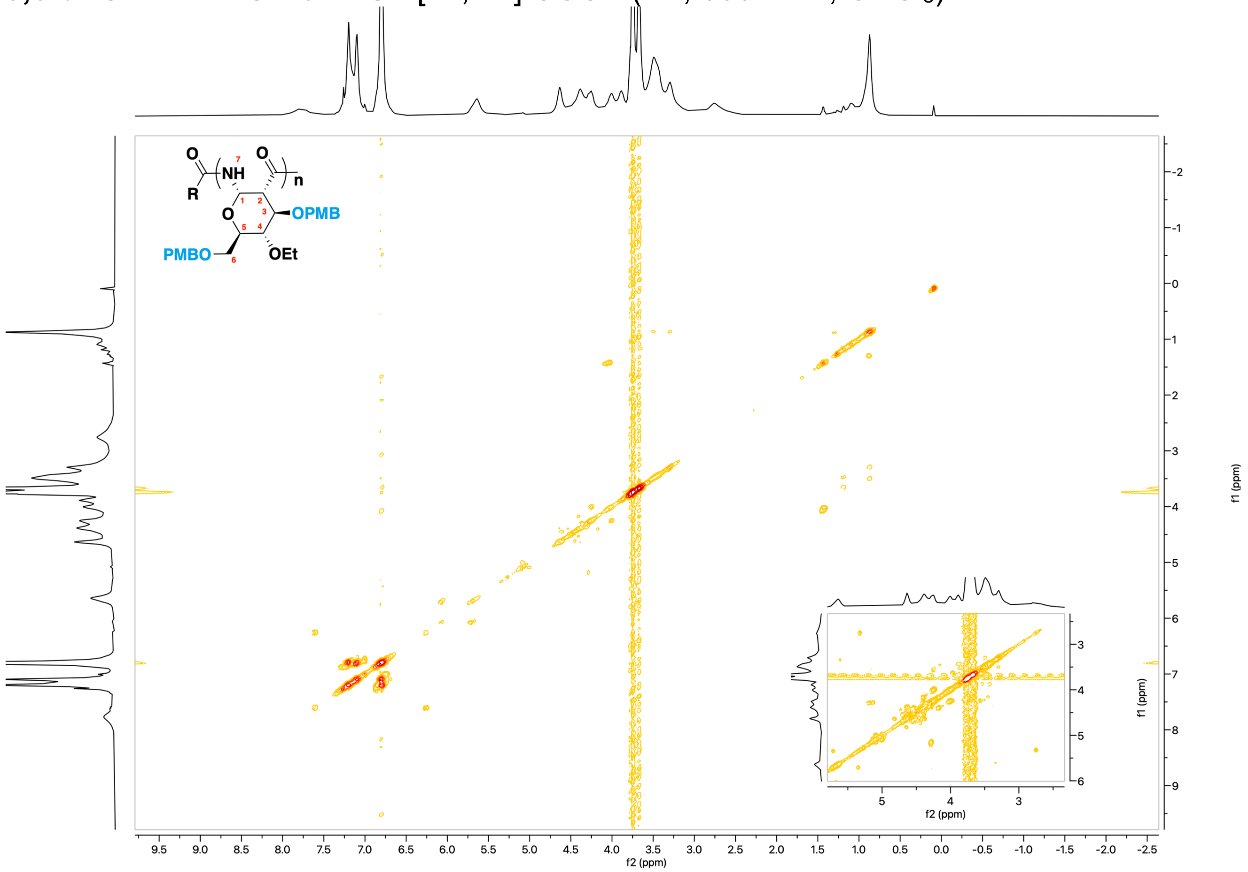


**3,6-di-*O*-PMB-4-*O*-Et-PAS** - [1H, 13C] HSQC (2D, 151 MHz, $CDCl_3$)

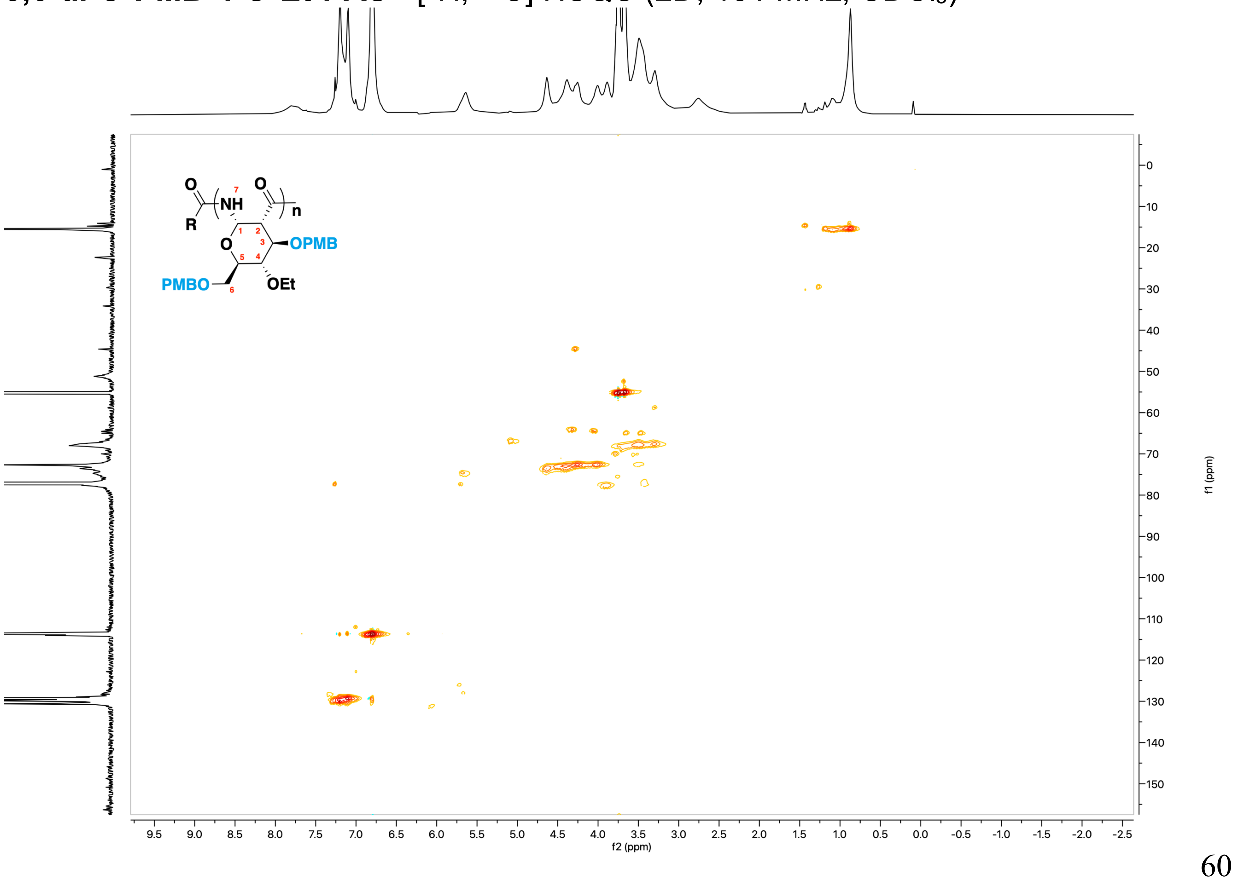

$^{1}$H NMR of **3,6-di-OH-4-*O*-Bn-PAS.**

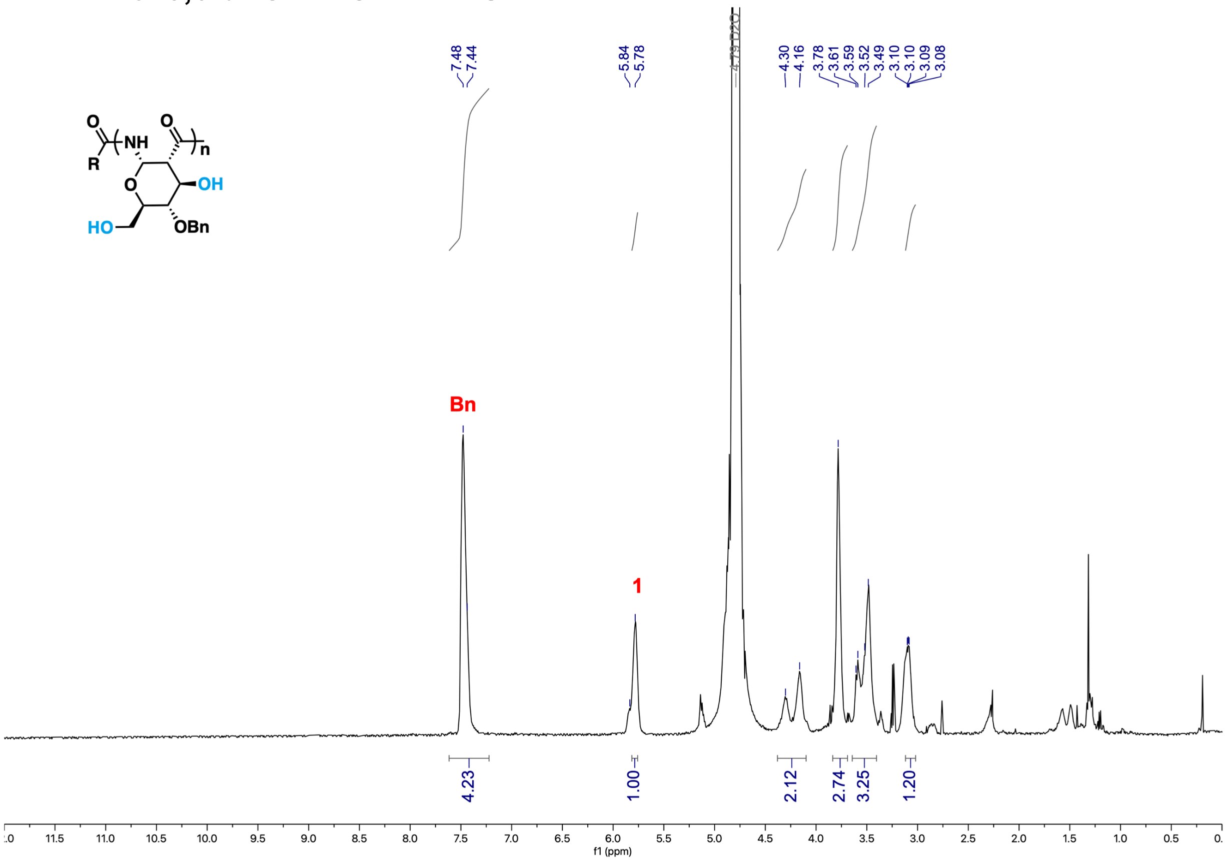

$^{13}$C NMR of **3,6-di-OH-4-*O*-Bn-PAS.**

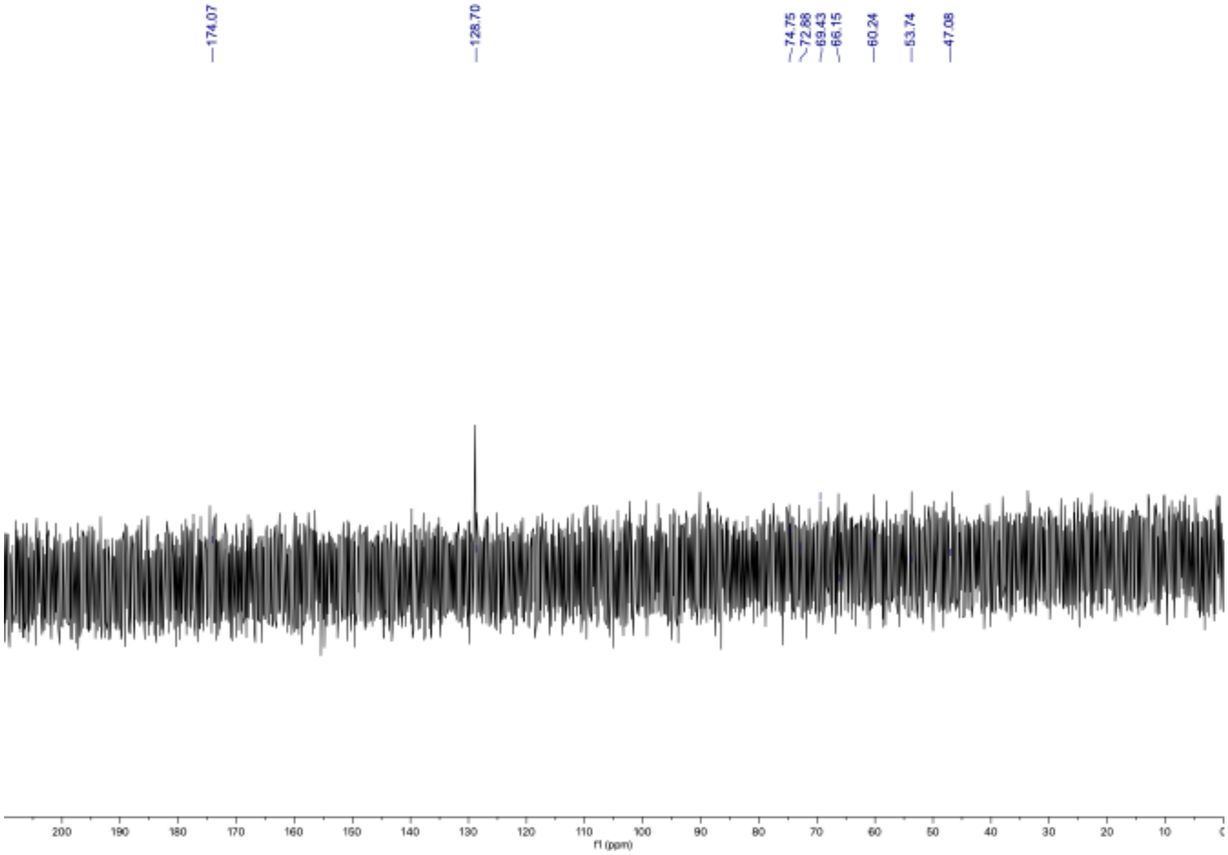

$^{1}$H NMR of **3,6-di-OH-4-*O*-Et-PAS.**

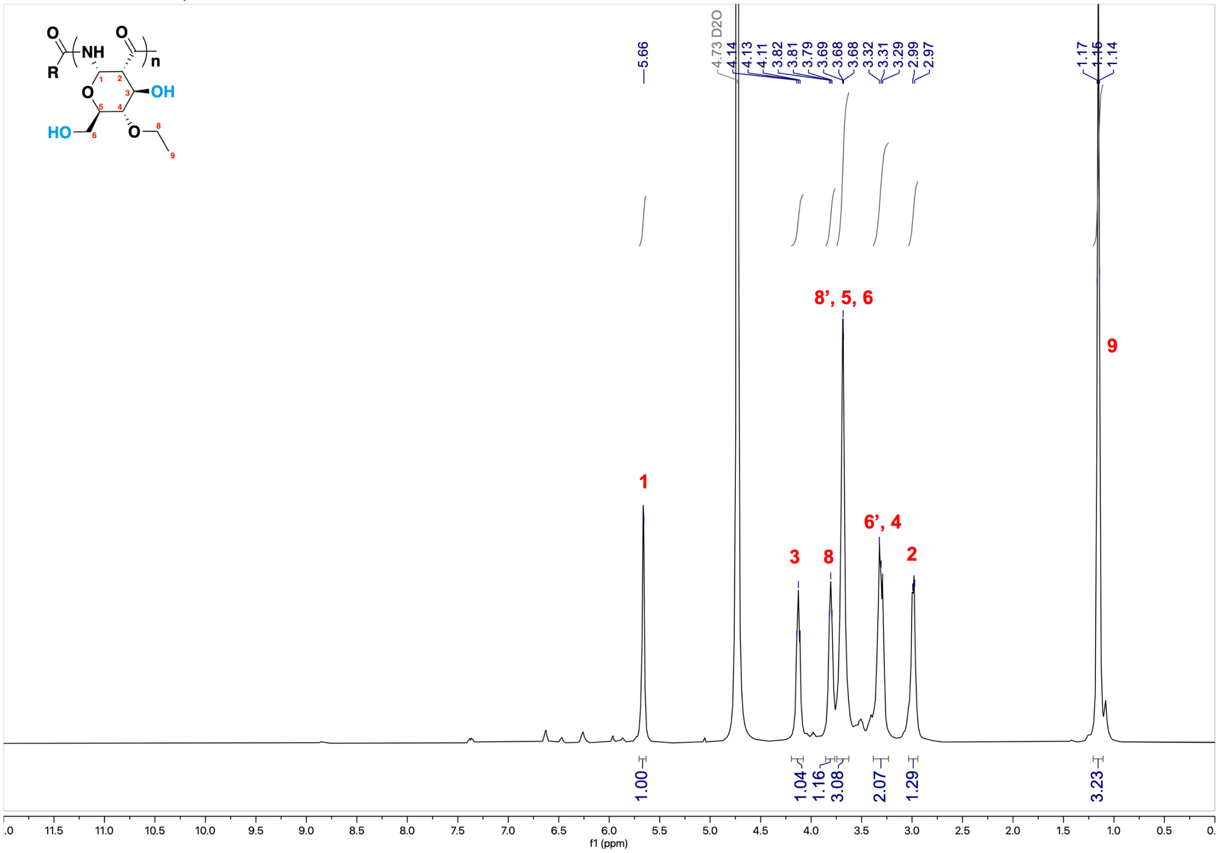


$^{13}$C NMR of **3,6-di-OH-4-*O*-Et-PAS.**

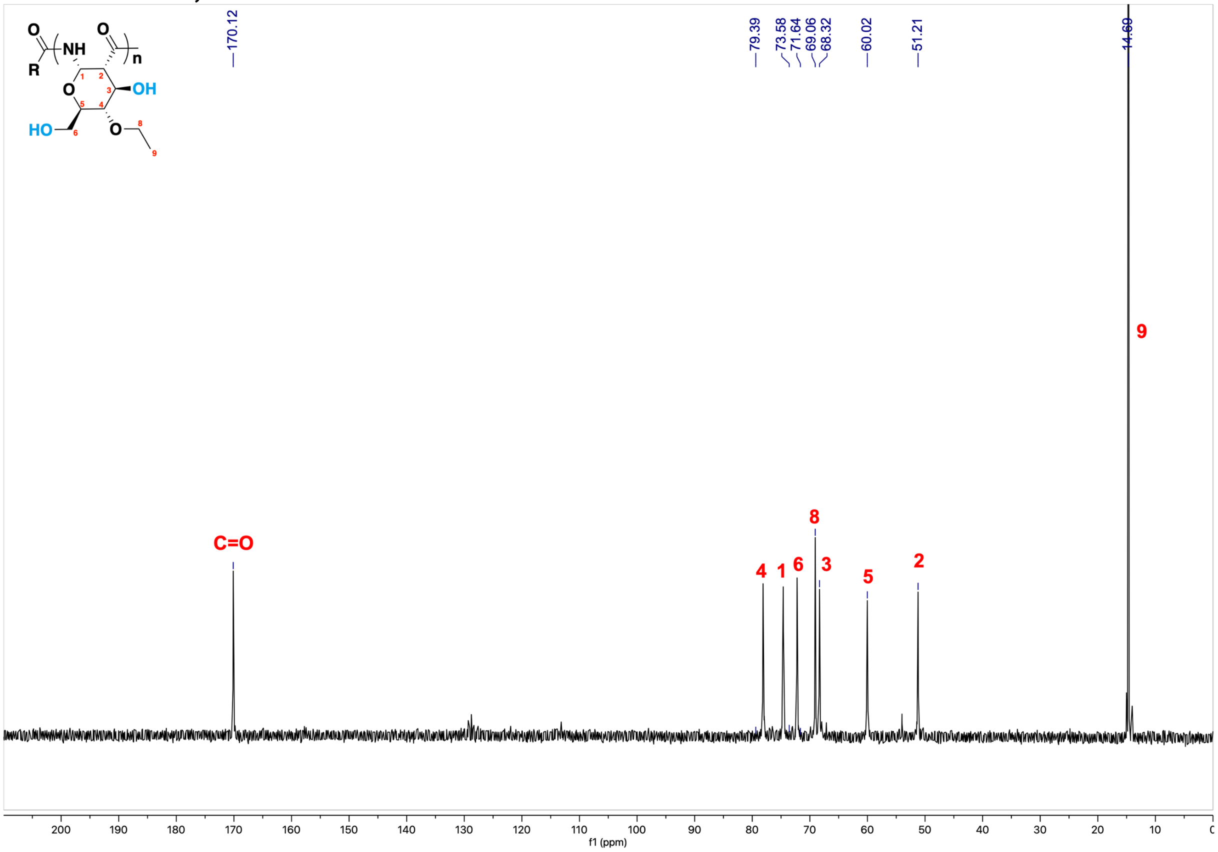

**3,6-di-OH-4-*O*-Et-PAS** - [$^1$H, $^1$H] COSY (2D, 600 MHz, $CDCl_3$)

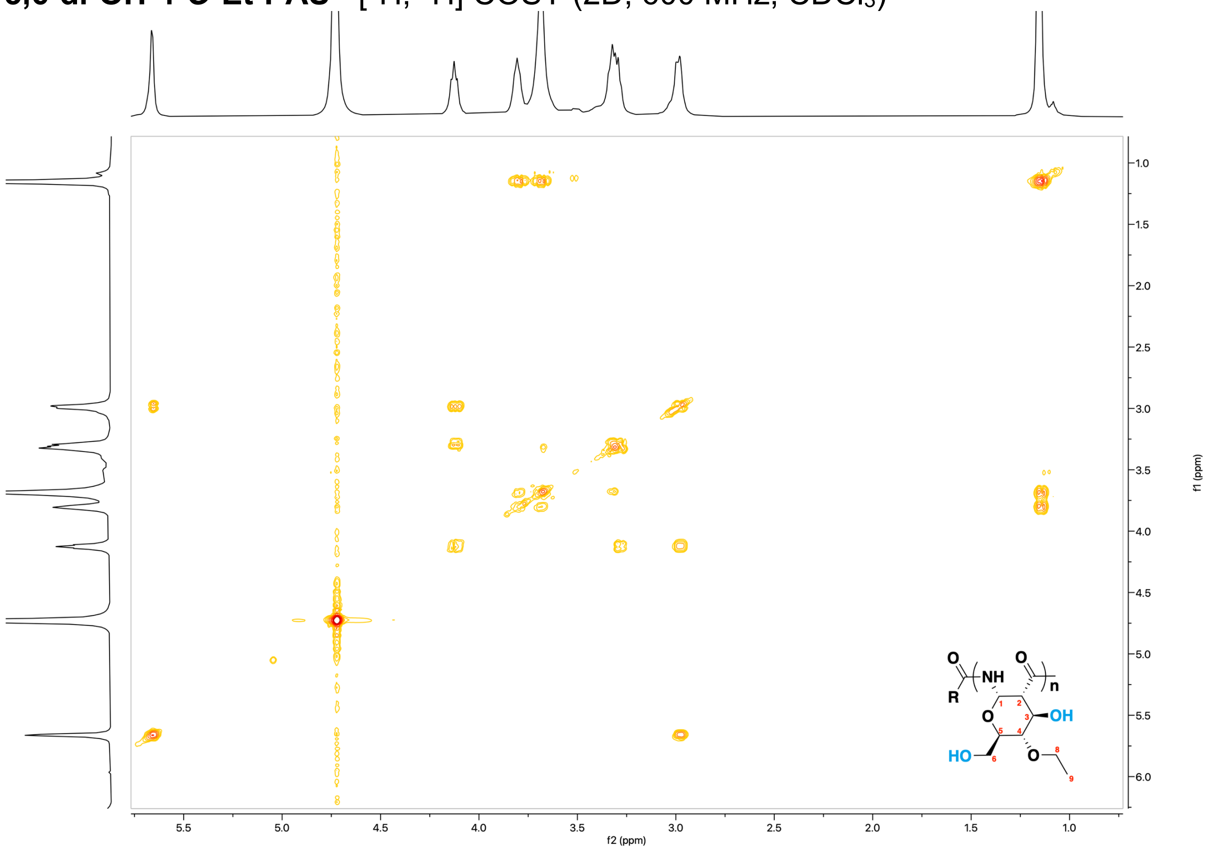

**3,6-di-OH-4-*O*-Et-PAS** - [$^1$H, $^{13}$C] HSQC (2D, 151 MHz, $CDCl_3$)

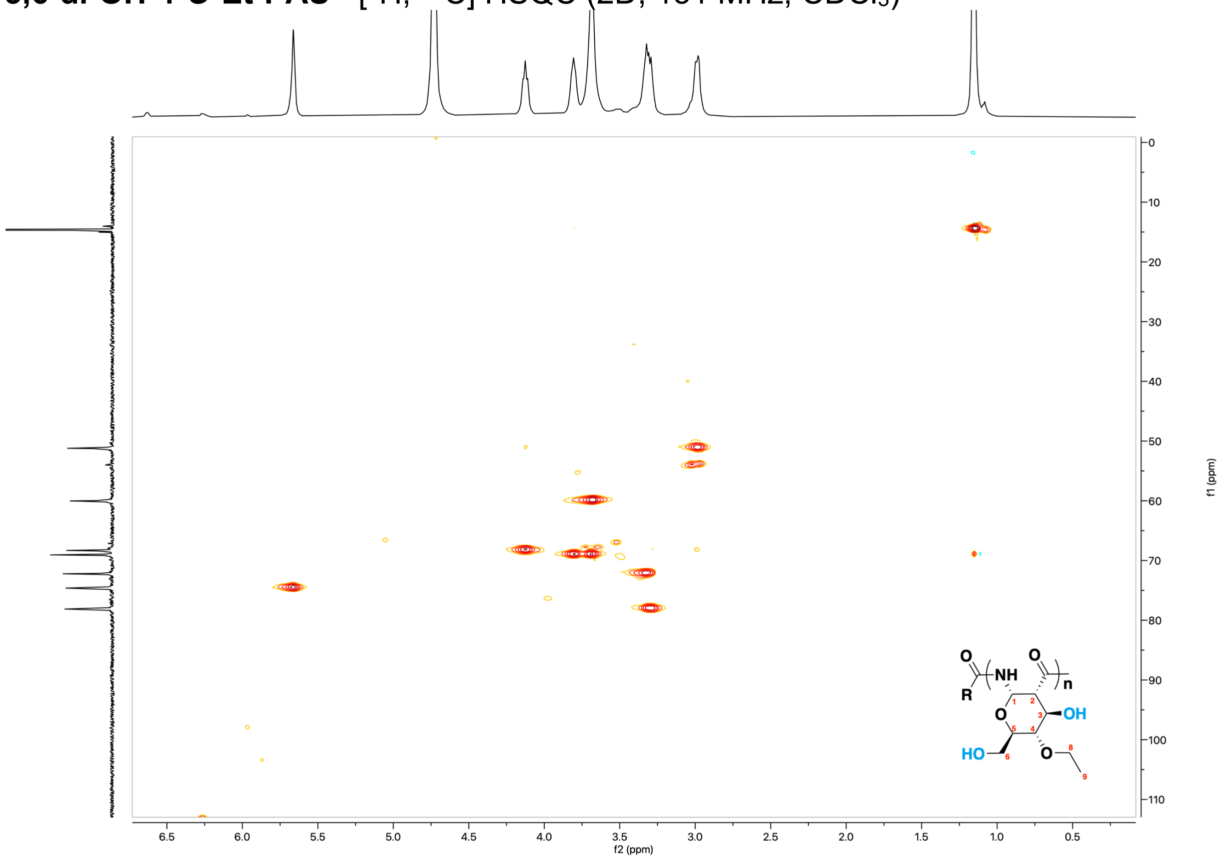

$^{1}$H NMR of **3,6-di-sulfate-4-*O*-Bn-PAS.**

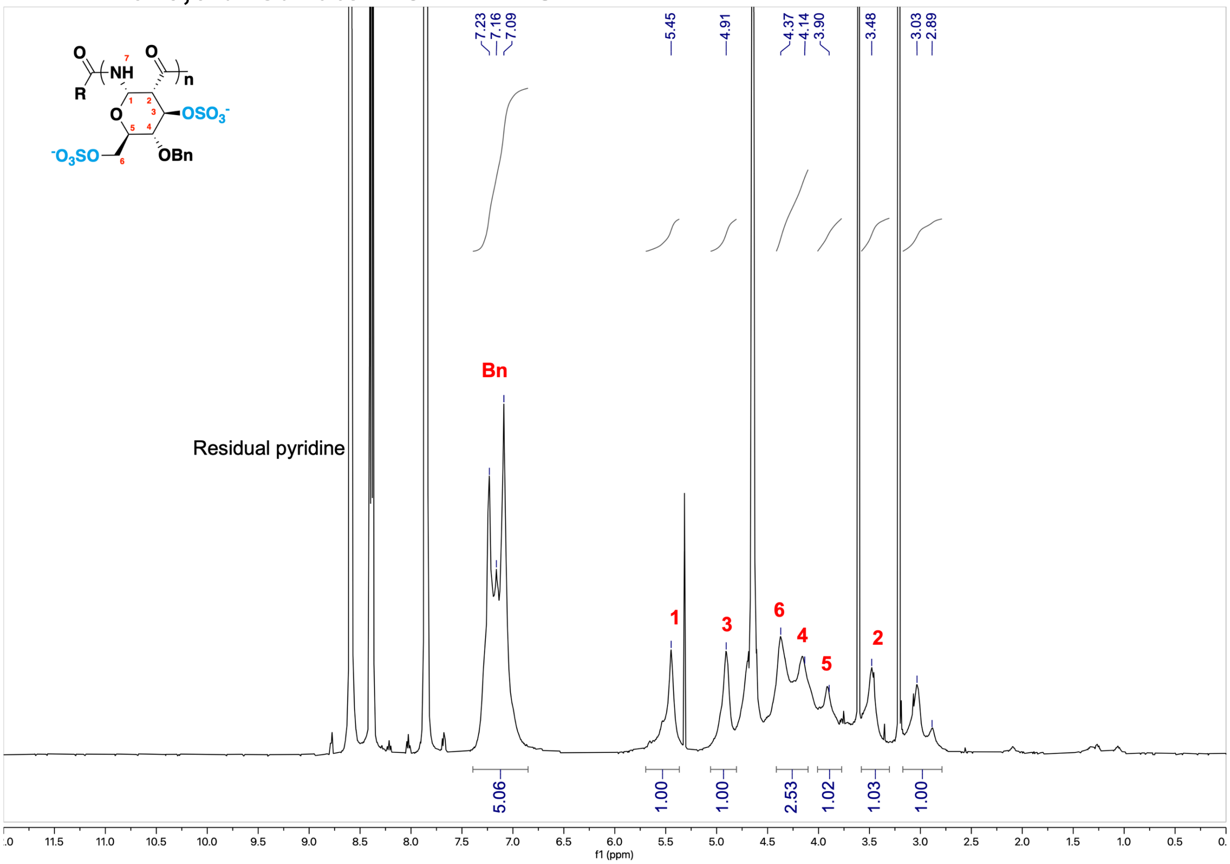


$^{13}$C NMR of **3,6-di-sulfate-4-*O*-Bn-PAS.**

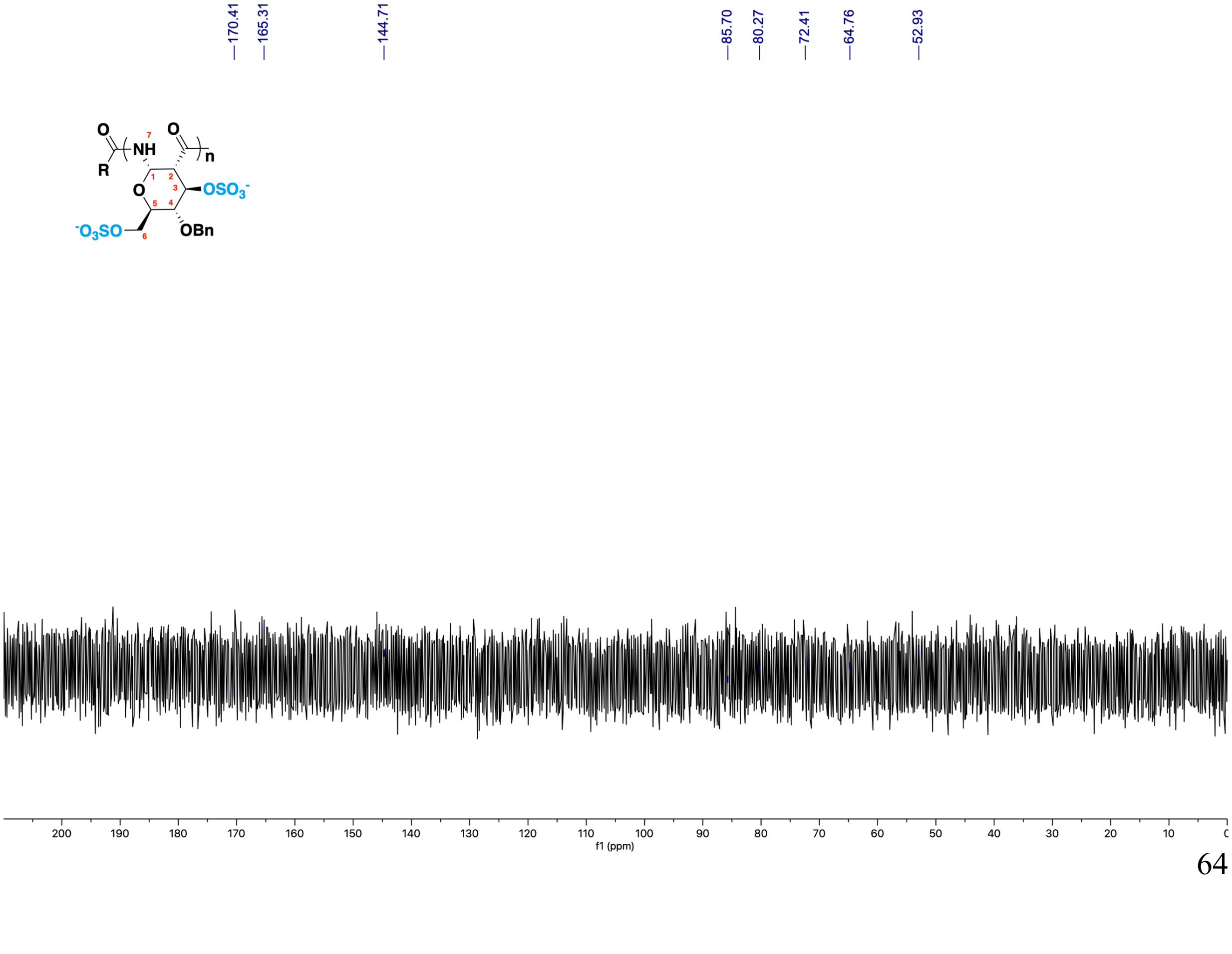

$^{1}$H NMR of **3,6-di-sulfate-4-*O*-Et-PAS** *before* NaOH treatment.

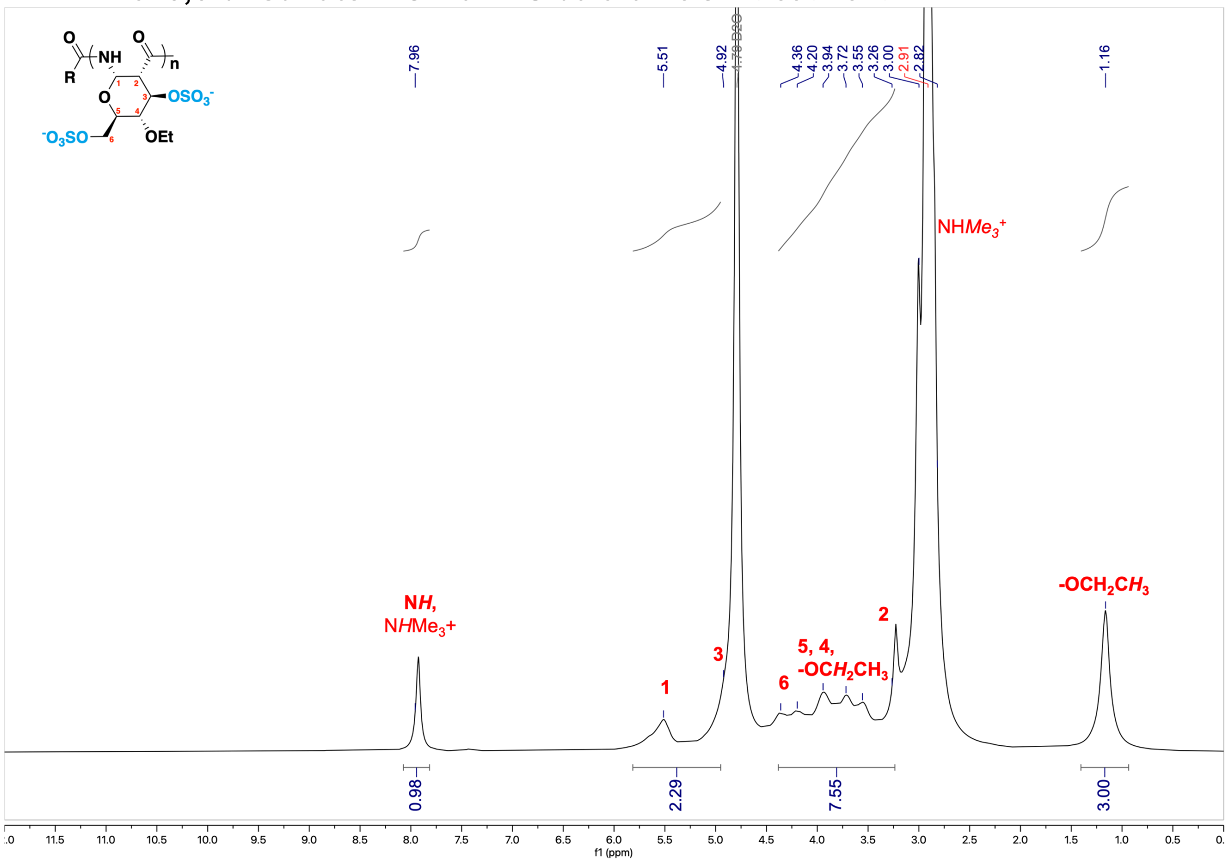


$^{13}$C NMR of **3,6-di-sulfate-4-*O*-Et-PAS** *before* NaOH treatment.

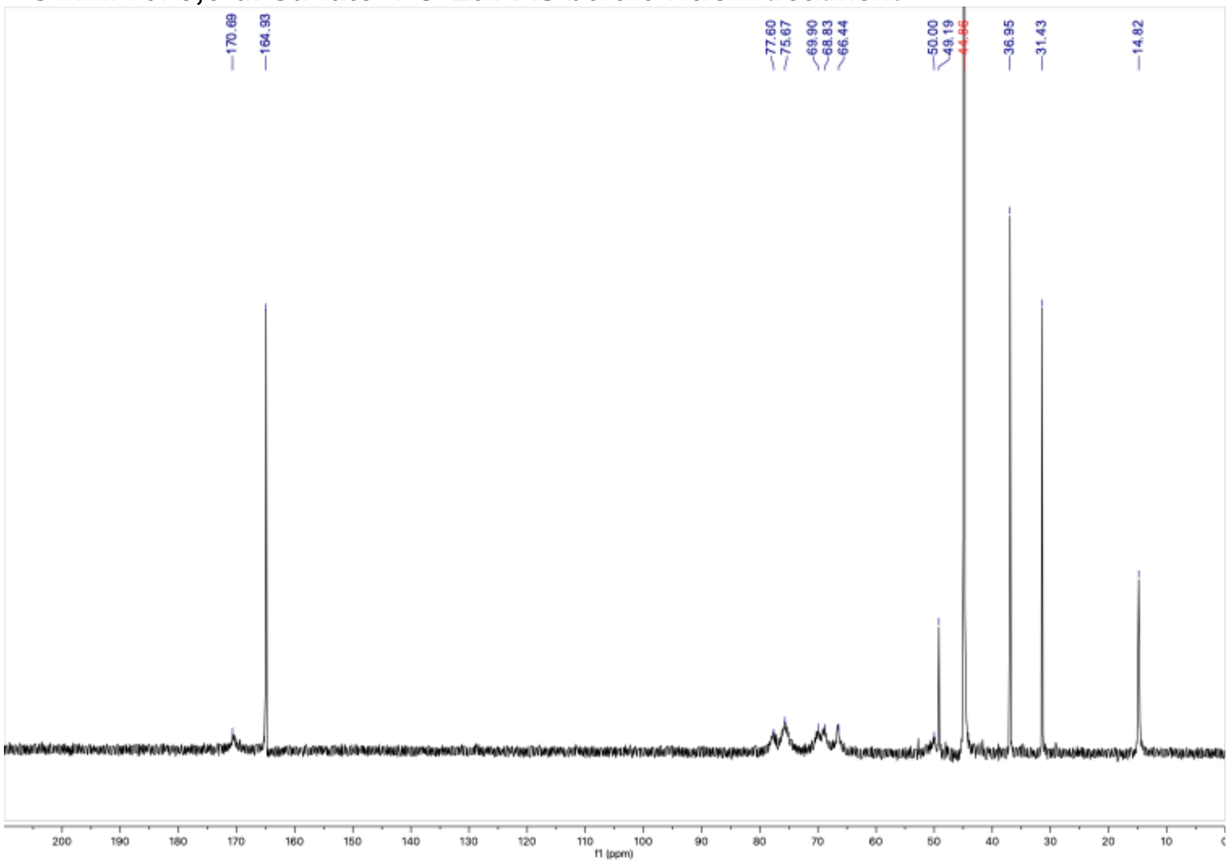

[$^1$H, 1H] COSY of **3,6-di-sulfate-4-*O*-Et-PAS** *before* NaOH treatment.

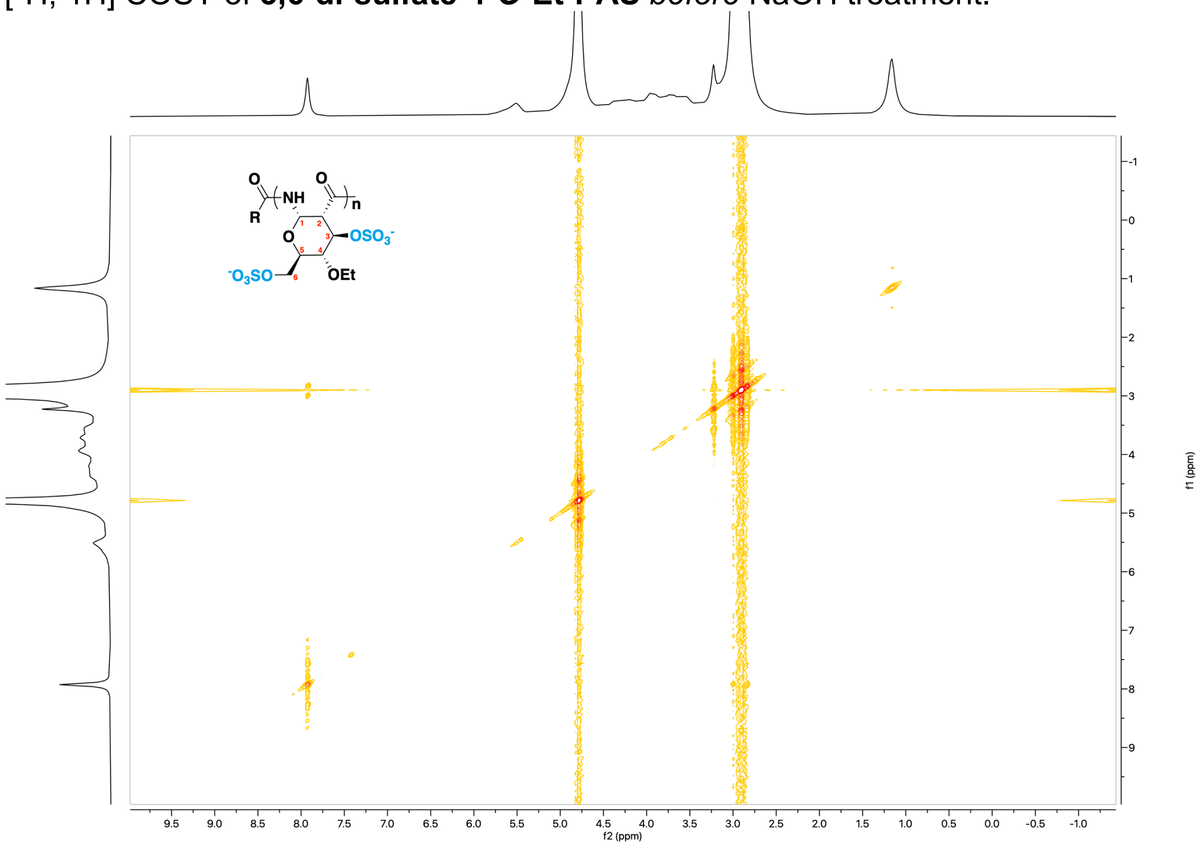

[$^1$H, $^{13}$C] HSQC of **3,6-di-sulfate-4-*O*-Et-PAS** *before* NaOH treatment.

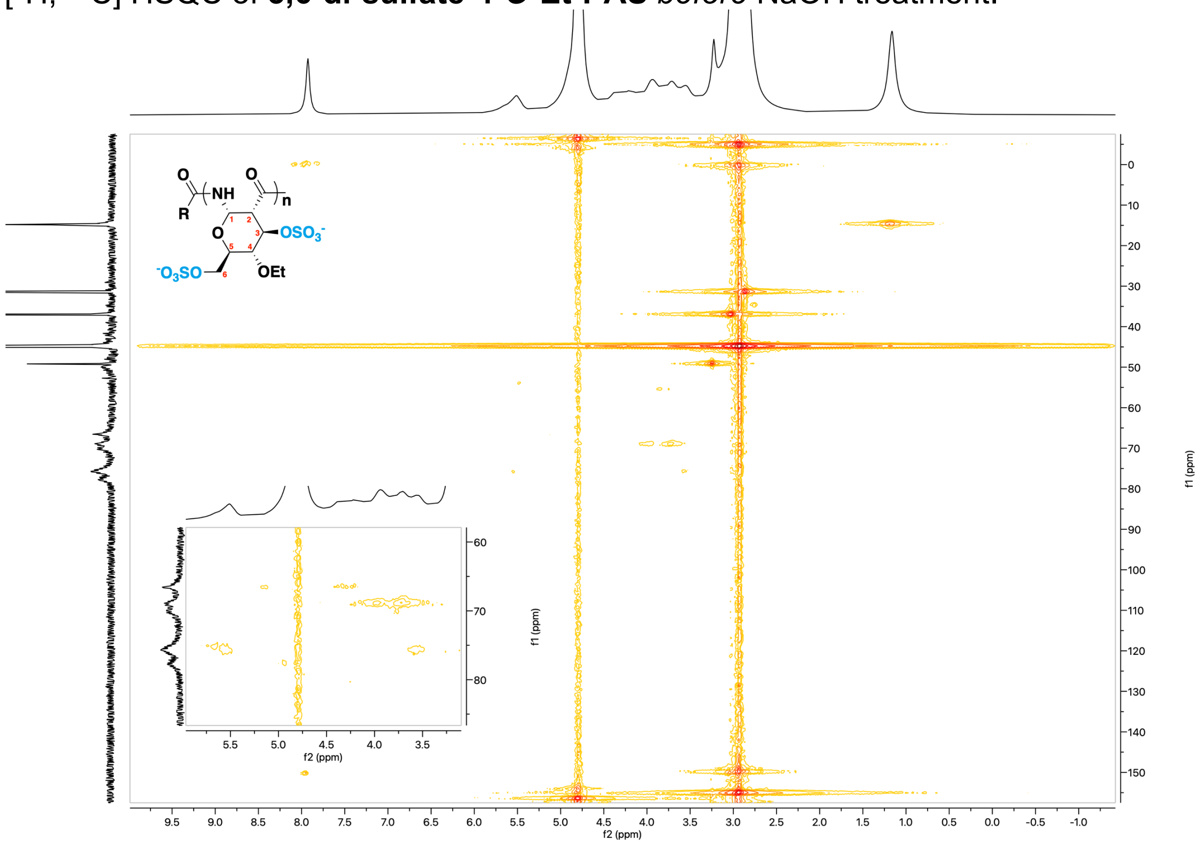

[1]H NMR of **3,6-di-sulfate-4-*O*-Et-PAS** *after* NaOH treatment.

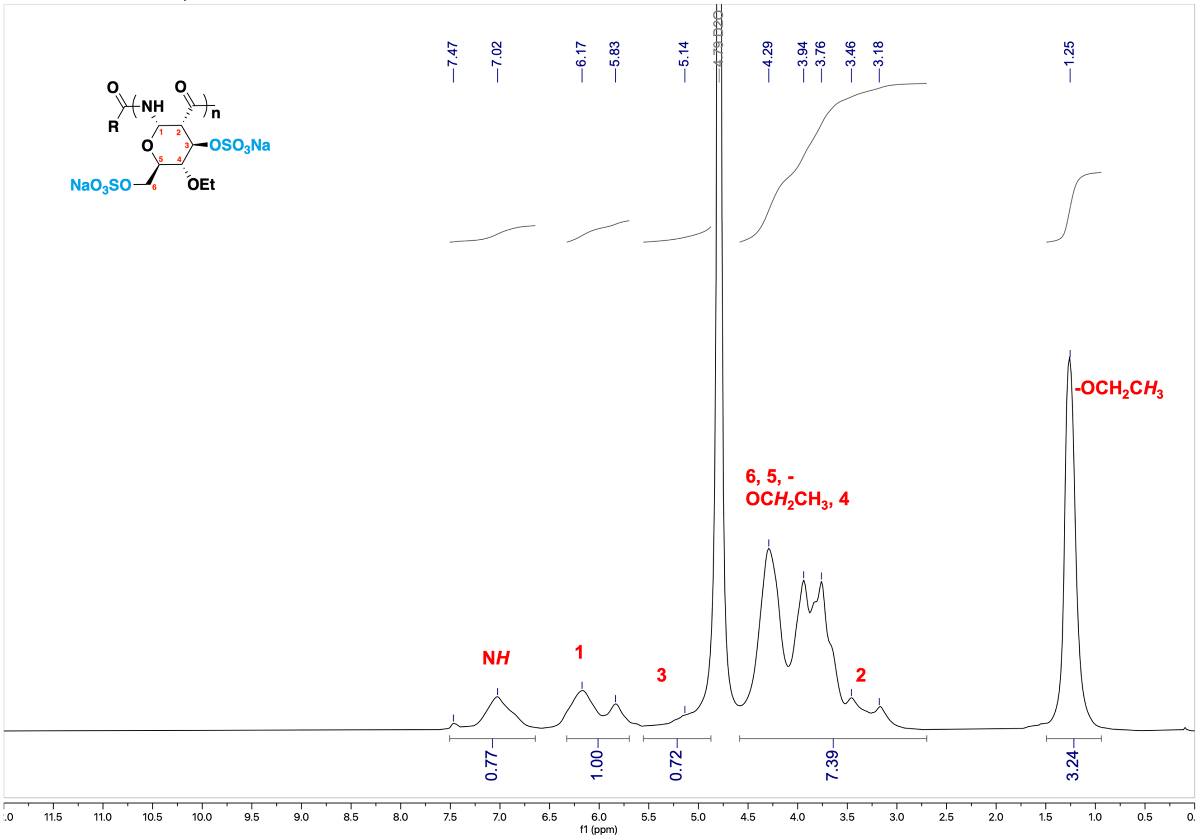


[13]C NMR of **3,6-di-sulfate-4-*O*-Et-PAS** *after* NaOH treatment.

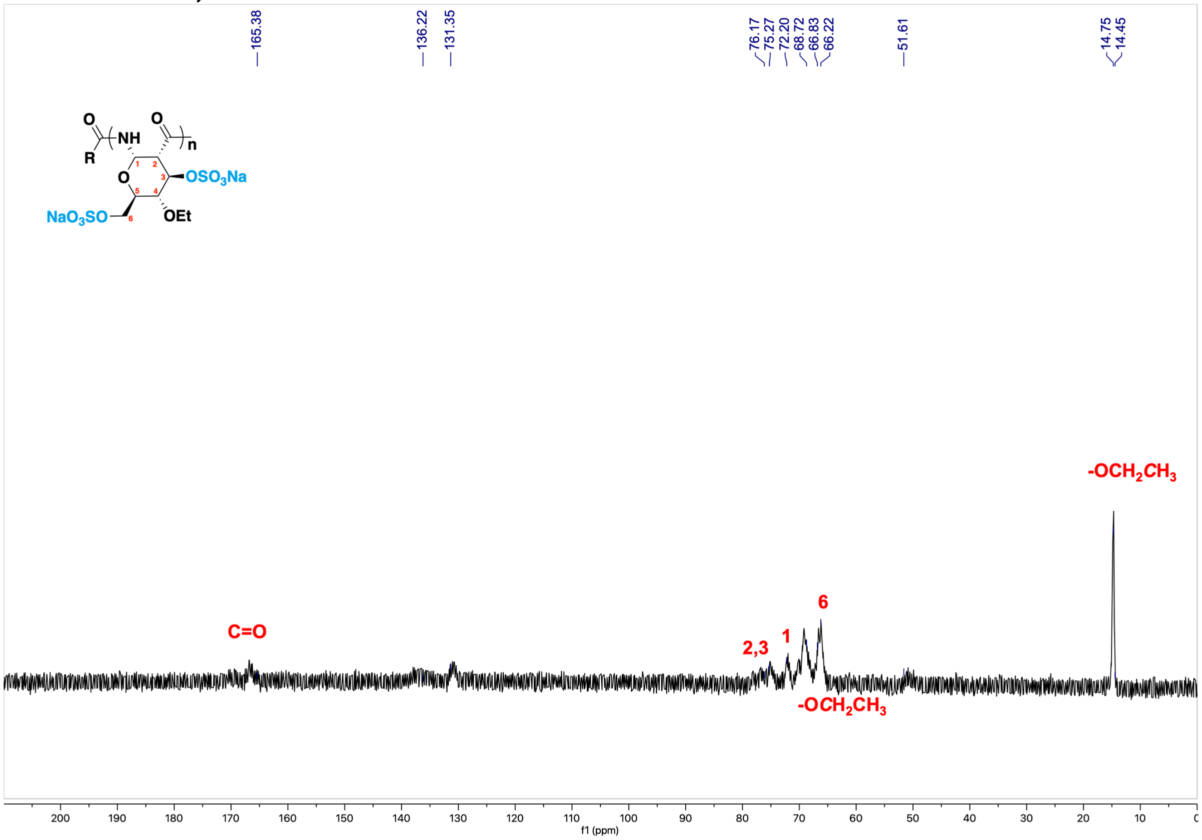

[$^1$H, 1H] COSY of **3,6-di-sulfate-4-*O*-Et-PAS** after NaOH treatment.

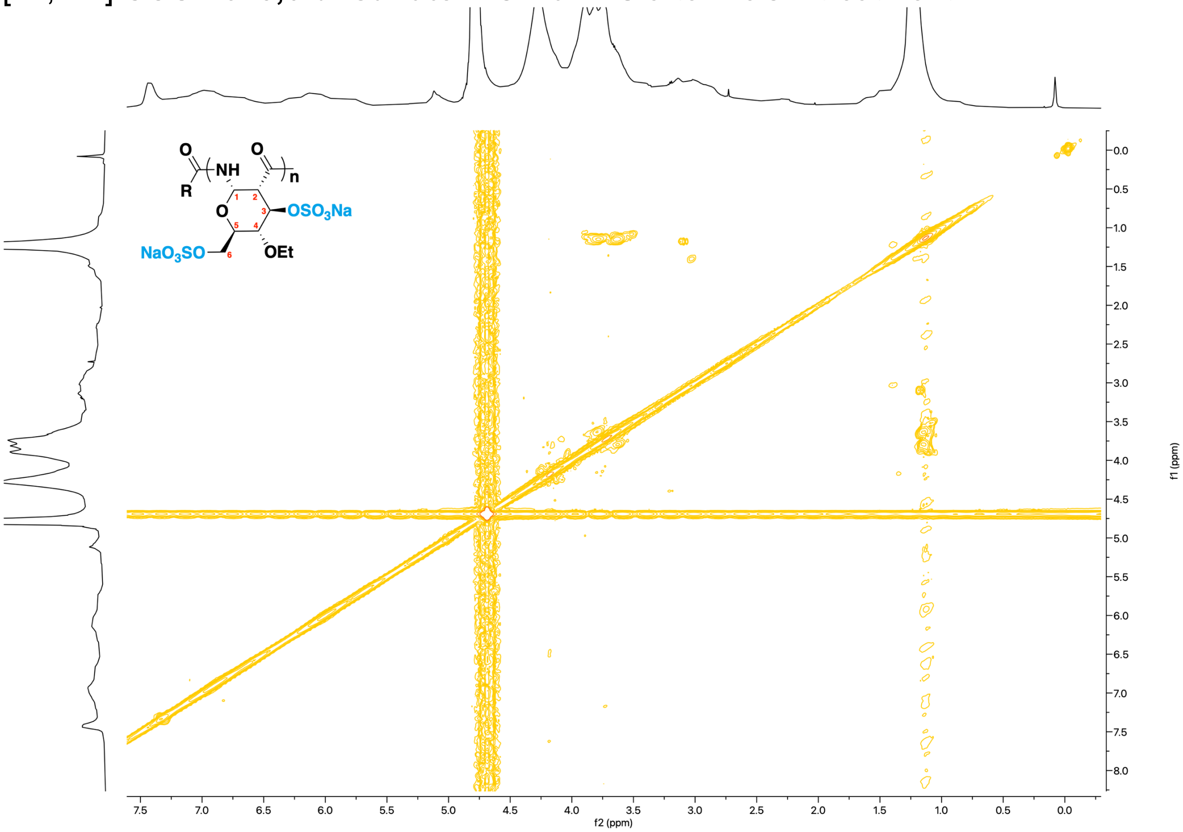


[$^1$H, $^{13}$C] HSQC of **3,6-di-sulfate-4-*O*-Et-PAS** after NaOH treatment.

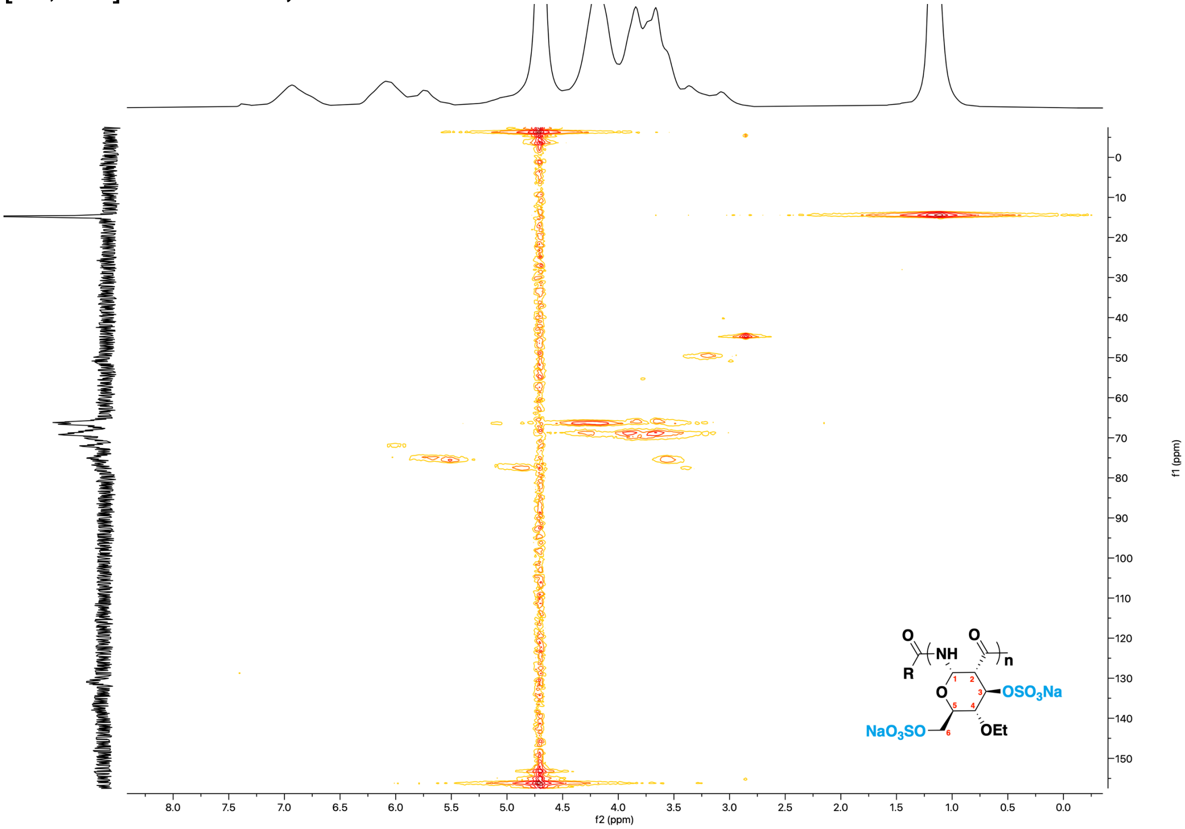
